# The First Complete Zoroastrian-Parsi Mitochondrial Reference Genome and genetic signatures of an endogamous non-smoking population

**DOI:** 10.1101/2020.06.05.124891

**Authors:** Villoo Morawala Patell, Naseer Pasha, Kashyap Krishnasamy, Bharti Mittal, Chellappa Gopalakrishnan, Raja Mugasimangalam, Naveen Sharma, Arati-Khanna Gupta, Perviz Bhote-Patell, Sudha Rao, Renuka Jain, The Avestagenome Project

## Abstract

The present-day Zoroastrian-Parsis have roots in ancient pastoralist migrations from circumpolar regions leading to their settlement on the Eurasian Steppes and later, as Indo-Iranians in the Fertile Crescent. After migrating from the Persian province of Pars to India, the Zoroastrians from Pars (“Parsis”) practiced endogamy, thereby preserving their genetic identity and social practices. The study was undertaken to gain an insight into the genetic consequences of migration on the community, the practice of endogamy, to decipher the phylogenetic relationships with other groups, and elucidate the disease linkages to their individual haplotypes

We generated the *de novo* the Zoroastrian-Parsi Mitochondrial Reference Genome (AGENOME-ZPMS-HV2a-1), which is the first complete mitochondrial reference genome assembled for this group. Phylogenetic analysis of an additional 99 Parsi mitochondrial genome sequences showed the presence of HV, U, T, A and F (belonging to the macrohaplogroup N) and Z and other M descendents of the macrohaplogroup M (M5, M39, M33, M44’52, M24, M3, M30, M2, M4’30, M2, M35 and M27) and a largely Persian origin for the Parsi community. We assembled individual reference genomes for each major haplogroup and the Zoroastrian-Parsi Mitochondrial Consensus Genome (AGENOME-ZPMCG V1.0), which is the first consensus genome assembled for this group. We report the existence of 420 mitochondrial genetic variants, including 12 unique variants, in the 100 Zoroastrian-Parsi mitochondrial genome sequences. Disease association mapping showed 217 unique variants linked to longevity and 41 longevity-associated disease phenotypes across the majority of haplogroups.

Analysis of the coding genes, tRNA genes, and the D-loop region revealed haplogroup-specific disease associations for Parkinson’s disease, Alzheimer’s disease, cancers, and rare diseases. No known mutations linked to lung cancer were found in our study. Mutational signatures linked to tobacco carcinogens, specifically, the C>A and G>T transitions, were observed at extremely low frequencies in the Parsi cohort, suggestive of an association between the cultural norm prohibiting smoking and its reflection in the genetic signatures. In sum, the Parsi mitochondrial genome provides an exceptional resource for determining details of their migration and uncovering novel genetic signatures for wellness and disease.

## Introduction

### The Travelogue of the Zoroastrian-Parsi Mitochondrion

Human mitochondrial DNA (mtDNA) is a double-stranded, circular (16,569 kb) genome of bacterial origin^1,2^, primarily encoding 22 tRNAs and 2 rRNAs and the genes encoding subunits of the energy-generating oxidative phosphorylation and electron transport chain (ETC) pathway^3,4^. Analysis of the variability of mtDNA is commonly used to reconstruct the history of populations, especially with respect to maternal inheritance.

The accumulation over time of maternally inherited mitochondrial variants creates haplotypes^4^ characteristic of different mtDNA lineages and can be used to follow populations through history and trace their migrations. Such an approach has also provided insights into the origins and disease etiologies associated with endogamous communities, such as the Icelandic population^5^, island communities of Andaman and Nicobar^6^ and Polynesia^7^.

Until the fall of the Zoroastrian Persian Empire in the seventh century AD, the Zoroastrian-Parsis resided in what is now the province of Pars in present-day Iran^8,9^. To escape the persecution that ensued, the Zoroastrians from Pars^10,11^ (referred to here as the “Parsis”) migrated to India in the 8^th^ century AD. Because they practiced endogamy^12,13^ among their Indian neighbors, Parsi genetic identity was retained to a large extent as well as certain social practices. As fire was considered sacred in the Zoroastrian religion^14,15^, strict social ostracism has long been maintained against smokers within the Parsi community. Today, the Parsis, are a small community of <52,000 in India (2011 Census, Govt of India). We present the genetic data for the conserved Parsi mitochondrion, which has survived largely intact for over 1300 years.

In this study, our aim was two fold: 1) to determine the consequences for the Parsi mitochondrial genome of the historic Parsi migration from Persia to India and their subsequent practice of endogamy and 2) to discover any linkages between the mtDNA variants observed in the Parsis and their predispositions to various diseases. To address these questions, we generated *de novo* the Zoroastrian Parsi Mitochondrial Genome (AGENOME-ZPMS-HV2a-1; Genbank ID, MT506314), which is complete and the first of its kind, and used it as our starting point to determine the mitochondrial haplogroup-specific reference genomes for 100 Parsi individuals. We also assembled the Zoroastrian Parsi Mitochondrial Consensus Genome (AGENOME-ZPMCG V1.0; Genbank ID MT506339), which is also the first of its kind.

Our phylogenetic analysis confirmed that the present-day Parsis are closely related to Persians and, like most endogamous communities, have comparatively low genetic diversity and are predisposed to several inherited genetic disorders^16,17^. Interestingly, the Parsis also possess longevity as a trait and are a long-lived community^18^, with lower incidences of lung cancer^19^. Overall, the Parsi community is a unique genetic resource for understanding the linkage between mtDNA variation and disease.

## Results

### Assembly of the first complete Zoroastrian-Parsi mitochondrial sequence, AGENOME-ZPMS-HV2a-1

The first complete *de novo* non-smoking Zoroastrian-Parsi mitochondrial sequence, AGENOME-ZPMS-HV2a-1 (Genbank ID, MT506314), was assembled from a healthy Parsi female by combining the sequence data generated from two next-generation sequencing (NGS) platforms using the protocol outlined in Materials and Methods. Our approach combines the sequencing depth and accuracy of short-read technology (Illumina) with the coverage of long-read technology (Nanopore). QC parameters for mitochondrial reads, mitochondrial coverage, and the extent of coverage were found to be optimal (**Supplementary Figure 1**). The hybrid Zoroastrian-Parsi mitochondrial genome was assembled as a single contig of 16.6 kb (with 99.82% sequence identity), resulting in the first *de novo* Zoroastrian-Parsi mitochondrialsequence, with 99.84% sequence identity with the revised Cambridge Reference Sequence (rCRS^21^).

### Identification of 28 unique variants in AGENOME-ZPMS-HV2a-1

A total of 28 significant variants were identified by BLAST alignment between the Parsi mitochondrial hybrid assembly and the rCRS^20^ (**Figure 1A, B**). To confirm their authenticity, we selected a total of 7 identified variants from the D-loop region and one SNP from the *COI* gene (m.C7028T) and subjected them to Sanger sequencing. All 8 predicted variants were confirmed (**Supplementary Figure 2**).

**Figure 1A:**
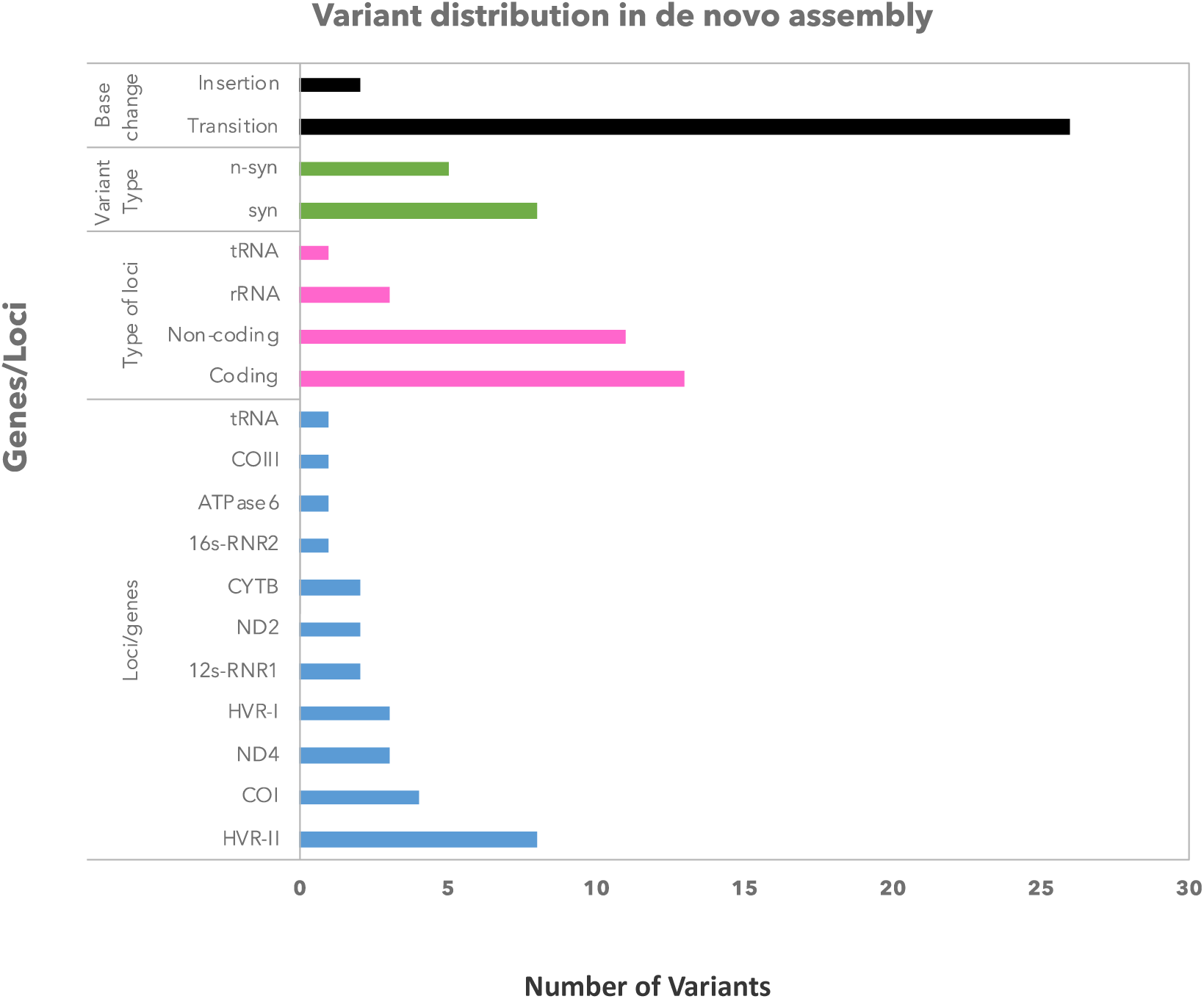
Identification of 28 variants in the de novo Parsi mitochondrial genome, AGENOME-ZPMS-HV2a-1

**Figure 1B:**
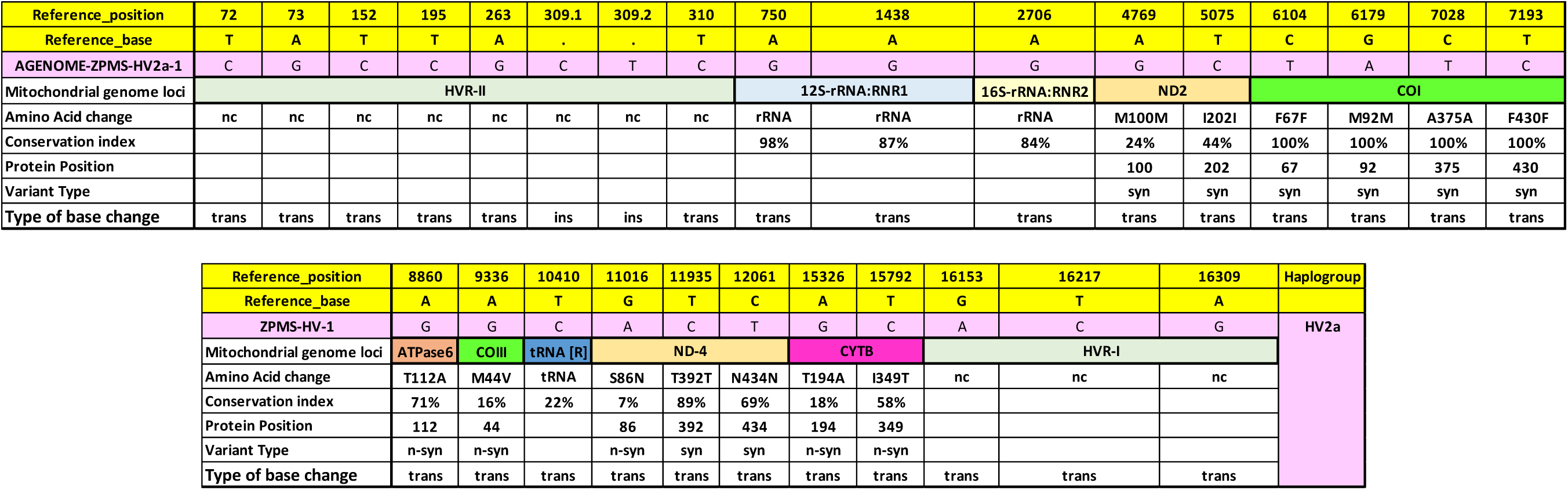
Annotation of 28 variants in the AGENOME-ZPMS-HV2a-1

**Figure 1.** Characterization of 28 variants identified in the *de novo* Parsi mitochondrial reference genome (AGENOME-ZPMS-HV2a-1). **A**, Classification and distribution of the variants. B, Annotation of the variants in relation to the revised Cambridge Reference Sequence (rCRS).

The majority of the variants identified in the AGENOME-ZPMS-HV2a-1 (n=11) were found in the hypervariable regions (HVRI and HVRII) of the D-loop in comparison to other individual regions in the mitochondrial genome sequence. Of the remaining 17 variants, eight were found to represent synonymous variants, while four were in genes for 12S rRNA, 16S rRNA (n=3), and tRNA (n=1) (**Figure 1A**). The remaining 5 nonsynonymous variants were located (one each) within the genes for *ATPase6* (m8860G>A), *COIII* (m.9336 A>G), and *ND4* (m.11016 G>A), while two were located in the *CytB* gene (m15326 A>G and m15792 T>C) (**Figure 1B**). Except for the *ATPase6* gene variant, whose occurrence is associated mitochondrial degenerative diseases like Alzheimers, Lebers Hereditary Optic Neuropathy (LHON) and idiopathic cardiomyopathy^21^, no other disease associations were found in the published literature.

Given that the Parsis are known to have originated in Persia (present day Iran) and have practiced endogamy since their arrival on the Indian subcontinent, we wished to determine the mitochondrial haplogroup associated with the AGENOME-ZPMS-HV2a-1. We therefore compared the variants associated with this sequence to standard haplogroups obtained from MITOMAP and determined the haplogroup to be HV2a (**Figure 1B**). This haplogroup is known to have originated in Iran^25^, suggesting Persian ancestry for this Parsi individual.

### Seven major haplogroups identified in the 100 Parsi individuals

Keeping in mind the endogamous customs of the Indian Parsis and to understand the extent of the diversity of the mitochondrial haplogroups in this population, we analyzed mitochondrial genomes from 100 consenting Parsi individuals. Our study had an equal representation of both genders, and 60% of the subjects were of age 30–59 (mean age 50±1.6, **Figure 2A**). Complete analysis of the variants in the 100 Parsi samples identified a total of 420 distinct variants (**Figure 2B**, **Appendix 1**). QC analysis of the 100 mitochondrial genomes sequenced determined them to be optimal (PHRED>30, **Supplementary Figure 3**). Variant distribution in the coding region normalized to gene length showed that the *ND6* gene had the greatest number of variants (**Supplementary Figure 4**).

**Figure 2A:**
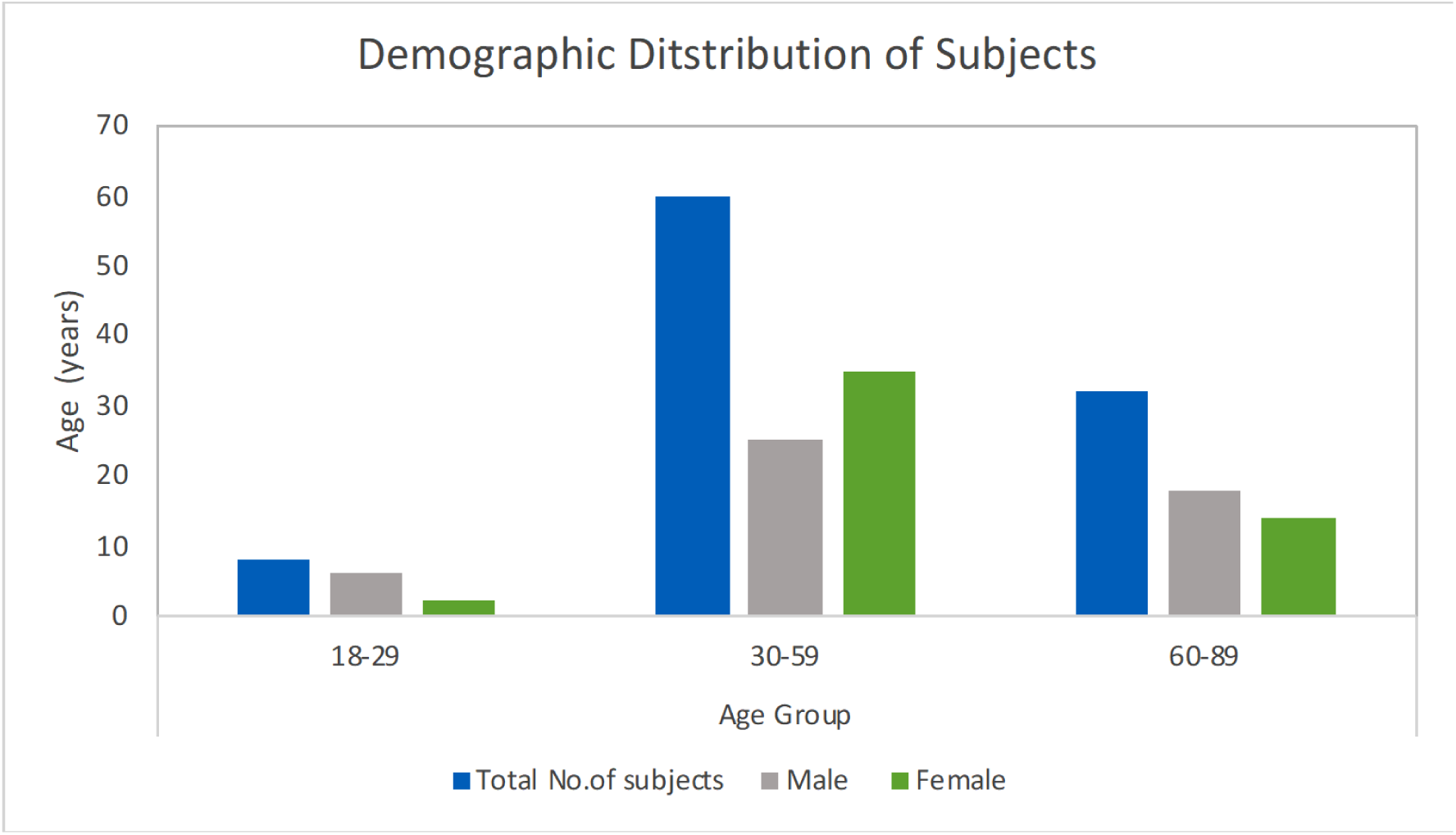
Representation of Males and Females in the 100 Zoroastrian-Parsi whole mitogenome study

**Figure 2B:**
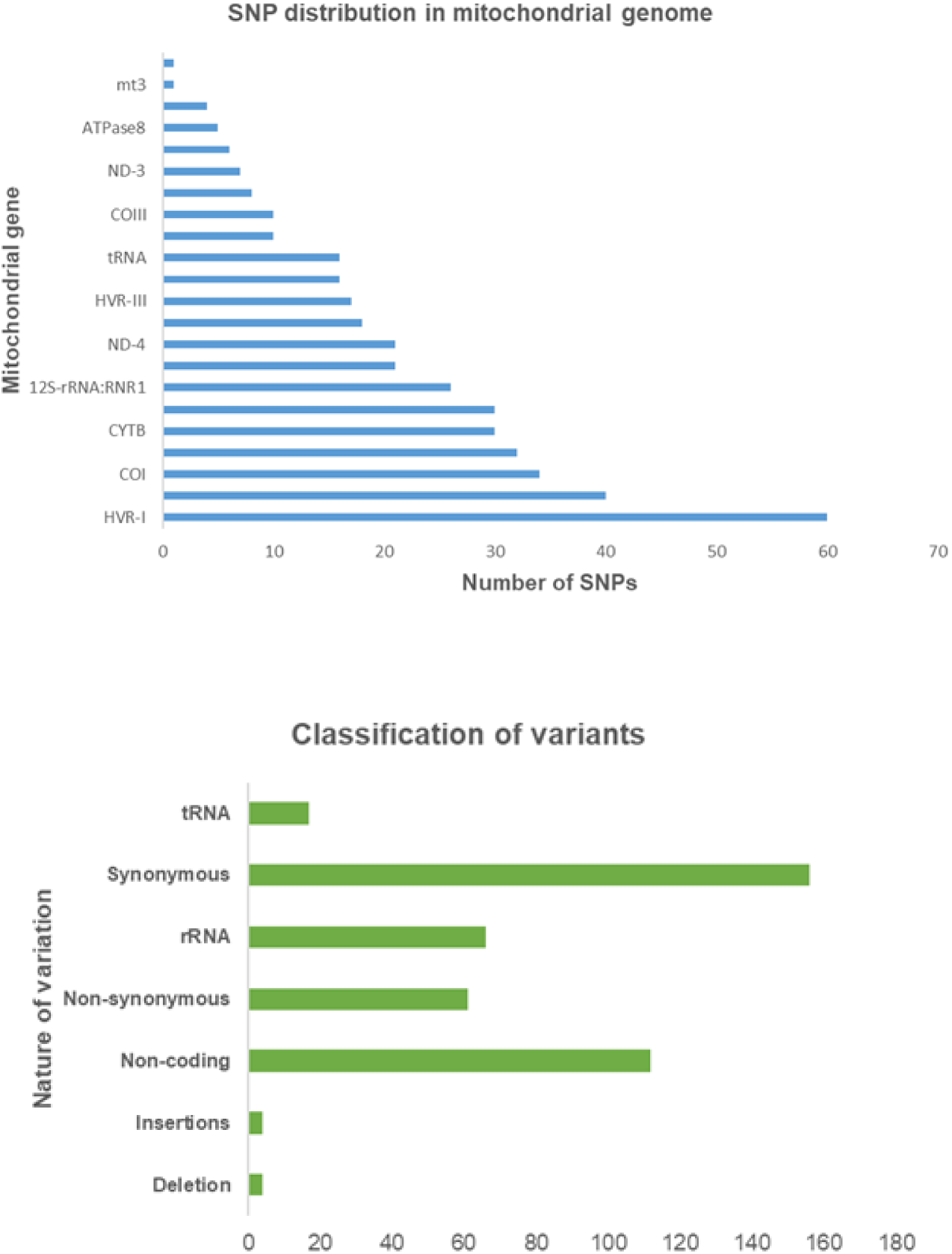
Distribution of 420 variants across gene loci in the 100 Zoroastrian-Parsi whole mitogenomes

**Figure 2:** Characterization of the 100 study participants and the variants identified in their mitochondrial genomes. **A,** Demographic distribution of the 100 Zoroastrian-Parsi subjects in this study. **B,** (upper) Distribution of the 420 SNPs identified in the genes of the 100 mitochondrial genomes; (lower) classification of the 420 variants identified in the 100 mitochondrial genomes.

The 100 Parsi mitochondrial genomes were subjected to haplogroup analysis using a haplogroup-specific variant assignment matrix from MITOMAP (**Appendix 4**). The variant-based haplogroup assignments classified the genomes as HV, U, T, A and F (belonging to the macrohaplogroup N) and Z and other M descendants of the macrohaplogroup M (M5, M39, M33, M44’52, M24, M3, M30, M2, M4’30, M2, M35 and M27) (HV, U, T, M, A, F, and Z), and 25 sub-haplogroups were identified within these principal haplogroups (**Table 1**). Additional analysis of haplogroup classification indicated alternate haplogroup calls for A2v (Alternate call: H; A2v:7 variants, H:7 variants), M24a (Alternate call:M37; M24a:25 variants, M37:25 variants), M27b (Alternate call:M30b; M27b:25 variants, M30b:25 variants), T2b (Alternate call: R30b; T2b:14 variants, R30b:13 variants), Z1a (Alternate call:M37a; Z1a:26 variants, M37a:25 variants). The variant count across all sub-haplogroups was in the range 14–64 (**Supplementary Figure 5A**). Analysis of the sub-haplogroups demonstrated that HV2a was the single largest sub-haplogroup within the Parsi population (n=14, n=9 females, n=5 males, **Supplementary Figure 5B**), including the AGENOME-ZPMS-HV2a-1 subject.

**Table 1:**
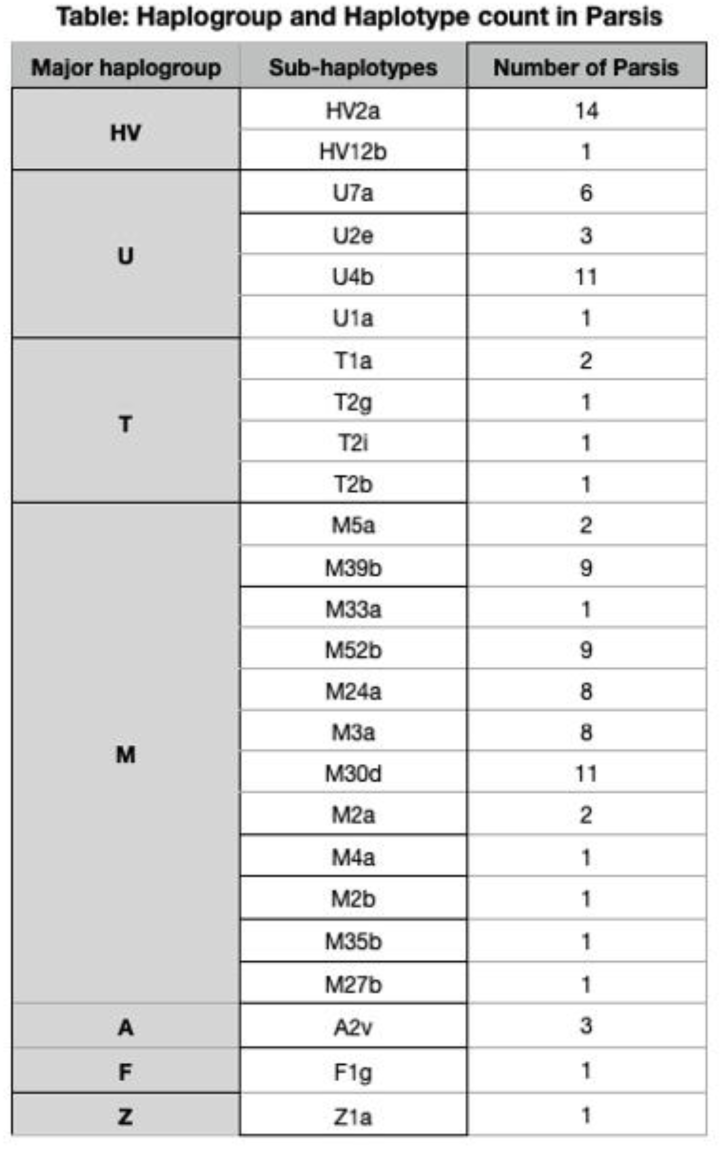
Identification of 25 sub-haplogroups in the 100 Zoroastrian-Parsi study group. Distribution of the 100 Parsi subjects across 7 major haplogroups and 25 sub-haplogroups.

All subjects of sub-haplogroup HV2a (n=14) contained the 27/28 variants observed in the AGENOME-ZPMS-HV2a-1 sequence. In total, the HV2a sub-haplogroup had 38 variants, with the highest number in the HVRII region (n=8). Coding region mutations constituted 20/38 variants, with an equal distribution between synonymous (n=10) and nonsynonymous (n=10) substitutions observed for this sub-haplogroup. Among the coding regions, the greatest number of variants was found in the gene encoding *COI* (n=6, **Supplementary Figure 6A**). We found a variant in the gene encoding tRNA[R] at m.10410 T>C (n=14 subjects), but no mutations were observed in the D-loop region for the entire group under analysis.

Further analysis of the other sub-haplogroups revealed that the majority of the variants in the noncoding region occurred in HVRII and HVRI, while in the gene-coding regions, the majority of variants occurred in the *CYTB* gene, followed by variants in the *ND5, ND2*, 12S RNR1, and 16S RNR2 genes (**Supplementary Figure 6A–E**).

### Comparative phylogenetic analysis of the Parsi mitochondrial genomes

A comparative analysis of 100 Parsi mitochondrial genomes with 352 Iranian^22^ and 100 random Indian mitochondrial genome sequences^23–25^ was undertaken. The rationale for selection of the Iranian and Indian populations for comparative analysis was centered around their shared ancestral migration history^26,27^.

We compared the haplogroups identified in the Parsi population with those in the Iranian mitogenome dataset. The Persians (n=180) and the Qashqais (n=112) were the most frequently represented in the Iranian population in the 352 Iranian mitogenome study^22^ when compared with the Iranian population haplogroups, we found that a) all Parsi haplogroups (HV, U, T, A, F, and Z) and lineages of the macrohaplogroup M observed in the Parsis were also seen in the Iranian population and b) there was a marked lack of haplogroup diversity in the Parsi datatset (25 sub-haplogroups) compared to the Persians (125 sub-haplogroups) and Qashqais (77 sub-haplogroups) (**Figure 3A, B, Appendix 7**). The reason for the lack of haplotype diversity may lie in the practice of endogamy, which has been strictly followed by the Parsi community for centuries. Contemporary Iranians belong to a broader range of haplogroups, perhaps due to admixture events following political upheavals in the region^27^.

**Figure 3.**
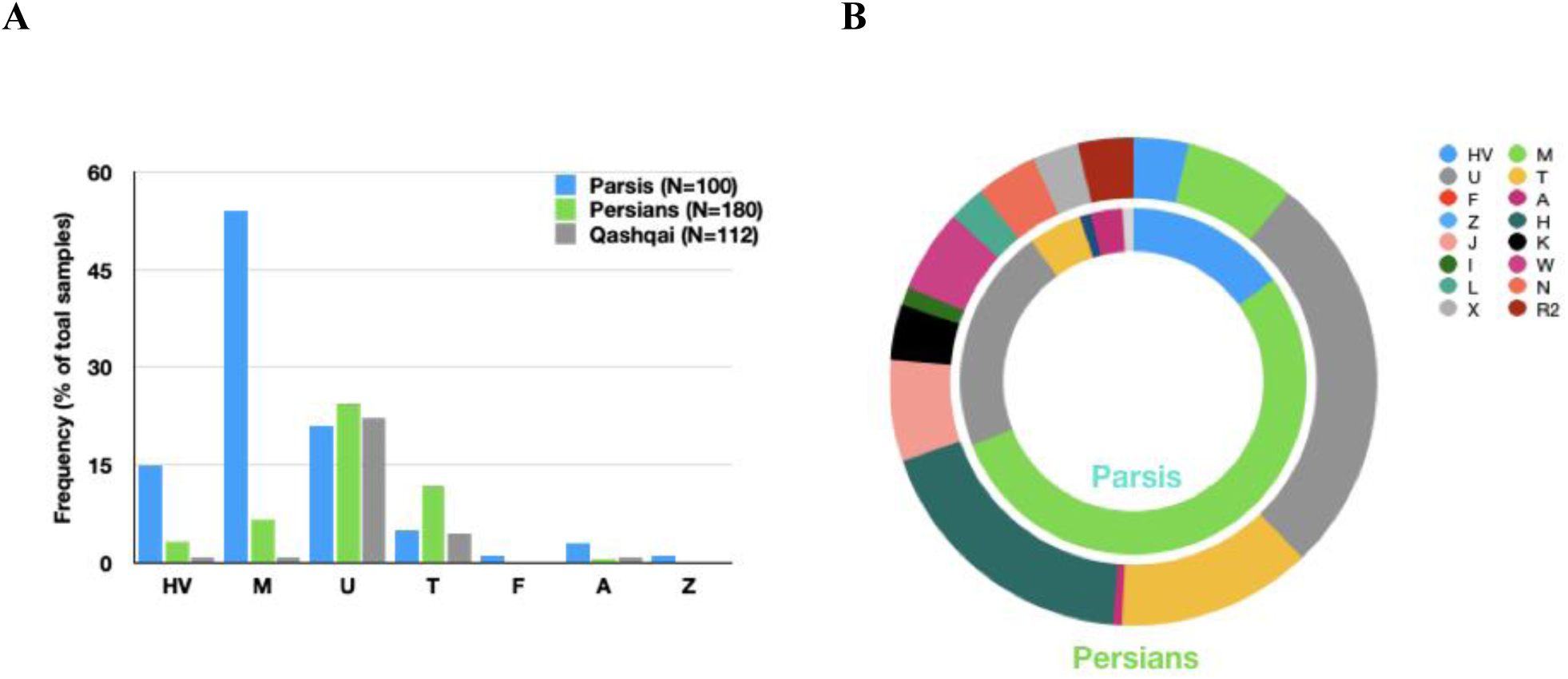
Lack of haplogroup diversity in the Parsi cohort suggesting endogamy. A lack of haplogroup diversity in the Parsi cohort is consistent with endogamy. **A,** Distribution of the seven major haplogroups identified in the Parsi cohort for Parsis, Persians, and Qashqais. **B,** Distribution of all the major haplogroups identified in either the Parsi or Persian cohorts.

Our analysis revealed that the Parsis predominantly cluster with populations from Iran (Persians and people of Persian descent, **Figure 4A, E**). For example, the most common HV sub-haplogroup (HV2a, n=14) clustered with Persians (neighbour-joining tree weight >72%, **Figure 4A and Supplementary Table 3**), while the single Parsi in the HV12b sub-haplotype (n=1) clustered with with other Iranian ethnic groups in the dataset of 352 Iranian mitogenomes, including the Khorasanis and Mazandaranis, in addition to the Qashqais and Persians (**Supplementary Table 3**). The Parsis in the macro-haplogroups U, T, A, F, and Z also cluster with Persians, while there were secondary associations with Kurds, Turkmen, Mazandaranis, Armenians, Azeris, and Khorasanis (**Figure 4B, C**), all of whom claim descent from Mesopotamia and the older Persian empire^22^

**Figure 4A, B:**
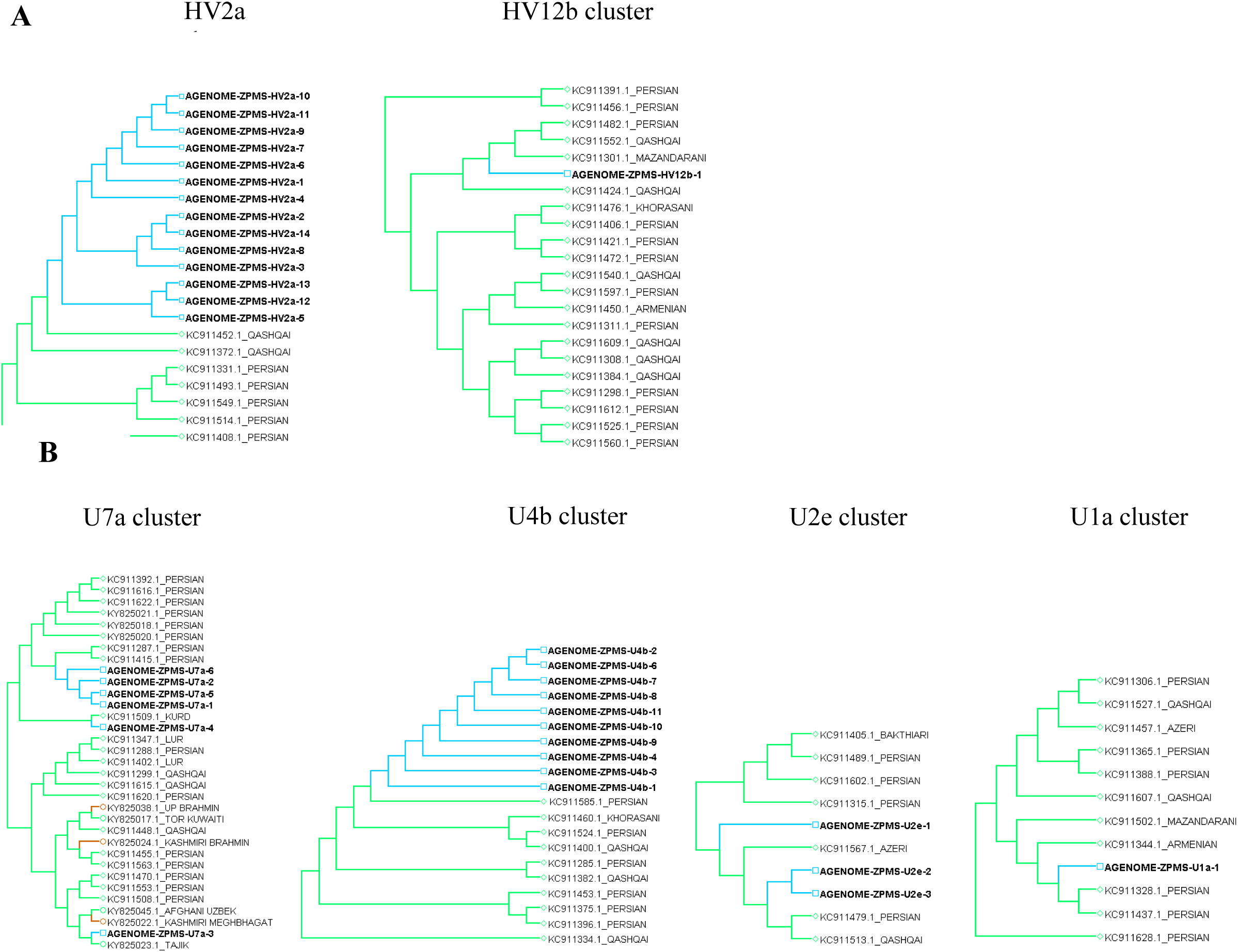
Phylogenetic analysis depicting individual sub-haplogroup clusters of 97 Parsis, 352 Iranian and 100 relic tribes of Indian origin. (A) Representative cladograms of the HV sub-haplogroup (B) Representative cladograms of the U sub-haplogroup

**Figure 4C:**
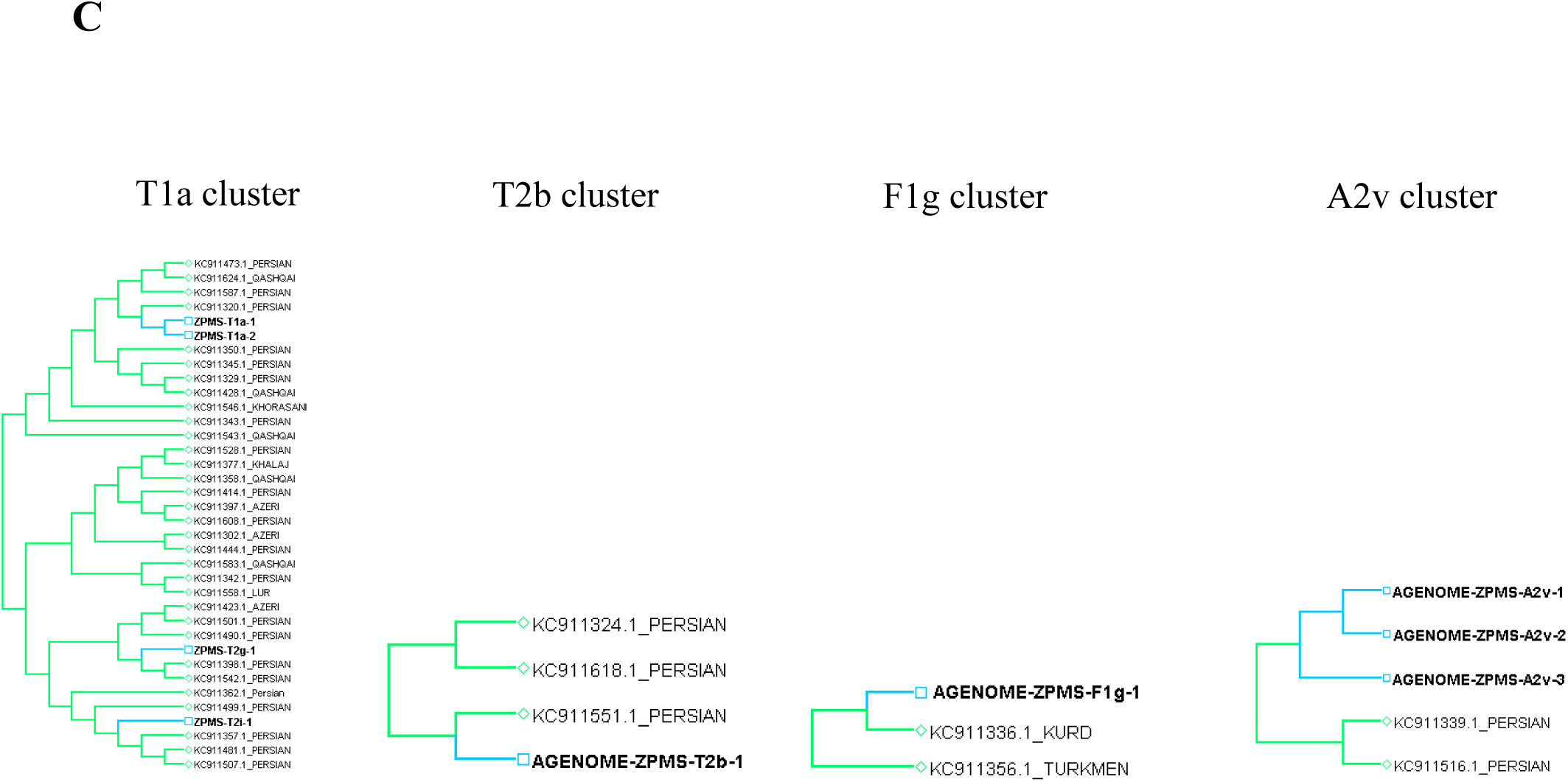
Phylogenetic analysis depicting individual sub-haplogroup clusters of 97 Parsis, 352 Iranian and 100 relic tribes of Indian origin. (C) Representative cladograms of the T, F and A sub-haplogroup

**Figure 4D:**
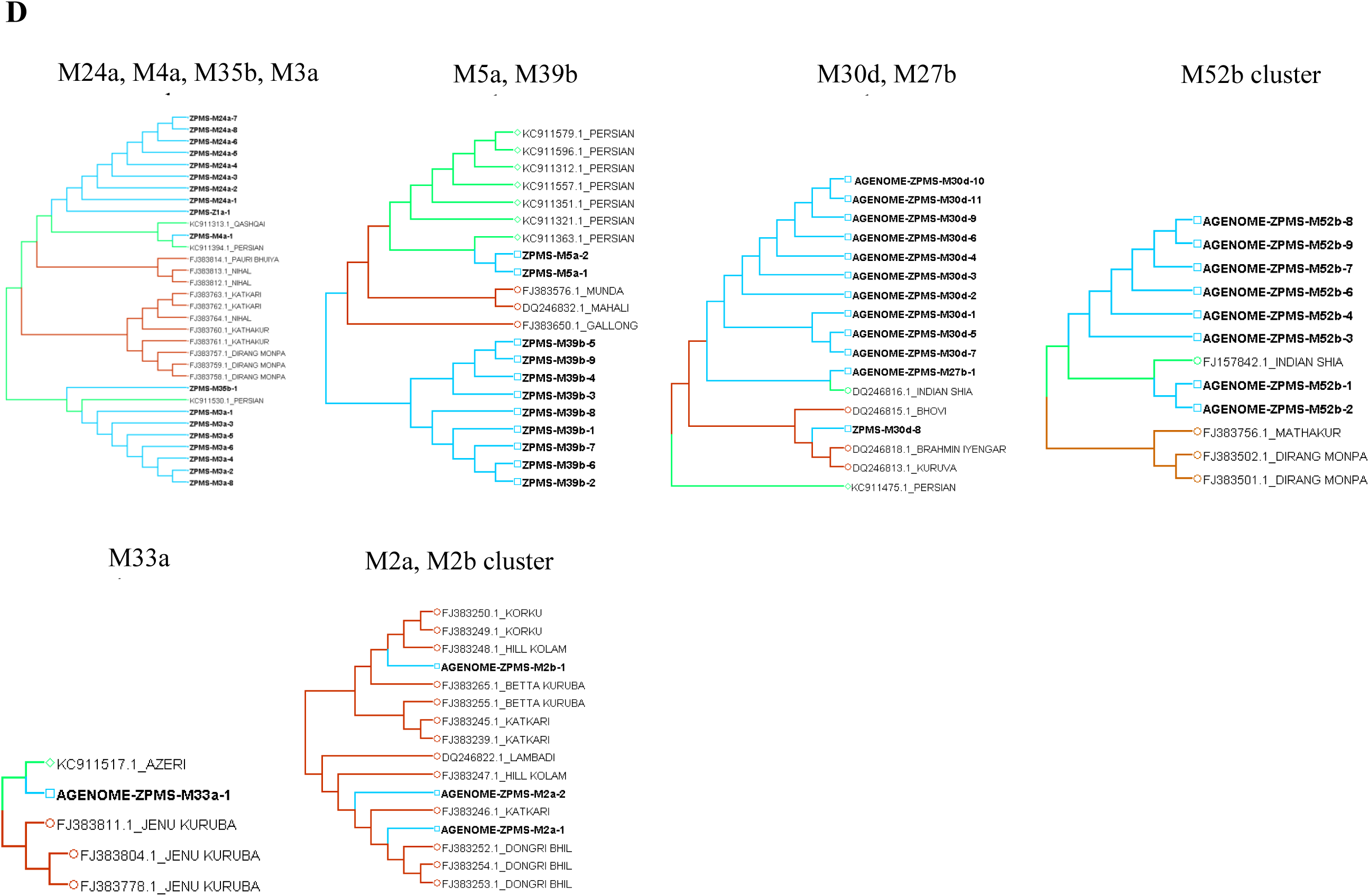
Phylogenetic analysis depicting individual sub-haplogroup clusters of 97 Parsis, 352 Iranian and 100 relic tribes of Indian origin. (A-D) Representative cladograms of the each sub-haplogroup

**Figure 4.**
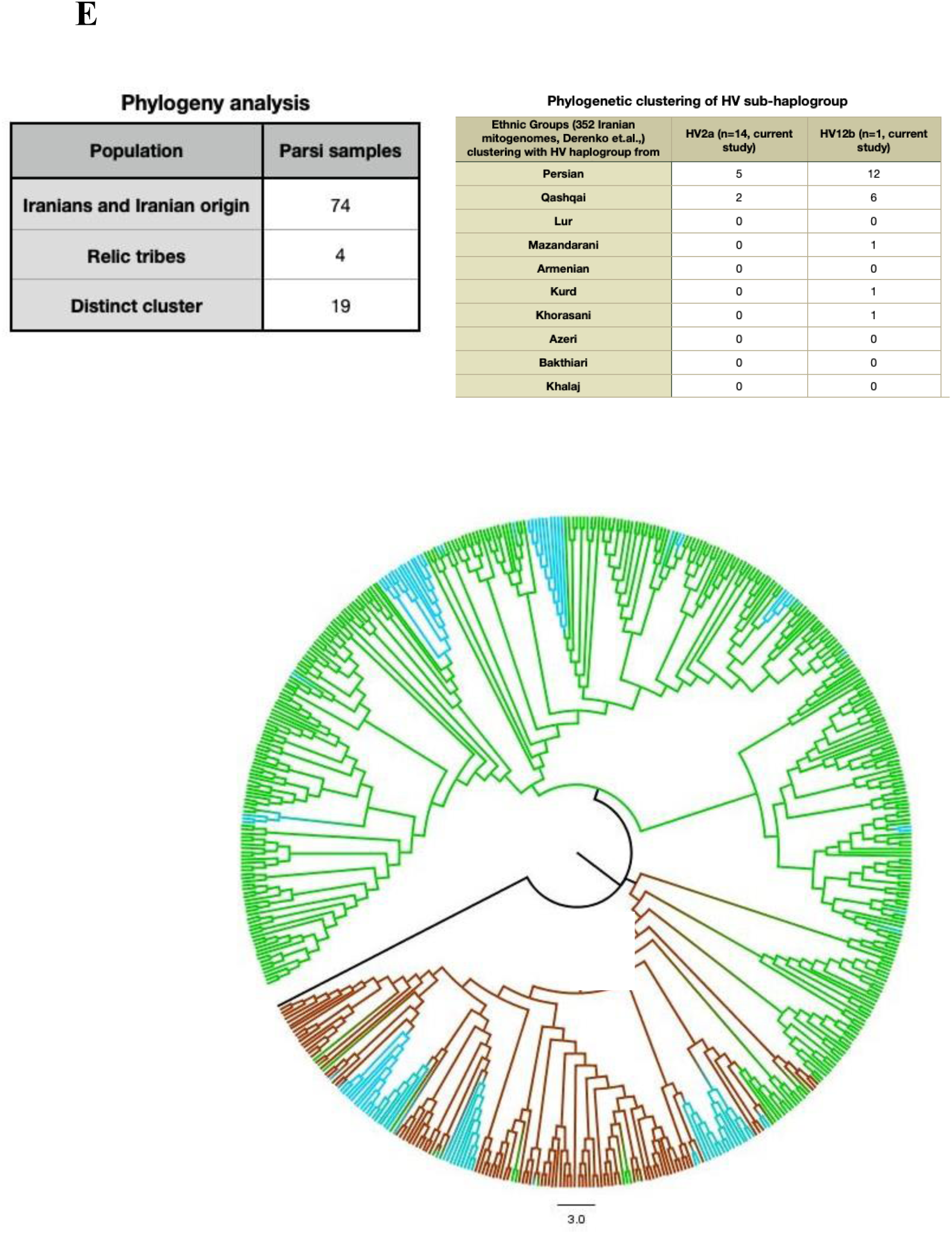
Comparative phylogenetic analysis of individual sub-haplogroup clusters for 97 Parsis, 352 Iranians, and 100 individuals from relic tribes of Indian origin. **A,** HV sub-haplogroup. **B,** U sub-haplogroup. **C,** T, F, and A sub-haplogroups. **D,** M sub-haplogroup.Figure 4A-D represent the zoomed in version of the clustering represented in the complete circular representation in Figure 4E. **E,** Parsi clustering with Iranians, relic tribes of Indian origin, or forming a unique cluster (Table, top left). Results of clustering of the HV2 Parsis with other ethnic groups in the Iranian mitogenome (Table, top right). Whole-mitochondrial genome clustering of Parsis (blue), Iranians (green), and Indians (brown). The outgroup is indicated by the black line

Unlike the HV, U, and T haplogroups, for which the Parsis cluster closely with Persians, the Parsis harboring the M haplogroup appear to demonstrate more diversity in their mitochondrial genomes. This study showed the following breakdown: 8/12 M sub-haplogroups of the 29 Parsi M haplotypes (M24a [n= 8], M33a [n=1], M5a [n=2], M4a [n=1)], M3a [n=7], M52b [n=8], M27b [n=1], and M35b [n=1]) clustered with the Persians, Qashqais, Azeris of Iranian ethnicity, and others of Persian descent (**Figure 4D, Supplementary Table 3**). Only two sub-haplogroups in our study (M2a and M2b [n=21], M30d [n=1], **Figure 4D**) clustered with relic tribes of Indian origin. Our phylogenetic analyses further showed that 19 Parsi individuals belonging to the M30d (n=10) and M39d (n=9) haplogroups did not cluster either with Indian or Iranian ethnic groups (**Figure 4D**) but remained clustered within their own subgroups.

Outgroup sampling is of primary importance in phylogenetic analyses, affecting in-group relationships, and, by correctly placing the root, determining the sequence of branching events. Accordingly, we used the AGENOME-OUTGROUP-Y2b sequence to root the phylogenetic tree. This sequence did not associate with the Parsis, Indians, or Iranians, attesting to the robustness of this method employed for phylogenetic analysis (**Figure 4E**, black line).

### Assembly of the Zoroastrian Parsi Mitochondrial Consensus Genome (AGENOME-ZPMCG-V1.0) and Parsi haplogroup-specific reference sequences

To better understand the nuances of disease and wellness in this unique community, we generated the Zoroastrian Parsi Mitochondrial Consensus Genome (AGENOME-ZPMCG V1.0; Genbank ID, MT506339). We also assembled seven individual haplogroup-based reference genomes, including AGENOME-ZPMRG-HV-V1.0 (n=15; Genbank ID, MT506342), AGENOME-ZPMRG-U-V1.0 (n=20; Genbank ID, MT506345), AGENOME-ZPMRG-T-V1.0 (n=5; Genbank ID, MT506344), AGENOME-ZPMRG-M-V1.0 (n=52; Genbank ID, MT506343), AGENOME-ZPMRG-A2v-V1.0 (Genbank ID, MT506340), AGENOME-ZPMRG-F1a-V1.0 (Genbank ID, MT506341), and AGENOME-ZPMRG-Z-V1.0 (Genbank ID, MT506346) (**Supplementary Table 4, Appendix 2**).

Additionally, using all 100 Parsi mitochondrial genome sequences generated in this study (see Materials and Methods), we built the first Zoroastrian-Parsi mitochondrial consensus genome (AGENOME-ZPMCG-V1.0). The consensus Parsi mtDNA sequence was found to have 31 unique variants (**Supplementary Table 5**), of which five (A263G, A750G, A1438G, A4769G, and A15326G) were found to be common to the reference sequences of all seven haplogroups considered (**Supplementary Table 5**). While the number of variants unique to each of the seven haplogroups ranged from 11 to 33, haplogroup M did not appear to have any unique variants when compared with the overall consensus sequence (AGENOME-ZPMCG-V1.0).

### mtDNA variant-specific disease associations in the non-smoking Parsi cohort

Comparison of mitochondrial sequence data from the WGS of 100 Parsi subjects with the revised Cambridge Reference Sequence (rCRS) standard resulted in identification of 420 distinct variants. Further analysis with VarDiG^®^-R, a database of genes and disease variants, identified 217 unique variants associated with 41 disease phenotypes, which were further classified according to the seven major haplogroups and their 25 sub-haplogroups.

### Haplogroup and disease linkage

Principal component analysis (PCA) showed the assocations between variants and haplogroups. Longevity variants in the Parsi sub-haplogroups were found to be associated with Parkinson’s disease (PD), Alzheimer’s disease (AD), breast cancer, and cardiomyopathy in 23/25 sub-haplogroups (HV2a, U7a, U4b, T1a, T2g, T2i, T2b, M5a, M39b, M33a, M52b, M24a, M3a, M30d, M2a, M4a, M2b, M35b, M27b, A2v, F1g, and Z1a). Longevity variants were absent in only 2/25 sub-haplogroups (HV12b and U1a, **Figure 5A**). We found a close association between variants and PD in most haplogroups (**Appendix 3**), while further analysis revealed linkages to colon cancer in 13/23 longevity-linked sub-haplogroups. Previously reported lung cancer and non-small cell lung cancer-associated variants^28,29^ that were found occurring in the 16S RNR2, *ND5, ND6,* and tRNA genes were absent in the 420 variants in the Parsi population (**Appendix 6**).

**Figure 5A:**
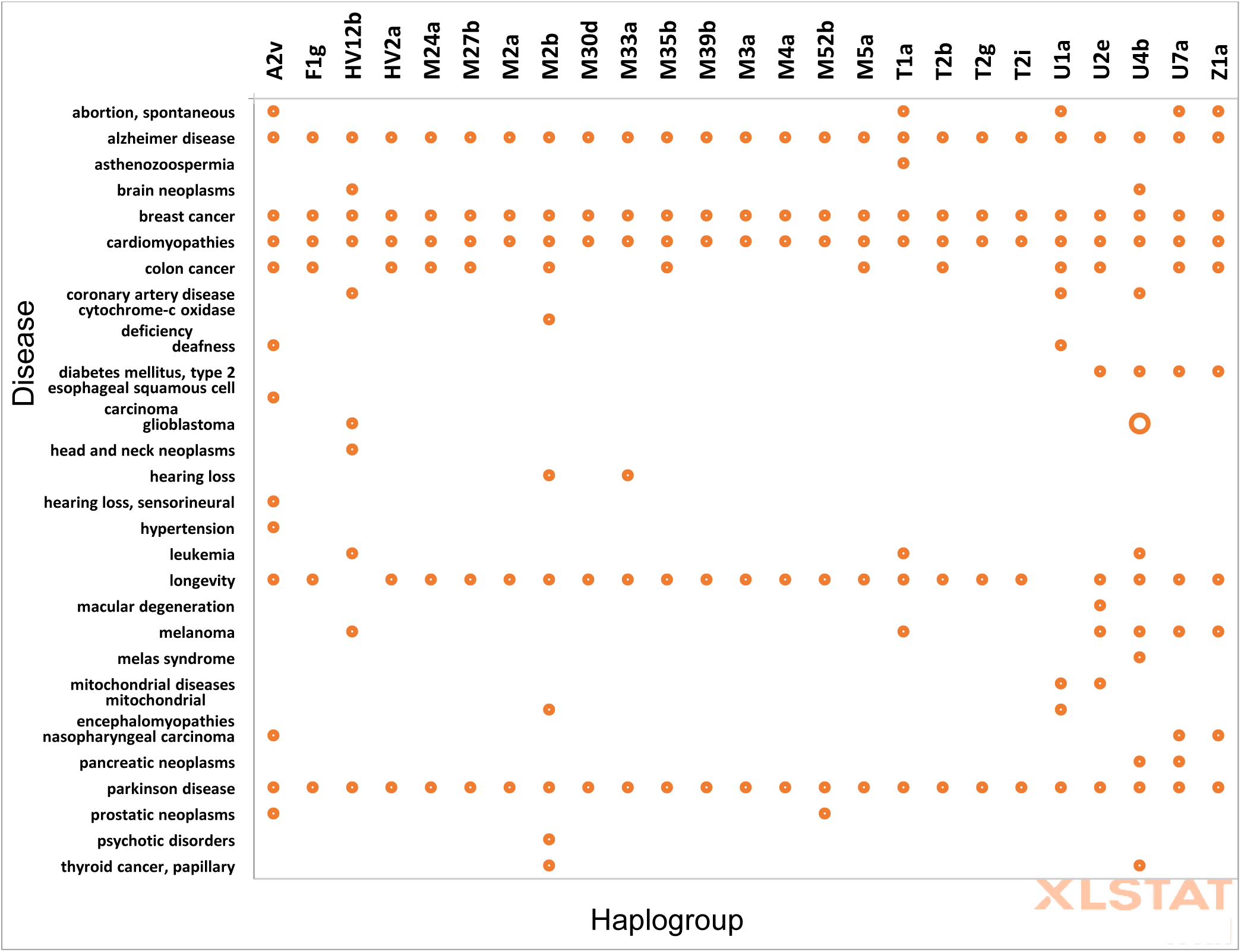
PCA analysis shows absence of Longevity variants in U1a and HV12bsub-haplogroups

**Figure 5B:**
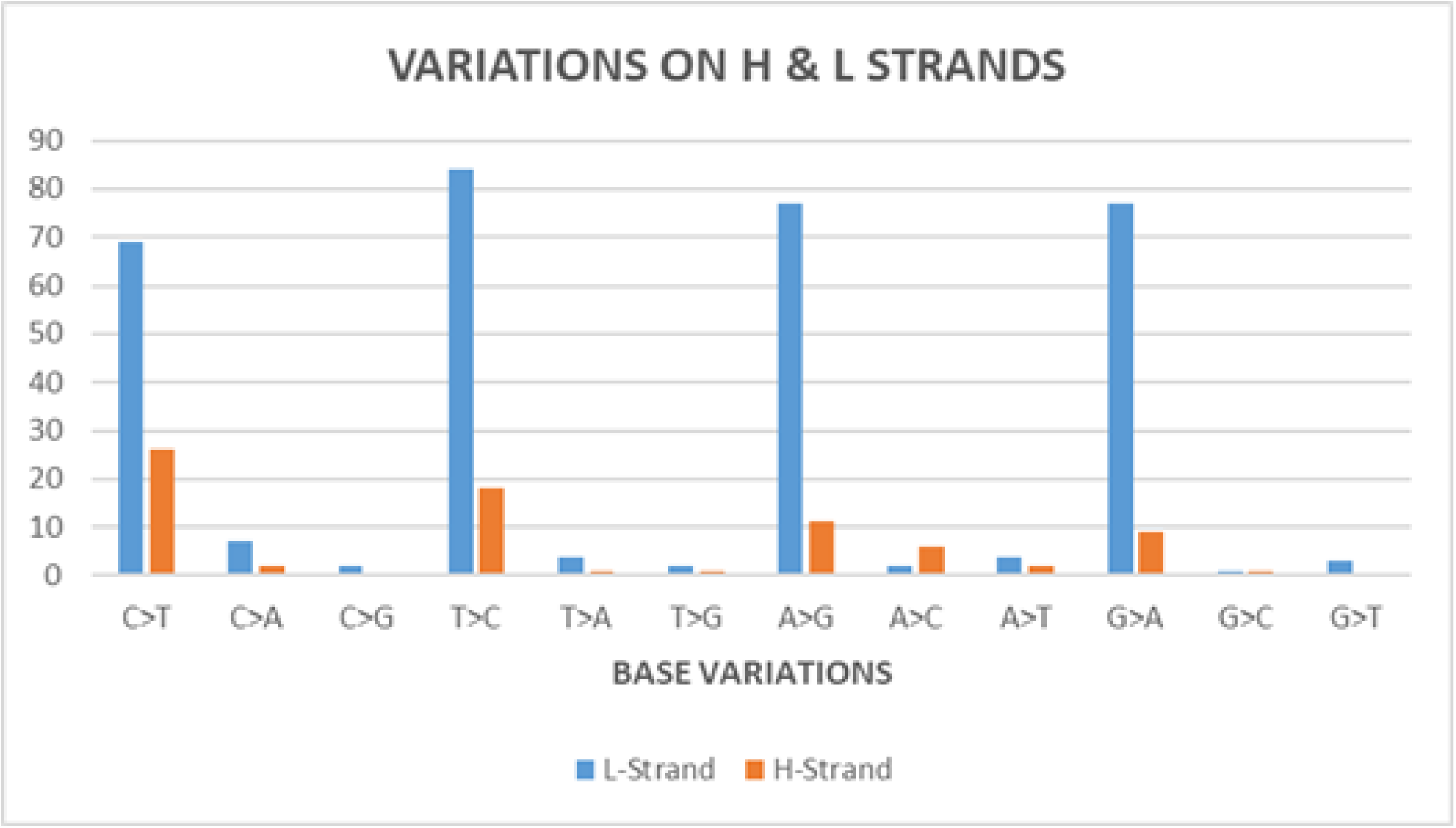
Lack of smoking induced mutational signatures in the Parsi cohort

**Figure 5.** Haplogroup specific disease associations and smoking-related mutational signatures for the 100 Parsi mitochondrial genomes in this study. **A,** Principal component analysis of disease associations with sub-haplogroups. The U1a and F1g sub-haplogroups show an absence of longevity-related disease associations. **B,** Transitions and transversions on the heavy (H) and light (L) strands for the 100 Parsi mitochondrial genomes in this study.

### Variant analysis

Given the importance of mitochondrial heteroplasmy in the etiology of diseases, we implemented a bioinformatic pipeline to detect heteroplasmies in our sample set using Mutserver (mtDNA-Server Version 1.0.7) variant caller for the mitochondrial genome with a minimum heteroplasmy level with a stringent threshold value of 0.05 (5%). Our analysis detected 24 unique high confidence heteroplasmies from the 420 distinct variants across the 100 samples at a minimum heteroplasmy level threshold ≥ 0.05 (5%) and mean coverage ≥ 500X (**Appendix 8**)

Further analysis of the 420 variants revealed a putative association between PD and our variants (**Supplementary Figure 7**), neurodegenerative diseases, rare diseases of mitochondrial origin, and cardiovascular and metabolic diseases in our study. (**Supplementary Figure 7**).

While predispositions for 41 diseases were spread across 25 sub-haplogroups, many disease variants were found to recur across haplogroups, totalling 188 instances of disease variants (**Supplementary Figure 8A**). Haplogroup U4b harbored 15 disease-associated variants, while the majority of M and T groups had 5 variants (Figure 6B). Some of the mitochondrial rare diseases, such as mitochondrial encephalomyopathies, Mitochondrial Encephalopathy, Lactic Acidosis, and Stroke-like episodes (MELAS syndrome), and cytochrome c oxidase deficiency were found to be associated with the M2a and U1a; U4b; and M2b sub-haplogroups, respectively (**Supplementary Figure 8B**).

**Figure 6A:**
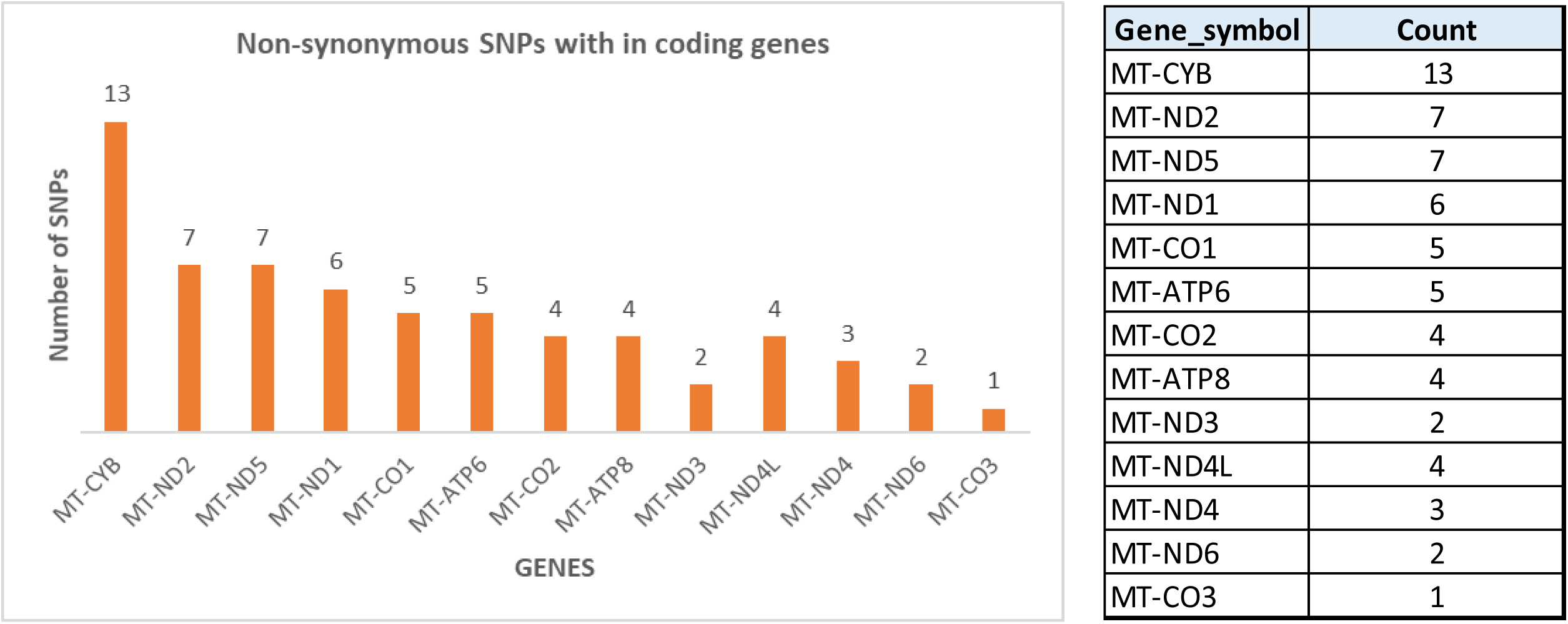
CYTB gene has the highest occurrence of non-synonymous variants in this study

**Figure 6B:**
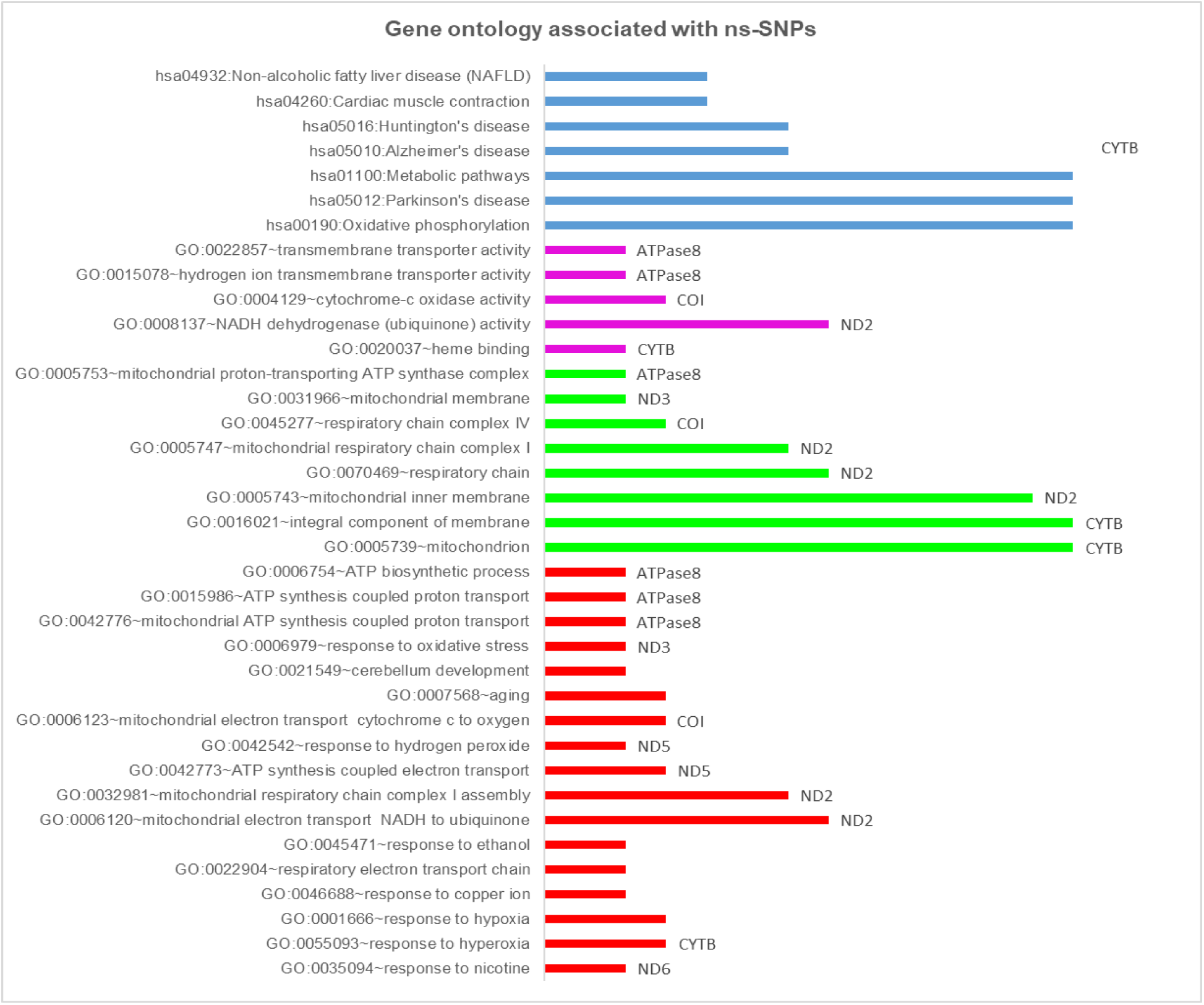
Gene ontology associated with non-synonymous variants among 420 variants

**Figure 6.** Analysis of the nonsynonymous variants in the 420 variants in the 100 Parsi mitochondrial genome sequences. **A,** Occurrence of the nonsynomymous variants within coding gene loci of the mitochondrial genome, as analyzed with the MitImpact database. Note that the *CYTB* gene has the highest occurrence. **B,** Gene ontology analysis of the nonsynonymous variants using the DAVID and UNITPROT annotation tools.

Further analysis of the nucleotide transitions and transversions that constitute the 420 variants revealed that the mutational signatures (C>A and G>T) found in tobacco smoke-derived cancers^30^ were found at an extremely low frequency (<6% compared with other mutational signatures) on both the heavy (H) and light (L) strands of the mitochondrial genomes of the Parsi population (**Figure 5B**), who are known to refrain from smoking due to their religious and social habits.

### Analysis of the variants in tRNA genes and the D-loop region in the mitochondrial genome

In order to determine whether diseases known to be prevalent in the Parsi community could in fact be predicted by association using the collective mitochondrial variants discovered in this study, we first analyzed variants identified in tRNA genes that have previously been implicated in rare and degenerative diseases. We found a total of 17 tRNA-associated variants, with a pathogenic variant (G1644A) implicated significantly in adult onset-Leigh Syndrome (LS)/Hypertrophic CardioMyopathy (HCM)/MELAS, a genetically inherited mitochondrial disease^31^. We also found a total of six tRNA mutations associated with nonsyndromic hearing loss, hypertension, breast/prostate cancer risk, and progressive encephalopathies in the analysis of our 100 Zoroastrian-Parsi individuals (**Supplementary Table 6**).

While synonymous/neutral variants in mtDNA genome sequences do not affect mitochondrial function, nonsynonymous/non-neutral variants may have functional consequences. We therefore analyzed the 420 variants from 100 Parsi subjects for nonsynonymous mutations and identified 63 such variants located within different mitochondrial genes (**Figure 6A**). Twenty of 63 variants were found in the genes encoding *CYTB* (n=13) and *ND2* (n=7), followed by *ND5* and *ND1*. Annotation of disease pathway-association analysis with MitImpact server, showed the association of non-synonymous variants in our study with disease pathways for neurodegenerative conditions, such as AD and PD; cancers of colorectal and prostate origin; metabolic diseases, such as type 2 diabetes; and rare diseases, such as Lebers Hereditary Optic Neuropathy (LHON) (*CYTB* and *ND2*) (**Supplementary Figures 9 and 10**). Variants implicated in longevity were observed in our study and distributed across the *ND2* gene (**Supplementary Figure 8B**). As mentioned above, we found no association between the nonsynonymous variants in our data set and lung cancer.

To understand the mitochondrial pathways affected by the non-synonymous variants in our study, we annotated the variants with DAVID and UNIPROT and found that the major genes *CYTB* and *ND2* were implicated in pathways that include the mitochondrial respiratory complex (*COI/COII/COIII/COIV*), OXPHOS, and metabolic pathways implicated in mitochondrial bioenergetics. Critical disease-related pathways in PD, AD, and cardiac muscle contraction were also associated with *CYTB*- and *ND2*-specific variants, which possibly explains the high incidence of these diseases in the Parsi population (**Supplementary Figure 10**).

A total of 87 variants, including 6 unique variants, were observed in the D-loop region across all 25 sub-haplogroups (n=100 subjects, **Supplementary Table 2**). Seventy-four of 100 Parsis in our study were found to have the polymorphism m.16519 T>C. Six subjects of the M52 sub-haplogroup were found to have the m.16525 A>G substitution. The rest of the variants were m.16390 G>A (n=4 subjects) and m.16399 A>G, m.16401 C>T, and m.16497 A>G (all with n=1 subject each).

### Identification of unique, unreported variants from the mitogenome analysis of 100 Parsi subjects

We performed a comparative analysis of the 420 variants in the Parsi community with MITOMASTER^32^, a database that contains all known pathogenic mtDNA mutations and common haplogroup polymorphisms, to identify unique variants in our population that were not previously reported. Our analysis showed the presence of 12 unique variants distributed across 27 subjects that were not observed in MITOMASTER nor in the VarDIG^®^-R disease-association dataset (**Figure 7, Appendix 5**). These unique variants were observed at different gene loci, including 12S rRNA (2 variants), 16S rRNA (5 variants), and 1 variant each in the *ND1*, *COII*, *COIII*, *ND4*, and *ND6* genes. SNP haplogroup-association analysis showed that they fell into four major haplogroups and 13 sub-haplogroups: HV2a (n=1), M24a (n=4), M2a (n=1), M30d (n=3), M35b (n=1), M39b (n=2), M3a (n=1), M4a (n=1), M52b (n=4), M5a (n=1), T2b (n=1), U4b (n=6), and U7a (n=1). Of the 12 variants identified, no disease associations were observed by analysis with MITOMASTER or VarDIG^®^-R.

**Figure 7:**
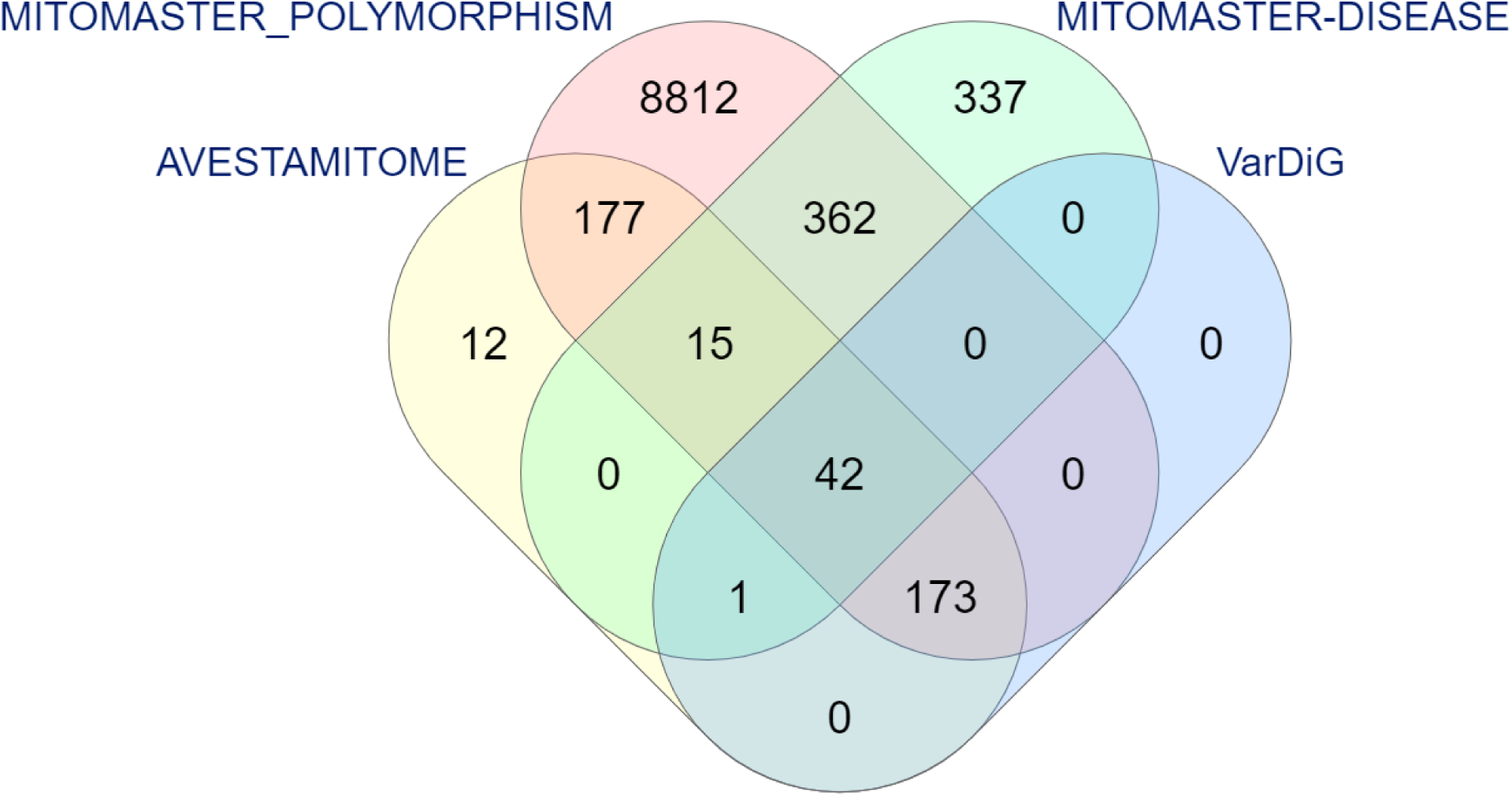
12 unique variants found in the current study. Comparative analysis of the 420 variants in the AVESTAMITOME^™^ Zoroastrian-Parsi community dataset with common and disease-associated polymorphisms in the MITOMASTER database and the VarDiG-R search engine. Twelve unique variants were found in the current study.

## Discussion

The first *de novo* Parsi mitochondrial genome, AGENOME-ZPMS-HV2a-1 (Genbank accession, MT506314), from a healthy, non-smoking female of haplogroup HV2a showed 28 unique variants compared with the revised Cambridge Reference Standard (rCRS). Upon extending our mitochondrial genome analyses to an additional 99 Parsi individuals, we found that 94 individuals belonged to four major mitochondrial haplogroupsHV, U, T, A (belonging to the macrohaplogroup N) and other M descendents of the macrohaplogroup M (M5, M39, M33, M44’52, M24, M3, M30, M2, M4’30, M2, M35 and M27), while 5 individuals belonged to the rarer haplogroups A, F, and Z. The largest sub-haplogroup was found to be HV2a (n=14).

Phylogenetic analysis of the major mitochondrial haplogroups in our Parsi cohort with 352 Iranian^22^ and 100 Indian mitochondrial genomes^23–25^, revealed that the Parsi genomes are phylogenetically related to the Persians and Qashqais^22^ in the HV, T, U, F, A, and Z haplogroups, which are those associated with the peopling of western Europe, Central Asia, and the Iranian plateau.

The haplogroup HV2 most likely arose in Persia, and the subclade HV2a has a demonstrated Persian ancestry. HV12b, a branch of the HV12 clade, is one of the oldest HV subclades and has been found in western Iran, India, and sporadically as far away as Central and Southeast Asia. It has strong associations with the Qashqais, who are Turkic-speaking nomadic pastoralists of southern Iran and who previously resided in the Iranian region of the South Caucasus^33,36^. Among the U haplogroup, the U4b and U7a haplotypes are distributed throughout the Central Asia in the Volga–Ural region^34^, South Asia^25^, and with lower frequencies in populations around the Baltic Sea^33^. Haplogroup U2 is found primarily in South Asia, whereas U2d and U2e are confined to the Near East and Europe^24^. The T haplogroup is also widely distributed in Eastern and Northern Europe, the Indus Valley, and the Arabian Peninsula following expansion during the Neolithic transition^34^. The presence of the predominantly Eurasian mtDNA haplotypes (HV, T, U, F, A, and Z) in our Parsi cohort attests to their practice of endogamy, given that the Parsis have resided on the Indian subcontinent for over 1300 years.

Despite the high frequency of the M haplogroup (the largest haplogroup in the Indian subcontinent^35^) in our Parsi cohort, phylogenetic analysis showed that 47/51 Parsis belonging to the M haplogroups in our study cluster with the Persians, suggesting Persian descent, with a small minority of Parsis found to be related to relic tribes of India. This observation suggests minimal gene flow from indigenous Indian females into the Parsi gene pool, as was previously proposed^26^. Phylogenetic analysis also revealed that two Parsi M sub-haplogroups, M30d and M39b, formed a unique cluster that needs further resolution.

We further present the first complete Zoroastrian Parsi mitochondrial consensus genome (AGENOME-ZPMCG V1.0), built from the mitochondrial genomes of 100 non-smoking Parsi individuals, representing seven mitochondrial haplogroups. The generation of a unique population-specific consensus genome for the Parsis is useful for comparative analyses and in reconstructing their population history, migration pattern, and disease associations.

We found that the *CYTB* gene contained the greatest number of variants (n≥5) in the coding region of haplogroup M, besides having the greatest representation in the F1g, T, and HV12b haplogroups. Haplogroups U, A2v, and Z1a showed a predominance of the variants linked to the ND complex genes *ND5* and *ND2*, while the *COI* gene variants were the most highly represented in HV2a and U4b. Variants in the *CYTB* gene are associated with Alzheimer’s disease (AD), diabetes mellitus, cognitive ability, breast cancer, hearing loss, and asthenozoospermia and are associated with changes in metabolic pathways, cardiac contraction, and rare diseases, such as Huntington’s disease, whereas the *ND2* and *ND5* variants are associated with prostate cancer; ovarian cancer; rare mitochondrial neuronal diseases, such as LHON; cardiomyopathy; AD; and Parkinson’s disease (PD).

Interrogation of the 420 variants across seven haplogroups in the Parsi cohort using the VarDIG^®^-R database revealed that PD, known to be prevalent in the Parsi community^37^, was the most prevalent, with 178 of the 420 variants represented. Not surprisingly, longevity, which often co-occurs with PD, was also predicted to be highly prevalent in the Parsi cohort, but with a notable absence in the U1 sub-haplogroup, an interesting observation that warrants further investigation.

Analysis of additional disease associations revealed that variants related to AD (also related to ageing), breast cancer, and cardiomyopathies^38–40^, were all the 25 Parsi sub-haplogroups. Additionally, the presence of variants associated with asthenozoospermia^41^ in the T1a sub-haplogroup, a condition associated with reduced sperm motility. The ‘T1a’ is a rare group in our analysis of the 100 mitogenomes sequenced (2/100) perhaps indicative of a slow decline of this particular haplogroup in the population moving to a possible extinction as it is a documented that the fertility rates in the community is on a steady decline.

It is noteworthy that previously published epidemiological studies demonstrating lower rates of lung cancer among the Parsis^42^, appears to have a genetic basis, given that no haplogroup in the Parsi cohort displayed known lung cancer-associated variants. The low frequency of mutational signatures for tobacco smoke-derived cancers, is in line with the non-smoking customs of the Parsi community.

The tRNA disease-association analysis in our study showed that these genes were implicated in the onset of neurodegenerative conditions, such as AD; PD; cancers of colorectal and prostate origin; metabolic diseases, such as type 2 diabetes; and rare diseases, such as LHON (*CYTB* and *ND2*). The D-loop SNP analysis showed the prevalence (74/100 subjects) of the m.16519 T>C polymorphism, which has been implicated in chronic kidney disease^43^, an increased risk of Huntington’s disease, cyclic vomiting syndrome^44^, schizophrenia, and bipolar disorder^45^. Taken together, these results warrant a deeper investigation into the tRNA and the D-loop variants in the Parsi community.

Our Parsi population genetics study has shown for the first time the existence of haplogroup-specific variants and their disease associations with longevity, neurodegenerative diseases, cancers, and rare disorders. The Parsis represent a small, unique, non-smoking community in which genetic signatures maintained by generations of endogamy, provide an exceptional opportunity to understand genetic predispositions to various diseases.

## Methods

### Sample collection and ethics statement

One hundred healthy, non-smoking Parsi volunteers residing in the cities of Hyderabad-Secunderabad and Bangalore, India were invited to attend blood collection camps at the Zoroastrian centers in their respective cities under the auspices of The Avestagenome Project™. Each adult participant (>18 years) underwent height and weight measurements and answered an extensive questionnaire designed to capture their medical, dietary, and life history. All subjects provided written informed consent for the collection of samples and subsequent analysis. All health-related data collected from the cohort questionnaire were secured in The Avestagenome Project™ database to ensure data privacy.

### Genomic DNA extraction

Genomic DNA from the buffy coat of peripheral blood was extracted using the Qiagen Whole Blood and Tissue Genomic DNA Extraction kit (cat. #69504). Extracted DNA samples were assessed for quality using the Agilent Tape Station and quantified using the Qubit™ dsDNA BR Assay kit (cat. #Q32850) with the Qubit 2.0^®^ fluorometer (Life Technologies™). Purified DNA was subjected to both long-read (Nanopore GridION-X5 sequencer, Oxford Nanopore Technologies, Oxford, UK) and short-read (Illumina sequencer) sequencing.

### Library preparation for sequencing on the Nanopore platform

Libraries of long reads from genomic DNA were generated using standard protocols from Oxford Nanopore Technology (ONT) using the SQK-LSK109 ligation sequencing kit. Briefly, 1.5 µg of high-molecular-weight genomic DNA was subjected to end repair using the NEBNext Ultra II End Repair kit (NEB, cat. #E7445) and purified using 1x AmPure beads (Beckman Coulter Life Sciences, cat. #A63880). Sequencing adaptors were ligated using NEB Quick T4 DNA ligase (cat. #M0202S) and purified using 0.6x AmPure beads. The final libraries were eluted in 15 µl of elution buffer. Sequencing was performed on a GridION X5 sequencer (Oxford Nanopore Technologies, Oxford, UK) using a SpotON R9.4 flow cell (FLO-MIN106) in a 48-hr sequencing protocol. Nanopore raw reads (fast5 format) were base called (fastq5 format) using Guppy v2.3.4 software. Samples were run on two flow cells and generated a dataset of ∼14 GB.

### Library preparation and sequencing on the Illumina platform

Genomic DNA samples were quantified using the Qubit fluorometer. For each sample, 100 ng of DNA was fragmented to an average size of 350 bp by ultrasonication (Covaris ME220 ultrasonicator). DNA sequencing libraries were prepared using dual-index adapters with the TruSeq Nano DNA Library Prep kit (Illumina) as per the manufacturer’s protocol. The amplified libraries were checked on a Tape Station (Agilent Technologies) and quantified by real-time PCR using the KAPA Library Quantification kit (Roche) with the QuantStudio-7flex Real-Time PCR system (Thermo). Equimolar pools of sequencing libraries were sequenced using S4 flow cells in a Novaseq 6000 sequencer (Illumina) to generate 2 x 150-bp sequencing reads for 30x genome coverage per sample.

### Generation of the d*e novo* Parsi mitochondrial genome (AGENOME-ZPMS-HV2a-1)

#### a) Retrieval of mitochondrial reads from whole-genome sequencing (WGS) data

A total of 16 GB of raw data (.fasta) was generated from a GridION-X5 Nanopore sequencer for AGENOME-ZPMS-HV2a-1 from WGS. About 320 million paired-end raw reads were generated for AGENOME-ZPMS-HV2a-1 by Illumina sequencing.

Long Nanopore reads (. fastaq5) were generated from the GridION-X5 samples. The high-quality reads were filtered (PHRED score =>20) and trimmed for adapters using Porechop (v0.2.3). The high-quality reads were then aligned to the human mitochondrial reference sequence (rCRS) NC_12920.1 using Minimap2 software. The aligned SAM file was then converted to a BAM file using SAMtools. The paired aligned reads from the BAM file were extracted using Picard tools (v1.102).

The short Illumina high-quality reads were filtered (PHRED score =>30). The adapters were trimmed using Trimgalore (v0.4.4) for both forward and reverse reads, respectively. The filtered reads were then aligned against a human mitochondrial reference (rCRS^21^) using the Bowtie2 (v2.2.5) aligner with default parameters. The mapped SAM file was converted to a BAM file using SAMtools, and the mapped paired reads were extracted using Picard tools (v1.102).

#### b) *De novo* mitochondrial genome assembly

Mapped reads were used for *de novo* hybrid assembly using the Maryland Super-Read Celera Assembler (MaSuRCA-3.2.8) tool. The configuration file from the MaSuRCA tool was edited by adding appropriate Illumina and Nanopore read files. The MaSuRCA tool uses a hybrid approach that has the computational efficiency of the de Bruijn graph methods and the flexibility of overlap-based assembly strategies. It significantly improves assemblies when the original data are augmented with long reads. AGENOME-ZPMS-HV2a-1 was generated by realigning the mapped mitochondrial reads from Illumina as well as Nanopore data with the initial assembly.

### Confirmation of variants in the *de novo* Parsi mitochondrial genome using Sanger sequencing

To validate the *de novo* Parsi mitochondrial sequence (AGENOME-ZPMS-HV2a-1), selected variants were identified and subjected to PCR amplification. Genomic DNA (20 ng) was PCR amplified using LongAmpTaq 2X master mix (NEB). The PCR amplicons of selected regions were subjected to Sanger sequencing and BLAST analysis to confirm the presence of eight variants using the primers listed in Supplemental Table 1.

### Generation of the Zoroastrian-Parsi Mitochondrial Consensus Genome (AGENOME-ZPMCG-V1.0) and Parsi haplogroup-specific consensus sequences

#### a) Retrieving mitochondrial reads from 100 Parsi whole-genome sequences

The whole-genome data from 100 Parsi samples were processed for quality assessment. The adapters were removed using the Trimgalore 0.4.4 tool for paired end reads (R1 and R2), and sites with PHRED scores less than 30 and reads shorter than 20 bp in length were removed. The processed Illumina reads were aligned against a human mitochondrial reference sequence (rCRS^21^, NC_012920.1) using the Bowtie 2 (v2.4.1) aligner with default parameters. Mapped reads were further used for the *de novo* assembly using SPAdes (v3.11.1), Velvet, and IVA (v1.0.8). Comparison of the assembly and statistics were obtained using Quast (v5.0.2). The assembled scaffolds were subjected to BLASTn against the NCBI nonredundant nucleotide database for validation.

Additionally, we have implemented an extra QC step to deal numt sequences by implementing RtN pipeline^46^ that retains reads that map using sequence similarity to an extensive database of publicly available mitochondrial genomes. RTN uses annotated genomes from HmtDB. RtN! removes low-level sequencing noise and mitochondrial paralogs while not impacting variant calling. It retains mitochondrial reads from the input .bam file that are an exact match to known mitochondrial genome sequences in the HmtDB, otherwise it’s mapping quality is set to 0. RTN also maps to database of annotated allele

#### b) Variant calling, hetroplasmy detection and haplogroup classification

Sequencing reads were mapped to the human mitochondrial genome (rCRS^21^) assembly using the MEM algorithm of the Burrows–Wheeler aligner (v0.7.17-r1188) with default parameters. Variants were called using SAMtools (v1.3.1) to transpose the mapped data in a sorted BAM file and calculate the Bayesian prior probability. Next, Bcftools (v1.10.2) was used to calculate the prior probability distribution to obtain the actual genotype of the variants detected. The classification and haplogroup assignment were performed for each of the 100 Parsi mtDNAs after variant calling and after mapping reference and alternate alleles to the standard haplogroups obtained from MITOMAP (**Appendix 4**).

For the mitochondrial heteroplasmy analysis, we implemented a bioinformatic pipeline to detect heteroplasmies in our sample set using Mutserver run locally (mtDNA-Server Version 1.0.7) variant caller for the mitochondrial genome with a Minimum heteroplasmy level with a stringent threshold value of 0.05 (5%). Our threshold/cutoff was based on literature evidence that indicated a cut off of 50–60% for high levels of mutant mtDNA alleles for the emergence of mitochondrial pathology while further evidence of lower levels of heteroplasmy (not exceeding 30–40%) of certain mtDNA mutations increase the risk of age-related chronic diseases^47^.

#### c) Haplogroup-based consensus sequence

Ninety-seven of 100 full-length Parsi mitogenome sequences were segregated based on haplogroups and separately aligned using the MUSCLE program to obtain the multiple sequence alignments. The Zoroastrian-Parsi Mitochondrial Reference Genome (ZPMRG) and the Parsi haplogroup-specific consensus sequences were generated after calculation of the ATGC base frequency by comparison of the nucleotides in an alignment column to all other nucleotides in the same column called for other samples at the same position. The highest frequency (%) was taken to build seven Parsi haplogroup ZPMRGs and the seven Parsi haplogroup-specific consensus sequences.

### Phylogeny build and analysis

Ninety-seven of 100 full-length Parsi mitogenome sequences generated as described above were compared with 100 randomly chosen Indian mtDNA sequences derived from NCBI Genbank under the accession codes FJ383174.1-FJ 383814.1^23^, DQ246811.1-DQ246833.1^24^, and KY824818.1-KY825084.1^25^ and from previously published data on 352 complete Iranian mtDNA sequences^22^. All mtDNA sequences were aligned using MUSCLE software^48^ using the “maxiters 2” and “diags 1” options, followed by manual verification using BioEdit (v7.0.0). Following alignment, the neighbor-joining method, implemented in MEGAX^49^, was employed to reconstruct the haplotype-based phylogeny. This method was used, because it is more efficient for large data sets^50^.

### Variant disease analysis

One hundred Parsi mitochondria sequences extracted from the WGS were uploaded into the VarDiG^®^-R search engine (https://vardigrviz.genomatics.life/vardig-r-viz/) on AmazonWeb Services. VarDiG-R, developed by Genomatics Private Ltd, connects variants, diseases, and genes in the human genome. Currently, the VarDiG-R knowledgebase contains manually curated information on 330,000+ variants and >20 K genes covering >4500 phenotypes, including nuclear and mitochondrial regions for 150,000+ published articles from 388+ journals. Variants obtained from Parsi mitochondria were mapped against all the published variants in VarDiG-R. Associations with putative diseases were ascertained for each variant through VarDIG-R.

Seventeen tRNA SNP sites were identified in the 100 Parsi mitochondrial SNP data. The PON-mt-tRNA database^51^ was downloaded to annotate the tRNA variants for their impact and disease associations. This database employs a posterior probability-based method for classification of mitochondrial tRNA variations. PON-mt-tRNA integrates the machine learning-based probability of pathogenicity and the evidence-based likelihood of pathogenicity to predict the posterior probability of pathogenicity. In the absence of evidence, it classifies the variations based on the machine learning-based probability of pathogenicity.

For annotation of disease pathways associated with variants, we employed MitImpact (https://mitimpact.css-mendel.it/) to predict the functional impact of the nonsynonymous variants on their pathogenicity. This database is a collection of nonsynonymous mitochondrial variants and their functional impact according to various databases, including SIFT, Polyphen, Clinvar, Mutationtester, dbSNP, APOGEE, and others. The disease associations, functional classifications, and engagement in different pathways were determined using the DAVID and UNIPROT annotation tools.

### Haplogroup and disease linkage

Principal component analysis (PCA) was performed to visualize the linkage of the haplogroup with disease. XLSTAT (Addinsoft 2020, New York, USA. https://www.xlstat.com) was used for statistical and data analysis, including PCA.

## List of abbrevations

mtDNA: mitochondrial DNA
rCRS: revised Cambridge Reference Sequence
NGS: next-generation sequencing
ZPMS: Zoroastrian Parsi Mitochondrial Sequence
ZPMRG: Zoroastrian Parsi Mitochondrial Reference Genome
ZPMCG: Zoroastrian Parsi Mitochondrial Consensus Genome
PCA: Principal Component Analysis
AD: Alzheimer’s disease
PD: Parkinson’s disease
LHON: Lebers Hereditary Optic Neuropathy
MELAS: Mitochondrial Encephalopathy Lactic Acidosis and Stroke-like episodes

## Declarations

### Ethics approval and consent to participate

The samples of peripheral blood collected in this study involve human healthy donors and were obtained with their informed consent. were in accordance with the ethical standards of the institution (Avesthagen Limited, Bangalore, India) and in line with the 1964 Helsinki declaration and its later amendments. One hundred healthy, non-smoking Parsi volunteers residing in the cities of Hyderabad-Secunderabad and Bangalore, India were invited to attend blood collection camps at the Zoroastrian centers in their respective cities under the auspices of The Avestagenome Project™. Each adult participant (>18 years) underwent height and weight measurements and answered an extensive questionnaire designed to capture their medical, dietary, and life history. All subjects provided written informed consent for the collection of samples and subsequent analysis. This study was approved by the Avesthagen Ethics Committee constituted under the Department of Biotechnology, Government of India (BLAG-CSP-033).

### Consent for publication

All subjects have provided written informed consent for the collection of samples and subsequent analysis.

### Availability of data and materials

The GenBank (http://www.ncbi.nlm.nih.gov/genbank) accession numbers for the 105 novel and complete mtDNA sequences (97 ZPMS, 7 ZPMRG, and 1 ZPMCG) reported in this article are numbered MT506242–MT506346 sequentially. The raw reads for 97 ZPMS mitochondrial genome sequences have been deposited with BioProject ID: PRJNA636291. The SRA accession numbers for the 97 ZMPS sequences is SRR11888826-SRR11888922.

### Competing interests

The authors declare that they have no competing interests

### Funding

The project was funded by the grant awarded to Dr.Villoo Morawala-Patell “Cancer risk in smoking subjects assessed by next-generation sequencing profile of circulating free DNA and RNA” (GG-0005) by the Foundation for a Smoke-Free World, New York, USA.

### Authors contributions

VMP conceptualized, designed, guided the experiments and analysis; VMP founded The Avestagenome Project™ and provided access to the dataset for this study; NP and CG analysed the sequences, performed bioinformatics analysis, and interpreted the results; RM, SR, and NS coordinated wet-lab work flows and data analysis; VMP, NP, BM, and KK analysed the data and plotted the graphs and figures; VMP, AKG, PB, KK, and RJ drafted the manuscript, with input from RM, SR, NS, and CG. All authors reviewed the manuscript.

## Acknowledgements

We thank the Foundation for Smoke-Free World, who is advancing global progress in smoking cessation and harm reduction, for funding this project and Dr. Derek Yach for his support of this project. We thank National Institute of Bio Medical Genetics, [NIBMG], Kolkata and Center for Cellular and Molecular Biology [CCMB], Hyderabad for their excellent sequencing services. We thank the Zoroastrian-Parsi community of India for their enthusiastic cooperation. We thank Dr.Sami Gazder, Kouser Sonnekhan, and the The Avestagenome Project™ project team.

## Supplementary Figures and Tables

**Supplementary Figure 1:**
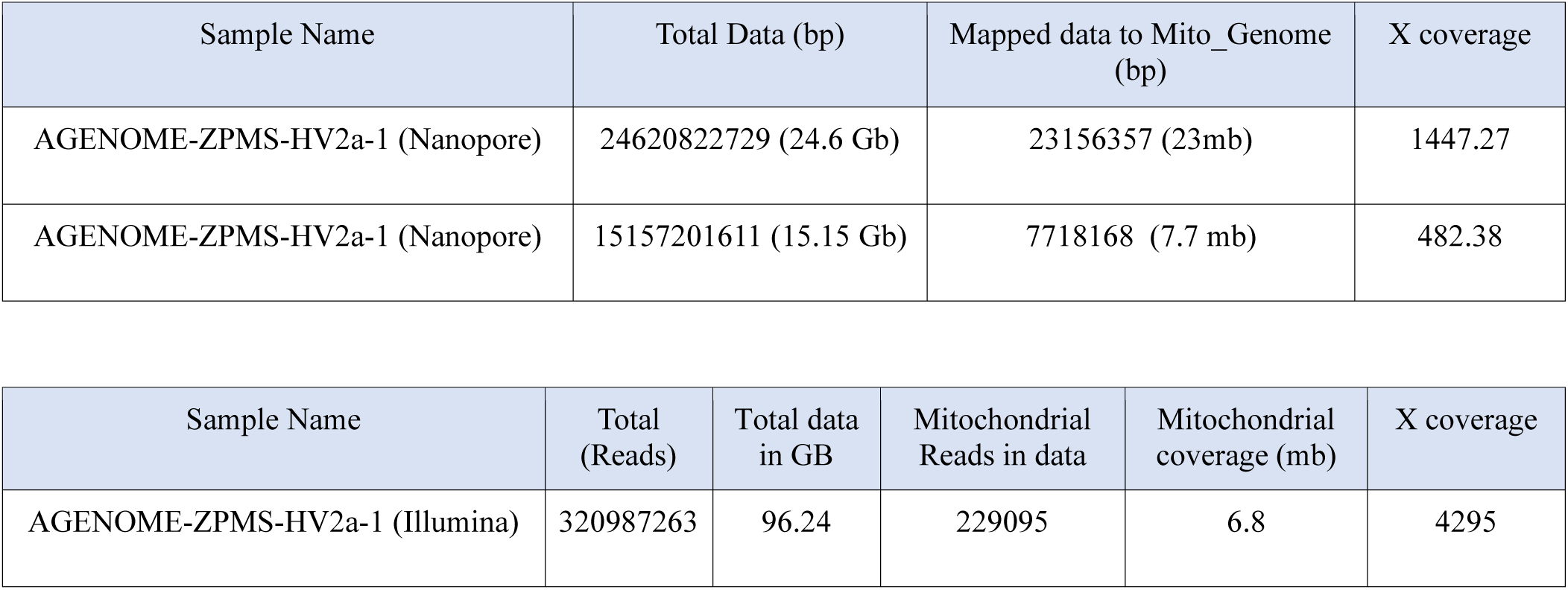
QC data of the *de novo* Zoroastrian Parsi Mitochondrial Reference Genome (AGENOME-ZPMRG-HV2a-1)

**Supplementary Figure 2:**
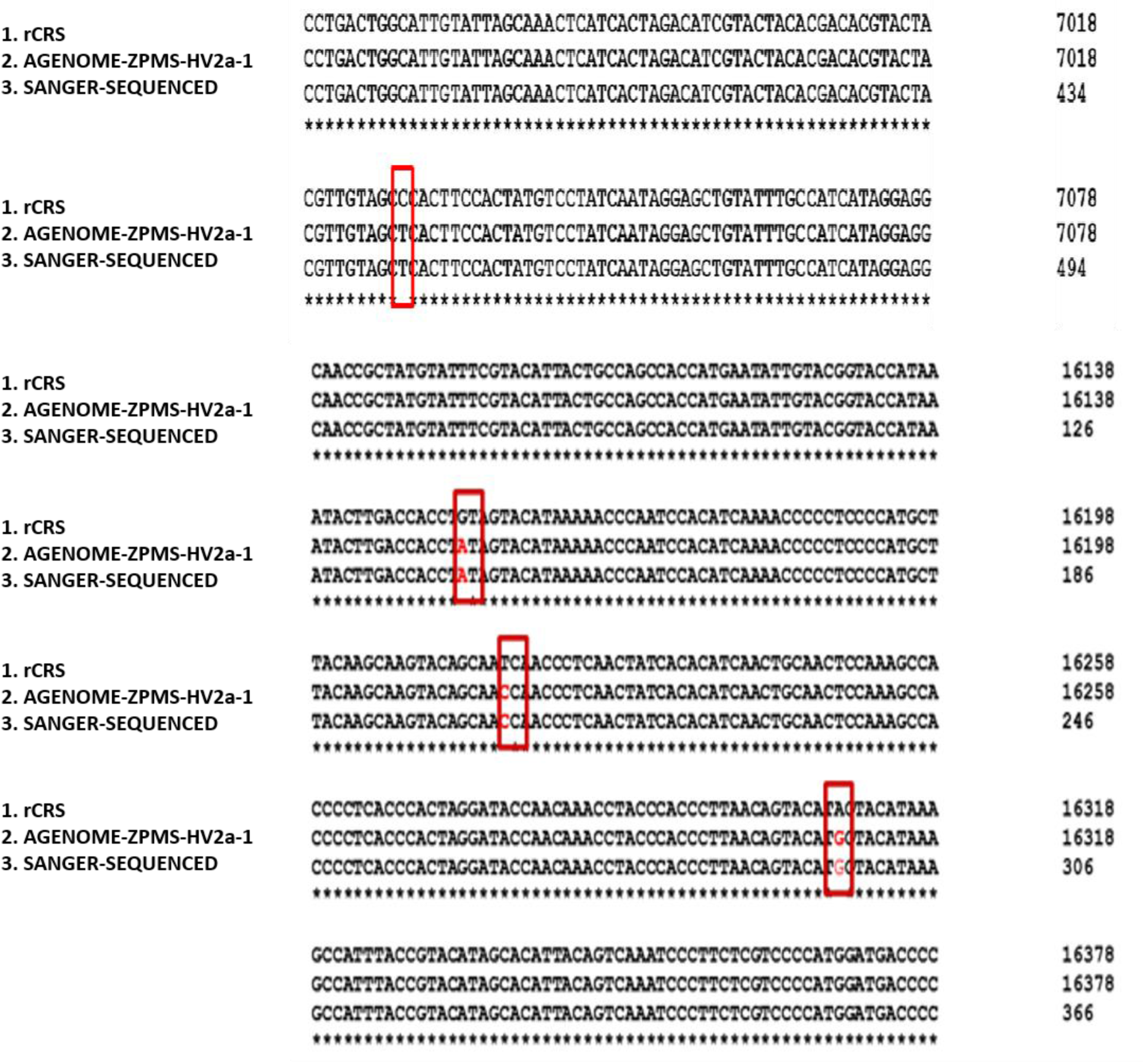
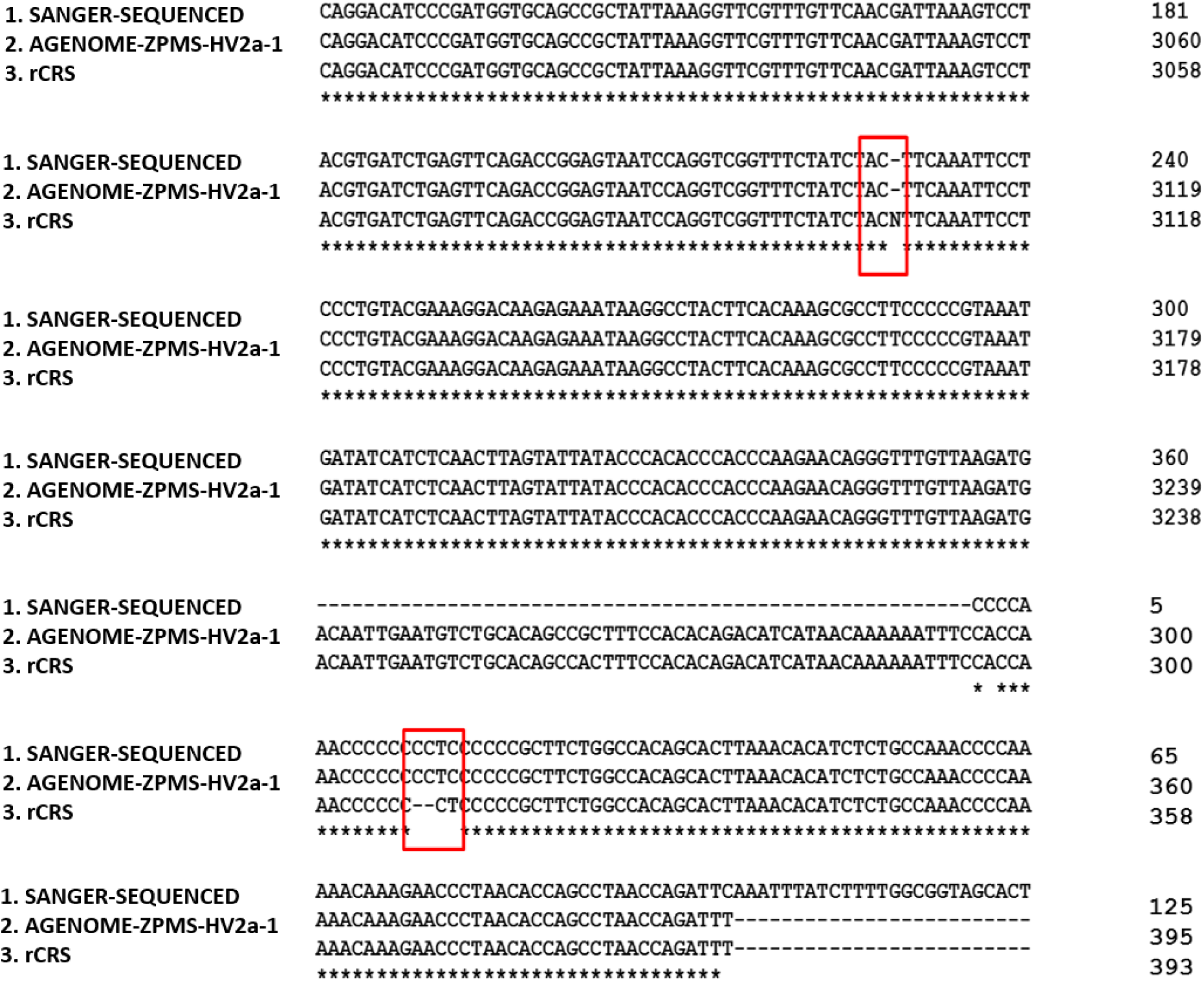
Validation of variants in the AGENOME-ZPMS-HV2a-1 by Sanger sequencing. Confirmation of variants identified with next-generation sequencing (NGS) data and confirmation by Sanger sequencing. Sequences obtained from desired regions were analyzed for presence of variants/Variants. Low quality bases were trimmed from both ends of the sequences and used for alignment with the reference Mitochondrial Genome (rCRS). A total of 13 variants/Variants from D-loop and internal region of mitochondrial genome were verified.

**Supplementary Figure 3:**
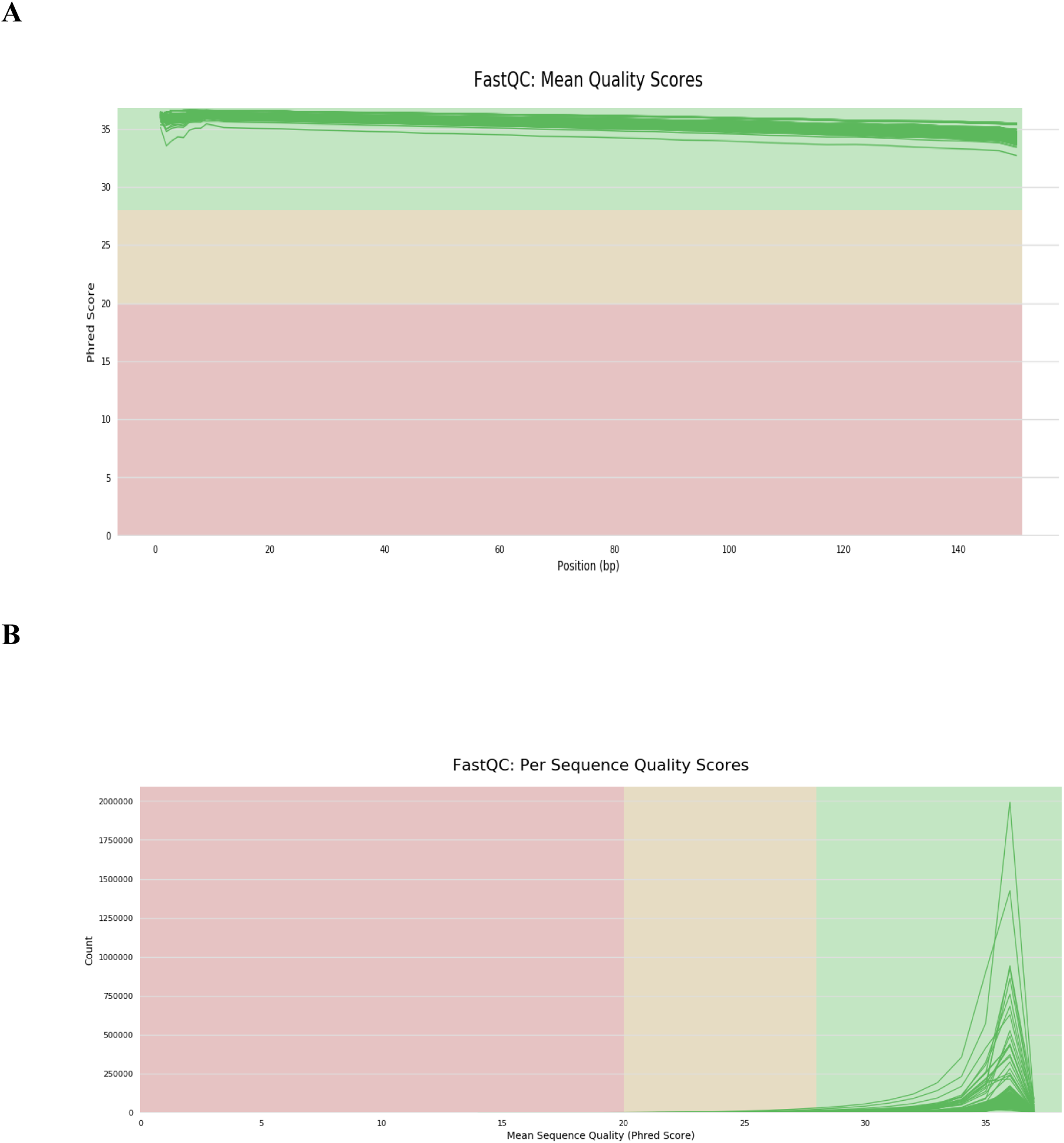
QC analysis of 100 Zoroastrian-Parsi mitochondrial genome sequences. QC analysis of 100 Parsi mitochondrial genomes (A) Frequency of mean PHRED score per read (150 read length) for 100 mitochondrial sample (B) Frequency of mean PHRED score per sequence for 100 mitochondrial samples

**Supplementary Figure 4:**
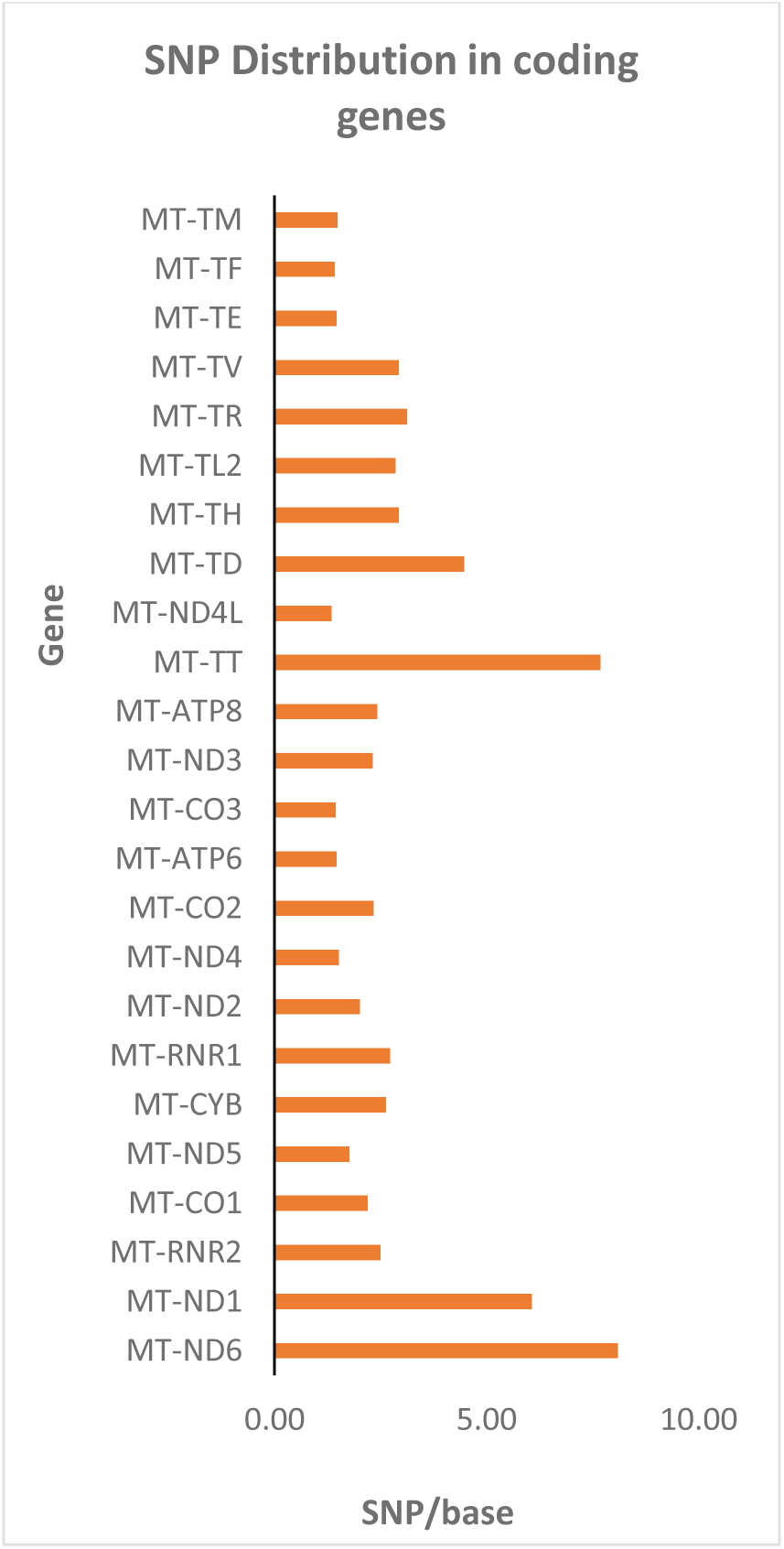
Distribution of 420 variants across coding genes normalized for gene length. Distribution of 420 variants across coding genes normalized to gene length (variants/gene length (in kb)

**Supplementary Figure 5:**
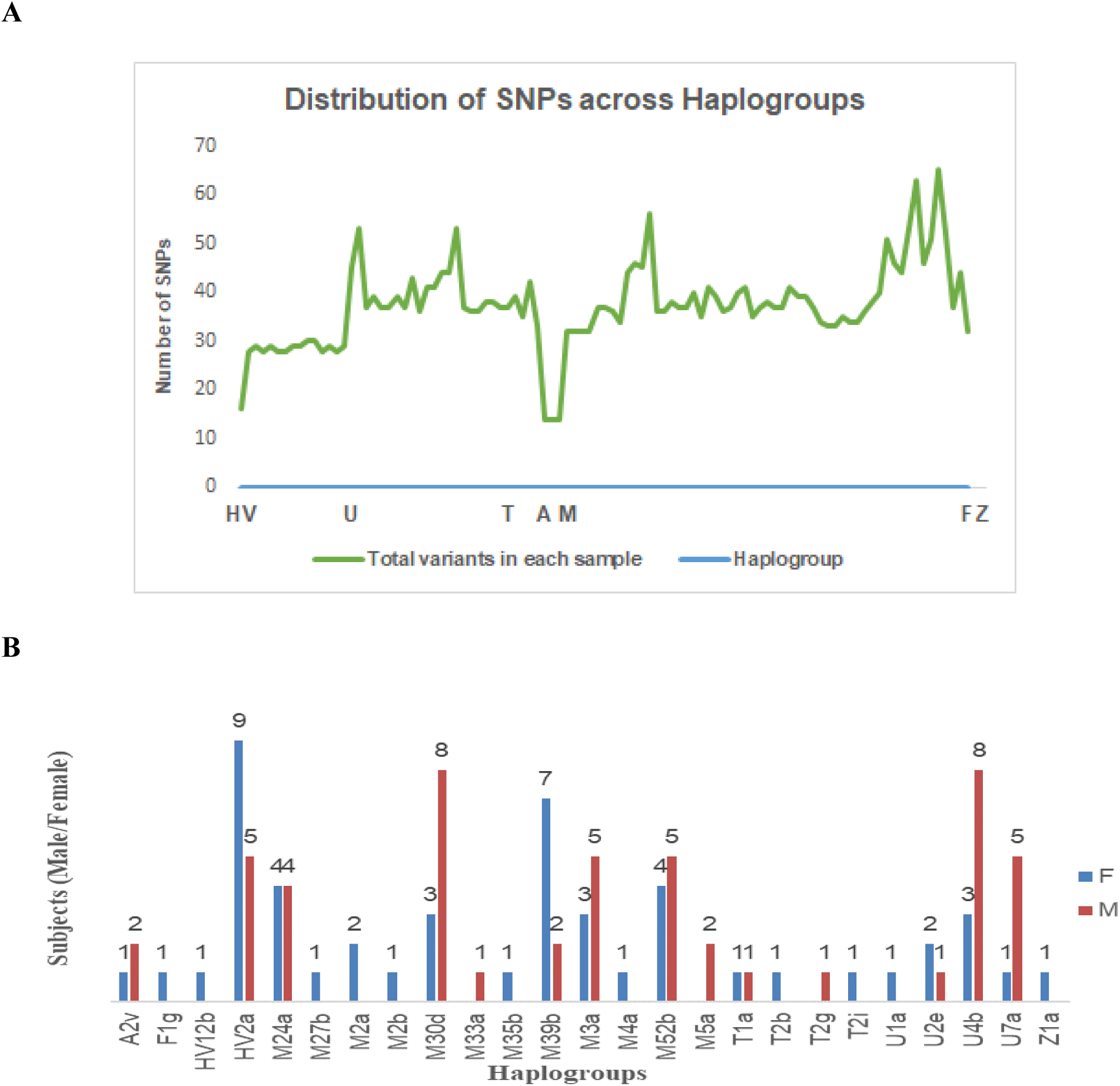
Distribution of variants across haplogroups and demographic classification of the 100 Parsi study group. Distribution across the 100 Zoroastrian-Parsi subjects. (A) Representative graph depicting the distribution of SNP’s count across the 7 major haplogroups (B) Graph depicts the distribution of the subjects classified based on gender across 25 sub-haplogroups

**Supplementary Figure 6: Sub-haplogroup specific breakdown of 420 variants** The sub-haplogroup HV12b (n=1 subject) contained 17 Variants. HVR II harbors four Variants, while the coding genes together contain six Variants that encode three synonymous and three nonsynonymous substitutions. No Variants were observed in the genes coding for tRNAs in the HV12b sub-haplogroup.

In the four U sub-haplogroups analyzed, U1a contained 44 Variants. Two Variants were found in regions coding for tRNA[D] and tRNA[L:CUN].

64 variants were observed for U4b, the most common sub-haplogroup, (n=20) found in the gene encoding 16S-RNR2 (**Supplementary Figure 6B**). Twenty-one Variants were found in coding regions (14 synonymous and 7 nonsynonymous substitutions), with the highest number seen in the gene coding for *COI* (n=6 Variants). Four tRNA mutations were observed in this sub-haplogroup and one mutation in the D-loop region.

A total of 52 variants were observed across all samples in the U7a subgroup (**Supplementary Figure 6B**). Twenty-seven Variants were found in noncoding regions, 12S-RNR1, 16S-RNR2, and the D-loop region. Twenty-five Variants were found in the coding region (17 synonymous and 8 nonsynonymous substitutions), with 17/25 distributed among the *ND* genes coding for *ND1–6*. *ND5* (n=6 Variants) encodes five synonymous mutations, with a nonsynonymous mutation observed at m.14110 T>C (F592L, in 4/6 subjects).

A total of 55 Variants was observed for U2e, with the majority (n=33 Variants) falling in the noncoding regions (HVRI-III and D-loop) and the 12S-RNR1, 16S-RNR2, and tRNA genes. Twenty-two Variants fell within the coding region (15 synonymous and 7 nonsynonymous substitutions), of which 8 fell in the ND gene complex (four *ND2*, four *ND5*) and four in the *CYTB* gene. While all the Variants in the *ND2* and *ND4* genes are synonymous substitutions, all the Variants in the *CYTB* gene encoded nonsynonymous mutations (m.14766 C>T; T7I in 3/3 subjects, m.15326 A>G; T194A in 3/3 subjects; m.14831 G>A; A29T and m.15479 C>T; F245L, both in 1/3 subjects).

Five subjects in our analysis (n=100) fell within the T haplogroup. We found four sub-haplogroups within this haplogroup (T1a, 2 subjects: T2b, T2i, and T2g, with 1 subject each). Our analysis indicated a total of 39 Variants (**Supplementary Figure 6C**) for T1a, with 21/39 Variants found in noncoding regions, including 12S-rRNA, 16S-rRNA, tRNAs, and control regions, including the D-loop. Eighteen Variants were observed in the coding region, with the greatest number occurring in the *CYTB* gene (n=5 Variants). Three Variants within the *CYTB* gene coded for nonsynonymous mutations, including m.14776 C>T, m.14905 G>A, and m.15452 C>A, coding for T7I, T194A, and L236I substitutions, respectively.

The T2b, T2g, and T2i sub-haplogroups contained 35, 42, and 34 Variants, respectively, in total. We found that *CYTB* contained the majority of the Variants found in the coding regions in these sub-haplogroups, except for the T2i group in which the *CYTB* Variants (n=5) constituted the majority of the Variants found in coding and noncoding regions of the genome. Two Variants, m.14766 C>T and m.15326 A>G, seen in all three groups code for nonsynonymous substitutions, and m.15452 C>A was seen in T2g and T2i and codes for a nonsynonymous mutation. Single mutations were seen for m.15497 G>A and m.14798 T>C and code for nonsynonymous substitutions and need further investigation.

The A haplogroup in our study consists of the sub-haplogroup A2v (n=3 subjects). The subjects in the A2v sub-haplogroup had a total of 17 Variants (**Supplementary Figure 6D**) distributed across the mitochondrial genome. Twelve of seventeen Variants were found in the noncoding regions (HVR I, II) and in the 12S rRNA and 16S rRNA genes. Five Variants were distributed in the coding region across *ND2* (m.4769 A>G and m.6095 A>G), *ATPase6* (m.8860 A>G), *ND4* (m.11881 C>T), and *CYTB* (m.15326 A>G). Two nonsynonymous substitutions were observed in the *ATPase6* and *CYTB* genes that need further investigation.

F1g (n=1 subject) is a sub-haplogroup, along with Z1a (n=1 subject). A total of 33 and 32 Variants, respectively, were identified in these groups. Nine *CYTB* Variants were observed in total for both groups. Two encoded nonsynonymous substitutions, m.14766 C>T (T7I) and m.15326 A>G (T194A), while the seven other Variants resulted in synonymous mutations. Variants for *ND4L* are seen only across Z1a and F1g, with the m.10609 T>C SNP in F1g resulting in a nonsynonymous shift (M47T), while the Z1a SNP resulted in a synonymous substitution (**Supplementary Figure 6D**).

The M haplogroup (n=52 subjects) consists of 12 sub-haplogroups, the most number for a haplogroup in our study (**Supplementary Figure 6E**). M30d is the sub-haplogroups with the highest number of subjects in the M haplogroup (n=11 subjects). Fifty-one Variants were identified in this sub-haplogroup in total, of which 28 Variants were seen in the noncoding regions (HVR I, II, III), the D-loop region, and the 12S-RNR1 and 16S-RNR2 genes. The remaining 23 Variants were part of the coding region within *CYTB* (n=8 Variants) and *ND4* (n=5 Variants) and formed a majority. Nine of thirteen Variants in *CYTB* and *ND4* code for synonymous substitutions, while four Variants in *CYTB* resulted in nonsynonymous substitutions (m.14766 C>T; T7I, m.15218 A>G; T158A, m.15326 G>A; T194A, and m.15420 G>A; A229T).

M39b (n=10 subjects) is one of the largest sub-haplogroups, and a total of 59 Variants were seen for this sub-haplolgroup. The noncoding regions, 12S, 16S, and control regions, together constitute 33/59 of the Variants. Of the remaining 26 Variants, the 5 Variants in the *CYTB* complex constitute the greatest number, while the ND gene complex accounts for 12 Variants (2 *ND1*, 1 *ND2*, 2 *ND3*, 2 *ND4*, 3 *ND5*, and 2 *ND6*). Of the nine remaining Variants, six are seen in the *COI*, *II*, and *III* genes (two each), while three Variants are found in the *ATPase6* gene.

The M2 sub-haplogroup consists of M2a (n=2 subjects) and M2b (n=1 subject). A total of 110 Variants was observed in total for M2a and M2b (**Supplementary Figure 6E**). In M2a, 23/53 Variants occurred in noncoding regions (HVR I, II, III), the 12S-RNR1 and 16S-RNR2 genes, the control region (OL), and the D-loop region. Thirty Variants occurred in the coding regions, making this one of the sub-haplogroups in which Variants in the coding region outnumber the Variants in the noncoding region. *CYTB* harbors seven Variants, followed by three Variants in *ND4* and three Variants in *ATPase8*, *ATPase6*, and *COI*. A total of 55 Variants was observed for M2b, in which 31/55 Variants occurred in the noncoding regions. Twenty-four Variants were observed in genes coding for COI, III; *ND1,2,3,4,5; ATPase6,8*; and *CYTB*. The six Variants in *CYTB* constitute the greatest number of Variants in the coding region. The M2a/b sub-haplogroup is also conspicuous by the presence of Variants in the *ATPase8* gene, which is not observed in any sub-haplogroup besides U4b. The complete distribution of the Variants across all the sub-haplogroups is presented in **Table 2**.

The M3a sub-haplogroup (n=8 subjects) consists of 38 variants, with 12/38 variants in the HVR I, II, III, D-loop regions (**Supplementary Figure 6E**). 19/38 variants were observed in the protein coding regions, with the most variants in this region occurring in *CYTB* (n=5). We found 15 coding for synonymous substitutions and 5 for non-synonymous variants (**Supplementary Figure 4E**)

M52b sub haplogroup (n=9 subjects) contained a total of 90 variants. 29/90 variants were observed in HVR I, II, III and the D-loop (**Supplementary Figure 6E**). 31 variants were observed for protein coding genes. *CYTB* (n=9 variants) contains the most variants for this region. 2 variants were found in t-RNA coding genes. 22 variants coded for synonymous substitutions while 9 variants coded for non-synonymous substitutions.

M24a subhaplogroup (n=8 subjects) contains a total of 48 variants, 12/48 are seen in HVR I, II, III and D-loop (Supplementary Figure 6E). 22/48 are found in protein encoding genes with the most on *CYTB* (n=5 variants). 13 synonymous variants and 7 non-synonymous variants are seen in this sub-haplogroup. The rest of the variants are seen in 12S, 16S-rRNA. No variants for t-RNA genes were observed in this sub-haplogroup.

M27b (n=1 subject) has a total of 41 variants (**Supplementary Figure 6E**). 16/41 are seen in HVR I, II, III and the D-loop. 22/41 variants are seen in protein encoding genes with the highest variant count in *CYTB* (n=6 variants). 14 synonymous and 8 non-synonymous variants are observed for this sub-haplogroup and 1 variant for t-RNA coding gene.

M4a (n=1 subject) contains a total of 40 variants. 15/40 variants are seen in the non-coding regions of HVRI, II, III and D-loop (**Supplementary Figure 6E**). 21 variants are seen in the protein coding region with *CYTB* gene (n=5 variants) containing the highest variant count. Like M27b, M4a contains 14 synonymous and 7 non-synonymous variants and 1 variant on the t-RNA coding gene.

A total of 45 variants was seen in M5a sub-haplogroup (n=2 subjects) (**Supplementary Figure 6E**). 19/45 was seen in protein coding genes with *CYTB* (n=7 variants) representing the highest variants in the protein coding region. 13 variants code for synonymous substitutions while 6 code for non-synonymous variants. 1 variant is observed for a t-RNA coding gene.

M35b sub-haplogroup (1 subject) contains a total of 40 variants (**Supplementary Figure 6E**). 15/40 variants are seen in HVR I, II, III and D-loop and 20/40 variants are found in protein encoding regions with the most variants observed in *CYTB* gene (n=5 variants). 14 code for synonymous substitution while 7 code for non-synonymous substitutions. 1 variant is observed for a t-RNA coding gene.

M33a sub-haplogroup (n=1 subject) contains 39 variants (**Supplementary Figure 6E**). 15/39 variants are observed in HVR I, II, III and D-loop, 19/39 variants are seen in the protein coding region, with the highest count seen for *CYTB* (n=5 variants) for this region. 12 are synonymous and 7 are non-synonymous substitutions.1 variant for t-RNA coding gene is also observed in this sub-haplogroup. This haplogroup is unique amongst the 25 sub-haplogroups owing to the presence of a variant (m.8562 C>T) at *ATPase6/8* gene.

**Supplementary Figure 6A:**
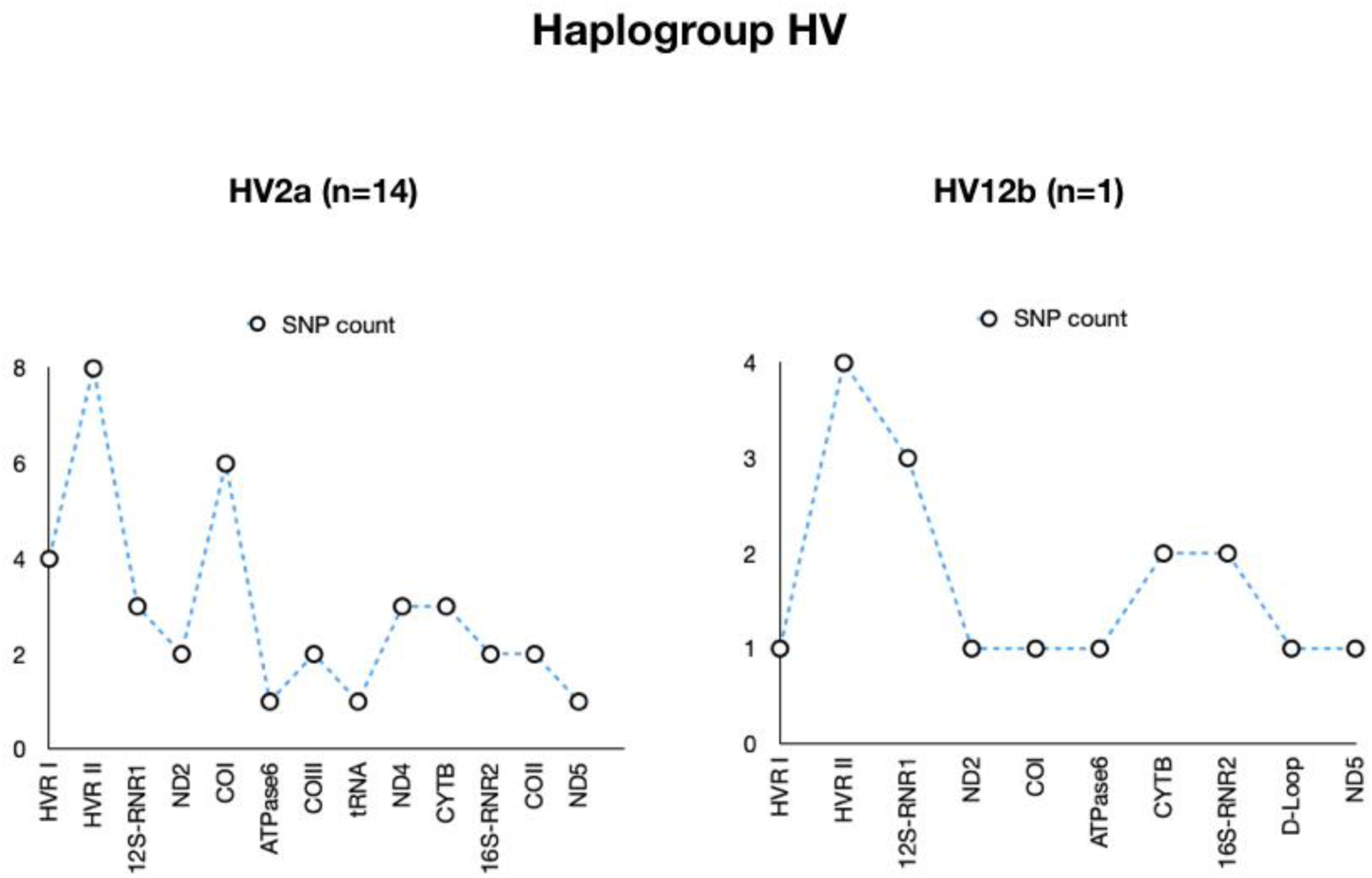
Distribution of Variants across gene loci in the HV haplogroup consisting of HV2a (n=14 subjects and HV12b (n=1 subject)

**Supplementary Figure 6B:**
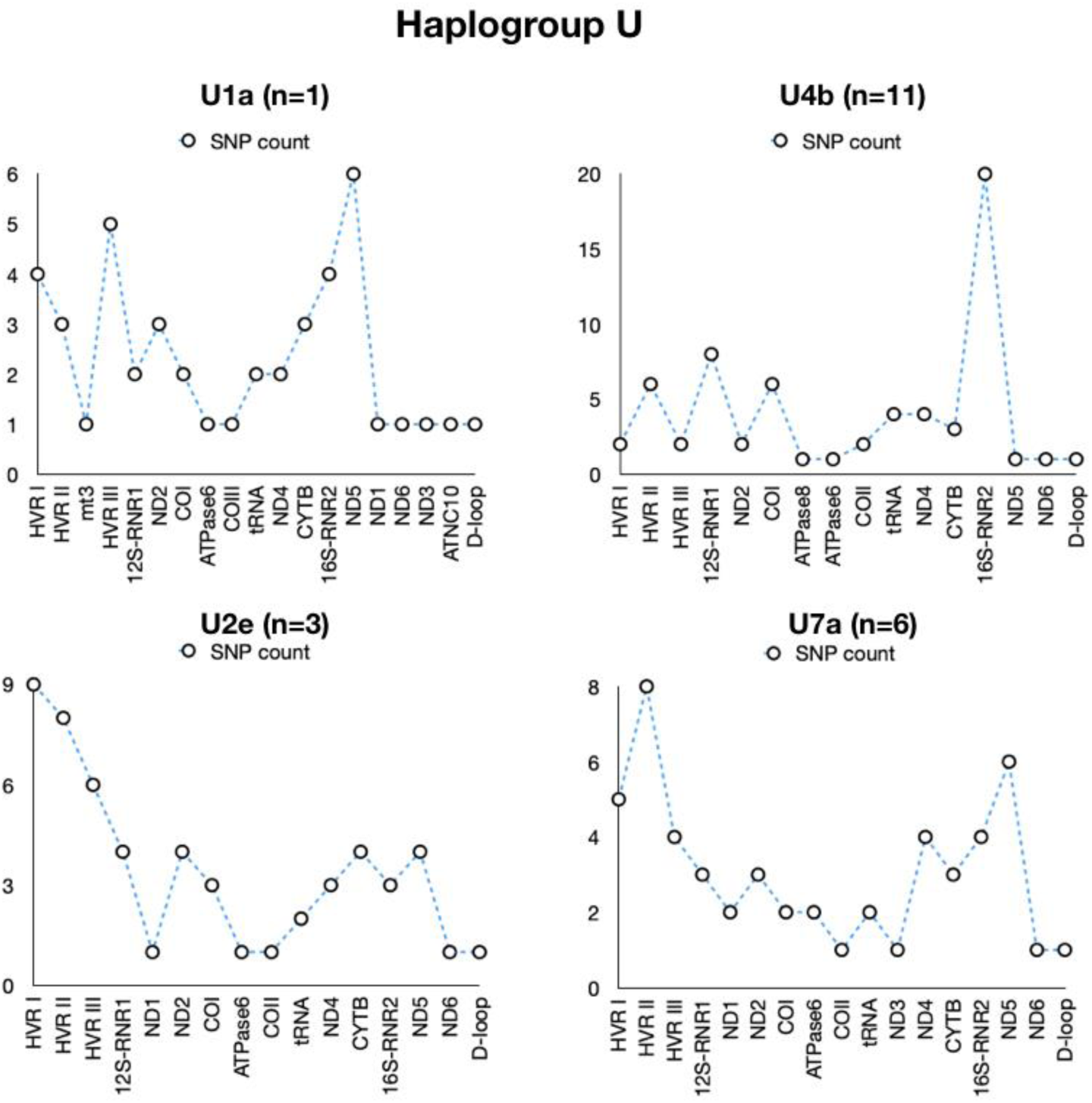
Distribution of Variants across gene loci in the U haplogroup consisting of U1a, U4b, U2e and U7a

**Supplementary Figure 6C:**
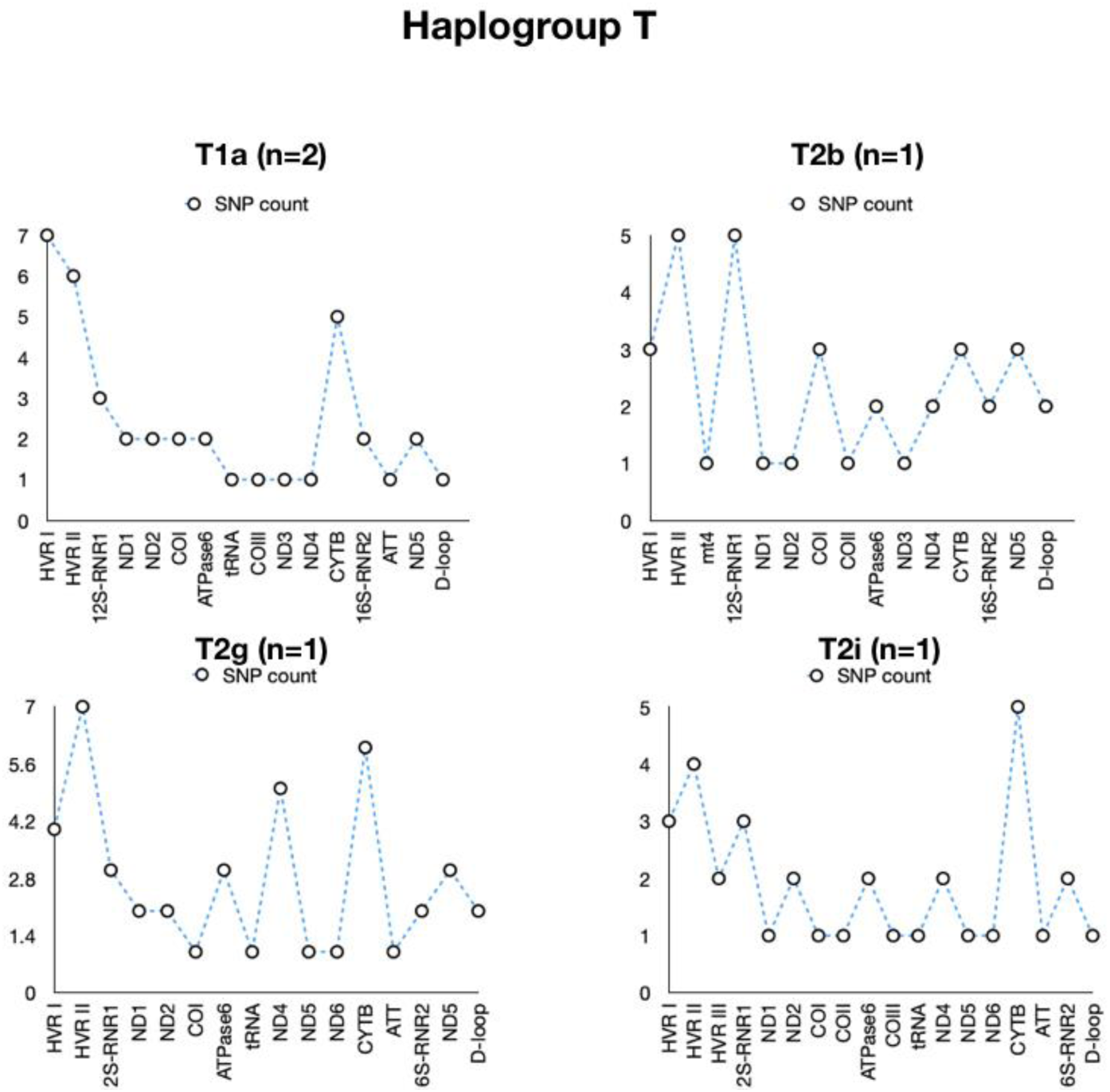
Distribution of Variants across gene loci in the T haplogroup consisting of T1a, T2b, T2g and T2i

**Supplementary Figure 6D:**
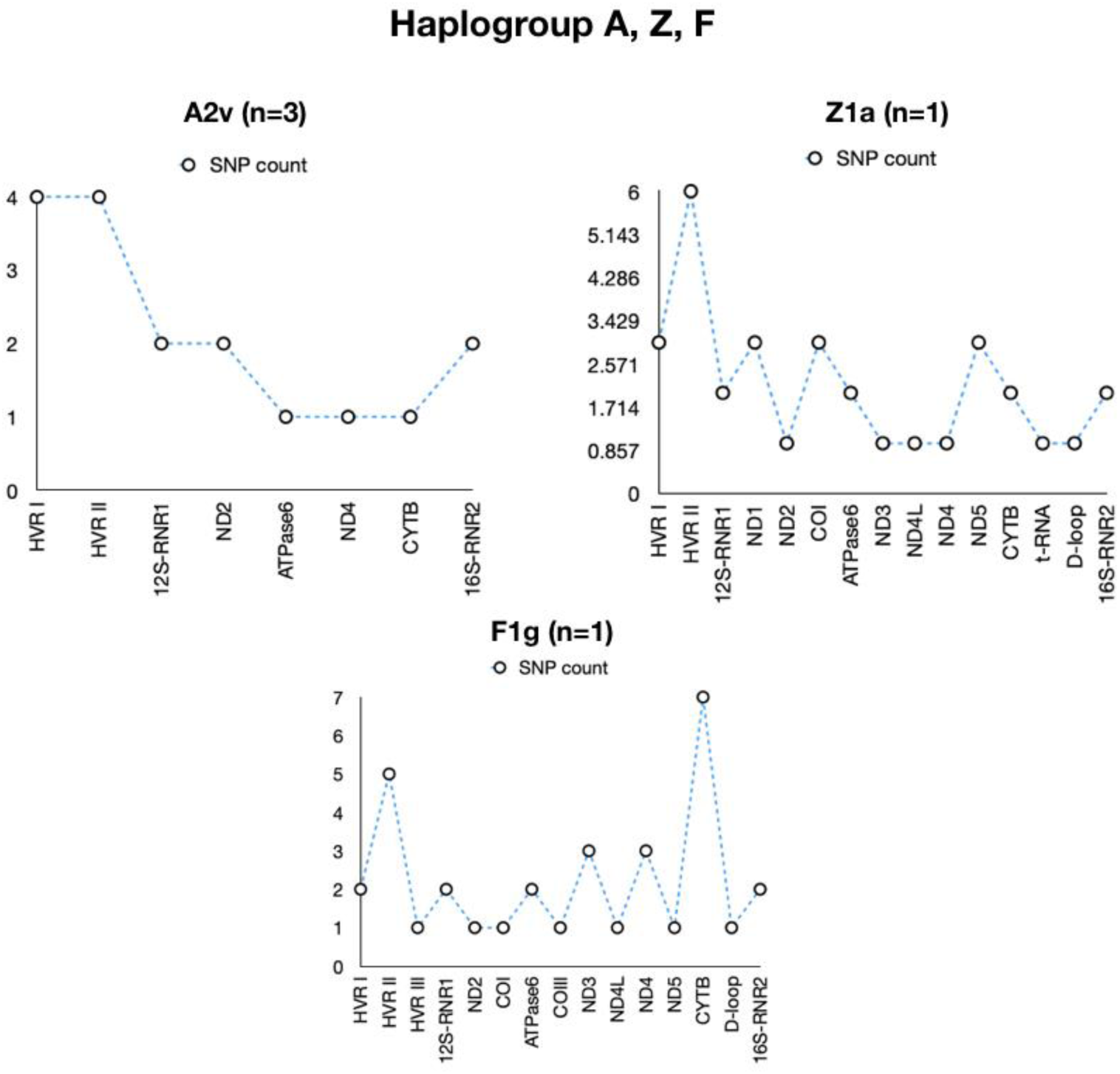
Distribution of Variants across gene loci in the A, Z and F haplogroup consisting of A2v, Z1a and F1g

**Supplementary Figure 6E:**
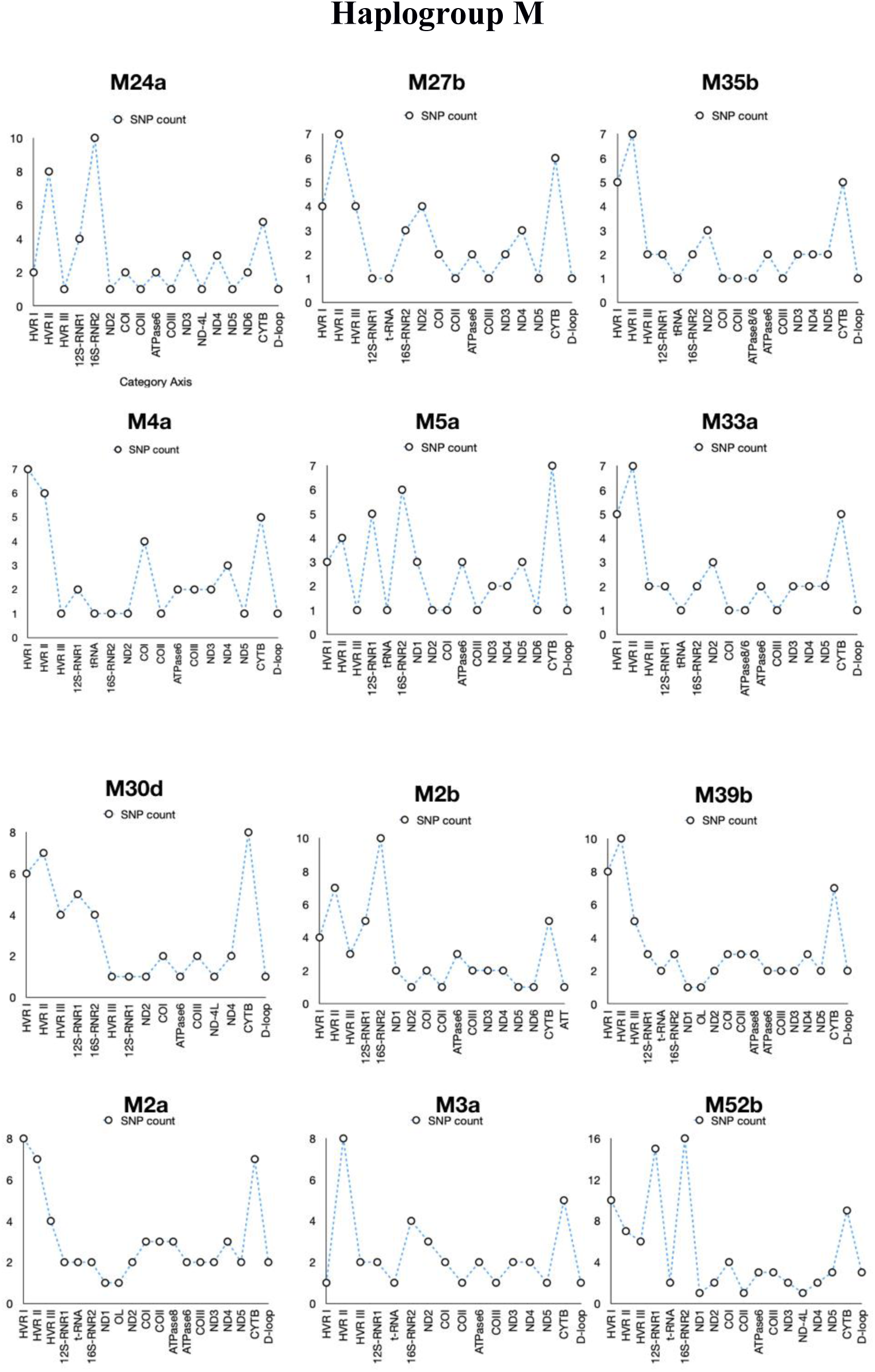
Distribution of variants across gene loci across the M sub-haplogroups

**Supplementary Figure 7:**
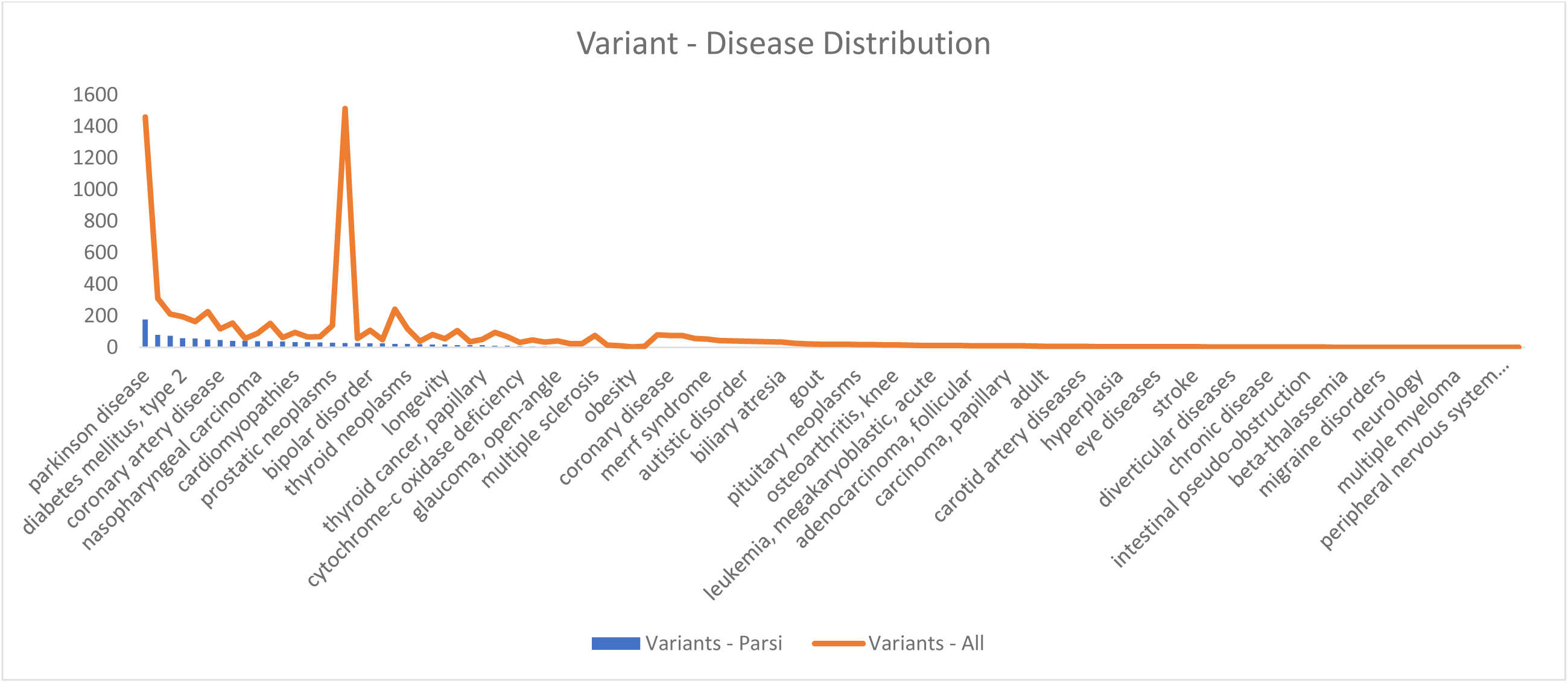
VarDiG^®^ -R analysis of 420 variants indicates high association of Parsi specific variants with Parkinsons diseases. Variant-disease distribution of 420 Parsi variants. Graph depicts the variant-disease distribution between Parsis (blue) and VarDiG^®^-R (Brown)

**Supplementary Figure 8:**
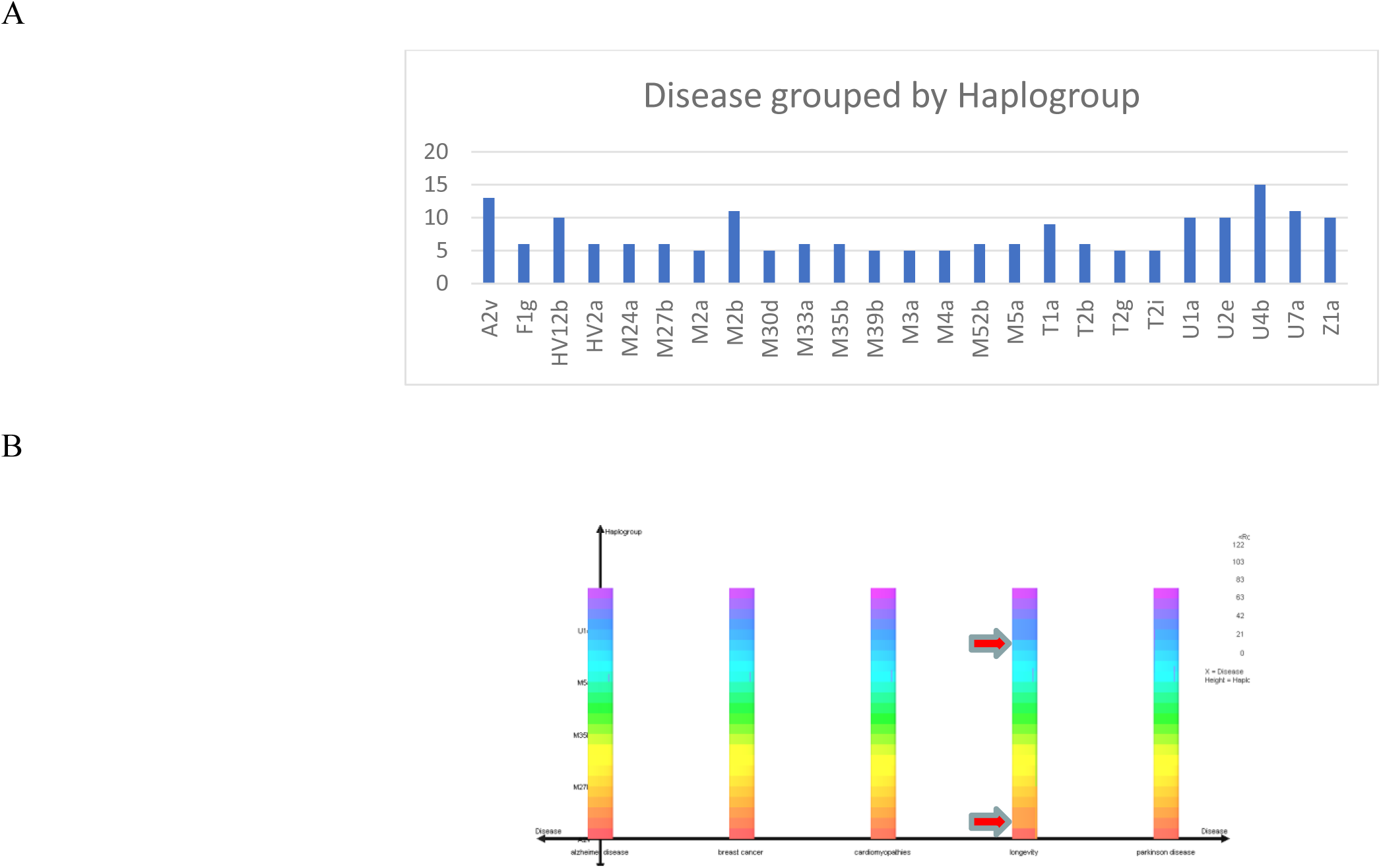
Observation of Longevity variants across all sub-haplogroups and predisposition of U and M haplogroups to diseases. Haplogroup specific distribution of diseases. (A) Distribution of 188 diseases across 25 sub-haplogroups of the 100 Parsi subjects analyzed in this study (B) Histogram depicting longevity and disease prevalence across U1a, M52b, M35b, M27b

**Supplementary Figure 9:**
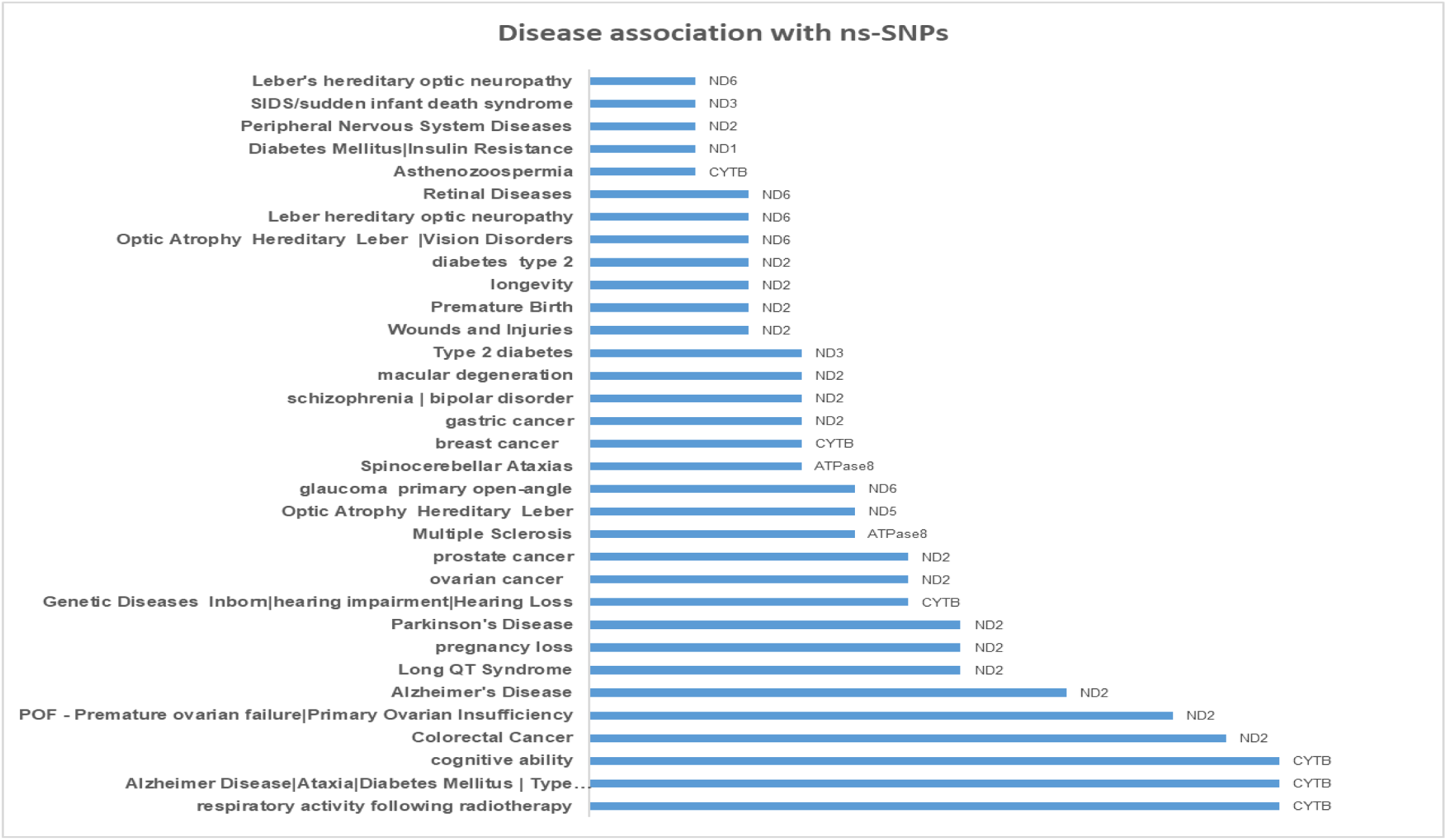
Non-synonymous variants among 420 variants and their disease associations. Analysis of the non-synonymous variants within 420 variants in the 100 Parsi mitochondrial genome sequences for and their disease associations.

**Supplementary Figure 10:**
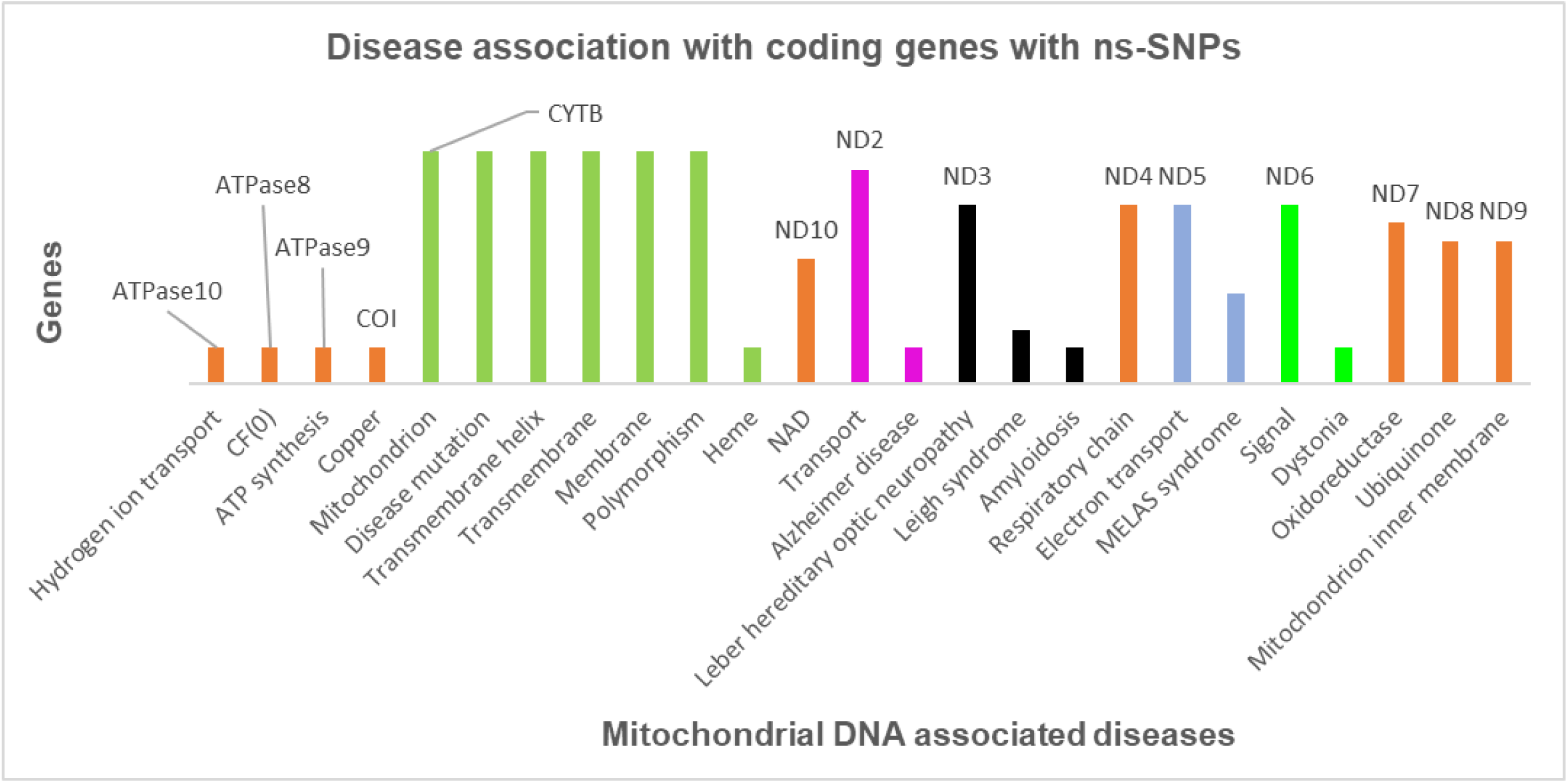
Non-synonymous variants among 420 variants and their associations with mitochondrial function. Distribution of non-synonymous Variants across coding genes. Analysis was performed on the 420 Variants linked to the 100 Parsi mitochondrial genomes.

**Supplementary Table 1:**
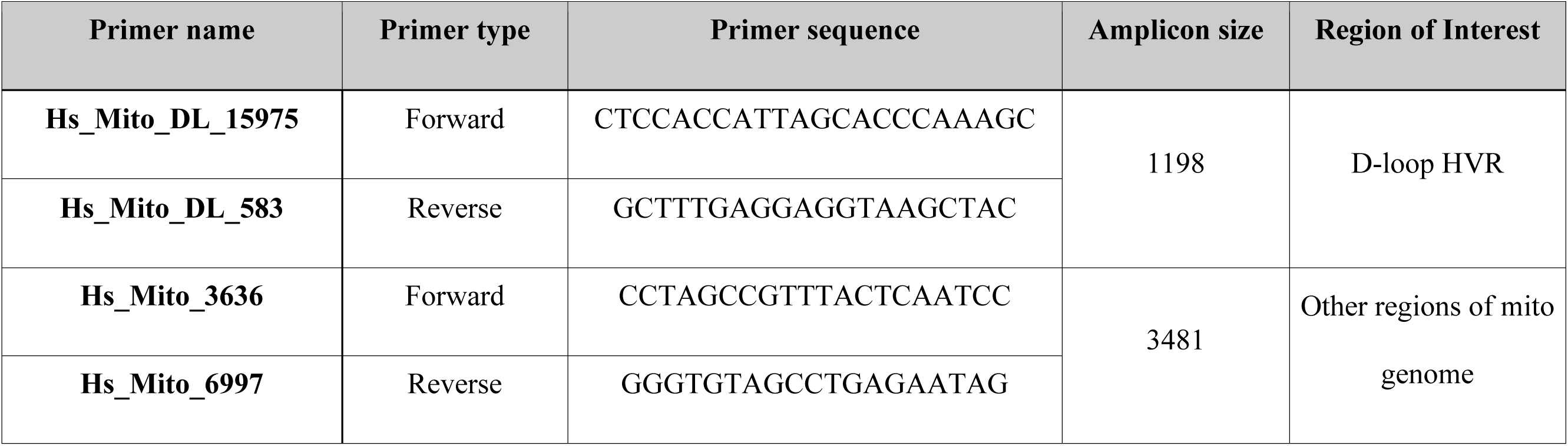
Description of primers used in validation of AGENOME-ZPMS-HV2a-1 by Sanger sequencing. Table shows the list of primers sequences used for Sanger sequencing for validation of selected variants in the AGENOME-ZPMS-HV2a-1

**Supplementary Table 2:**
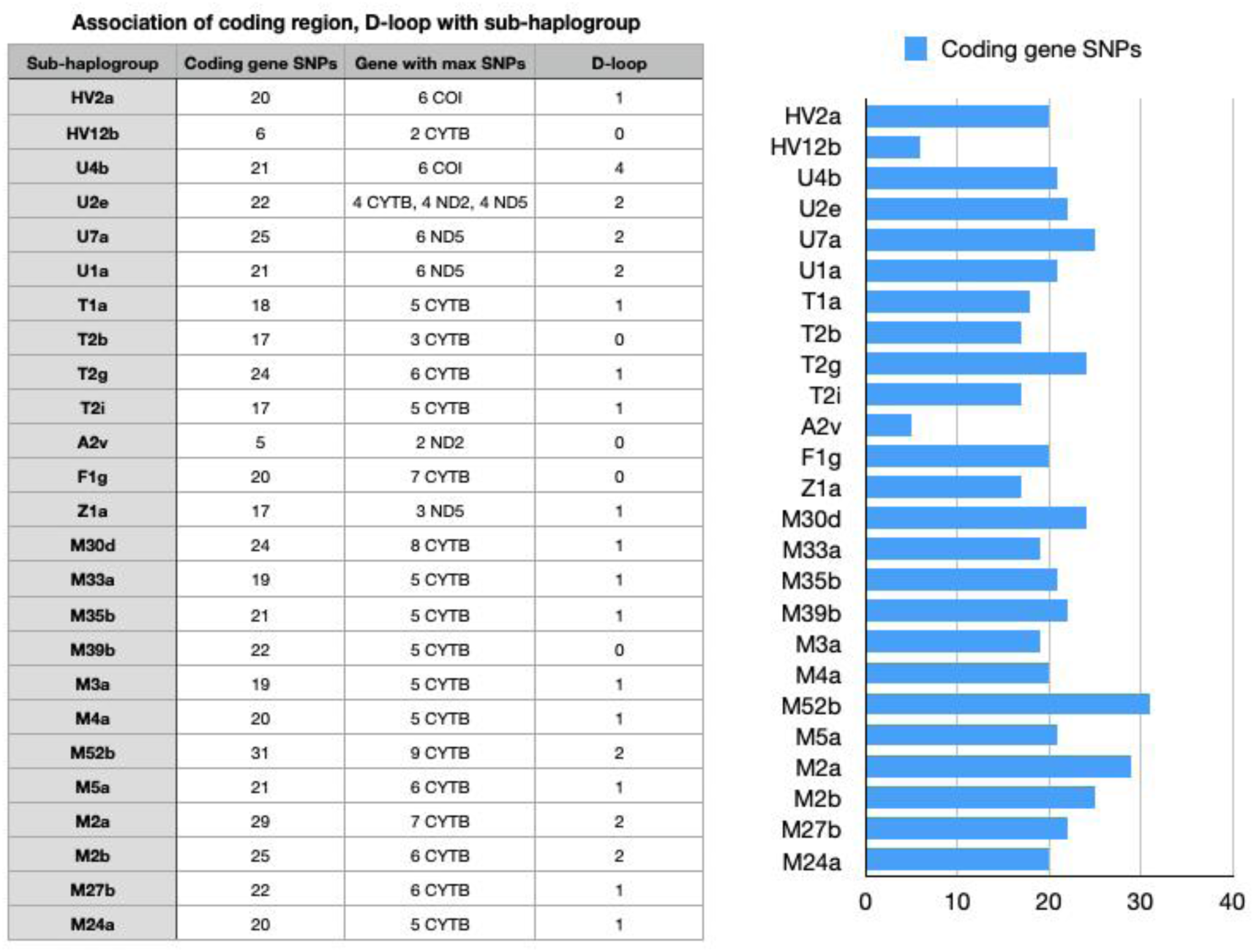
Distribution of 420 variants for each sub-haplogroup for protein coding regions, D-loop of 100 Parsi mitogenomes. Distribution of Variants across coding genes, D-loop across all the 25 sub-haplogroup

**Supplementary Table 3:** Phylogenetic clustering of complete mitogenomes of Parsis with 352 Iranian and 100 relic tribes of Indian origin. Results of the Phylogenetic clustering of the 100 Parsis mitochondrial genomes with 352 mitochondrial genomes of Iranian origin and 100 mitochondrial genomes of relic tribes of Indian origin through Neighbour Joining method. BS indicates Boot-Strap values between each sample. *N.A. indicates *No Association,* indicating a lack of representation of samples in the specific sub-haplogroup

**Table: Clustering of Parsis with population of Persian and Indian descent**
| Major haplogroup | Sub-haplogroups | People of Persian origin (PO) | People of Indian & Relic tribal origin (IO) | Max BS value to nearest PO | Max BS value to nearest IO |
| --- | --- | --- | --- | --- | --- |
| HV | HV2a | Persian | N.A | 0.7270 | 0 |
|  | HV12b | Persian, Qashqai, Mazandarani | N.A | 0.6550 | 0 |
| U | U7a | Persian, Kurd, Tajik | N.A | 0.8980 | 0 |
|  | U2e | Persian, Qashqai, Azeri | N.A | 1.000 | 0 |
|  | U4b | Persian, Khorasani, Qashqai | N.A | 0.5100 | 0 |
|  | U1a | Persian, Armenian | N.A | 0.6850 | 0 |
| T | T1a | Persian | N.A | 0.7320 | 0 |
|  | T2g | Persian | N.A | 0.4880 | 0 |
|  | T2i | Persian | N.A | 0.4480 | 0 |
|  | T2b | Persian | N.A | 0.4320 | 0 |
| M | M5a | Persian | Munda, Mahali | 0.9860 | 0.6270 |
|  | M39b | Unique cluster |  |  |  |
|  | M33a | Azeri | Jenu Kuruba | 0.2250 | 0.0960 |
|  | M52b | Indian Shia Muslim | Mathakur, Dirang Monpa | 0.7950 | 0.1170 |
|  | M24a | Persian, Qashqai | Pauri Bhaiya, Nihal | 0.8560 | 0.0200 |
|  | M3a | Persian | N.A | 0.9380 | 0 |
|  | M30d | Unique cluster | 1 M30d with Brahmin lyengar, Bhovi | 0 | 0.4020 |
|  | M2a | N.A | Lambadi, Hill Kolam, Katkari, Dongri Bhil | 0 | 0.6110 |
|  | M4a | Persian | N.A | 0.8560 | 0 |
|  | M2b | N.R | Korku, Hill Kolam | 0 | 0.9400 |
|  | M35b | Persian | N.A | 0.3860 | 0 |
|  | M27b | Indian Shia Muslim | N.A | 0.4220 | 0 |
| A | A2v | Persian | N.A | 0.4690 | 0 |
| F | F1g | Kurd, Turkmen | N.A | 0.9970 | 0 |
| Z | Z1a | Qashqai, Persian | N.A | 0.2470 | 0 |

**Supplementary Table 4:**
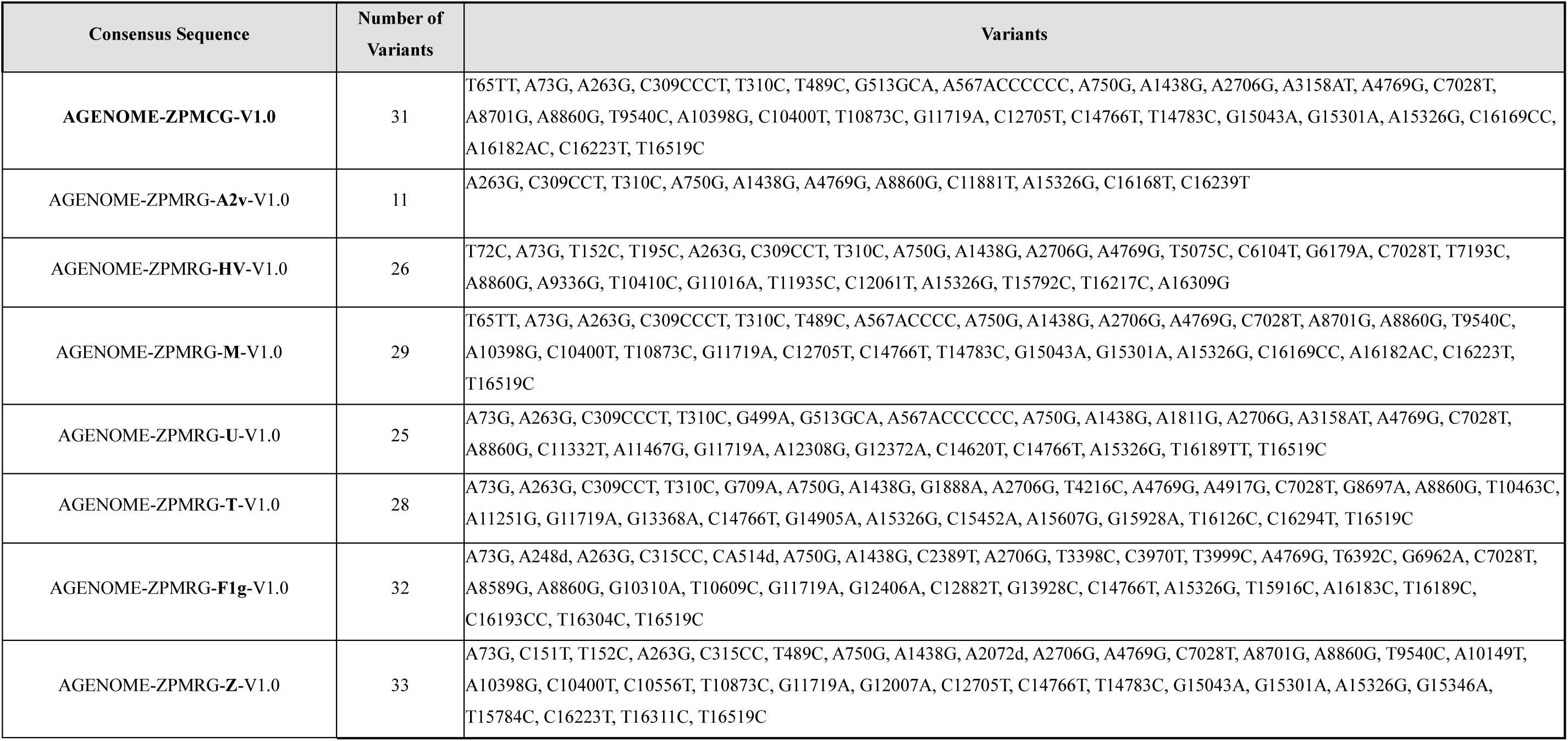
Variants associated with haplogroup specific Zoroastrian Parsi Mitochondrial Reference Genome (n=7) and Zoroastrian Parsi Mitochondrial Consensus Genome (n=1) mitochondrial genome sequences. List of unique variants associated with the Haplogroup specific Zoroastrian Parsi Mitochondrial Reference Genomes (ZPMRG) for A2v, HV, M, U, T, F1g, Z and overall unique variants in the Zoroastrian Parsi Mitochondrial Consensus Genome (ZPMCG)

**Supplementary Table 5:** Variants associated with Zoroastrian Parsi Mitochondrial Reference Genome (ZPMRG) and unique variants of each ZPMRG compared to Zoroastrian Parsi Mitochondrial Consensus Genome (ZPMCG) (A) Unique Variants found in the haplogroup specific Reference Genomes (ZPMRG) compared to the Zoroastrian-Parsi Consensus Genome (AGENOME-ZPMCG-V1). The histogram (right) lists the exact number of variants in each ZPMRG compared to ZPMCG

| AGENOME-ZPMRG-A2v-V1.0 | AGENOME-ZPMRG-HV-V1.0 | AGENOME-ZPMRG-M-V1.0 | AGENOME-ZPMRG-T-V1.0 | AGENOME-ZPMRG-U-V1.0 | AGENOME-ZPMRG-F-V1.0 | AGENOME-ZPMRG-F-V1.0 |
| --- | --- | --- | --- | --- | --- | --- |
| C11881T | A16G |  | C6G | A21G | A248d | C151T |
| C16168T | T72C |  | G709A | G499A | CA514d | A2072d |
| C16239T | T195C |  | G1888A | A1811G | C2389T | C10556T |
|  | T5075C |  | T4216C | C11332T | T3398C | G12007A |
|  | C6104T |  | A4917G | A11467G | C3970T | G15346A |
|  | G6179A |  | G8697A | A12308G | T3999C | T15784C |
|  | T7193C |  | T10463C | G12372A | T6392C | T16311C |
|  | A9336G |  | A11251G | C14620T | G6962A |  |
|  | T10410C |  | G13368A |  | A8589G |  |
|  | G11016A |  | G14905A |  | G10310A |  |
|  | T11935C |  | C15452A |  | T10609C |  |
|  | C12061T |  | A15607G |  | G12406A |  |
|  | T15792C |  | G15928A |  | C12882T |  |
|  | T16217C |  | T16126C |  | G13928C |  |
|  | A16309G |  | C16294T |  | T15916C |  |
|  |  |  |  |  | A16183C |  |
|  |  |  |  |  | C16193CC |  |
|  |  |  |  |  | T16304C |  |

**Supplementary Table 6:** mt-t-RNA variants in our study and their disease association. Analysis of the occurrence of the 420 variants in the tRNA and their disease associations annotated with the PON-mt-tRNA database. A frequency score ≥0.5 – pathogenic, =0.5 – likely pathogenic, <0.5 – neutra

| mt-tRNA | Variation | Probability_of_pathogenicity | Classification | Frequency % | Haplogroup | Disease association |
| --- | --- | --- | --- | --- | --- | --- |
| Phe | T593C | 0.16 | Neutral | 0.06 | M52b | Non-syndromic hearing loss (Reported) |
| Val | G1644A | 0.67 | Pathogenic | 0.01 | U4b | LS/HCM/MELAS (Reported) |
| Val | T1654C | 0.12 | Neutral | 0.01 | M3a |  |
| Met | T4454C | 0.13 | Neutral | 0.02 | M5a | Possible contributor to mito dysfunction / Hypertension (Reported) |
| Asp | G7521A | 0.46 | Likely neutral | 0.01 | U4b |  |
| Asp | T7561C | 0.33 | Neutral | 0.01 | U7a |  |
| Asp | T7581C | 0.42 | Likely neutral | 0.01 | U1a |  |
| Arg | T10410C | 0.17 | Neutral | 0.14 | Hv2a |  |
| Arg | T10463C | 0.31 | Neutral | 0.04 | T1a,T2g,T2i |  |
| His | A12172G | 0.53 | Likely pathogenic | 0.01 | U4b |  |
| His | C12191G | 0.11 | Neutral | 0.01 | M27b |  |
| Leu(CUN) | A12279G | 0.37 | Likely neutral | 0.06 | M52b |  |
| Leu(CUN) | A12308G | 0.41 | Likely neutral | 0.21 | U4b,U7a | Stroke, CM, CPEO, Breast/Renal/Prostate cancer risk, Altered brain pH(Reported) |
| Glu | A14696G | 0.26 | Neutral | 0.01 | A2v | Progressive Encephalopathy (Reported) |
| Thr | A15907G | 0.23 | Neutral | 0.03 | U2e |  |
| Thr | T15908C | 0.5 | Likely pathogenic | 0.01 | M33a | Deaf Helper mutation (Reported) |
| Thr | T15916C | 0.33 | Likely neutral | 0.01 | F1g |  |

## Notes

### Competing Interest Statement

The authors have declared no competing interest.

### Summary of Updates

1) we have implemented an extra QC step to deal numt sequences by implementing RtN pipeline (Updated in "Materials and Methods" section, sub-heading "Retrieving mitochondrial reads from 100 Parsi whole-genome sequences") 2)For the mitochondrial heteroplasmy analysis, we implemented a bioinformatic pipeline to detect heteroplasmies in our sample set using Mutserver run locally ((Updated in "Materials and Methods" section, sub-heading "Retrieving mitochondrial reads from 100 Parsi whole-genome sequences"). The analysis is attached in the "Supplementary Data" as "Appendix 8" 3) In depth haplogroup and haplotype based comparison between the Parsi population with those in the 352 Iranian mitogenome dataset. The results of the analysis is attached in the "Supplementary Data" as "Appendix 7"

## References

1. Wallace, D. C. Mitochondrial DNA Variation in Human Radiation and Disease. Cell (2015) doi:10.1016/j.cell.2015.08.067.

2. Roger, A. J., Muñoz-Gómez, S. A. & Kamikawa, R. The Origin and Diversification of Mitochondria. Current Biology (2017) doi:10.1016/j.cub.2017.09.015.

3. Garcia, I., Jones, E., Ramos, M., Innis-Whitehouse, W. & Gilkerson, R. The little big genome: The organization of mitochondrial DNA. Front. Biosci. - Landmark (2017) doi:10.2741/4511.

4. Wallace, D. C., Brown, M. D. & Lott, M. T. Mitochondrial DNA variation in human evolution and disease. Gene (1999) doi:10.1016/S0378-1119(99)00295-4.

5. Helgason, A., Sigurǒardóttir, S., Gulcher, J. R., Ward, R. & Stefánsson, K. mtDNA and the origin of the Icelanders: Deciphering signals of recent population history. Am. J. Hum. Genet. (2000) doi:10.1086/302816.

6. Thangaraj K, Chaubey G, Kivisild T, Reddy AG, Singh VK, Rasalkar AA, Singh L. Reconstructing the origin of Andaman Islanders. Science. 2005 May 13;308(5724):996. doi: 10.1126/science.1109987. PMID: 15890876.

7. Benton M, Macartney-Coxson D, Eccles D, Griffiths L, Chambers G, et al. Complete Mitochondrial Genome Sequencing Reveals Novel Haplotypes in a Polynesian Population. PLOS ONE 2012, 7(4) e35026. https://doi.org/10.1371/journal.pone.0035026

8. Mistry, R. K. Glimpses of Parsi history, Insights Into The Zarathustrian Religion, p.20.

9. Nariman, R. F. The Inner Fire – Faith, Choice, and Modern Day Living in Zoroastrianism, p. 20–21

10. Anthony, DW, (2007), The Horse, The Wheel, And Language. How Bronze-Age Riders from the Eurasian Steppes Shaped the Modern World, Princeton University Press. p. 9.

11. Alizadeh, A. The Rise of the Highland Elamite State in Southwestern Iran. Current. Curr Anthropol. 51, 353–383 (2010).

12. Shroff Z, C. M. The potential impact of intermarriage on the population decline of the Parsis of Mumbai, India. Demogr Res. 25, 545–564 (2011).

13. Karkal, M. Marriage among Parsis. Demogr. India 4, 128 (1975).

14. The Vendidad: The Zoroastrian Book Of The Law Paperback – September 10, 2010. I, 1-2 & II, 5. Charles. F. Horne. ISBN-10: 1162910089; ISBN-13: 978-1162910086. Kessinger Publishing, LLC (September 10, 2010)

15. Bennet, J. G. The Hyperborean Origin of the Indo-European Culture, Journal Systematics. J Syst. 1, (1963).

16. Jussawalla, D. J., Yeole, B. B. & Natekar, M. V. Histological and epidemiological features of breast cancer in different religious groups in greater bombay. J. Surg. Oncol. (1981) doi:10.1002/jso.2930180309.

17. Barnabas-Sohi, N. et al. Breast carcinoma in a high-risk population: Structural alterations in neu, int-2, and p-53 genes. Breast Dis. (1993).

18. Jussawalla, D. J. The persistance of differences in cancer incidence at various anatomical sites 1300 years after immigration. Recent Results Cancer Res. (1975) doi:10.1007/978-3-642-80880-7_22.

19. Jussawalla, D. J. & Jain, D. K. Lung cancer in Greater Bombay: Correlations with religion and smoking habits. Br. J. Cancer (1979) doi:10.1038/bjc.1979.199.

20. Andrews, R. M. et al. Reanalysis and revision of the cambridge reference sequence for human mitochondrial DNA [5]. Nature Genetics (1999) doi:10.1038/13779.

21. Houshmand, M. et al. Is 8860 variation a rare polymorphism or associated as a secondary effect in HCM disease? Arch. Med. Sci. (2011) doi:10.5114/aoms.2011.22074.

22. Derenko, M. et al. Complete mitochondrial DNA diversity in Iranians. PLoS One (2013) doi:10.1371/journal.pone.0080673.

23. Chandrasekar, A. et al. Updating phylogeny of mitochondrial DNA macrohaplogroup m in India: dispersal of modern human in South Asian corridor. PLoS One 4, e7447–e7447 (2009).

24. Rajkumar, R., Banerjee, J., Gunturi, H. B., Trivedi, R. & Kashyap, V. K. Phylogeny and antiquity of M macrohaplogroup inferred from complete mt DNA sequence of Indian specific lineages. BMC Evol. Biol. 5, 26 (2005).

25. Sahakyan, H. et al. Origin and spread of human mitochondrial DNA haplogroup U7. Sci. Rep. 7, 46044 (2017).

26. Chaubey, G. et al. ‘Like sugar in milk’: Reconstructing the genetic history of the Parsi population. Genome Biol. (2017) doi:10.1186/s13059-017-1244-9.

27. López, S. et al. The Genetic Legacy of Zoroastrianism in Iran and India: Insights into Population Structure, Gene Flow, and Selection. Am. J. Hum. Genet. (2017) doi:10.1016/j.ajhg.2017.07.013.

28. Brandon, M., Baldi, P. & Wallace, D. Mitochondrial mutations in cancer. Oncogene 25, 4647–4662 (2006). https://doi.org/10.1038/sj.onc.1209607

29. Koshikawa N, Akimoto M, Hayashi JI, Nagase H, Takenaga K. Association of predicted pathogenic mutations in mitochondrial ND genes with distant metastasis in NSCLC and colon cancer. Sci Rep. 2017 Nov 14;7(1):15535. doi: 10.1038/s41598-017-15592-2.

30. Alexandrov LB, Ju YS, Haase K, et al. Mutational signatures associated with tobacco smoking in human cancer. Science. 2016;354(6312):618–622.

31. Menotti F, Brega A, Diegoli M, Grasso M, Modena MG, Arbustini E. A novel mtDNA point mutation in tRNA(Val) is associated with hypertrophic cardiomyopathy and MELAS. Ital Heart J. 2004;5(6):460–465.

32. Brandon MC, Ruiz-Pesini E, Mishmar D, et al. MITOMASTER: a bioinformatics tool for the analysis of mitochondrial DNA sequences. Hum Mutat. 2009;30(1):1–6. doi:10.1002/humu.20801.

33. Quintana-Murci, L. et al. Where west meets east: the complex mtDNA landscape of the southwest and Central Asian corridor. Am. J. Hum. Genet. 74, 827–845 (2004).

34. Shamoon-Pour, M., Li, M. & Merriwether, D. A. Rare human mitochondrial HV lineages spread from the Near East and Caucasus during post-LGM and Neolithic expansions. Sci. Rep. 9, 14751 (2019).

35. Farjadian, S. et al. Discordant Patterns of mtDNA and Ethno-Linguistic Variation in 14 Iranian Ethnic Groups. Hum. Hered. 72, 73–84 (2011).

36. Thangaraj, K. et al. In situ origin of deep rooting lineages of mitochondrial Macrohaplogroup ‘M’ in India. BMC Genomics 7, 151 (2006).

37. Bharucha NE, Bharucha EP, Bharucha AE, Bhise AV, Schoenberg BS. Prevalence of Parkinson’s Disease in the Parsi Community of Bombay, India. Arch Neurol. 1988;45(12):1321–1323. doi:10.1001/archneur.1988.00520360039008

38. Fang, H., Shen, L., Chen, T. et al. Cancer type-specific modulation of mitochondrial haplogroups in breast, colorectal and thyroid cancer. BMC Cancer 10, 421 (2010).

39. Van der Walt JM, Dementieva YA, Martin ER, Scott WK, Nicodemus KK, Kroner CC, Welsh-Bohmer KA, Saunders AM, Roses AD, Small GW, Schmechel DE, Murali Doraiswamy P, Gilbert JR, Haines JL, Vance JM, Pericak-Vance MA. Analysis of European mitochondrial haplogroups with Alzheimer disease risk. Neurosci Lett. 2004 Jul 15; 365(1):28–32.

40. van Oven M, Kayser M Hum Mutat. Updated comprehensive phylogenetic tree of global human mitochondrial DNA variation. 2009 Feb; 30(2): E386–94.

41. E. Ruiz-Pesini, A.C. Lapeña, C. Díez, E. Alvarez, J.A. Enríquez, M.J. López-Pérez Seminal quality correlates with mitochondrial functionality. Clin. Chim. Acta., 300 (2000), p. 97 105.

42. Balkrishna Bhika Yeole, AP Kurkure, SH Advani, Sunny Lizzy; An Assessment of Cancer Incidence Patterns in Parsi and Non Parsi Populations, Greater Mumbai. Asian Pacific Journal of Cancer Prevention, Vol 2, 2001; 293–298

43. Chen JB, Yang YH, Lee WC, et al. Sequence-based polymorphisms in the mitochondrial D-loop and potential SNP predictors for chronic dialysis. PLoS One. 2012;7(7):e41125. doi:10.1371/journal.pone.0041125

44. Zaki EA, Freilinger T, Klopstock T, et al. Two common mitochondrial DNA polymorphisms are highly associated with migraine headache and cyclic vomiting syndrome. Cephalalgia. 2009;29(7):719–728. doi:10.1111/j.1468-2982.2008.01793.x

45. Schulmann A, Ryu E, Goncalves V, et al. Novel Complex Interactions between Mitochondrial and Nuclear DNA in Schizophrenia and Bipolar Disorder. Mol Neuropsychiatry. 2019;5(1):13–27. doi:10.1159/000495658

46. August E Woerner, Jennifer Churchill Cihlar, Utpal Smart, Bruce Budowle, Numt identification and removal with RtN!, Bioinformatics, btaa642, https://doi.org/10.1093/bioinformatics/btaa642

47. Sobenin IA, Mitrofanov KY, Zhelankin AV, et al. Quantitative assessment of heteroplasmy of mitochondrial genome: perspectives in diagnostics and methodological pitfalls. Biomed Res Int. 2014;2014:292017. doi:10.1155/2014/292017

48. Edgar, R. C. MUSCLE: multiple sequence alignment with high accuracy and high throughput. Nucleic Acids Res. 32, 1792–1797 (2004).

49. Kumar, S., Stecher, G., Li, M., Knyaz, C. & Tamura, K. MEGA X: Molecular Evolutionary Genetics Analysis across Computing Platforms. Mol. Biol. Evol. 35, 1547–1549 (2018).

50. Tamura, K., Nei, M. & Kumar, S. Prospects for inferring very large phylogenies by using the neighbor-joining method. Proc. Natl. Acad. Sci. U. S. A. 101, 11030–11035 (2004).

51. Niroula A, Vihinen M. PON-mt-tRNA: a multifactorial probability-based method for classification of mitochondrial tRNA variations. Nucleic Acids Res. 2016;44(5):2020–2027. doi:10.1093/nar/gkw046

