## Appendix 2 for "The First Complete Zoroastrian-Parsi Mitochondrial Reference Genome and genetic signatures of an endogamous non-smoking population"

```
Reference sequence (1): AGENOME-ZPMRG-U-V1.0
Identities normalised by aligned length.
Colored by: identity

                             cov    pid     1 [         .         .         .         .         :         .         .         . 80
1 AGENOME-ZPMRG-U-V1.0    100.0% 100.0%       GATCACAGGTCTATCACCCTGTTAACCACTCACGGGAGCTCTCCATGCATTTGGTATTTTCGTC-TGGGGGGTGTGCACG
2 AGENOME-ZPMCG-V1.0      100.0%  99.9%       GATCACAGGTCTATCACCCTATTAACCACTCACGGGAGCTCTCCATGCATTTGGTATTTTCGTCTTGGGGGGTGTGCACG
3 AGENOME-ZPMRG-M-V1.0    100.0%  99.8%       GATCACAGGTCTATCACCCTATTAACCACTCACGGGAGCTCTCCATGCATTTGGTATTTTCGTCTTGGGGGGTGTGCACG
4 AGENOME-ZPMRG-F-V1.0     99.9%  99.8%       GATCACAGGTCTATCACCCTATTAACCACTCACGGGAGCTCTCCATGCATTTGGTATTTTCGTC-TGGGGGGTGTGCACG
5 AGENOME-ZPMRG-Z-V1.0     99.9%  99.7%       GATCACAGGTCTATCACCCTATTAACCACTCACGGGAGCTCTCCATGCATTTGGTATTTTCGTC-TGGGGGGTGTGCACG
6 AGENOME-ZPMRG-T-V1.0     99.9%  99.8%       GATCAGAGGTCTATCACCCTATTAACCACTCACGGGAGCTCTCCATGCATTTGGTATTTTCGTC-TGGGGGGTGTGCACG
7 AGENOME-ZPMRG-HV-V1.0    99.9%  99.8%       GATCACAGGTCTATCGCCCTATTAACCACTCACGGGAGCTCTCCATGCATTTGGTATTTTCGTC-TGGGGGGCGTGCACG
8 AGENOME-ZPMRG-A-V1.0     99.9%  99.8%       GATGACAGGTCTATCACCCTATTAACCACTCACGGGAGCTCTCCATGCATTTGGTATTTTCGTC-TGGGGGGTATGCACG
9 NC_012920.1              99.9%  99.8%       GATCACAGGTCTATCACCCTATTAACCACTCACGGGAGCTCTCCATGCATTTGGTATTTTCGTC-TGGGGGGTATGCACG
  consensus/100%                              GATsAsAGGTCTATCuCCCTuTTAACCACTCACGGGAGCTCTCCATGCATTTGGTATTTTCGTC.TGGGGGGsuTGCACG
  consensus/90%                               GATsAsAGGTCTATCuCCCTuTTAACCACTCACGGGAGCTCTCCATGCATTTGGTATTTTCGTC.TGGGGGGsuTGCACG
  consensus/80%                               GATCACAGGTCTATCACCCTATTAACCACTCACGGGAGCTCTCCATGCATTTGGTATTTTCGTC.TGGGGGGTuTGCACG
  consensus/70%                               GATCACAGGTCTATCACCCTATTAACCACTCACGGGAGCTCTCCATGCATTTGGTATTTTCGTC.TGGGGGGTGTGCACG

                             cov    pid    81         .         1         .         .         .         .         :         . 160
1 AGENOME-ZPMRG-U-V1.0    100.0% 100.0%       CGATAGCATTGCGAGACGCTGGAGCCGGAGCACCCTATGTCGCAGTATCTGTCTTTGATTCCTGCCTCATCCTATTATTT
2 AGENOME-ZPMCG-V1.0      100.0%  99.9%       CGATAGCATTGCGAGACGCTGGAGCCGGAGCACCCTATGTCGCAGTATCTGTCTTTGATTCCTGCCTCATCCTATTATTT
3 AGENOME-ZPMRG-M-V1.0    100.0%  99.8%       CGATAGCATTGCGAGACGCTGGAGCCGGAGCACCCTATGTCGCAGTATCTGTCTTTGATTCCTGCCTCATCCTATTATTT
4 AGENOME-ZPMRG-F-V1.0     99.9%  99.8%       CGATAGCATTGCGAGACGCTGGAGCCGGAGCACCCTATGTCGCAGTATCTGTCTTTGATTCCTGCCTCATCCTATTATTT
5 AGENOME-ZPMRG-Z-V1.0     99.9%  99.7%       CGATAGCATTGCGAGACGCTGGAGCCGGAGCACCCTATGTCGCAGTATCTGTCTTTGATTCCTGCCTCATCTCATTATTT
6 AGENOME-ZPMRG-T-V1.0     99.9%  99.8%       CGATAGCATTGCGAGACGCTGGAGCCGGAGCACCCTATGTCGCAGTATCTGTCTTTGATTCCTGCCTCATCCTATTATTT
7 AGENOME-ZPMRG-HV-V1.0    99.9%  99.8%       CGATAGCATTGCGAGACGCTGGAGCCGGAGCACCCTATGTCGCAGTATCTGTCTTTGATTCCTGCCTCATCCCATTATTT
8 AGENOME-ZPMRG-A-V1.0     99.9%  99.8%       CGATAGCATTGCGAGACGCTGGAGCCGGAGCACCCTATGTCGCAGTATCTGTCTTTGATTCCTGCCTCATCCTATTATTT
9 NC_012920.1              99.9%  99.8%       CGATAGCATTGCGAGACGCTGGAGCCGGAGCACCCTATGTCGCAGTATCTGTCTTTGATTCCTGCCTCATCCTATTATTT
  consensus/100%                              CGATAGCATTGCGAGACGCTGGAGCCGGAGCACCCTATGTCGCAGTATCTGTCTTTGATTCCTGCCTCATCssATTATTT
  consensus/90%                               CGATAGCATTGCGAGACGCTGGAGCCGGAGCACCCTATGTCGCAGTATCTGTCTTTGATTCCTGCCTCATCssATTATTT
  consensus/80%                               CGATAGCATTGCGAGACGCTGGAGCCGGAGCACCCTATGTCGCAGTATCTGTCTTTGATTCCTGCCTCATCCsATTATTT
  consensus/70%                               CGATAGCATTGCGAGACGCTGGAGCCGGAGCACCCTATGTCGCAGTATCTGTCTTTGATTCCTGCCTCATCCTATTATTT

                             cov    pid   161         .         .         .         2         .         .         .         . 240
1 AGENOME-ZPMRG-U-V1.0    100.0% 100.0%       ATCGCACCTACGTTCAATATTACAGGCGAACATACTTACTAAAGTGTGTTAATTAATTAATGCTTGTAGGACATAATAAT
2 AGENOME-ZPMCG-V1.0      100.0%  99.9%       ATCGCACCTACGTTCAATATTACAGGCGAACATACTTACTAAAGTGTGTTAATTAATTAATGCTTGTAGGACATAATAAT
3 AGENOME-ZPMRG-M-V1.0    100.0%  99.8%       ATCGCACCTACGTTCAATATTACAGGCGAACATACTTACTAAAGTGTGTTAATTAATTAATGCTTGTAGGACATAATAAT
4 AGENOME-ZPMRG-F-V1.0     99.9%  99.8%       ATCGCACCTACGTTCAATATTACAGGCGAACATACTTACTAAAGTGTGTTAATTAATTAATGCTTGTAGGACATAATAAT
5 AGENOME-ZPMRG-Z-V1.0     99.9%  99.7%       ATCGCACCTACGTTCAATATTACAGGCGAACATACTTACTAAAGTGTGTTAATTAATTAATGCTTGTAGGACATAATAAT
6 AGENOME-ZPMRG-T-V1.0     99.9%  99.8%       ATCGCACCTACGTTCAATATTACAGGCGAACATACTTACTAAAGTGTGTTAATTAATTAATGCTTGTAGGACATAATAAT
7 AGENOME-ZPMRG-HV-V1.0    99.9%  99.8%       ATCGCACCTACGTTCAATATTACAGGCGAACATACCTACTAAAGTGTGTTAATTAATTAATGCTTGTAGGACATAATAAT
8 AGENOME-ZPMRG-A-V1.0     99.9%  99.8%       ATCGCACCTACGTTCAATATTACAGGCGAACATACTTACTAAAGTGTGTTAATTAATTAATGCTTGTAGGACATAATAAT
9 NC_012920.1              99.9%  99.8%       ATCGCACCTACGTTCAATATTACAGGCGAACATACTTACTAAAGTGTGTTAATTAATTAATGCTTGTAGGACATAATAAT
  consensus/100%                              ATCGCACCTACGTTCAATATTACAGGCGAACATACsTACTAAAGTGTGTTAATTAATTAATGCTTGTAGGACATAATAAT
  consensus/90%                               ATCGCACCTACGTTCAATATTACAGGCGAACATACsTACTAAAGTGTGTTAATTAATTAATGCTTGTAGGACATAATAAT
  consensus/80%                               ATCGCACCTACGTTCAATATTACAGGCGAACATACTTACTAAAGTGTGTTAATTAATTAATGCTTGTAGGACATAATAAT
  consensus/70%                               ATCGCACCTACGTTCAATATTACAGGCGAACATACTTACTAAAGTGTGTTAATTAATTAATGCTTGTAGGACATAATAAT

                             cov    pid   241         :         .         .         .         .         3         .         . 320
1 AGENOME-ZPMRG-U-V1.0    100.0% 100.0%       AACAATTGAATGTCTGCACAGCCGCTTTCCACACAGACATCATAACAAAAAATTTCCACCAAACCCCCCCCTCCCCCCG
2 AGENOME-ZPMCG-V1.0      100.0%  99.9%       AACAATTGAATGTCTGCACAGCCGCTTTCCACACAGACATCATAACAAAAAATTTCCACCAAACCCCCCCCTCCCCCCG
3 AGENOME-ZPMRG-M-V1.0    100.0%  99.8%       AACAATTGAATGTCTGCACAGCCGCTTTCCACACAGACATCATAACAAAAAATTTCCACCAAACCCCCCCCTCCCCCCG
4 AGENOME-ZPMRG-F-V1.0     99.9%  99.8%       AACAATTG-ATGTCTGCACAGCCGCTTTCCACACAGACATCATAACAAAAAATTTCCACCAAACCCCCC--TCCCCCCG
5 AGENOME-ZPMRG-Z-V1.0     99.9%  99.7%       AACAATTGAATGTCTGCACAGCCGCTTTCCACACAGACATCATAACAAAAAATTTCCACCAAACCCCCC--TCCCCCCG
6 AGENOME-ZPMRG-T-V1.0     99.9%  99.8%       AACAATTGAATGTCTGCACAGCCGCTTTCCACACAGACATCATAACAAAAAATTTCCACCAAACCCCCC-CTCCCCCCG
7 AGENOME-ZPMRG-HV-V1.0    99.9%  99.8%       AACAATTGAATGTCTGCACAGCCGCTTTCCACACAGACATCATAACAAAAAATTTCCACCAAACCCCCC-CTCCCCCCG
8 AGENOME-ZPMRG-A-V1.0     99.9%  99.8%       AACAATTGAATGTCTGCACAGCCGCTTTCCACACAGACATCATAACAAAAAATTTCCACCAAACCCCCC-CTCCCCCCG
9 NC_012920.1              99.9%  99.8%       AACAATTGAATGTCTGCACAGCCACTTTCCACACAGACATCATAACAAAAAATTTCCACCAAACCCCCC---TCCCCCG
  consensus/100%                              AACAATTG.ATGTCTGCACAGCCuCTTTCCACACAGACATCATAACAAAAAATTTCCACCAAACCCCCC...sCCCCCG
  consensus/90%                               AACAATTG.ATGTCTGCACAGCCuCTTTCCACACAGACATCATAACAAAAAATTTCCACCAAACCCCCC...sCCCCCG
  consensus/80%                               AACAATTGAATGTCTGCACAGCCGCTTTCCACACAGACATCATAACAAAAAATTTCCACCAAACCCCCC..TCCCCCCG
  consensus/70%                               AACAATTGAATGTCTGCACAGCCGCTTTCCACACAGACATCATAACAAAAAATTTCCACCAAACCCCCC..TCCCCCCG

                             cov    pid   321         .         .         :         .         .         .         .         4 400
1 AGENOME-ZPMRG-U-V1.0    100.0% 100.0%       CTTCTGGCCACAGCACTTAAACACATCTCTGCCAAACCCCAAAAACAAAGAACCCTAACACCAGCCTAACCAGATTTCAA
2 AGENOME-ZPMCG-V1.0      100.0%  99.9%       CTTCTGGCCACAGCACTTAAACACATCTCTGCCAAACCCCAAAAACAAAGAACCCTAACACCAGCCTAACCAGATTTCAA
3 AGENOME-ZPMRG-M-V1.0    100.0%  99.8%       CTTCTGGCCACAGCACTTAAACACATCTCTGCCAAACCCCAAAAACAAAGAACCCTAACACCAGCCTAACCAGATTTCAA
4 AGENOME-ZPMRG-F-V1.0     99.9%  99.8%       CTTCTGGCCACAGCACTTAAACACATCTCTGCCAAACCCCAAAAACAAAGAACCCTAACACCAGCCTAACCAGATTTCAA
5 AGENOME-ZPMRG-Z-V1.0     99.9%  99.7%       CTTCTGGCCACAGCACTTAAACACATCTCTGCCAAACCCCAAAAACAAAGAACCCTAACACCAGCCTAACCAGATTTCAA
6 AGENOME-ZPMRG-T-V1.0     99.9%  99.8%       CTTCTGGCCACAGCACTTAAACACATCTCTGCCAAACCCCAAAAACAAAGAACCCTAACACCAGCCTAACCAGATTTCAA
7 AGENOME-ZPMRG-HV-V1.0    99.9%  99.8%       CTTCTGGCCACAGCACTTAAACACATCTCTGCCAAACCCCAAAAACAAAGAACCCTAACACCAGCCTAACCAGATTTCAA
8 AGENOME-ZPMRG-A-V1.0     99.9%  99.8%       CTTCTGGCCACAGCACTTAAACACATCTCTGCCAAACCCCAAAAACAAAGAACCCTAACACCAGCCTAACCAGATTTCAA
9 NC_012920.1              99.9%  99.8%       CTTCTGGCCACAGCACTTAAACACATCTCTGCCAAACCCCAAAAACAAAGAACCCTAACACCAGCCTAACCAGATTTCAA
  consensus/100%                              CTTCTGGCCACAGCACTTAAACACATCTCTGCCAAACCCCAAAAACAAAGAACCCTAACACCAGCCTAACCAGATTTCAA
  consensus/90%                               CTTCTGGCCACAGCACTTAAACACATCTCTGCCAAACCCCAAAAACAAAGAACCCTAACACCAGCCTAACCAGATTTCAA
  consensus/80%                               CTTCTGGCCACAGCACTTAAACACATCTCTGCCAAACCCCAAAAACAAAGAACCCTAACACCAGCCTAACCAGATTTCAA
  consensus/70%                               CTTCTGGCCACAGCACTTAAACACATCTCTGCCAAACCCCAAAAACAAAGAACCCTAACACCAGCCTAACCAGATTTCAA

                             cov    pid   401         .         .         .         .         :         .         .         . 480
1 AGENOME-ZPMRG-U-V1.0    100.0% 100.0%       ATTTTATCTTTTGGCGGTATGCACTTTTAACAGTCACCCCCCAACTAACACATTATTTTCCCCTCCCACTCCCATACTAC
2 AGENOME-ZPMCG-V1.0      100.0%  99.9%       ATTTTATCTTTTGGCGGTATGCACTTTTAACAGTCACCCCCCAACTAACACATTATTTTCCCCTCCCACTCCCATACTAC
3 AGENOME-ZPMRG-M-V1.0    100.0%  99.8%       ATTTTATCTTTTGGCGGTATGCACTTTTAACAGTCACCCCCCAACTAACACATTATTTTCCCCTCCCACTCCCATACTAC
4 AGENOME-ZPMRG-F-V1.0     99.9%  99.8%       ATTTTATCTTTTGGCGGTATGCACTTTTAACAGTCACCCCCCAACTAACACATTATTTTCCCCTCCCACTCCCATACTAC
5 AGENOME-ZPMRG-Z-V1.0     99.9%  99.7%       ATTTTATCTTTTGGCGGTATGCACTTTTAACAGTCACCCCCCAACTAACACATTATTTTCCCCTCCCACTCCCATACTAC
6 AGENOME-ZPMRG-T-V1.0     99.9%  99.8%       ATTTTATCTTTTGGCGGTATGCACTTTTAACAGTCACCCCCCAACTAACACATTATTTTCCCCTCCCACTCCCATACTAC
7 AGENOME-ZPMRG-HV-V1.0    99.9%  99.8%       ATTTTATCTTTTGGCGGTATGCACTTTTAACAGTCACCCCCCAACTAACACATTATTTTCCCCTCCCACTCCCATACTAC
8 AGENOME-ZPMRG-A-V1.0     99.9%  99.8%       ATTTTATCTTTTGGCGGTATGCACTTTTAACAGTCACCCCCCAACTAACACATTATTTTCCCCTCCCACTCCCATACTAC
9 NC_012920.1              99.9%  99.8%       ATTTTATCTTTTGGCGGTATGCACTTTTAACAGTCACCCCCCAACTAACACATTATTTTCCCCTCCCACTCCCATACTAC
```

```
   consensus/100%                     ATTTTATCTTTTGGCGGTATGCACTTTTAACAGTCACCCCCCAACTAACACATTATTTTCCCCTCCCACTCCCATACTAC
   consensus/90%                      ATTTTATCTTTTGGCGGTATGCACTTTTAACAGTCACCCCCCAACTAACACATTATTTTCCCCTCCCACTCCCATACTAC
   consensus/80%                      ATTTTATCTTTTGGCGGTATGCACTTTTAACAGTCACCCCCCAACTAACACATTATTTTCCCCTCCCACTCCCATACTAC
   consensus/70%                      ATTTTATCTTTTGGCGGTATGCACTTTTAACAGTCACCCCCCAACTAACACATTATTTTCCCCTCCCACTCCCATACTAC

                          cov    pid  481       .         5         .         .         .         .         :         . 560
 1 AGENOME-ZPMRG-U-V1.0  100.0% 100.0%     TAATCTCATCAATACAACCCCCACCCATCCTACCCAGCACACACACACACCGCTGCTAACCCCATACCCCGAACCAACCA
 2 AGENOME-ZPMCG-V1.0   100.0%  99.9%     TAATCTCATCAACACAACCCCCGCCCATCCTACCCAGCACACACACACACCGCTGCTAACCCCATACCCCGAACCAACCA
 3 AGENOME-ZPMRG-M-V1.0 100.0%  99.8%     TAATCTCATCAACACAACCCCCGCCCATCCTACCCAG--CACACACACACCGCTGCTAACCCCATACCCCGAACCAACCA
 4 AGENOME-ZPMRG-F-V1.0  99.9%  99.8%     TAATCTCATCAATACAACCCCCGCCCATCCTACCCAG----CACACACACCGCTGCTAACCCCATACCCCGAACCAACCA
 5 AGENOME-ZPMRG-Z-V1.0  99.9%  99.7%     TAATCTCATCAACACAACCCCCGCCCATCCTACCCAG--CACACACACACCGCTGCTAACCCCATACCCCGAACCAACCA
 6 AGENOME-ZPMRG-T-V1.0  99.9%  99.8%     TAATCTCATCAATACAACCCCCGCCCATCCTACCCAG--CACACACACACCGCTGCTAACCCCATACCCCGAACCAACCA
 7 AGENOME-ZPMRG-HV-V1.0 99.9%  99.8%     TAATCTCATCAATACAACCCCCGCCCATCCTACCCAG--CACACACACACCGCTGCTAACCCCATACCCCGAACCAACCA
 8 AGENOME-ZPMRG-A-V1.0  99.9%  99.8%     TAATCTCATCAATACAACCCCCGCCCATCCTACCCAG--CACACACACACCGCTGCTAACCCCATACCCCGAACCAACCA
 9 NC_012920.1           99.9%  99.8%     TAATCTCATCAATACAACCCCCGCCCATCCTACCCAG--CACACACACACCGCTGCTAACCCCATACCCCGAACCAACCA
   consensus/100%                         TAATCTCATCAAsACAACCCCCuCCCATCCTACCCAG....CACACACACCGCTGCTAACCCCATACCCCGAACCAACCA
   consensus/90%                          TAATCTCATCAAsACAACCCCCuCCCATCCTACCCAG....CACACACACCGCTGCTAACCCCATACCCCGAACCAACCA
   consensus/80%                          TAATCTCATCAAsACAACCCCCGCCCATCCTACCCAG..CACACACACACCGCTGCTAACCCCATACCCCGAACCAACCA
   consensus/70%                          TAATCTCATCAAsACAACCCCCGCCCATCCTACCCAG..CACACACACACCGCTGCTAACCCCATACCCCGAACCAACCA

                          cov    pid  561       .         .         .         6         .         .         .         . 640
 1 AGENOME-ZPMRG-U-V1.0  100.0% 100.0%     AACCCCAAAGACACCCCCCCCCCCCACAGTTTATGTAGCTTACCTCCTCAAAGCAATACACTGAAAATGTTTAGACGGGC
 2 AGENOME-ZPMCG-V1.0   100.0%  99.9%     AACCCCAAAGACACCCCCCCCCCCCACAGTTTATGTAGCTTACCTCCTCAAAGCAATACACTGAAAATGTTTAGACGGGC
 3 AGENOME-ZPMRG-M-V1.0 100.0%  99.8%     AACCCCAAAGACA--CCCCCCCCCCACAGTTTATGTAGCTTACCTCCTCAAAGCAATACACTGAAAATGTTTAGACGGGC
 4 AGENOME-ZPMRG-F-V1.0  99.9%  99.8%     AACCCCAAAGACA------CCCCCCACAGTTTATGTAGCTTACCTCCTCAAAGCAATACACTGAAAATGTTTAGACGGGC
 5 AGENOME-ZPMRG-Z-V1.0  99.9%  99.7%     AACCCCAAAGACA------CCCCCCACAGTTTATGTAGCTTACCTCCTCAAAGCAATACACTGAAAATGTTTAGACGGGC
 6 AGENOME-ZPMRG-T-V1.0  99.9%  99.8%     AACCCCAAAGACA------CCCCCCACAGTTTATGTAGCTTACCTCCTCAAAGCAATACACTGAAAATGTTTAGACGGGC
 7 AGENOME-ZPMRG-HV-V1.0 99.9%  99.8%     AACCCCAAAGACA------CCCCCCACAGTTTATGTAGCTTACCTCCTCAAAGCAATACACTGAAAATGTTTAGACGGGC
 8 AGENOME-ZPMRG-A-V1.0  99.9%  99.8%     AACCCCAAAGACA------CCCCCCACAGTTTATGTAGCTTACCTCCTCAAAGCAATACACTGAAAATGTTTAGACGGGC
 9 NC_012920.1           99.9%  99.8%     AACCCCAAAGACA------CCCCCCACAGTTTATGTAGCTTACCTCCTCAAAGCAATACACTGAAAATGTTTAGACGGGC
   consensus/100%                         AACCCCAAAGACA......CCCCCCACAGTTTATGTAGCTTACCTCCTCAAAGCAATACACTGAAAATGTTTAGACGGGC
   consensus/90%                          AACCCCAAAGACA......CCCCCCACAGTTTATGTAGCTTACCTCCTCAAAGCAATACACTGAAAATGTTTAGACGGGC
   consensus/80%                          AACCCCAAAGACA......CCCCCCACAGTTTATGTAGCTTACCTCCTCAAAGCAATACACTGAAAATGTTTAGACGGGC
   consensus/70%                          AACCCCAAAGACA......CCCCCCACAGTTTATGTAGCTTACCTCCTCAAAGCAATACACTGAAAATGTTTAGACGGGC

                          cov    pid  641       :         .         .         .         .         7         .         . 720
 1 AGENOME-ZPMRG-U-V1.0  100.0% 100.0%     TCACATCACCCCATAAACAAATAGGTTTGGTCCTAGCCTTTCTATTAGCTCTTAGTAAGATTACACATGCAAGCATCCCC
 2 AGENOME-ZPMCG-V1.0   100.0%  99.9%     TCACATCACCCCATAAACAAATAGGTTTGGTCCTAGCCTTTCTATTAGCTCTTAGTAAGATTACACATGCAAGCATCCCC
 3 AGENOME-ZPMRG-M-V1.0 100.0%  99.8%     TCACATCACCCCATAAACAAATAGGTTTGGTCCTAGCCTTTCTATTAGCTCTTAGTAAGATTACACATGCAAGCATCCCC
 4 AGENOME-ZPMRG-F-V1.0  99.9%  99.8%     TCACATCACCCCATAAACAAATAGGTTTGGTCCTAGCCTTTCTATTAGCTCTTAGTAAGATTACACATGCAAGCATCCCC
 5 AGENOME-ZPMRG-Z-V1.0  99.9%  99.7%     TCACATCACCCCATAAACAAATAGGTTTGGTCCTAGCCTTTCTATTAGCTCTTAGTAAGATTACACATGCAAGCATCCCC
 6 AGENOME-ZPMRG-T-V1.0  99.9%  99.8%     TCACATCACCCCATAAACAAATAGGTTTGGTCCTAGCCTTTCTATTAGCTCTTAGTAAGATTACACATGCAAGCATCCCC
 7 AGENOME-ZPMRG-HV-V1.0 99.9%  99.8%     TCACATCACCCCATAAACAAATAGGTTTGGTCCTAGCCTTTCTATTAGCTCTTAGTAAGATTACACATGCAAGCATCCCC
 8 AGENOME-ZPMRG-A-V1.0  99.9%  99.8%     TCACATCACCCCATAAACAAATAGGTTTGGTCCTAGCCTTTCTATTAGCTCTTAGTAAGATTACACATGCAAGCATCCCC
 9 NC_012920.1           99.9%  99.8%     TCACATCACCCCATAAACAAATAGGTTTGGTCCTAGCCTTTCTATTAGCTCTTAGTAAGATTACACATGCAAGCATCCCC
   consensus/100%                         TCACATCACCCCATAAACAAATAGGTTTGGTCCTAGCCTTTCTATTAGCTCTTAGTAAGATTACACATGCAAGCATCCCC
   consensus/90%                          TCACATCACCCCATAAACAAATAGGTTTGGTCCTAGCCTTTCTATTAGCTCTTAGTAAGATTACACATGCAAGCATCCCC
   consensus/80%                          TCACATCACCCCATAAACAAATAGGTTTGGTCCTAGCCTTTCTATTAGCTCTTAGTAAGATTACACATGCAAGCATCCCC
   consensus/70%                          TCACATCACCCCATAAACAAATAGGTTTGGTCCTAGCCTTTCTATTAGCTCTTAGTAAGATTACACATGCAAGCATCCCC

                          cov    pid  721       .         .         :         .         .         .         .         8 800
 1 AGENOME-ZPMRG-U-V1.0  100.0% 100.0%     GTTCCAGTGAGTTCACCCTCTAAATCACCACGATCAAAAGGGACAAGCATCAAGCACGCAGCAATGCAGCTCAAAACGCT
 2 AGENOME-ZPMCG-V1.0   100.0%  99.9%     GTTCCAGTGAGTTCACCCTCTAAATCACCACGATCAAAAGGGACAAGCATCAAGCACGCAGCAATGCAGCTCAAAACGCT
 3 AGENOME-ZPMRG-M-V1.0 100.0%  99.8%     GTTCCAGTGAGTTCACCCTCTAAATCACCACGATCAAAAGGGACAAGCATCAAGCACGCAGCAATGCAGCTCAAAACGCT
 4 AGENOME-ZPMRG-F-V1.0  99.9%  99.8%     GTTCCAGTGAGTTCACCCTCTAAATCACCACGATCAAAAGGGACAAGCATCAAGCACGCAGCAATGCAGCTCAAAACGCT
 5 AGENOME-ZPMRG-Z-V1.0  99.9%  99.7%     GTTCCAGTGAGTTCACCCTCTAAATCACCACGATCAAAAGGGACAAGCATCAAGCACGCAGCAATGCAGCTCAAAACGCT
 6 AGENOME-ZPMRG-T-V1.0  99.9%  99.8%     ATTCCAGTGAGTTCACCCTCTAAATCACCACGATCAAAAGGGACAAGCATCAAGCACGCAGCAATGCAGCTCAAAACGCT
 7 AGENOME-ZPMRG-HV-V1.0 99.9%  99.8%     GTTCCAGTGAGTTCACCCTCTAAATCACCACGATCAAAAGGGACAAGCATCAAGCACGCAGCAATGCAGCTCAAAACGCT
 8 AGENOME-ZPMRG-A-V1.0  99.9%  99.8%     GTTCCAGTGAGTTCACCCTCTAAATCACCACGATCAAAAGGGACAAGCATCAAGCACGCAGCAATGCAGCTCAAAACGCT
 9 NC_012920.1           99.9%  99.8%     GTTCCAGTGAGTTCACCCTCTAAATCACCACGATCAAAAGGAACAAGCATCAAGCACGCAGCAATGCAGCTCAAAACGCT
   consensus/100%                         uTTCCAGTGAGTTCACCCTCTAAATCACCACGATCAAAAGGuACAAGCATCAAGCACGCAGCAATGCAGCTCAAAACGCT
   consensus/90%                          uTTCCAGTGAGTTCACCCTCTAAATCACCACGATCAAAAGGuACAAGCATCAAGCACGCAGCAATGCAGCTCAAAACGCT
   consensus/80%                          GTTCCAGTGAGTTCACCCTCTAAATCACCACGATCAAAAGGGACAAGCATCAAGCACGCAGCAATGCAGCTCAAAACGCT
   consensus/70%                          GTTCCAGTGAGTTCACCCTCTAAATCACCACGATCAAAAGGGACAAGCATCAAGCACGCAGCAATGCAGCTCAAAACGCT

                          cov    pid  801       .         .         .         .         :         .         .         . 880
 1 AGENOME-ZPMRG-U-V1.0  100.0% 100.0%     TAGCCTAGCCACACCCCCACGGGAAACAGCAGTGATTAACCTTTAGCAATAAACGAAGTTTAACTAAGCTATACTAACC
 2 AGENOME-ZPMCG-V1.0   100.0%  99.9%     TAGCCTAGCCACACCCCCACGGGAAACAGCAGTGATTAACCTTTAGCAATAAACGAAGTTTAACTAAGCTATACTAACC
 3 AGENOME-ZPMRG-M-V1.0 100.0%  99.8%     TAGCCTAGCCACACCCCCACGGGAAACAGCAGTGATTAACCTTTAGCAATAAACGAAGTTTAACTAAGCTATACTAACC
 4 AGENOME-ZPMRG-F-V1.0  99.9%  99.8%     TAGCCTAGCCACACCCCCACGGGAAACAGCAGTGATTAACCTTTAGCAATAAACGAAGTTTAACTAAGCTATACTAACC
 5 AGENOME-ZPMRG-Z-V1.0  99.9%  99.7%     TAGCCTAGCCACACCCCCACGGGAAACAGCAGTGATTAACCTTTAGCAATAAACGAAGTTTAACTAAGCTATACTAACC
 6 AGENOME-ZPMRG-T-V1.0  99.9%  99.8%     TAGCCTAGCCACACCCCCACGGGAAACAGCAGTGATTAACCTTTAGCAATAAACGAAGTTTAACTAAGCTATACTAACC
 7 AGENOME-ZPMRG-HV-V1.0 99.9%  99.8%     TAGCCTAGCCACACCCCCACGGGAAACAGCAGTGATTAACCTTTAGCAATAAACGAAGTTTAACTAAGCTATACTAACC
 8 AGENOME-ZPMRG-A-V1.0  99.9%  99.8%     TAGCCTAGCCACACCCCCACGGGAAACAGCAGTGATTAACCTTTAGCAATAAACGAAGTTTAACTAAGCTATACTAACC
 9 NC_012920.1           99.9%  99.8%     TAGCCTAGCCACACCCCCACGGGAAACAGCAGTGATTAACCTTTAGCAATAAACGAAGTTTAACTAAGCTATACTAACC
   consensus/100%                         TAGCCTAGCCACACCCCCACGGGAAACAGCAGTGATTAACCTTTAGCAATAAACGAAGTTTAACTAAGCTATACTAACC
   consensus/90%                          TAGCCTAGCCACACCCCCACGGGAAACAGCAGTGATTAACCTTTAGCAATAAACGAAGTTTAACTAAGCTATACTAACC
   consensus/80%                          TAGCCTAGCCACACCCCCACGGGAAACAGCAGTGATTAACCTTTAGCAATAAACGAAGTTTAACTAAGCTATACTAACC
   consensus/70%                          TAGCCTAGCCACACCCCCACGGGAAACAGCAGTGATTAACCTTTAGCAATAAACGAAGTTTAACTAAGCTATACTAACC

                          cov    pid  881       .         9         .         .         .         .         :         . 960
 1 AGENOME-ZPMRG-U-V1.0  100.0% 100.0%     CCAGGGTTGGTCAATTTCGTGCCAGCCACCGCGGTCACACGATTAACCCAAGTCAATAGAAGCCGGCGTAAAGAGTGTTT
 2 AGENOME-ZPMCG-V1.0   100.0%  99.9%     CCAGGGTTGGTCAATTTCGTGCCAGCCACCGCGGTCACACGATTAACCCAAGTCAATAGAAGCCGGCGTAAAGAGTGTTT
 3 AGENOME-ZPMRG-M-V1.0 100.0%  99.8%     CCAGGGTTGGTCAATTTCGTGCCAGCCACCGCGGTCACACGATTAACCCAAGTCAATAGAAGCCGGCGTAAAGAGTGTTT
 4 AGENOME-ZPMRG-F-V1.0  99.9%  99.8%     CCAGGGTTGGTCAATTTCGTGCCAGCCACCGCGGTCACACGATTAACCCAAGTCAATAGAAGCCGGCGTAAAGAGTGTTT
 5 AGENOME-ZPMRG-Z-V1.0  99.9%  99.7%     CCAGGGTTGGTCAATTTCGTGCCAGCCACCGCGGTCACACGATTAACCCAAGTCAATAGAAGCCGGCGTAAAGAGTGTTT
 6 AGENOME-ZPMRG-T-V1.0  99.9%  99.8%     CCAGGGTTGGTCAATTTCGTGCCAGCCACCGCGGTCACACGATTAACCCAAGTCAATAGAAGCCGGCGTAAAGAGTGTTT
 7 AGENOME-ZPMRG-HV-V1.0 99.9%  99.8%     CCAGGGTTGGTCAATTTCGTGCCAGCCACCGCGGTCACACGATTAACCCAAGTCAATAGAAGCCGGCGTAAAGAGTGTTT
 8 AGENOME-ZPMRG-A-V1.0  99.9%  99.8%     CCAGGGTTGGTCAATTTCGTGCCAGCCACCGCGGTCACACGATTAACCCAAGTCAATAGAAGCCGGCGTAAAGAGTGTTT
 9 NC_012920.1           99.9%  99.8%     CCAGGGTTGGTCAATTTCGTGCCAGCCACCGCGGTCACACGATTAACCCAAGTCAATAGAAGCCGGCGTAAAGAGTGTTT
   consensus/100%                         CCAGGGTTGGTCAATTTCGTGCCAGCCACCGCGGTCACACGATTAACCCAAGTCAATAGAAGCCGGCGTAAAGAGTGTTT
```

```
    consensus/90%                               CCAGGGTTGGTCAATTTCGTGCCAGCCACCGCGGTCACACGATTAACCCAAGTCAATAGAAGCCGGCGTAAAGAGTGTTT
    consensus/80%                               CCAGGGTTGGTCAATTTCGTGCCAGCCACCGCGGTCACACGATTAACCCAAGTCAATAGAAGCCGGCGTAAAGAGTGTTT
    consensus/70%                               CCAGGGTTGGTCAATTTCGTGCCAGCCACCGCGGTCACACGATTAACCCAAGTCAATAGAAGCCGGCGTAAAGAGTGTTT

                             cov    pid   961          .         .         .         0         .         .         .         . 1040
  1 AGENOME-ZPMRG-U-V1.0  100.0% 100.0%        TAGATCACCCCCTCCCCAATAAAGCTAAAACTCACCTGAGTTGTAAAAAACTCCAGTTGACACAAAATAGACTACGAAAG
  2 AGENOME-ZPMCG-V1.0    100.0%  99.9%        TAGATCACCCCCTCCCCAATAAAGCTAAAACTCACCTGAGTTGTAAAAAACTCCAGTTGACACAAAATAGACTACGAAAG
  3 AGENOME-ZPMRG-M-V1.0  100.0%  99.8%        TAGATCACCCCCTCCCCAATAAAGCTAAAACTCACCTGAGTTGTAAAAAACTCCAGTTGACACAAAATAGACTACGAAAG
  4 AGENOME-ZPMRG-F-V1.0   99.9%  99.8%        TAGATCACCCCCTCCCCAATAAAGCTAAAACTCACCTGAGTTGTAAAAAACTCCAGTTGACACAAAATAGACTACGAAAG
  5 AGENOME-ZPMRG-Z-V1.0   99.9%  99.7%        TAGATCACCCCCTCCCCAATAAAGCTAAAACTCACCTGAGTTGTAAAAAACTCCAGTTGACACAAAATAGACTACGAAAG
  6 AGENOME-ZPMRG-T-V1.0   99.9%  99.8%        TAGATCACCCCCTCCCCAATAAAGCTAAAACTCACCTGAGTTGTAAAAAACTCCAGTTGACACAAAATAGACTACGAAAG
  7 AGENOME-ZPMRG-HV-V1.0  99.9%  99.8%        TAGATCACCCCCTCCCCAATAAAGCTAAAACTCACCTGAGTTGTAAAAAACTCCAGTTGACACAAAATAGACTACGAAAG
  8 AGENOME-ZPMRG-A-V1.0   99.9%  99.8%        TAGATCACCCCCTCCCCAATAAAGCTAAAACTCACCTGAGTTGTAAAAAACTCCAGTTGACACAAAATAGACTACGAAAG
  9 NC_012920.1            99.9%  99.8%        TAGATCACCCCCTCCCCAATAAAGCTAAAACTCACCTGAGTTGTAAAAAACTCCAGTTGACACAAAATAGACTACGAAAG
    consensus/100%                             TAGATCACCCCCTCCCCAATAAAGCTAAAACTCACCTGAGTTGTAAAAAACTCCAGTTGACACAAAATAGACTACGAAAG
    consensus/90%                              TAGATCACCCCCTCCCCAATAAAGCTAAAACTCACCTGAGTTGTAAAAAACTCCAGTTGACACAAAATAGACTACGAAAG
    consensus/80%                              TAGATCACCCCCTCCCCAATAAAGCTAAAACTCACCTGAGTTGTAAAAAACTCCAGTTGACACAAAATAGACTACGAAAG
    consensus/70%                              TAGATCACCCCCTCCCCAATAAAGCTAAAACTCACCTGAGTTGTAAAAAACTCCAGTTGACACAAAATAGACTACGAAAG

                             cov    pid  1041          :         .         .         .         .         1         .         . 1120
  1 AGENOME-ZPMRG-U-V1.0  100.0% 100.0%        TGGCTTTAACATATCTGAACACACAATAGCTAAGACCCAAACTGGGATTAGATACCCCACTATGCTTAGCCCTAAACCTC
  2 AGENOME-ZPMCG-V1.0    100.0%  99.9%        TGGCTTTAACATATCTGAACACACAATAGCTAAGACCCAAACTGGGATTAGATACCCCACTATGCTTAGCCCTAAACCTC
  3 AGENOME-ZPMRG-M-V1.0  100.0%  99.8%        TGGCTTTAACATATCTGAACACACAATAGCTAAGACCCAAACTGGGATTAGATACCCCACTATGCTTAGCCCTAAACCTC
  4 AGENOME-ZPMRG-F-V1.0   99.9%  99.8%        TGGCTTTAACATATCTGAACACACAATAGCTAAGACCCAAACTGGGATTAGATACCCCACTATGCTTAGCCCTAAACCTC
  5 AGENOME-ZPMRG-Z-V1.0   99.9%  99.7%        TGGCTTTAACATATCTGAACACACAATAGCTAAGACCCAAACTGGGATTAGATACCCCACTATGCTTAGCCCTAAACCTC
  6 AGENOME-ZPMRG-T-V1.0   99.9%  99.8%        TGGCTTTAACATATCTGAACACACAATAGCTAAGACCCAAACTGGGATTAGATACCCCACTATGCTTAGCCCTAAACCTC
  7 AGENOME-ZPMRG-HV-V1.0  99.9%  99.8%        TGGCTTTAACATATCTGAACACACAATAGCTAAGACCCAAACTGGGATTAGATACCCCACTATGCTTAGCCCTAAACCTC
  8 AGENOME-ZPMRG-A-V1.0   99.9%  99.8%        TGGCTTTAACATATCTGAACACACAATAGCTAAGACCCAAACTGGGATTAGATACCCCACTATGCTTAGCCCTAAACCTC
  9 NC_012920.1            99.9%  99.8%        TGGCTTTAACATATCTGAACACACAATAGCTAAGACCCAAACTGGGATTAGATACCCCACTATGCTTAGCCCTAAACCTC
    consensus/100%                             TGGCTTTAACATATCTGAACACACAATAGCTAAGACCCAAACTGGGATTAGATACCCCACTATGCTTAGCCCTAAACCTC
    consensus/90%                              TGGCTTTAACATATCTGAACACACAATAGCTAAGACCCAAACTGGGATTAGATACCCCACTATGCTTAGCCCTAAACCTC
    consensus/80%                              TGGCTTTAACATATCTGAACACACAATAGCTAAGACCCAAACTGGGATTAGATACCCCACTATGCTTAGCCCTAAACCTC
    consensus/70%                              TGGCTTTAACATATCTGAACACACAATAGCTAAGACCCAAACTGGGATTAGATACCCCACTATGCTTAGCCCTAAACCTC

                             cov    pid  1121          .         .         :         .         .         .         .         2 1200
  1 AGENOME-ZPMRG-U-V1.0  100.0% 100.0%        AACAGTTAAATCAACAAAACTGCTCGCCAGAACACTACGAGCCACAGCTTAAAACTCAAAGGACCTGGCGGTGCTTCATA
  2 AGENOME-ZPMCG-V1.0    100.0%  99.9%        AACAGTTAAATCAACAAAACTGCTCGCCAGAACACTACGAGCCACAGCTTAAAACTCAAAGGACCTGGCGGTGCTTCATA
  3 AGENOME-ZPMRG-M-V1.0  100.0%  99.8%        AACAGTTAAATCAACAAAACTGCTCGCCAGAACACTACGAGCCACAGCTTAAAACTCAAAGGACCTGGCGGTGCTTCATA
  4 AGENOME-ZPMRG-F-V1.0   99.9%  99.8%        AACAGTTAAATCAACAAAACTGCTCGCCAGAACACTACGAGCCACAGCTTAAAACTCAAAGGACCTGGCGGTGCTTCATA
  5 AGENOME-ZPMRG-Z-V1.0   99.9%  99.7%        AACAGTTAAATCAACAAAACTGCTCGCCAGAACACTACGAGCCACAGCTTAAAACTCAAAGGACCTGGCGGTGCTTCATA
  6 AGENOME-ZPMRG-T-V1.0   99.9%  99.8%        AACAGTTAAATCAACAAAACTGCTCGCCAGAACACTACGAGCCACAGCTTAAAACTCAAAGGACCTGGCGGTGCTTCATA
  7 AGENOME-ZPMRG-HV-V1.0  99.9%  99.8%        AACAGTTAAATCAACAAAACTGCTCGCCAGAACACTACGAGCCACAGCTTAAAACTCAAAGGACCTGGCGGTGCTTCATA
  8 AGENOME-ZPMRG-A-V1.0   99.9%  99.8%        AACAGTTAAATCAACAAAACTGCTCGCCAGAACACTACGAGCCACAGCTTAAAACTCAAAGGACCTGGCGGTGCTTCATA
  9 NC_012920.1            99.9%  99.8%        AACAGTTAAATCAACAAAACTGCTCGCCAGAACACTACGAGCCACAGCTTAAAACTCAAAGGACCTGGCGGTGCTTCATA
    consensus/100%                             AACAGTTAAATCAACAAAACTGCTCGCCAGAACACTACGAGCCACAGCTTAAAACTCAAAGGACCTGGCGGTGCTTCATA
    consensus/90%                              AACAGTTAAATCAACAAAACTGCTCGCCAGAACACTACGAGCCACAGCTTAAAACTCAAAGGACCTGGCGGTGCTTCATA
    consensus/80%                              AACAGTTAAATCAACAAAACTGCTCGCCAGAACACTACGAGCCACAGCTTAAAACTCAAAGGACCTGGCGGTGCTTCATA
    consensus/70%                              AACAGTTAAATCAACAAAACTGCTCGCCAGAACACTACGAGCCACAGCTTAAAACTCAAAGGACCTGGCGGTGCTTCATA

                             cov    pid  1201          .         .         .         .         :         .         .         . 1280
  1 AGENOME-ZPMRG-U-V1.0  100.0% 100.0%        TCCCTCTAGAGGAGCCTGTTCTGTAATCGATAAACCCCGATCAACCTCACCACCTCTTGCTCAGCCTATATACCGCCATC
  2 AGENOME-ZPMCG-V1.0    100.0%  99.9%        TCCCTCTAGAGGAGCCTGTTCTGTAATCGATAAACCCCGATCAACCTCACCACCTCTTGCTCAGCCTATATACCGCCATC
  3 AGENOME-ZPMRG-M-V1.0  100.0%  99.8%        TCCCTCTAGAGGAGCCTGTTCTGTAATCGATAAACCCCGATCAACCTCACCACCTCTTGCTCAGCCTATATACCGCCATC
  4 AGENOME-ZPMRG-F-V1.0   99.9%  99.8%        TCCCTCTAGAGGAGCCTGTTCTGTAATCGATAAACCCCGATCAACCTCACCACCTCTTGCTCAGCCTATATACCGCCATC
  5 AGENOME-ZPMRG-Z-V1.0   99.9%  99.7%        TCCCTCTAGAGGAGCCTGTTCTGTAATCGATAAACCCCGATCAACCTCACCACCTCTTGCTCAGCCTATATACCGCCATC
  6 AGENOME-ZPMRG-T-V1.0   99.9%  99.8%        TCCCTCTAGAGGAGCCTGTTCTGTAATCGATAAACCCCGATCAACCTCACCACCTCTTGCTCAGCCTATATACCGCCATC
  7 AGENOME-ZPMRG-HV-V1.0  99.9%  99.8%        TCCCTCTAGAGGAGCCTGTTCTGTAATCGATAAACCCCGATCAACCTCACCACCTCTTGCTCAGCCTATATACCGCCATC
  8 AGENOME-ZPMRG-A-V1.0   99.9%  99.8%        TCCCTCTAGAGGAGCCTGTTCTGTAATCGATAAACCCCGATCAACCTCACCACCTCTTGCTCAGCCTATATACCGCCATC
  9 NC_012920.1            99.9%  99.8%        TCCCTCTAGAGGAGCCTGTTCTGTAATCGATAAACCCCGATCAACCTCACCACCTCTTGCTCAGCCTATATACCGCCATC
    consensus/100%                             TCCCTCTAGAGGAGCCTGTTCTGTAATCGATAAACCCCGATCAACCTCACCACCTCTTGCTCAGCCTATATACCGCCATC
    consensus/90%                              TCCCTCTAGAGGAGCCTGTTCTGTAATCGATAAACCCCGATCAACCTCACCACCTCTTGCTCAGCCTATATACCGCCATC
    consensus/80%                              TCCCTCTAGAGGAGCCTGTTCTGTAATCGATAAACCCCGATCAACCTCACCACCTCTTGCTCAGCCTATATACCGCCATC
    consensus/70%                              TCCCTCTAGAGGAGCCTGTTCTGTAATCGATAAACCCCGATCAACCTCACCACCTCTTGCTCAGCCTATATACCGCCATC

                             cov    pid  1281          .         3         .         .         .         .         :         . 1360
  1 AGENOME-ZPMRG-U-V1.0  100.0% 100.0%        TTCAGCAAACCCTGATGAAGGCTACAAAGTAAGCGCAAGTACCCACGTAAAGACGTTAGGTCAAGGTGTAGCCCATGAGG
  2 AGENOME-ZPMCG-V1.0    100.0%  99.9%        TTCAGCAAACCCTGATGAAGGCTACAAAGTAAGCGCAAGTACCCACGTAAAGACGTTAGGTCAAGGTGTAGCCCATGAGG
  3 AGENOME-ZPMRG-M-V1.0  100.0%  99.8%        TTCAGCAAACCCTGATGAAGGCTACAAAGTAAGCGCAAGTACCCACGTAAAGACGTTAGGTCAAGGTGTAGCCCATGAGG
  4 AGENOME-ZPMRG-F-V1.0   99.9%  99.8%        TTCAGCAAACCCTGATGAAGGCTACAAAGTAAGCGCAAGTACCCACGTAAAGACGTTAGGTCAAGGTGTAGCCCATGAGG
  5 AGENOME-ZPMRG-Z-V1.0   99.9%  99.7%        TTCAGCAAACCCTGATGAAGGCTACAAAGTAAGCGCAAGTACCCACGTAAAGACGTTAGGTCAAGGTGTAGCCCATGAGG
  6 AGENOME-ZPMRG-T-V1.0   99.9%  99.8%        TTCAGCAAACCCTGATGAAGGCTACAAAGTAAGCGCAAGTACCCACGTAAAGACGTTAGGTCAAGGTGTAGCCCATGAGG
  7 AGENOME-ZPMRG-HV-V1.0  99.9%  99.8%        TTCAGCAAACCCTGATGAAGGCTACAAAGTAAGCGCAAGTACCCACGTAAAGACGTTAGGTCAAGGTGTAGCCCATGAGG
  8 AGENOME-ZPMRG-A-V1.0   99.9%  99.8%        TTCAGCAAACCCTGATGAAGGCTACAAAGTAAGCGCAAGTACCCACGTAAAGACGTTAGGTCAAGGTGTAGCCCATGAGG
  9 NC_012920.1            99.9%  99.8%        TTCAGCAAACCCTGATGAAGGCTACAAAGTAAGCGCAAGTACCCACGTAAAGACGTTAGGTCAAGGTGTAGCCCATGAGG
    consensus/100%                             TTCAGCAAACCCTGATGAAGGCTACAAAGTAAGCGCAAGTACCCACGTAAAGACGTTAGGTCAAGGTGTAGCCCATGAGG
    consensus/90%                              TTCAGCAAACCCTGATGAAGGCTACAAAGTAAGCGCAAGTACCCACGTAAAGACGTTAGGTCAAGGTGTAGCCCATGAGG
    consensus/80%                              TTCAGCAAACCCTGATGAAGGCTACAAAGTAAGCGCAAGTACCCACGTAAAGACGTTAGGTCAAGGTGTAGCCCATGAGG
    consensus/70%                              TTCAGCAAACCCTGATGAAGGCTACAAAGTAAGCGCAAGTACCCACGTAAAGACGTTAGGTCAAGGTGTAGCCCATGAGG

                             cov    pid  1361          .         .         .         4         .         .         .         . 1440
  1 AGENOME-ZPMRG-U-V1.0  100.0% 100.0%        TGGCAAGAAATGGGCTACATTTTCTACCCCAGAAAACTACGATAGCCCTTATGAAACTTAAGGGTCGAAGGTGGATTTAG
  2 AGENOME-ZPMCG-V1.0    100.0%  99.9%        TGGCAAGAAATGGGCTACATTTTCTACCCCAGAAAACTACGATAGCCCTTATGAAACTTAAGGGTCGAAGGTGGATTTAG
  3 AGENOME-ZPMRG-M-V1.0  100.0%  99.8%        TGGCAAGAAATGGGCTACATTTTCTACCCCAGAAAACTACGATAGCCCTTATGAAACTTAAGGGTCGAAGGTGGATTTAG
  4 AGENOME-ZPMRG-F-V1.0   99.9%  99.8%        TGGCAAGAAATGGGCTACATTTTCTACCCCAGAAAACTACGATAGCCCTTATGAAACTTAAGGGTCGAAGGTGGATTTAG
  5 AGENOME-ZPMRG-Z-V1.0   99.9%  99.7%        TGGCAAGAAATGGGCTACATTTTCTACCCCAGAAAACTACGATAGCCCTTATGAAACTTAAGGGTCGAAGGTGGATTTAG
  6 AGENOME-ZPMRG-T-V1.0   99.9%  99.8%        TGGCAAGAAATGGGCTACATTTTCTACCCCAGAAAACTACGATAGCCCTTATGAAACTTAAGGGTCGAAGGTGGATTTAG
  7 AGENOME-ZPMRG-HV-V1.0  99.9%  99.8%        TGGCAAGAAATGGGCTACATTTTCTACCCCAGAAAACTACGATAGCCCTTATGAAACTTAAGGGTCGAAGGTGGATTTAG
  8 AGENOME-ZPMRG-A-V1.0   99.9%  99.8%        TGGCAAGAAATGGGCTACATTTTCTACCCCAGAAAACTACGATAGCCCTTATGAAACTTAAGGGTCGAAGGTGGATTTAG
  9 NC_012920.1            99.9%  99.8%        TGGCAAGAAATGGGCTACATTTTCTACCCCAGAAAACTACGATAGCCCTTATGAAACTTAAGGGTCGAAGGTGGATTTAG
    consensus/100%                             TGGCAAGAAATGGGCTACATTTTCTACCCCAGAAAACTACGATAGCCCTTATGAAACTTAAGGGTCGAAGGTGGATTTAG
    consensus/90%                              TGGCAAGAAATGGGCTACATTTTCTACCCCAGAAAACTACGATAGCCCTTATGAAACTTAAGGGTCGAAGGTGGATTTAG
```

```
   consensus/80%                        TGGCAAGAAATGGGCTACATTTTCTACCCCAGAAAACTACGATAGCCCTTATGAAACTTAAGGGTCGAAGGTGGATTTAG
   consensus/70%                        TGGCAAGAAATGGGCTACATTTTCTACCCCAGAAAACTACGATAGCCCTTATGAAACTTAAGGGTCGAAGGTGGATTTAG

                         cov    pid 1441         :         .         .         .         .         5         .         . 1520
 1 AGENOME-ZPMRG-U-V1.0  100.0% 100.0%     CAGTAAACTGAGAGTAGAGTGCTTAGTTGAACAGGGCCCTGAAGCGCGTACACACCGCCCGTCACCCTCCTCAAGTATAC
 2 AGENOME-ZPMCG-V1.0   100.0%  99.9%     CAGTAAACTGAGAGTAGAGTGCTTAGTTGAACAGGGCCCTGAAGCGCGTACACACCGCCCGTCACCCTCCTCAAGTATAC
 3 AGENOME-ZPMRG-M-V1.0 100.0%  99.8%     CAGTAAACTGAGAGTAGAGTGCTTAGTTGAACAGGGCCCTGAAGCGCGTACACACCGCCCGTCACCCTCCTCAAGTATAC
 4 AGENOME-ZPMRG-F-V1.0  99.9%  99.8%     CAGTAAACTGAGAGTAGAGTGCTTAGTTGAACAGGGCCCTGAAGCGCGTACACACCGCCCGTCACCCTCCTCAAGTATAC
 5 AGENOME-ZPMRG-Z-V1.0  99.9%  99.7%     CAGTAAACTGAGAGTAGAGTGCTTAGTTGAACAGGGCCCTGAAGCGCGTACACACCGCCCGTCACCCTCCTCAAGTATAC
 6 AGENOME-ZPMRG-T-V1.0  99.9%  99.8%     CAGTAAACTGAGAGTAGAGTGCTTAGTTGAACAGGGCCCTGAAGCGCGTACACACCGCCCGTCACCCTCCTCAAGTATAC
 7 AGENOME-ZPMRG-HV-V1.0 99.9%  99.8%     CAGTAAACTGAGAGTAGAGTGCTTAGTTGAACAGGGCCCTGAAGCGCGTACACACCGCCCGTCACCCTCCTCAAGTATAC
 8 AGENOME-ZPMRG-A-V1.0  99.9%  99.8%     CAGTAAACTGAGAGTAGAGTGCTTAGTTGAACAGGGCCCTGAAGCGCGTACACACCGCCCGTCACCCTCCTCAAGTATAC
 9 NC_012920.1           99.9%  99.8%     CAGTAAACTAAGAGTAGAGTGCTTAGTTGAACAGGGCCCTGAAGCGCGTACACACCGCCCGTCACCCTCCTCAAGTATAC
   consensus/100%                       CAGTAAACTuAGAGTAGAGTGCTTAGTTGAACAGGGCCCTGAAGCGCGTACACACCGCCCGTCACCCTCCTCAAGTATAC
   consensus/90%                        CAGTAAACTuAGAGTAGAGTGCTTAGTTGAACAGGGCCCTGAAGCGCGTACACACCGCCCGTCACCCTCCTCAAGTATAC
   consensus/80%                        CAGTAAACTGAGAGTAGAGTGCTTAGTTGAACAGGGCCCTGAAGCGCGTACACACCGCCCGTCACCCTCCTCAAGTATAC
   consensus/70%                        CAGTAAACTGAGAGTAGAGTGCTTAGTTGAACAGGGCCCTGAAGCGCGTACACACCGCCCGTCACCCTCCTCAAGTATAC

                         cov    pid 1521         .         .         :         .         .         .         .         6 1600
 1 AGENOME-ZPMRG-U-V1.0  100.0% 100.0%     TTCAAAGGACATTTAACTAAAACCCCTACGCATTTATATAGAGGAGACAAGTCGTAACATGGTAAGTGTACTGGAAAGTG
 2 AGENOME-ZPMCG-V1.0   100.0%  99.9%     TTCAAAGGACATTTAACTAAAACCCCTACGCATTTATATAGAGGAGACAAGTCGTAACATGGTAAGTGTACTGGAAAGTG
 3 AGENOME-ZPMRG-M-V1.0 100.0%  99.8%     TTCAAAGGACATTTAACTAAAACCCCTACGCATTTATATAGAGGAGACAAGTCGTAACATGGTAAGTGTACTGGAAAGTG
 4 AGENOME-ZPMRG-F-V1.0  99.9%  99.8%     TTCAAAGGACATTTAACTAAAACCCCTACGCATTTATATAGAGGAGACAAGTCGTAACATGGTAAGTGTACTGGAAAGTG
 5 AGENOME-ZPMRG-Z-V1.0  99.9%  99.7%     TTCAAAGGACATTTAACTAAAACCCCTACGCATTTATATAGAGGAGACAAGTCGTAACATGGTAAGTGTACTGGAAAGTG
 6 AGENOME-ZPMRG-T-V1.0  99.9%  99.8%     TTCAAAGGACATTTAACTAAAACCCCTACGCATTTATATAGAGGAGACAAGTCGTAACATGGTAAGTGTACTGGAAAGTG
 7 AGENOME-ZPMRG-HV-V1.0 99.9%  99.8%     TTCAAAGGACATTTAACTAAAACCCCTACGCATTTATATAGAGGAGACAAGTCGTAACATGGTAAGTGTACTGGAAAGTG
 8 AGENOME-ZPMRG-A-V1.0  99.9%  99.8%     TTCAAAGGACATTTAACTAAAACCCCTACGCATTTATATAGAGGAGACAAGTCGTAACATGGTAAGTGTACTGGAAAGTG
 9 NC_012920.1           99.9%  99.8%     TTCAAAGGACATTTAACTAAAACCCCTACGCATTTATATAGAGGAGACAAGTCGTAACATGGTAAGTGTACTGGAAAGTG
   consensus/100%                       TTCAAAGGACATTTAACTAAAACCCCTACGCATTTATATAGAGGAGACAAGTCGTAACATGGTAAGTGTACTGGAAAGTG
   consensus/90%                        TTCAAAGGACATTTAACTAAAACCCCTACGCATTTATATAGAGGAGACAAGTCGTAACATGGTAAGTGTACTGGAAAGTG
   consensus/80%                        TTCAAAGGACATTTAACTAAAACCCCTACGCATTTATATAGAGGAGACAAGTCGTAACATGGTAAGTGTACTGGAAAGTG
   consensus/70%                        TTCAAAGGACATTTAACTAAAACCCCTACGCATTTATATAGAGGAGACAAGTCGTAACATGGTAAGTGTACTGGAAAGTG

                         cov    pid 1601         .         .         .         .         :         .         .         . 1680
 1 AGENOME-ZPMRG-U-V1.0  100.0% 100.0%     CACTTGGACGAACCAGAGTGTAGCTTAACACAAAGCACCCAACTTACACTTAGGAGATTTCAACTTAACTTGACCGCTCT
 2 AGENOME-ZPMCG-V1.0   100.0%  99.9%     CACTTGGACGAACCAGAGTGTAGCTTAACACAAAGCACCCAACTTACACTTAGGAGATTTCAACTTAACTTGACCGCTCT
 3 AGENOME-ZPMRG-M-V1.0 100.0%  99.8%     CACTTGGACGAACCAGAGTGTAGCTTAACACAAAGCACCCAACTTACACTTAGGAGATTTCAACTTAACTTGACCGCTCT
 4 AGENOME-ZPMRG-F-V1.0  99.9%  99.8%     CACTTGGACGAACCAGAGTGTAGCTTAACACAAAGCACCCAACTTACACTTAGGAGATTTCAACTTAACTTGACCGCTCT
 5 AGENOME-ZPMRG-Z-V1.0  99.9%  99.7%     CACTTGGACGAACCAGAGTGTAGCTTAACACAAAGCACCCAACTTACACTTAGGAGATTTCAACTTAACTTGACCGCTCT
 6 AGENOME-ZPMRG-T-V1.0  99.9%  99.8%     CACTTGGACGAACCAGAGTGTAGCTTAACACAAAGCACCCAACTTACACTTAGGAGATTTCAACTTAACTTGACCGCTCT
 7 AGENOME-ZPMRG-HV-V1.0 99.9%  99.8%     CACTTGGACGAACCAGAGTGTAGCTTAACACAAAGCACCCAACTTACACTTAGGAGATTTCAACTTAACTTGACCGCTCT
 8 AGENOME-ZPMRG-A-V1.0  99.9%  99.8%     CACTTGGACGAACCAGAGTGTAGCTTAACACAAAGCACCCAACTTACACTTAGGAGATTTCAACTTAACTTGACCGCTCT
 9 NC_012920.1           99.9%  99.8%     CACTTGGACGAACCAGAGTGTAGCTTAACACAAAGCACCCAACTTACACTTAGGAGATTTCAACTTAACTTGACCGCTCT
   consensus/100%                       CACTTGGACGAACCAGAGTGTAGCTTAACACAAAGCACCCAACTTACACTTAGGAGATTTCAACTTAACTTGACCGCTCT
   consensus/90%                        CACTTGGACGAACCAGAGTGTAGCTTAACACAAAGCACCCAACTTACACTTAGGAGATTTCAACTTAACTTGACCGCTCT
   consensus/80%                        CACTTGGACGAACCAGAGTGTAGCTTAACACAAAGCACCCAACTTACACTTAGGAGATTTCAACTTAACTTGACCGCTCT
   consensus/70%                        CACTTGGACGAACCAGAGTGTAGCTTAACACAAAGCACCCAACTTACACTTAGGAGATTTCAACTTAACTTGACCGCTCT

                         cov    pid 1681         .         7         .         .         .         .         :         . 1760
 1 AGENOME-ZPMRG-U-V1.0  100.0% 100.0%     GAGCTAAACCTAGCCCCAAACCCACTCCACCTTACTACCAGACAACCTTAGCCAAACCATTTACCCAAATAAAGTATAGG
 2 AGENOME-ZPMCG-V1.0   100.0%  99.9%     GAGCTAAACCTAGCCCCAAACCCACTCCACCTTACTACCAGACAACCTTAGCCAAACCATTTACCCAAATAAAGTATAGG
 3 AGENOME-ZPMRG-M-V1.0 100.0%  99.8%     GAGCTAAACCTAGCCCCAAACCCACTCCACCTTACTACCAGACAACCTTAGCCAAACCATTTACCCAAATAAAGTATAGG
 4 AGENOME-ZPMRG-F-V1.0  99.9%  99.8%     GAGCTAAACCTAGCCCCAAACCCACTCCACCTTACTACCAGACAACCTTAGCCAAACCATTTACCCAAATAAAGTATAGG
 5 AGENOME-ZPMRG-Z-V1.0  99.9%  99.7%     GAGCTAAACCTAGCCCCAAACCCACTCCACCTTACTACCAGACAACCTTAGCCAAACCATTTACCCAAATAAAGTATAGG
 6 AGENOME-ZPMRG-T-V1.0  99.9%  99.8%     GAGCTAAACCTAGCCCCAAACCCACTCCACCTTACTACCAGACAACCTTAGCCAAACCATTTACCCAAATAAAGTATAGG
 7 AGENOME-ZPMRG-HV-V1.0 99.9%  99.8%     GAGCTAAACCTAGCCCCAAACCCACTCCACCTTACTACCAGACAACCTTAGCCAAACCATTTACCCAAATAAAGTATAGG
 8 AGENOME-ZPMRG-A-V1.0  99.9%  99.8%     GAGCTAAACCTAGCCCCAAACCCACTCCACCTTACTACCAGACAACCTTAGCCAAACCATTTACCCAAATAAAGTATAGG
 9 NC_012920.1           99.9%  99.8%     GAGCTAAACCTAGCCCCAAACCCACTCCACCTTACTACCAGACAACCTTAGCCAAACCATTTACCCAAATAAAGTATAGG
   consensus/100%                       GAGCTAAACCTAGCCCCAAACCCACTCCACCTTACTACCAGACAACCTTAGCCAAACCATTTACCCAAATAAAGTATAGG
   consensus/90%                        GAGCTAAACCTAGCCCCAAACCCACTCCACCTTACTACCAGACAACCTTAGCCAAACCATTTACCCAAATAAAGTATAGG
   consensus/80%                        GAGCTAAACCTAGCCCCAAACCCACTCCACCTTACTACCAGACAACCTTAGCCAAACCATTTACCCAAATAAAGTATAGG
   consensus/70%                        GAGCTAAACCTAGCCCCAAACCCACTCCACCTTACTACCAGACAACCTTAGCCAAACCATTTACCCAAATAAAGTATAGG

                         cov    pid 1761         .         .         .         8         .         .         .         . 1840
 1 AGENOME-ZPMRG-U-V1.0  100.0% 100.0%     CGATAGAAATTGAAACCTGGCGCAATAGATATAGTACCGCAAGGGAAAGATGAAAAATTATAGCCAAGCATAATATAGCA
 2 AGENOME-ZPMCG-V1.0   100.0%  99.9%     CGATAGAAATTGAAACCTGGCGCAATAGATATAGTACCGCAAGGGAAAGATGAAAAATTATAACCAAGCATAATATAGCA
 3 AGENOME-ZPMRG-M-V1.0 100.0%  99.8%     CGATAGAAATTGAAACCTGGCGCAATAGATATAGTACCGCAAGGGAAAGATGAAAAATTATAACCAAGCATAATATAGCA
 4 AGENOME-ZPMRG-F-V1.0  99.9%  99.8%     CGATAGAAATTGAAACCTGGCGCAATAGATATAGTACCGCAAGGGAAAGATGAAAAATTATAACCAAGCATAATATAGCA
 5 AGENOME-ZPMRG-Z-V1.0  99.9%  99.7%     CGATAGAAATTGAAACCTGGCGCAATAGATATAGTACCGCAAGGGAAAGATGAAAAATTATAACCAAGCATAATATAGCA
 6 AGENOME-ZPMRG-T-V1.0  99.9%  99.8%     CGATAGAAATTGAAACCTGGCGCAATAGATATAGTACCGCAAGGGAAAGATGAAAAATTATAACCAAGCATAATATAGCA
 7 AGENOME-ZPMRG-HV-V1.0 99.9%  99.8%     CGATAGAAATTGAAACCTGGCGCAATAGATATAGTACCGCAAGGGAAAGATGAAAAATTATAACCAAGCATAATATAGCA
 8 AGENOME-ZPMRG-A-V1.0  99.9%  99.8%     CGATAGAAATTGAAACCTGGCGCAATAGATATAGTACCGCAAGGGAAAGATGAAAAATTATAACCAAGCATAATATAGCA
 9 NC_012920.1           99.9%  99.8%     CGATAGAAATTGAAACCTGGCGCAATAGATATAGTACCGCAAGGGAAAGATGAAAAATTATAACCAAGCATAATATAGCA
   consensus/100%                       CGATAGAAATTGAAACCTGGCGCAATAGATATAGTACCGCAAGGGAAAGATGAAAAATTATAuCCAAGCATAATATAGCA
   consensus/90%                        CGATAGAAATTGAAACCTGGCGCAATAGATATAGTACCGCAAGGGAAAGATGAAAAATTATAuCCAAGCATAATATAGCA
   consensus/80%                        CGATAGAAATTGAAACCTGGCGCAATAGATATAGTACCGCAAGGGAAAGATGAAAAATTATAACCAAGCATAATATAGCA
   consensus/70%                        CGATAGAAATTGAAACCTGGCGCAATAGATATAGTACCGCAAGGGAAAGATGAAAAATTATAACCAAGCATAATATAGCA

                         cov    pid 1841         :         .         .         .         .         9         .         . 1920
 1 AGENOME-ZPMRG-U-V1.0  100.0% 100.0%     AGGACTAACCCCTATACCTTCTGCATAATGAATTAACTAGAAATAACTTTGCAAGGAGAGCCAAAGCTAAGACCCCCGAA
 2 AGENOME-ZPMCG-V1.0   100.0%  99.9%     AGGACTAACCCCTATACCTTCTGCATAATGAATTAACTAGAAATAACTTTGCAAGGAGAGCCAAAGCTAAGACCCCCGAA
 3 AGENOME-ZPMRG-M-V1.0 100.0%  99.8%     AGGACTAACCCCTATACCTTCTGCATAATGAATTAACTAGAAATAACTTTGCAAGGAGAGCCAAAGCTAAGACCCCCGAA
 4 AGENOME-ZPMRG-F-V1.0  99.9%  99.8%     AGGACTAACCCCTATACCTTCTGCATAATGAATTAACTAGAAATAACTTTGCAAGGAGAGCCAAAGCTAAGACCCCCGAA
 5 AGENOME-ZPMRG-Z-V1.0  99.9%  99.7%     AGGACTAACCCCTATACCTTCTGCATAATGAATTAACTAGAAATAACTTTGCAAGGAGAGCCAAAGCTAAGACCCCCGAA
 6 AGENOME-ZPMRG-T-V1.0  99.9%  99.8%     AGGACTAACCCCTATACCTTCTGCATAATGAATTAACTAGAAATAACTTTGCAAGGAGAACCAAAGCTAAGACCCCCGAA
 7 AGENOME-ZPMRG-HV-V1.0 99.9%  99.8%     AGGACTAACCCCTATACCTTCTGCATAATGAATTAACTAGAAATAACTTTGCAAGGAGAGCCAAAGCTAAGACCCCCGAA
 8 AGENOME-ZPMRG-A-V1.0  99.9%  99.8%     AGGACTAACCCCTATACCTTCTGCATAATGAATTAACTAGAAATAACTTTGCAAGGAGAGCCAAAGCTAAGACCCCCGAA
 9 NC_012920.1           99.9%  99.8%     AGGACTAACCCCTATACCTTCTGCATAATGAATTAACTAGAAATAACTTTGCAAGGAGAGCCAAAGCTAAGACCCCCGAA
   consensus/100%                       AGGACTAACCCCTATACCTTCTGCATAATGAATTAACTAGAAATAACTTTGCAAGGAGAuCCAAAGCTAAGACCCCCGAA
   consensus/90%                        AGGACTAACCCCTATACCTTCTGCATAATGAATTAACTAGAAATAACTTTGCAAGGAGAuCCAAAGCTAAGACCCCCGAA
   consensus/80%                        AGGACTAACCCCTATACCTTCTGCATAATGAATTAACTAGAAATAACTTTGCAAGGAGAGCCAAAGCTAAGACCCCCGAA
```

```
   consensus/70%                                AGGACTAACCCCTATACCTTCTGCATAATGAATTAACTAGAAATAACTTTGCAAGGAGAGCCAAAGCTAAGACCCCCGAA

                              cov    pid   1921          .         .         :         .         .         .         .         0 2000
 1 AGENOME-ZPMRG-U-V1.0  100.0% 100.0%         ACCAGACGAGCTACCTAAGAACAGCTAAAAGAGCACACCCGTCTATGTAGCAAAATAGTGGGAAGATTTATAGGTAGAGG
 2 AGENOME-ZPMCG-V1.0    100.0%  99.9%         ACCAGACGAGCTACCTAAGAACAGCTAAAAGAGCACACCCGTCTATGTAGCAAAATAGTGGGAAGATTTATAGGTAGAGG
 3 AGENOME-ZPMRG-M-V1.0  100.0%  99.8%         ACCAGACGAGCTACCTAAGAACAGCTAAAAGAGCACACCCGTCTATGTAGCAAAATAGTGGGAAGATTTATAGGTAGAGG
 4 AGENOME-ZPMRG-F-V1.0   99.9%  99.8%         ACCAGACGAGCTACCTAAGAACAGCTAAAAGAGCACACCCGTCTATGTAGCAAAATAGTGGGAAGATTTATAGGTAGAGG
 5 AGENOME-ZPMRG-Z-V1.0   99.9%  99.7%         ACCAGACGAGCTACCTAAGAACAGCTAAAAGAGCACACCCGTCTATGTAGCAAAATAGTGGGAAGATTTATAGGTAGAGG
 6 AGENOME-ZPMRG-T-V1.0   99.9%  99.8%         ACCAGACGAGCTACCTAAGAACAGCTAAAAGAGCACACCCGTCTATGTAGCAAAATAGTGGGAAGATTTATAGGTAGAGG
 7 AGENOME-ZPMRG-HV-V1.0  99.9%  99.8%         ACCAGACGAGCTACCTAAGAACAGCTAAAAGAGCACACCCGTCTATGTAGCAAAATAGTGGGAAGATTTATAGGTAGAGG
 8 AGENOME-ZPMRG-A-V1.0   99.9%  99.8%         ACCAGACGAGCTACCTAAGAACAGCTAAAAGAGCACACCCGTCTATGTAGCAAAATAGTGGGAAGATTTATAGGTAGAGG
 9 NC_012920.1            99.9%  99.8%         ACCAGACGAGCTACCTAAGAACAGCTAAAAGAGCACACCCGTCTATGTAGCAAAATAGTGGGAAGATTTATAGGTAGAGG
   consensus/100%                               ACCAGACGAGCTACCTAAGAACAGCTAAAAGAGCACACCCGTCTATGTAGCAAAATAGTGGGAAGATTTATAGGTAGAGG
   consensus/90%                                ACCAGACGAGCTACCTAAGAACAGCTAAAAGAGCACACCCGTCTATGTAGCAAAATAGTGGGAAGATTTATAGGTAGAGG
   consensus/80%                                ACCAGACGAGCTACCTAAGAACAGCTAAAAGAGCACACCCGTCTATGTAGCAAAATAGTGGGAAGATTTATAGGTAGAGG
   consensus/70%                                ACCAGACGAGCTACCTAAGAACAGCTAAAAGAGCACACCCGTCTATGTAGCAAAATAGTGGGAAGATTTATAGGTAGAGG

                              cov    pid   2001          .         .         .         .         :         .         .         . 2080
 1 AGENOME-ZPMRG-U-V1.0  100.0% 100.0%         CGACAAACCTACCGAGCCTGGTGATAGCTGGTTGTCCAAGATAGAATCTTAGTTCAACTTTAAATTTGCCCACAGAACCC
 2 AGENOME-ZPMCG-V1.0    100.0%  99.9%         CGACAAACCTACCGAGCCTGGTGATAGCTGGTTGTCCAAGATAGAATCTTAGTTCAACTTTAAATTTGCCCACAGAACCC
 3 AGENOME-ZPMRG-M-V1.0  100.0%  99.8%         CGACAAACCTACCGAGCCTGGTGATAGCTGGTTGTCCAAGATAGAATCTTAGTTCAACTTTAAATTTGCCCACAGAACCC
 4 AGENOME-ZPMRG-F-V1.0   99.9%  99.8%         CGACAAACCTACCGAGCCTGGTGATAGCTGGTTGTCCAAGATAGAATCTTAGTTCAACTTTAAATTTGCCCACAGAACCC
 5 AGENOME-ZPMRG-Z-V1.0   99.9%  99.7%         CGACAAACCTACCGAGCCTGGTGATAGCTGGTTGTCCAAGATAGAATCTTAGTTCAACTTTAAATTTGCCCACAGAACCC
 6 AGENOME-ZPMRG-T-V1.0   99.9%  99.8%         CGACAAACCTACCGAGCCTGGTGATAGCTGGTTGTCCAAGATAGAATCTTAGTTCAACTTTAAATTTGCCCACAGAACCC
 7 AGENOME-ZPMRG-HV-V1.0  99.9%  99.8%         CGACAAACCTACCGAGCCTGGTGATAGCTGGTTGTCCAAGATAGAATCTTAGTTCAACTTTAAATTTGCCCACAGAACCC
 8 AGENOME-ZPMRG-A-V1.0   99.9%  99.8%         CGACAAACCTACCGAGCCTGGTGATAGCTGGTTGTCCAAGATAGAATCTTAGTTCAACTTTAAATTTGCCCACAGAACCC
 9 NC_012920.1            99.9%  99.8%         CGACAAACCTACCGAGCCTGGTGATAGCTGGTTGTCCAAGATAGAATCTTAGTTCAACTTTAAATTTGCCCACAGAACCC
   consensus/100%                               CGACAAACCTACCGAGCCTGGTGATAGCTGGTTGTCCAAGATAGAATCTTAGTTCAACTTTAAATTTGCCCACAGAACCC
   consensus/90%                                CGACAAACCTACCGAGCCTGGTGATAGCTGGTTGTCCAAGATAGAATCTTAGTTCAACTTTAAATTTGCCCACAGAACCC
   consensus/80%                                CGACAAACCTACCGAGCCTGGTGATAGCTGGTTGTCCAAGATAGAATCTTAGTTCAACTTTAAATTTGCCCACAGAACCC
   consensus/70%                                CGACAAACCTACCGAGCCTGGTGATAGCTGGTTGTCCAAGATAGAATCTTAGTTCAACTTTAAATTTGCCCACAGAACCC

                              cov    pid   2081          .         1         .         .         .         .         :         . 2160
 1 AGENOME-ZPMRG-U-V1.0  100.0% 100.0%         TCTAAATCCCCTTGTAAATTTAACTGTTAGTCCAAAGAGGAACAGCTCTTTGGACACTAGGAAAAAACCTTGTAGAGAGA
 2 AGENOME-ZPMCG-V1.0    100.0%  99.9%         TCTAAATCCCCTTGTAAATTTAACTGTTAGTCCAAAGAGGAACAGCTCTTTGGACACTAGGAAAAAACCTTGTAGAGAGA
 3 AGENOME-ZPMRG-M-V1.0  100.0%  99.8%         TCTAAATCCCCTTGTAAATTTAACTGTTAGTCCAAAGAGGAACAGCTCTTTGGACACTAGGAAAAAACCTTGTAGAGAGA
 4 AGENOME-ZPMRG-F-V1.0   99.9%  99.8%         TCTAAATCCCCTTGTAAATTTAACTGTTAGTCCAAAGAGGAACAGCTCTTTGGACACTAGGAAAAAACCTTGTAGAGAGA
 5 AGENOME-ZPMRG-Z-V1.0   99.9%  99.7%         TCT-AATCCCCTTGTAAATTTAACTGTTAGTCCAAAGAGGAACAGCTCTTTGGACACTAGGAAAAAACCTTGTAGAGAGA
 6 AGENOME-ZPMRG-T-V1.0   99.9%  99.8%         TCTAAATCCCCTTGTAAATTTAACTGTTAGTCCAAAGAGGAACAGCTCTTTGGACACTAGGAAAAAACCTTGTAGAGAGA
 7 AGENOME-ZPMRG-HV-V1.0  99.9%  99.8%         TCTAAATCCCCTTGTAAATTTAACTGTTAGTCCAAAGAGGAACAGCTCTTTGGACACTAGGAAAAAACCTTGTAGAGAGA
 8 AGENOME-ZPMRG-A-V1.0   99.9%  99.8%         TCTAAATCCCCTTGTAAATTTAACTGTTAGTCCAAAGAGGAACAGCTCTTTGGACACTAGGAAAAAACCTTGTAGAGAGA
 9 NC_012920.1            99.9%  99.8%         TCTAAATCCCCTTGTAAATTTAACTGTTAGTCCAAAGAGGAACAGCTCTTTGGACACTAGGAAAAAACCTTGTAGAGAGA
   consensus/100%                               TCT.AATCCCCTTGTAAATTTAACTGTTAGTCCAAAGAGGAACAGCTCTTTGGACACTAGGAAAAAACCTTGTAGAGAGA
   consensus/90%                                TCT.AATCCCCTTGTAAATTTAACTGTTAGTCCAAAGAGGAACAGCTCTTTGGACACTAGGAAAAAACCTTGTAGAGAGA
   consensus/80%                                TCTAAATCCCCTTGTAAATTTAACTGTTAGTCCAAAGAGGAACAGCTCTTTGGACACTAGGAAAAAACCTTGTAGAGAGA
   consensus/70%                                TCTAAATCCCCTTGTAAATTTAACTGTTAGTCCAAAGAGGAACAGCTCTTTGGACACTAGGAAAAAACCTTGTAGAGAGA

                              cov    pid   2161          .         .         .         2         .         .         .         . 2240
 1 AGENOME-ZPMRG-U-V1.0  100.0% 100.0%         GTAAAAAATTTAACACCCATAGTAGGCCTAAAAGCAGCCACCAATTAAGAAAGCGTTCAAGCTCAACACCCACTACCTAA
 2 AGENOME-ZPMCG-V1.0    100.0%  99.9%         GTAAAAAATTTAACACCCATAGTAGGCCTAAAAGCAGCCACCAATTAAGAAAGCGTTCAAGCTCAACACCCACTACCTAA
 3 AGENOME-ZPMRG-M-V1.0  100.0%  99.8%         GTAAAAAATTTAACACCCATAGTAGGCCTAAAAGCAGCCACCAATTAAGAAAGCGTTCAAGCTCAACACCCACTACCTAA
 4 AGENOME-ZPMRG-F-V1.0   99.9%  99.8%         GTAAAAAATTTAACACCCATAGTAGGCCTAAAAGCAGCCACCAATTAAGAAAGCGTTCAAGCTCAACACCCACTACCTAA
 5 AGENOME-ZPMRG-Z-V1.0   99.9%  99.7%         GTAAAAAATTTAACACCCATAGTAGGCCTAAAAGCAGCCACCAATTAAGAAAGCGTTCAAGCTCAACACCCACTACCTAA
 6 AGENOME-ZPMRG-T-V1.0   99.9%  99.8%         GTAAAAAATTTAACACCCATAGTAGGCCTAAAAGCAGCCACCAATTAAGAAAGCGTTCAAGCTCAACACCCACTACCTAA
 7 AGENOME-ZPMRG-HV-V1.0  99.9%  99.8%         GTAAAAAATTTAACACCCATAGTAGGCCTAAAAGCAGCCACCAATTAAGAAAGCGTTCAAGCTCAACACCCACTACCTAA
 8 AGENOME-ZPMRG-A-V1.0   99.9%  99.8%         GTAAAAAATTTAACACCCATAGTAGGCCTAAAAGCAGCCACCAATTAAGAAAGCGTTCAAGCTCAACACCCACTACCTAA
 9 NC_012920.1            99.9%  99.8%         GTAAAAAATTTAACACCCATAGTAGGCCTAAAAGCAGCCACCAATTAAGAAAGCGTTCAAGCTCAACACCCACTACCTAA
   consensus/100%                               GTAAAAAATTTAACACCCATAGTAGGCCTAAAAGCAGCCACCAATTAAGAAAGCGTTCAAGCTCAACACCCACTACCTAA
   consensus/90%                                GTAAAAAATTTAACACCCATAGTAGGCCTAAAAGCAGCCACCAATTAAGAAAGCGTTCAAGCTCAACACCCACTACCTAA
   consensus/80%                                GTAAAAAATTTAACACCCATAGTAGGCCTAAAAGCAGCCACCAATTAAGAAAGCGTTCAAGCTCAACACCCACTACCTAA
   consensus/70%                                GTAAAAAATTTAACACCCATAGTAGGCCTAAAAGCAGCCACCAATTAAGAAAGCGTTCAAGCTCAACACCCACTACCTAA

                              cov    pid   2241          :         .         .         .         .         3         .         . 2320
 1 AGENOME-ZPMRG-U-V1.0  100.0% 100.0%         AAAATCCCAAACATATAACTGAACTCCTCACACCCAATTGGACCAATCTATCACCCTATAGAAGAACTAATGTTAGTATA
 2 AGENOME-ZPMCG-V1.0    100.0%  99.9%         AAAATCCCAAACATATAACTGAACTCCTCACACCCAATTGGACCAATCTATCACCCTATAGAAGAACTAATGTTAGTATA
 3 AGENOME-ZPMRG-M-V1.0  100.0%  99.8%         AAAATCCCAAACATATAACTGAACTCCTCACACCCAATTGGACCAATCTATCACCCTATAGAAGAACTAATGTTAGTATA
 4 AGENOME-ZPMRG-F-V1.0   99.9%  99.8%         AAAATCCCAAACATATAACTGAACTCCTCACACCCAATTGGACCAATCTATCACCCTATAGAAGAACTAATGTTAGTATA
 5 AGENOME-ZPMRG-Z-V1.0   99.9%  99.7%         AAAATCCCAAACATATAACTGAACTCCTCACACCCAATTGGACCAATCTATCACCCTATAGAAGAACTAATGTTAGTATA
 6 AGENOME-ZPMRG-T-V1.0   99.9%  99.8%         AAAATCCCAAACATATAACTGAACTCCTCACACCCAATTGGACCAATCTATCACCCTATAGAAGAACTAATGTTAGTATA
 7 AGENOME-ZPMRG-HV-V1.0  99.9%  99.8%         AAAATCCCAAACATATAACTGAACTCCTCACACCCAATTGGACCAATCTATCACCCTATAGAAGAACTAATGTTAGTATA
 8 AGENOME-ZPMRG-A-V1.0   99.9%  99.8%         AAAATCCCAAACATATAACTGAACTCCTCACACCCAATTGGACCAATCTATCACCCTATAGAAGAACTAATGTTAGTATA
 9 NC_012920.1            99.9%  99.8%         AAAATCCCAAACATATAACTGAACTCCTCACACCCAATTGGACCAATCTATCACCCTATAGAAGAACTAATGTTAGTATA
   consensus/100%                               AAAATCCCAAACATATAACTGAACTCCTCACACCCAATTGGACCAATCTATCACCCTATAGAAGAACTAATGTTAGTATA
   consensus/90%                                AAAATCCCAAACATATAACTGAACTCCTCACACCCAATTGGACCAATCTATCACCCTATAGAAGAACTAATGTTAGTATA
   consensus/80%                                AAAATCCCAAACATATAACTGAACTCCTCACACCCAATTGGACCAATCTATCACCCTATAGAAGAACTAATGTTAGTATA
   consensus/70%                                AAAATCCCAAACATATAACTGAACTCCTCACACCCAATTGGACCAATCTATCACCCTATAGAAGAACTAATGTTAGTATA

                              cov    pid   2321          .         .         :         .         .         .         .         4 2400
 1 AGENOME-ZPMRG-U-V1.0  100.0% 100.0%         AGTAACATGAAAACATTCTCCTCCGCATAAGCCTGCGTCAGATTAAAACACTGAACTGACAATTAACAGCCCAATATCTA
 2 AGENOME-ZPMCG-V1.0    100.0%  99.9%         AGTAACATGAAAACATTCTCCTCCGCATAAGCCTGCGTCAGATTAAAACACTGAACTGACAATTAACAGCCCAATATCTA
 3 AGENOME-ZPMRG-M-V1.0  100.0%  99.8%         AGTAACATGAAAACATTCTCCTCCGCATAAGCCTGCGTCAGATTAAAACACTGAACTGACAATTAACAGCCCAATATCTA
 4 AGENOME-ZPMRG-F-V1.0   99.9%  99.8%         AGTAACATGAAAACATTCTCCTCCGCATAAGCCTGCGTCAGATTAAAACACTGAACTGACAATTAACAGCCCAATATCTA
 5 AGENOME-ZPMRG-Z-V1.0   99.9%  99.7%         AGTAACATGAAAACATTCTCCTCCGCATAAGCCTGCGTCAGATTAAAACACTGAACTGACAATTAACAGCCCAATATCTA
 6 AGENOME-ZPMRG-T-V1.0   99.9%  99.8%         AGTAACATGAAAACATTCTCCTCCGCATAAGCCTGCGTCAGATTAAAACACTGAACTGACAATTAACAGCCCAATATCTA
 7 AGENOME-ZPMRG-HV-V1.0  99.9%  99.8%         AGTAACATGAAAACATTCTCCTCCGCATAAGCCTGCGTCAGATTAAAACACTGAACTGACAATTAACAGCCCAATATCTA
 8 AGENOME-ZPMRG-A-V1.0   99.9%  99.8%         AGTAACATGAAAACATTCTCCTCCGCATAAGCCTGCGTCAGATTAAAACACTGAACTGACAATTAACAGCCCAATATCTA
 9 NC_012920.1            99.9%  99.8%         AGTAACATGAAAACATTCTCCTCCGCATAAGCCTGCGTCAGATTAAAACACTGAACTGACAATTAACAGCCCAATATCTA
   consensus/100%                               AGTAACATGAAAACATTCTCCTCCGCATAAGCCTGCGTCAGATTAAAACACTGAACTGACAATTAACAGCCCAATATCTA
   consensus/90%                                AGTAACATGAAAACATTCTCCTCCGCATAAGCCTGCGTCAGATTAAAACACTGAACTGACAATTAACAGCCCAATATCTA
   consensus/80%                                AGTAACATGAAAACATTCTCCTCCGCATAAGCCTGCGTCAGATTAAAACACTGAACTGACAATTAACAGCCCAATATCTA
   consensus/70%                                AGTAACATGAAAACATTCTCCTCCGCATAAGCCTGCGTCAGATTAAAACACTGAACTGACAATTAACAGCCCAATATCTA
```

```
                              cov    pid  2401         .         .         .         .         :         .         .         . 2480
1 AGENOME-ZPMRG-U-V1.0   100.0% 100.0%       CAATCAACCAACAAGTCATTATTACCCTCACTGTCAACCCAACACAGGCATGCTCATAAGGAAAGGTTAAAAAAAGTAAA
2 AGENOME-ZPMCG-V1.0     100.0%  99.9%       CAATCAACCAACAAGTCATTATTACCCTCACTGTCAACCCAACACAGGCATGCTCATAAGGAAAGGTTAAAAAAAGTAAA
3 AGENOME-ZPMRG-M-V1.0   100.0%  99.8%       CAATCAACCAACAAGTCATTATTACCCTCACTGTCAACCCAACACAGGCATGCTCATAAGGAAAGGTTAAAAAAAGTAAA
4 AGENOME-ZPMRG-F-V1.0    99.9%  99.8%       TAATCAACCAACAAGTCATTATTACCCTCACTGTCAACCCAACACAGGCATGCTCATAAGGAAAGGTTAAAAAAAGTAAA
5 AGENOME-ZPMRG-Z-V1.0    99.9%  99.7%       CAATCAACCAACAAGTCATTATTACCCTCACTGTCAACCCAACACAGGCATGCTCATAAGGAAAGGTTAAAAAAAGTAAA
6 AGENOME-ZPMRG-T-V1.0    99.9%  99.8%       CAATCAACCAACAAGTCATTATTACCCTCACTGTCAACCCAACACAGGCATGCTCATAAGGAAAGGTTAAAAAAAGTAAA
7 AGENOME-ZPMRG-HV-V1.0   99.9%  99.8%       CAATCAACCAACAAGTCATTATTACCCTCACTGTCAACCCAACACAGGCATGCTCATAAGGAAAGGTTAAAAAAAGTAAA
8 AGENOME-ZPMRG-A-V1.0    99.9%  99.8%       CAATCAACCAACAAGTCATTATTACCCTCACTGTCAACCCAACACAGGCATGCTCATAAGGAAAGGTTAAAAAAAGTAAA
9 NC_012920.1             99.9%  99.8%       CAATCAACCAACAAGTCATTATTACCCTCACTGTCAACCCAACACAGGCATGCTCATAAGGAAAGGTTAAAAAAAGTAAA
  consensus/100%                             sAATCAACCAACAAGTCATTATTACCCTCACTGTCAACCCAACACAGGCATGCTCATAAGGAAAGGTTAAAAAAAGTAAA
  consensus/90%                              sAATCAACCAACAAGTCATTATTACCCTCACTGTCAACCCAACACAGGCATGCTCATAAGGAAAGGTTAAAAAAAGTAAA
  consensus/80%                              CAATCAACCAACAAGTCATTATTACCCTCACTGTCAACCCAACACAGGCATGCTCATAAGGAAAGGTTAAAAAAAGTAAA
  consensus/70%                              CAATCAACCAACAAGTCATTATTACCCTCACTGTCAACCCAACACAGGCATGCTCATAAGGAAAGGTTAAAAAAAGTAAA

                              cov    pid  2481         .         5         .         .         .         .         :         . 2560
1 AGENOME-ZPMRG-U-V1.0   100.0% 100.0%       AGGAACTCGGCAAATCTTACCCCGCCTGTTTACCAAAAACATCACCTCTAGCATCACCAGTATTAGAGGCACCGCCTGCC
2 AGENOME-ZPMCG-V1.0     100.0%  99.9%       AGGAACTCGGCAAATCTTACCCCGCCTGTTTACCAAAAACATCACCTCTAGCATCACCAGTATTAGAGGCACCGCCTGCC
3 AGENOME-ZPMRG-M-V1.0   100.0%  99.8%       AGGAACTCGGCAAATCTTACCCCGCCTGTTTACCAAAAACATCACCTCTAGCATCACCAGTATTAGAGGCACCGCCTGCC
4 AGENOME-ZPMRG-F-V1.0    99.9%  99.8%       AGGAACTCGGCAAATCTTACCCCGCCTGTTTACCAAAAACATCACCTCTAGCATCACCAGTATTAGAGGCACCGCCTGCC
5 AGENOME-ZPMRG-Z-V1.0    99.9%  99.7%       AGGAACTCGGCAAATCTTACCCCGCCTGTTTACCAAAAACATCACCTCTAGCATCACCAGTATTAGAGGCACCGCCTGCC
6 AGENOME-ZPMRG-T-V1.0    99.9%  99.8%       AGGAACTCGGCAAATCTTACCCCGCCTGTTTACCAAAAACATCACCTCTAGCATCACCAGTATTAGAGGCACCGCCTGCC
7 AGENOME-ZPMRG-HV-V1.0   99.9%  99.8%       AGGAACTCGGCAAATCTTACCCCGCCTGTTTACCAAAAACATCACCTCTAGCATCACCAGTATTAGAGGCACCGCCTGCC
8 AGENOME-ZPMRG-A-V1.0    99.9%  99.8%       AGGAACTCGGCAAATCTTACCCCGCCTGTTTACCAAAAACATCACCTCTAGCATCACCAGTATTAGAGGCACCGCCTGCC
9 NC_012920.1             99.9%  99.8%       AGGAACTCGGCAAATCTTACCCCGCCTGTTTACCAAAAACATCACCTCTAGCATCACCAGTATTAGAGGCACCGCCTGCC
  consensus/100%                             AGGAACTCGGCAAATCTTACCCCGCCTGTTTACCAAAAACATCACCTCTAGCATCACCAGTATTAGAGGCACCGCCTGCC
  consensus/90%                              AGGAACTCGGCAAATCTTACCCCGCCTGTTTACCAAAAACATCACCTCTAGCATCACCAGTATTAGAGGCACCGCCTGCC
  consensus/80%                              AGGAACTCGGCAAATCTTACCCCGCCTGTTTACCAAAAACATCACCTCTAGCATCACCAGTATTAGAGGCACCGCCTGCC
  consensus/70%                              AGGAACTCGGCAAATCTTACCCCGCCTGTTTACCAAAAACATCACCTCTAGCATCACCAGTATTAGAGGCACCGCCTGCC

                              cov    pid  2561         .         .         .         6         .         .         .         . 2640
1 AGENOME-ZPMRG-U-V1.0   100.0% 100.0%       CAGTGACACATGTTTAACGGCCGCGGTACCCTAACCGTGCAAAGGTAGCATAATCACTTGTTCCTTAAATAGGGACCTGT
2 AGENOME-ZPMCG-V1.0     100.0%  99.9%       CAGTGACACATGTTTAACGGCCGCGGTACCCTAACCGTGCAAAGGTAGCATAATCACTTGTTCCTTAAATAGGGACCTGT
3 AGENOME-ZPMRG-M-V1.0   100.0%  99.8%       CAGTGACACATGTTTAACGGCCGCGGTACCCTAACCGTGCAAAGGTAGCATAATCACTTGTTCCTTAAATAGGGACCTGT
4 AGENOME-ZPMRG-F-V1.0    99.9%  99.8%       CAGTGACACATGTTTAACGGCCGCGGTACCCTAACCGTGCAAAGGTAGCATAATCACTTGTTCCTTAAATAGGGACCTGT
5 AGENOME-ZPMRG-Z-V1.0    99.9%  99.7%       CAGTGACACATGTTTAACGGCCGCGGTACCCTAACCGTGCAAAGGTAGCATAATCACTTGTTCCTTAAATAGGGACCTGT
6 AGENOME-ZPMRG-T-V1.0    99.9%  99.8%       CAGTGACACATGTTTAACGGCCGCGGTACCCTAACCGTGCAAAGGTAGCATAATCACTTGTTCCTTAAATAGGGACCTGT
7 AGENOME-ZPMRG-HV-V1.0   99.9%  99.8%       CAGTGACACATGTTTAACGGCCGCGGTACCCTAACCGTGCAAAGGTAGCATAATCACTTGTTCCTTAAATAGGGACCTGT
8 AGENOME-ZPMRG-A-V1.0    99.9%  99.8%       CAGTGACACATGTTTAACGGCCGCGGTACCCTAACCGTGCAAAGGTAGCATAATCACTTGTTCCTTAAATAGGGACCTGT
9 NC_012920.1             99.9%  99.8%       CAGTGACACATGTTTAACGGCCGCGGTACCCTAACCGTGCAAAGGTAGCATAATCACTTGTTCCTTAAATAGGGACCTGT
  consensus/100%                             CAGTGACACATGTTTAACGGCCGCGGTACCCTAACCGTGCAAAGGTAGCATAATCACTTGTTCCTTAAATAGGGACCTGT
  consensus/90%                              CAGTGACACATGTTTAACGGCCGCGGTACCCTAACCGTGCAAAGGTAGCATAATCACTTGTTCCTTAAATAGGGACCTGT
  consensus/80%                              CAGTGACACATGTTTAACGGCCGCGGTACCCTAACCGTGCAAAGGTAGCATAATCACTTGTTCCTTAAATAGGGACCTGT
  consensus/70%                              CAGTGACACATGTTTAACGGCCGCGGTACCCTAACCGTGCAAAGGTAGCATAATCACTTGTTCCTTAAATAGGGACCTGT

                              cov    pid  2641         :         .         .         .         .         7         .         . 2720
1 AGENOME-ZPMRG-U-V1.0   100.0% 100.0%       ATGAATGGCTCCACGAGGGTTCAGCTGTCTCTTACTTTTAACCAGTGAAATTGACCTGCCCGTGAAGAGGCGGGCATGAC
2 AGENOME-ZPMCG-V1.0     100.0%  99.9%       ATGAATGGCTCCACGAGGGTTCAGCTGTCTCTTACTTTTAACCAGTGAAATTGACCTGCCCGTGAAGAGGCGGGCATGAC
3 AGENOME-ZPMRG-M-V1.0   100.0%  99.8%       ATGAATGGCTCCACGAGGGTTCAGCTGTCTCTTACTTTTAACCAGTGAAATTGACCTGCCCGTGAAGAGGCGGGCATGAC
4 AGENOME-ZPMRG-F-V1.0    99.9%  99.8%       ATGAATGGCTCCACGAGGGTTCAGCTGTCTCTTACTTTTAACCAGTGAAATTGACCTGCCCGTGAAGAGGCGGGCATGAC
5 AGENOME-ZPMRG-Z-V1.0    99.9%  99.7%       ATGAATGGCTCCACGAGGGTTCAGCTGTCTCTTACTTTTAACCAGTGAAATTGACCTGCCCGTGAAGAGGCGGGCATGAC
6 AGENOME-ZPMRG-T-V1.0    99.9%  99.8%       ATGAATGGCTCCACGAGGGTTCAGCTGTCTCTTACTTTTAACCAGTGAAATTGACCTGCCCGTGAAGAGGCGGGCATGAC
7 AGENOME-ZPMRG-HV-V1.0   99.9%  99.8%       ATGAATGGCTCCACGAGGGTTCAGCTGTCTCTTACTTTTAACCAGTGAAATTGACCTGCCCGTGAAGAGGCGGGCATGAC
8 AGENOME-ZPMRG-A-V1.0    99.9%  99.8%       ATGAATGGCTCCACGAGGGTTCAGCTGTCTCTTACTTTTAACCAGTGAAATTGACCTGCCCGTGAAGAGGCGGGCATAAC
9 NC_012920.1             99.9%  99.8%       ATGAATGGCTCCACGAGGGTTCAGCTGTCTCTTACTTTTAACCAGTGAAATTGACCTGCCCGTGAAGAGGCGGGCATAAC
  consensus/100%                             ATGAATGGCTCCACGAGGGTTCAGCTGTCTCTTACTTTTAACCAGTGAAATTGACCTGCCCGTGAAGAGGCGGGCATuAC
  consensus/90%                              ATGAATGGCTCCACGAGGGTTCAGCTGTCTCTTACTTTTAACCAGTGAAATTGACCTGCCCGTGAAGAGGCGGGCATuAC
  consensus/80%                              ATGAATGGCTCCACGAGGGTTCAGCTGTCTCTTACTTTTAACCAGTGAAATTGACCTGCCCGTGAAGAGGCGGGCATuAC
  consensus/70%                              ATGAATGGCTCCACGAGGGTTCAGCTGTCTCTTACTTTTAACCAGTGAAATTGACCTGCCCGTGAAGAGGCGGGCATGAC

                              cov    pid  2721         .         .         :         .         .         .         .         8 2800
1 AGENOME-ZPMRG-U-V1.0   100.0% 100.0%       ACAGCAAGACGAGAAGACCCTATGGAGCTTTAATTTATTAATGCAAACAGTACCTAACAAACCCACAGGTCCTAAACTAC
2 AGENOME-ZPMCG-V1.0     100.0%  99.9%       ACAGCAAGACGAGAAGACCCTATGGAGCTTTAATTTATTAATGCAAACAGTACCTAACAAACCCACAGGTCCTAAACTAC
3 AGENOME-ZPMRG-M-V1.0   100.0%  99.8%       ACAGCAAGACGAGAAGACCCTATGGAGCTTTAATTTATTAATGCAAACAGTACCTAACAAACCCACAGGTCCTAAACTAC
4 AGENOME-ZPMRG-F-V1.0    99.9%  99.8%       ACAGCAAGACGAGAAGACCCTATGGAGCTTTAATTTATTAATGCAAACAGTACCTAACAAACCCACAGGTCCTAAACTAC
5 AGENOME-ZPMRG-Z-V1.0    99.9%  99.7%       ACAGCAAGACGAGAAGACCCTATGGAGCTTTAATTTATTAATGCAAACAGTACCTAACAAACCCACAGGTCCTAAACTAC
6 AGENOME-ZPMRG-T-V1.0    99.9%  99.8%       ACAGCAAGACGAGAAGACCCTATGGAGCTTTAATTTATTAATGCAAACAGTACCTAACAAACCCACAGGTCCTAAACTAC
7 AGENOME-ZPMRG-HV-V1.0   99.9%  99.8%       ACAGCAAGACGAGAAGACCCTATGGAGCTTTAATTTATTAATGCAAACAGTACCTAACAAACCCACAGGTCCTAAACTAC
8 AGENOME-ZPMRG-A-V1.0    99.9%  99.8%       ACAGCAAGACGAGAAGACCCTATGGAGCTTTAATTTATTAATGCAAACAGTACCTAACAAACCCACAGGTCCTAAACTAC
9 NC_012920.1             99.9%  99.8%       ACAGCAAGACGAGAAGACCCTATGGAGCTTTAATTTATTAATGCAAACAGTACCTAACAAACCCACAGGTCCTAAACTAC
  consensus/100%                             ACAGCAAGACGAGAAGACCCTATGGAGCTTTAATTTATTAATGCAAACAGTACCTAACAAACCCACAGGTCCTAAACTAC
  consensus/90%                              ACAGCAAGACGAGAAGACCCTATGGAGCTTTAATTTATTAATGCAAACAGTACCTAACAAACCCACAGGTCCTAAACTAC
  consensus/80%                              ACAGCAAGACGAGAAGACCCTATGGAGCTTTAATTTATTAATGCAAACAGTACCTAACAAACCCACAGGTCCTAAACTAC
  consensus/70%                              ACAGCAAGACGAGAAGACCCTATGGAGCTTTAATTTATTAATGCAAACAGTACCTAACAAACCCACAGGTCCTAAACTAC

                              cov    pid  2801         .         .         .         .         :         .         .         . 2880
1 AGENOME-ZPMRG-U-V1.0   100.0% 100.0%       CAAACCTGCATTAAAAATTTCGGTTGGGGCGACCTCGGAGCAGAACCCAACCTCCGAGCAGTACATGCTAAGACTTCACC
2 AGENOME-ZPMCG-V1.0     100.0%  99.9%       CAAACCTGCATTAAAAATTTCGGTTGGGGCGACCTCGGAGCAGAACCCAACCTCCGAGCAGTACATGCTAAGACTTCACC
3 AGENOME-ZPMRG-M-V1.0   100.0%  99.8%       CAAACCTGCATTAAAAATTTCGGTTGGGGCGACCTCGGAGCAGAACCCAACCTCCGAGCAGTACATGCTAAGACTTCACC
4 AGENOME-ZPMRG-F-V1.0    99.9%  99.8%       CAAACCTGCATTAAAAATTTCGGTTGGGGCGACCTCGGAGCAGAACCCAACCTCCGAGCAGTACATGCTAAGACTTCACC
5 AGENOME-ZPMRG-Z-V1.0    99.9%  99.7%       CAAACCTGCATTAAAAATTTCGGTTGGGGCGACCTCGGAGCAGAACCCAACCTCCGAGCAGTACATGCTAAGACTTCACC
6 AGENOME-ZPMRG-T-V1.0    99.9%  99.8%       CAAACCTGCATTAAAAATTTCGGTTGGGGCGACCTCGGAGCAGAACCCAACCTCCGAGCAGTACATGCTAAGACTTCACC
7 AGENOME-ZPMRG-HV-V1.0   99.9%  99.8%       CAAACCTGCATTAAAAATTTCGGTTGGGGCGACCTCGGAGCAGAACCCAACCTCCGAGCAGTACATGCTAAGACTTCACC
8 AGENOME-ZPMRG-A-V1.0    99.9%  99.8%       CAAACCTGCATTAAAAATTTCGGTTGGGGCGACCTCGGAGCAGAACCCAACCTCCGAGCAGTACATGCTAAGACTTCACC
9 NC_012920.1             99.9%  99.8%       CAAACCTGCATTAAAAATTTCGGTTGGGGCGACCTCGGAGCAGAACCCAACCTCCGAGCAGTACATGCTAAGACTTCACC
  consensus/100%                             CAAACCTGCATTAAAAATTTCGGTTGGGGCGACCTCGGAGCAGAACCCAACCTCCGAGCAGTACATGCTAAGACTTCACC
  consensus/90%                              CAAACCTGCATTAAAAATTTCGGTTGGGGCGACCTCGGAGCAGAACCCAACCTCCGAGCAGTACATGCTAAGACTTCACC
  consensus/80%                              CAAACCTGCATTAAAAATTTCGGTTGGGGCGACCTCGGAGCAGAACCCAACCTCCGAGCAGTACATGCTAAGACTTCACC
  consensus/70%                              CAAACCTGCATTAAAAATTTCGGTTGGGGCGACCTCGGAGCAGAACCCAACCTCCGAGCAGTACATGCTAAGACTTCACC
```

```
                              cov    pid  2881          .         9         .         .         .         .         :         . 2960
  1 AGENOME-ZPMRG-U-V1.0  100.0% 100.0%       AGTCAAAGCGAACTACTATACTCAATTGATCCAATAACTTGACCAACGGAACAAGTTACCCTAGGGATAACAGCGCAATC
  2 AGENOME-ZPMCG-V1.0    100.0%  99.9%       AGTCAAAGCGAACTACTATACTCAATTGATCCAATAACTTGACCAACGGAACAAGTTACCCTAGGGATAACAGCGCAATC
  3 AGENOME-ZPMRG-M-V1.0  100.0%  99.8%       AGTCAAAGCGAACTACTATACTCAATTGATCCAATAACTTGACCAACGGAACAAGTTACCCTAGGGATAACAGCGCAATC
  4 AGENOME-ZPMRG-F-V1.0   99.9%  99.8%       AGTCAAAGCGAACTACTATACTCAATTGATCCAATAACTTGACCAACGGAACAAGTTACCCTAGGGATAACAGCGCAATC
  5 AGENOME-ZPMRG-Z-V1.0   99.9%  99.7%       AGTCAAAGCGAACTACTATACTCAATTGATCCAATAACTTGACCAACGGAACAAGTTACCCTAGGGATAACAGCGCAATC
  6 AGENOME-ZPMRG-T-V1.0   99.9%  99.8%       AGTCAAAGCGAACTACTATACTCAATTGATCCAATAACTTGACCAACGGAACAAGTTACCCTAGGGATAACAGCGCAATC
  7 AGENOME-ZPMRG-HV-V1.0  99.9%  99.8%       AGTCAAAGCGAACTACTATACTCAATTGATCCAATAACTTGACCAACGGAACAAGTTACCCTAGGGATAACAGCGCAATC
  8 AGENOME-ZPMRG-A-V1.0   99.9%  99.8%       AGTCAAAGCGAACTACTATACTCAATTGATCCAATAACTTGACCAACGGAACAAGTTACCCTAGGGATAACAGCGCAATC
  9 NC_012920.1            99.9%  99.8%       AGTCAAAGCGAACTACTATACTCAATTGATCCAATAACTTGACCAACGGAACAAGTTACCCTAGGGATAACAGCGCAATC
    consensus/100%                            AGTCAAAGCGAACTACTATACTCAATTGATCCAATAACTTGACCAACGGAACAAGTTACCCTAGGGATAACAGCGCAATC
    consensus/90%                             AGTCAAAGCGAACTACTATACTCAATTGATCCAATAACTTGACCAACGGAACAAGTTACCCTAGGGATAACAGCGCAATC
    consensus/80%                             AGTCAAAGCGAACTACTATACTCAATTGATCCAATAACTTGACCAACGGAACAAGTTACCCTAGGGATAACAGCGCAATC
    consensus/70%                             AGTCAAAGCGAACTACTATACTCAATTGATCCAATAACTTGACCAACGGAACAAGTTACCCTAGGGATAACAGCGCAATC

                              cov    pid  2961          .         .         .         0         .         .         .         . 3040
  1 AGENOME-ZPMRG-U-V1.0  100.0% 100.0%       CTATTCTAGAGTCCATATCAACAATAGGGTTTACGACCTCGATGTTGGATCAGGACATCCCGATGGTGCAGCCGCTATTA
  2 AGENOME-ZPMCG-V1.0    100.0%  99.9%       CTATTCTAGAGTCCATATCAACAATAGGGTTTACGACCTCGATGTTGGATCAGGACATCCCGATGGTGCAGCCGCTATTA
  3 AGENOME-ZPMRG-M-V1.0  100.0%  99.8%       CTATTCTAGAGTCCATATCAACAATAGGGTTTACGACCTCGATGTTGGATCAGGACATCCCGATGGTGCAGCCGCTATTA
  4 AGENOME-ZPMRG-F-V1.0   99.9%  99.8%       CTATTCTAGAGTCCATATCAACAATAGGGTTTACGACCTCGATGTTGGATCAGGACATCCCGATGGTGCAGCCGCTATTA
  5 AGENOME-ZPMRG-Z-V1.0   99.9%  99.7%       CTATTCTAGAGTCCATATCAACAATAGGGTTTACGACCTCGATGTTGGATCAGGACATCCCGATGGTGCAGCCGCTATTA
  6 AGENOME-ZPMRG-T-V1.0   99.9%  99.8%       CTATTCTAGAGTCCATATCAACAATAGGGTTTACGACCTCGATGTTGGATCAGGACATCCCGATGGTGCAGCCGCTATTA
  7 AGENOME-ZPMRG-HV-V1.0  99.9%  99.8%       CTATTCTAGAGTCCATATCAACAATAGGGTTTACGACCTCGATGTTGGATCAGGACATCCCGATGGTGCAGCCGCTATTA
  8 AGENOME-ZPMRG-A-V1.0   99.9%  99.8%       CTATTCTAGAGTCCATATCAACAATAGGGTTTACGACCTCGATGTTGGATCAGGACATCCCGATGGTGCAGCCGCTATTA
  9 NC_012920.1            99.9%  99.8%       CTATTCTAGAGTCCATATCAACAATAGGGTTTACGACCTCGATGTTGGATCAGGACATCCCGATGGTGCAGCCGCTATTA
    consensus/100%                            CTATTCTAGAGTCCATATCAACAATAGGGTTTACGACCTCGATGTTGGATCAGGACATCCCGATGGTGCAGCCGCTATTA
    consensus/90%                             CTATTCTAGAGTCCATATCAACAATAGGGTTTACGACCTCGATGTTGGATCAGGACATCCCGATGGTGCAGCCGCTATTA
    consensus/80%                             CTATTCTAGAGTCCATATCAACAATAGGGTTTACGACCTCGATGTTGGATCAGGACATCCCGATGGTGCAGCCGCTATTA
    consensus/70%                             CTATTCTAGAGTCCATATCAACAATAGGGTTTACGACCTCGATGTTGGATCAGGACATCCCGATGGTGCAGCCGCTATTA

                              cov    pid  3041          :         .         .         .         .         1         .         . 3120
  1 AGENOME-ZPMRG-U-V1.0  100.0% 100.0%       AAGGTTCGTTTGTTCAACGATTAAAGTCCTACGTGATCTGAGTTCAGACCGGAGTAATCCAGGTCGGTTTCTATCTAC-T
  2 AGENOME-ZPMCG-V1.0    100.0%  99.9%       AAGGTTCGTTTGTTCAACGATTAAAGTCCTACGTGATCTGAGTTCAGACCGGAGTAATCCAGGTCGGTTTCTATCTAC-T
  3 AGENOME-ZPMRG-M-V1.0  100.0%  99.8%       AAGGTTCGTTTGTTCAACGATTAAAGTCCTACGTGATCTGAGTTCAGACCGGAGTAATCCAGGTCGGTTTCTATCTAC-T
  4 AGENOME-ZPMRG-F-V1.0   99.9%  99.8%       AAGGTTCGTTTGTTCAACGATTAAAGTCCTACGTGATCTGAGTTCAGACCGGAGTAATCCAGGTCGGTTTCTATCTAC-T
  5 AGENOME-ZPMRG-Z-V1.0   99.9%  99.7%       AAGGTTCGTTTGTTCAACGATTAAAGTCCTACGTGATCTGAGTTCAGACCGGAGTAATCCAGGTCGGTTTCTATCTAC-T
  6 AGENOME-ZPMRG-T-V1.0   99.9%  99.8%       AAGGTTCGTTTGTTCAACGATTAAAGTCCTACGTGATCTGAGTTCAGACCGGAGTAATCCAGGTCGGTTTCTATCTAC-T
  7 AGENOME-ZPMRG-HV-V1.0  99.9%  99.8%       AAGGTTCGTTTGTTCAACGATTAAAGTCCTACGTGATCTGAGTTCAGACCGGAGTAATCCAGGTCGGTTTCTATCTAC-T
  8 AGENOME-ZPMRG-A-V1.0   99.9%  99.8%       AAGGTTCGTTTGTTCAACGATTAAAGTCCTACGTGATCTGAGTTCAGACCGGAGTAATCCAGGTCGGTTTCTATCTAC-T
  9 NC_012920.1            99.9%  99.8%       AAGGTTCGTTTGTTCAACGATTAAAGTCCTACGTGATCTGAGTTCAGACCGGAGTAATCCAGGTCGGTTTCTATCTACNT
    consensus/100%                            AAGGTTCGTTTGTTCAACGATTAAAGTCCTACGTGATCTGAGTTCAGACCGGAGTAATCCAGGTCGGTTTCTATCTAC.T
    consensus/90%                             AAGGTTCGTTTGTTCAACGATTAAAGTCCTACGTGATCTGAGTTCAGACCGGAGTAATCCAGGTCGGTTTCTATCTAC.T
    consensus/80%                             AAGGTTCGTTTGTTCAACGATTAAAGTCCTACGTGATCTGAGTTCAGACCGGAGTAATCCAGGTCGGTTTCTATCTAC.T
    consensus/70%                             AAGGTTCGTTTGTTCAACGATTAAAGTCCTACGTGATCTGAGTTCAGACCGGAGTAATCCAGGTCGGTTTCTATCTAC.T

                              cov    pid  3121          .         .         :         .         .         .         .         2 3200
  1 AGENOME-ZPMRG-U-V1.0  100.0% 100.0%       TCAAATTCCTCCCTGTACGAAAGGACAAGAGAAATAAGGCCTACTTCACATAAGCGCCTTCCCCCGTAAATGATATCATC
  2 AGENOME-ZPMCG-V1.0    100.0%  99.9%       TCAAATTCCTCCCTGTACGAAAGGACAAGAGAAATAAGGCCTACTTCACATAAGCGCCTTCCCCCGTAAATGATATCATC
  3 AGENOME-ZPMRG-M-V1.0  100.0%  99.8%       TCAAATTCCTCCCTGTACGAAAGGACAAGAGAAATAAGGCCTACTTCACA-AAGCGCCTTCCCCCGTAAATGATATCATC
  4 AGENOME-ZPMRG-F-V1.0   99.9%  99.8%       TCAAATTCCTCCCTGTACGAAAGGACAAGAGAAATAAGGCCTACTTCACA-AAGCGCCTTCCCCCGTAAATGATATCATC
  5 AGENOME-ZPMRG-Z-V1.0   99.9%  99.7%       TCAAATTCCTCCCTGTACGAAAGGACAAGAGAAATAAGGCCTACTTCACA-AAGCGCCTTCCCCCGTAAATGATATCATC
  6 AGENOME-ZPMRG-T-V1.0   99.9%  99.8%       TCAAATTCCTCCCTGTACGAAAGGACAAGAGAAATAAGGCCTACTTCACA-AAGCGCCTTCCCCCGTAAATGATATCATC
  7 AGENOME-ZPMRG-HV-V1.0  99.9%  99.8%       TCAAATTCCTCCCTGTACGAAAGGACAAGAGAAATAAGGCCTACTTCACA-AAGCGCCTTCCCCCGTAAATGATATCATC
  8 AGENOME-ZPMRG-A-V1.0   99.9%  99.8%       TCAAATTCCTCCCTGTACGAAAGGACAAGAGAAATAAGGCCTACTTCACA-AAGCGCCTTCCCCCGTAAATGATATCATC
  9 NC_012920.1            99.9%  99.8%       TCAAATTCCTCCCTGTACGAAAGGACAAGAGAAATAAGGCCTACTTCACA-AAGCGCCTTCCCCCGTAAATGATATCATC
    consensus/100%                            TCAAATTCCTCCCTGTACGAAAGGACAAGAGAAATAAGGCCTACTTCACA.AAGCGCCTTCCCCCGTAAATGATATCATC
    consensus/90%                             TCAAATTCCTCCCTGTACGAAAGGACAAGAGAAATAAGGCCTACTTCACA.AAGCGCCTTCCCCCGTAAATGATATCATC
    consensus/80%                             TCAAATTCCTCCCTGTACGAAAGGACAAGAGAAATAAGGCCTACTTCACA.AAGCGCCTTCCCCCGTAAATGATATCATC
    consensus/70%                             TCAAATTCCTCCCTGTACGAAAGGACAAGAGAAATAAGGCCTACTTCACA.AAGCGCCTTCCCCCGTAAATGATATCATC

                              cov    pid  3201          .         .         .         .         :         .         .         . 3280
  1 AGENOME-ZPMRG-U-V1.0  100.0% 100.0%       TCAACTTAGTATTATACCCACACCCACCCAAGAACAGGGTTTGTTAAGATGGCAGAGCCCGGTAATCGCATAAAACTTAA
  2 AGENOME-ZPMCG-V1.0    100.0%  99.9%       TCAACTTAGTATTATACCCACACCCACCCAAGAACAGGGTTTGTTAAGATGGCAGAGCCCGGTAATCGCATAAAACTTAA
  3 AGENOME-ZPMRG-M-V1.0  100.0%  99.8%       TCAACTTAGTATTATACCCACACCCACCCAAGAACAGGGTTTGTTAAGATGGCAGAGCCCGGTAATCGCATAAAACTTAA
  4 AGENOME-ZPMRG-F-V1.0   99.9%  99.8%       TCAACTTAGTATTATACCCACACCCACCCAAGAACAGGGTTTGTTAAGATGGCAGAGCCCGGTAATCGCATAAAACTTAA
  5 AGENOME-ZPMRG-Z-V1.0   99.9%  99.7%       TCAACTTAGTATTATACCCACACCCACCCAAGAACAGGGTTTGTTAAGATGGCAGAGCCCGGTAATCGCATAAAACTTAA
  6 AGENOME-ZPMRG-T-V1.0   99.9%  99.8%       TCAACTTAGTATTATACCCACACCCACCCAAGAACAGGGTTTGTTAAGATGGCAGAGCCCGGTAATCGCATAAAACTTAA
  7 AGENOME-ZPMRG-HV-V1.0  99.9%  99.8%       TCAACTTAGTATTATACCCACACCCACCCAAGAACAGGGTTTGTTAAGATGGCAGAGCCCGGTAATCGCATAAAACTTAA
  8 AGENOME-ZPMRG-A-V1.0   99.9%  99.8%       TCAACTTAGTATTATACCCACACCCACCCAAGAACAGGGTTTGTTAAGATGGCAGAGCCCGGTAATCGCATAAAACTTAA
  9 NC_012920.1            99.9%  99.8%       TCAACTTAGTATTATACCCACACCCACCCAAGAACAGGGTTTGTTAAGATGGCAGAGCCCGGTAATCGCATAAAACTTAA
    consensus/100%                            TCAACTTAGTATTATACCCACACCCACCCAAGAACAGGGTTTGTTAAGATGGCAGAGCCCGGTAATCGCATAAAACTTAA
    consensus/90%                             TCAACTTAGTATTATACCCACACCCACCCAAGAACAGGGTTTGTTAAGATGGCAGAGCCCGGTAATCGCATAAAACTTAA
    consensus/80%                             TCAACTTAGTATTATACCCACACCCACCCAAGAACAGGGTTTGTTAAGATGGCAGAGCCCGGTAATCGCATAAAACTTAA
    consensus/70%                             TCAACTTAGTATTATACCCACACCCACCCAAGAACAGGGTTTGTTAAGATGGCAGAGCCCGGTAATCGCATAAAACTTAA

                              cov    pid  3281          .         3         .         .         .         .         :         . 3360
  1 AGENOME-ZPMRG-U-V1.0  100.0% 100.0%       AACTTTACAGTCAGAGGTTCAATTCCTCTTCTTAACAACATACCCATGGCCAACCTCCTACTCCTCATTGTACCCATTCT
  2 AGENOME-ZPMCG-V1.0    100.0%  99.9%       AACTTTACAGTCAGAGGTTCAATTCCTCTTCTTAACAACATACCCATGGCCAACCTCCTACTCCTCATTGTACCCATTCT
  3 AGENOME-ZPMRG-M-V1.0  100.0%  99.8%       AACTTTACAGTCAGAGGTTCAATTCCTCTTCTTAACAACATACCCATGGCCAACCTCCTACTCCTCATTGTACCCATTCT
  4 AGENOME-ZPMRG-F-V1.0   99.9%  99.8%       AACTTTACAGTCAGAGGTTCAATTCCTCTTCTTAACAACATACCCATGGCCAACCTCCTACTCCTCATTGTACCCATTCT
  5 AGENOME-ZPMRG-Z-V1.0   99.9%  99.7%       AACTTTACAGTCAGAGGTTCAATTCCTCTTCTTAACAACATACCCATGGCCAACCTCCTACTCCTCATTGTACCCATTCT
  6 AGENOME-ZPMRG-T-V1.0   99.9%  99.8%       AACTTTACAGTCAGAGGTTCAATTCCTCTTCTTAACAACATACCCATGGCCAACCTCCTACTCCTCATTGTACCCATTCT
  7 AGENOME-ZPMRG-HV-V1.0  99.9%  99.8%       AACTTTACAGTCAGAGGTTCAATTCCTCTTCTTAACAACATACCCATGGCCAACCTCCTACTCCTCATTGTACCCATTCT
  8 AGENOME-ZPMRG-A-V1.0   99.9%  99.8%       AACTTTACAGTCAGAGGTTCAATTCCTCTTCTTAACAACATACCCATGGCCAACCTCCTACTCCTCATTGTACCCATTCT
  9 NC_012920.1            99.9%  99.8%       AACTTTACAGTCAGAGGTTCAATTCCTCTTCTTAACAACATACCCATGGCCAACCTCCTACTCCTCATTGTACCCATTCT
    consensus/100%                            AACTTTACAGTCAGAGGTTCAATTCCTCTTCTTAACAACATACCCATGGCCAACCTCCTACTCCTCATTGTACCCATTCT
    consensus/90%                             AACTTTACAGTCAGAGGTTCAATTCCTCTTCTTAACAACATACCCATGGCCAACCTCCTACTCCTCATTGTACCCATTCT
    consensus/80%                             AACTTTACAGTCAGAGGTTCAATTCCTCTTCTTAACAACATACCCATGGCCAACCTCCTACTCCTCATTGTACCCATTCT
    consensus/70%                             AACTTTACAGTCAGAGGTTCAATTCCTCTTCTTAACAACATACCCATGGCCAACCTCCTACTCCTCATTGTACCCATTCT

                              cov    pid  3361          .         .         .         4         .         .         .         . 3440
```

```
  1 AGENOME-ZPMRG-U-V1.0  100.0% 100.0%       AATCGCAATGGCATTCCTAATGCTTACCGAACGAAAAATTCTAGGCTATATACAACTACGCAAAGGCCCCAACGTTGTAG
  2 AGENOME-ZPMCG-V1.0    100.0%  99.9%       AATCGCAATGGCATTCCTAATGCTTACCGAACGAAAAATTCTAGGCTATATACAACTACGCAAAGGCCCCAACGTTGTAG
  3 AGENOME-ZPMRG-M-V1.0  100.0%  99.8%       AATCGCAATGGCATTCCTAATGCTTACCGAACGAAAAATTCTAGGCTATATACAACTACGCAAAGGCCCCAACGTTGTAG
  4 AGENOME-ZPMRG-F-V1.0   99.9%  99.8%       AATCGCAATGGCATTCCTAATGCTTACCGAACGAAAAATTCTAGGCTATACACAACTACGCAAAGGCCCCAACGTTGTAG
  5 AGENOME-ZPMRG-Z-V1.0   99.9%  99.7%       AATCGCAATGGCATTCCTAATGCTTACCGAACGAAAAATTCTAGGCTATATACAACTACGCAAAGGCCCCAACGTTGTAG
  6 AGENOME-ZPMRG-T-V1.0   99.9%  99.8%       AATCGCAATGGCATTCCTAATGCTTACCGAACGAAAAATTCTAGGCTATATACAACTACGCAAAGGCCCCAACGTTGTAG
  7 AGENOME-ZPMRG-HV-V1.0  99.9%  99.8%       AATCGCAATGGCATTCCTAATGCTTACCGAACGAAAAATTCTAGGCTATATACAACTACGCAAAGGCCCCAACGTTGTAG
  8 AGENOME-ZPMRG-A-V1.0   99.9%  99.8%       AATCGCAATGGCATTCCTAATGCTTACCGAACGAAAAATTCTAGGCTATATACAACTACGCAAAGGCCCCAACGTTGTAG
  9 NC_012920.1            99.9%  99.8%       AATCGCAATGGCATTCCTAATGCTTACCGAACGAAAAATTCTAGGCTATATACAACTACGCAAAGGCCCCAACGTTGTAG
    consensus/100%                            AATCGCAATGGCATTCCTAATGCTTACCGAACGAAAAATTCTAGGCTATAsACAACTACGCAAAGGCCCCAACGTTGTAG
    consensus/90%                             AATCGCAATGGCATTCCTAATGCTTACCGAACGAAAAATTCTAGGCTATAsACAACTACGCAAAGGCCCCAACGTTGTAG
    consensus/80%                             AATCGCAATGGCATTCCTAATGCTTACCGAACGAAAAATTCTAGGCTATATACAACTACGCAAAGGCCCCAACGTTGTAG
    consensus/70%                             AATCGCAATGGCATTCCTAATGCTTACCGAACGAAAAATTCTAGGCTATATACAACTACGCAAAGGCCCCAACGTTGTAG

                             cov    pid  3441          :         .         .         .         .         5         .         . 3520
  1 AGENOME-ZPMRG-U-V1.0  100.0% 100.0%       GCCCCTACGGGCTACTACAACCCTTCGCTGACGCCATAAAACTCTTCACCAAAGAGCCCCTAAAACCCGCCACATCTACC
  2 AGENOME-ZPMCG-V1.0    100.0%  99.9%       GCCCCTACGGGCTACTACAACCCTTCGCTGACGCCATAAAACTCTTCACCAAAGAGCCCCTAAAACCCGCCACATCTACC
  3 AGENOME-ZPMRG-M-V1.0  100.0%  99.8%       GCCCCTACGGGCTACTACAACCCTTCGCTGACGCCATAAAACTCTTCACCAAAGAGCCCCTAAAACCCGCCACATCTACC
  4 AGENOME-ZPMRG-F-V1.0   99.9%  99.8%       GCCCCTACGGGCTACTACAACCCTTCGCTGACGCCATAAAACTCTTCACCAAAGAGCCCCTAAAACCCGCCACATCTACC
  5 AGENOME-ZPMRG-Z-V1.0   99.9%  99.7%       GCCCCTACGGGCTACTACAACCCTTCGCTGACGCCATAAAACTCTTCACCAAAGAGCCCCTAAAACCCGCCACATCTACC
  6 AGENOME-ZPMRG-T-V1.0   99.9%  99.8%       GCCCCTACGGGCTACTACAACCCTTCGCTGACGCCATAAAACTCTTCACCAAAGAGCCCCTAAAACCCGCCACATCTACC
  7 AGENOME-ZPMRG-HV-V1.0  99.9%  99.8%       GCCCCTACGGGCTACTACAACCCTTCGCTGACGCCATAAAACTCTTCACCAAAGAGCCCCTAAAACCCGCCACATCTACC
  8 AGENOME-ZPMRG-A-V1.0   99.9%  99.8%       GCCCCTACGGGCTACTACAACCCTTCGCTGACGCCATAAAACTCTTCACCAAAGAGCCCCTAAAACCCGCCACATCTACC
  9 NC_012920.1            99.9%  99.8%       GCCCCTACGGGCTACTACAACCCTTCGCTGACGCCATAAAACTCTTCACCAAAGAGCCCCTAAAACCCGCCACATCTACC
    consensus/100%                            GCCCCTACGGGCTACTACAACCCTTCGCTGACGCCATAAAACTCTTCACCAAAGAGCCCCTAAAACCCGCCACATCTACC
    consensus/90%                             GCCCCTACGGGCTACTACAACCCTTCGCTGACGCCATAAAACTCTTCACCAAAGAGCCCCTAAAACCCGCCACATCTACC
    consensus/80%                             GCCCCTACGGGCTACTACAACCCTTCGCTGACGCCATAAAACTCTTCACCAAAGAGCCCCTAAAACCCGCCACATCTACC
    consensus/70%                             GCCCCTACGGGCTACTACAACCCTTCGCTGACGCCATAAAACTCTTCACCAAAGAGCCCCTAAAACCCGCCACATCTACC

                             cov    pid  3521          .         .         :         .         .         .         .         6 3600
  1 AGENOME-ZPMRG-U-V1.0  100.0% 100.0%       ATCACCCTCTACATCACCGCCCCGACCTTAGCTCTCACCATCGCTCTTCTACTATGAACCCCCCTCCCCATACCCAACCC
  2 AGENOME-ZPMCG-V1.0    100.0%  99.9%       ATCACCCTCTACATCACCGCCCCGACCTTAGCTCTCACCATCGCTCTTCTACTATGAACCCCCCTCCCCATACCCAACCC
  3 AGENOME-ZPMRG-M-V1.0  100.0%  99.8%       ATCACCCTCTACATCACCGCCCCGACCTTAGCTCTCACCATCGCTCTTCTACTATGAACCCCCCTCCCCATACCCAACCC
  4 AGENOME-ZPMRG-F-V1.0   99.9%  99.8%       ATCACCCTCTACATCACCGCCCCGACCTTAGCTCTCACCATCGCTCTTCTACTATGAACCCCCCTCCCCATACCCAACCC
  5 AGENOME-ZPMRG-Z-V1.0   99.9%  99.7%       ATCACCCTCTACATCACCGCCCCGACCTTAGCTCTCACCATCGCTCTTCTACTATGAACCCCCCTCCCCATACCCAACCC
  6 AGENOME-ZPMRG-T-V1.0   99.9%  99.8%       ATCACCCTCTACATCACCGCCCCGACCTTAGCTCTCACCATCGCTCTTCTACTATGAACCCCCCTCCCCATACCCAACCC
  7 AGENOME-ZPMRG-HV-V1.0  99.9%  99.8%       ATCACCCTCTACATCACCGCCCCGACCTTAGCTCTCACCATCGCTCTTCTACTATGAACCCCCCTCCCCATACCCAACCC
  8 AGENOME-ZPMRG-A-V1.0   99.9%  99.8%       ATCACCCTCTACATCACCGCCCCGACCTTAGCTCTCACCATCGCTCTTCTACTATGAACCCCCCTCCCCATACCCAACCC
  9 NC_012920.1            99.9%  99.8%       ATCACCCTCTACATCACCGCCCCGACCTTAGCTCTCACCATCGCTCTTCTACTATGAACCCCCCTCCCCATACCCAACCC
    consensus/100%                            ATCACCCTCTACATCACCGCCCCGACCTTAGCTCTCACCATCGCTCTTCTACTATGAACCCCCCTCCCCATACCCAACCC
    consensus/90%                             ATCACCCTCTACATCACCGCCCCGACCTTAGCTCTCACCATCGCTCTTCTACTATGAACCCCCCTCCCCATACCCAACCC
    consensus/80%                             ATCACCCTCTACATCACCGCCCCGACCTTAGCTCTCACCATCGCTCTTCTACTATGAACCCCCCTCCCCATACCCAACCC
    consensus/70%                             ATCACCCTCTACATCACCGCCCCGACCTTAGCTCTCACCATCGCTCTTCTACTATGAACCCCCCTCCCCATACCCAACCC

                             cov    pid  3601          .         .         .         .         :         .         .         . 3680
  1 AGENOME-ZPMRG-U-V1.0  100.0% 100.0%       CCTGGTCAACCTCAACCTAGGCCTCCTATTTATTCTAGCCACCTCTAGCCTAGCCGTTTACTCAATCCTCTGATCAGGGT
  2 AGENOME-ZPMCG-V1.0    100.0%  99.9%       CCTGGTCAACCTCAACCTAGGCCTCCTATTTATTCTAGCCACCTCTAGCCTAGCCGTTTACTCAATCCTCTGATCAGGGT
  3 AGENOME-ZPMRG-M-V1.0  100.0%  99.8%       CCTGGTCAACCTCAACCTAGGCCTCCTATTTATTCTAGCCACCTCTAGCCTAGCCGTTTACTCAATCCTCTGATCAGGGT
  4 AGENOME-ZPMRG-F-V1.0   99.9%  99.8%       CCTGGTCAACCTCAACCTAGGCCTCCTATTTATTCTAGCCACCTCTAGCCTAGCCGTTTACTCAATCCTCTGATCAGGGT
  5 AGENOME-ZPMRG-Z-V1.0   99.9%  99.7%       CCTGGTCAACCTCAACCTAGGCCTCCTATTTATTCTAGCCACCTCTAGCCTAGCCGTTTACTCAATCCTCTGATCAGGGT
  6 AGENOME-ZPMRG-T-V1.0   99.9%  99.8%       CCTGGTCAACCTCAACCTAGGCCTCCTATTTATTCTAGCCACCTCTAGCCTAGCCGTTTACTCAATCCTCTGATCAGGGT
  7 AGENOME-ZPMRG-HV-V1.0  99.9%  99.8%       CCTGGTCAACCTCAACCTAGGCCTCCTATTTATTCTAGCCACCTCTAGCCTAGCCGTTTACTCAATCCTCTGATCAGGGT
  8 AGENOME-ZPMRG-A-V1.0   99.9%  99.8%       CCTGGTCAACCTCAACCTAGGCCTCCTATTTATTCTAGCCACCTCTAGCCTAGCCGTTTACTCAATCCTCTGATCAGGGT
  9 NC_012920.1            99.9%  99.8%       CCTGGTCAACCTCAACCTAGGCCTCCTATTTATTCTAGCCACCTCTAGCCTAGCCGTTTACTCAATCCTCTGATCAGGGT
    consensus/100%                            CCTGGTCAACCTCAACCTAGGCCTCCTATTTATTCTAGCCACCTCTAGCCTAGCCGTTTACTCAATCCTCTGATCAGGGT
    consensus/90%                             CCTGGTCAACCTCAACCTAGGCCTCCTATTTATTCTAGCCACCTCTAGCCTAGCCGTTTACTCAATCCTCTGATCAGGGT
    consensus/80%                             CCTGGTCAACCTCAACCTAGGCCTCCTATTTATTCTAGCCACCTCTAGCCTAGCCGTTTACTCAATCCTCTGATCAGGGT
    consensus/70%                             CCTGGTCAACCTCAACCTAGGCCTCCTATTTATTCTAGCCACCTCTAGCCTAGCCGTTTACTCAATCCTCTGATCAGGGT

                             cov    pid  3681          .         7         .         .         .         .         :         . 3760
  1 AGENOME-ZPMRG-U-V1.0  100.0% 100.0%       GAGCATCAAACTCAAACTACGCCCTGATCGGCGCACTGCGAGCAGTAGCCCAAACAATCTCATATGAAGTCACCCTAGCC
  2 AGENOME-ZPMCG-V1.0    100.0%  99.9%       GAGCATCAAACTCAAACTACGCCCTGATCGGCGCACTGCGAGCAGTAGCCCAAACAATCTCATATGAAGTCACCCTAGCC
  3 AGENOME-ZPMRG-M-V1.0  100.0%  99.8%       GAGCATCAAACTCAAACTACGCCCTGATCGGCGCACTGCGAGCAGTAGCCCAAACAATCTCATATGAAGTCACCCTAGCC
  4 AGENOME-ZPMRG-F-V1.0   99.9%  99.8%       GAGCATCAAACTCAAACTACGCCCTGATCGGCGCACTGCGAGCAGTAGCCCAAACAATCTCATATGAAGTCACCCTAGCC
  5 AGENOME-ZPMRG-Z-V1.0   99.9%  99.7%       GAGCATCAAACTCAAACTACGCCCTGATCGGCGCACTGCGAGCAGTAGCCCAAACAATCTCATATGAAGTCACCCTAGCC
  6 AGENOME-ZPMRG-T-V1.0   99.9%  99.8%       GAGCATCAAACTCAAACTACGCCCTGATCGGCGCACTGCGAGCAGTAGCCCAAACAATCTCATATGAAGTCACCCTAGCC
  7 AGENOME-ZPMRG-HV-V1.0  99.9%  99.8%       GAGCATCAAACTCAAACTACGCCCTGATCGGCGCACTGCGAGCAGTAGCCCAAACAATCTCATATGAAGTCACCCTAGCC
  8 AGENOME-ZPMRG-A-V1.0   99.9%  99.8%       GAGCATCAAACTCAAACTACGCCCTGATCGGCGCACTGCGAGCAGTAGCCCAAACAATCTCATATGAAGTCACCCTAGCC
  9 NC_012920.1            99.9%  99.8%       GAGCATCAAACTCAAACTACGCCCTGATCGGCGCACTGCGAGCAGTAGCCCAAACAATCTCATATGAAGTCACCCTAGCC
    consensus/100%                            GAGCATCAAACTCAAACTACGCCCTGATCGGCGCACTGCGAGCAGTAGCCCAAACAATCTCATATGAAGTCACCCTAGCC
    consensus/90%                             GAGCATCAAACTCAAACTACGCCCTGATCGGCGCACTGCGAGCAGTAGCCCAAACAATCTCATATGAAGTCACCCTAGCC
    consensus/80%                             GAGCATCAAACTCAAACTACGCCCTGATCGGCGCACTGCGAGCAGTAGCCCAAACAATCTCATATGAAGTCACCCTAGCC
    consensus/70%                             GAGCATCAAACTCAAACTACGCCCTGATCGGCGCACTGCGAGCAGTAGCCCAAACAATCTCATATGAAGTCACCCTAGCC

                             cov    pid  3761          .         .         .         8         .         .         .         . 3840
  1 AGENOME-ZPMRG-U-V1.0  100.0% 100.0%       ATCATTCTACTATCAACATTACTAATAAGTGGCTCCTTTAACCTCTCCACCCTTATCACAACACAAGAACACCTCTGATT
  2 AGENOME-ZPMCG-V1.0    100.0%  99.9%       ATCATTCTACTATCAACATTACTAATAAGTGGCTCCTTTAACCTCTCCACCCTTATCACAACACAAGAACACCTCTGATT
  3 AGENOME-ZPMRG-M-V1.0  100.0%  99.8%       ATCATTCTACTATCAACATTACTAATAAGTGGCTCCTTTAACCTCTCCACCCTTATCACAACACAAGAACACCTCTGATT
  4 AGENOME-ZPMRG-F-V1.0   99.9%  99.8%       ATCATTCTACTATCAACATTACTAATAAGTGGCTCCTTTAACCTCTCCACCCTTATCACAACACAAGAACACCTCTGATT
  5 AGENOME-ZPMRG-Z-V1.0   99.9%  99.7%       ATCATTCTACTATCAACATTACTAATAAGTGGCTCCTTTAACCTCTCCACCCTTATCACAACACAAGAACACCTCTGATT
  6 AGENOME-ZPMRG-T-V1.0   99.9%  99.8%       ATCATTCTACTATCAACATTACTAATAAGTGGCTCCTTTAACCTCTCCACCCTTATCACAACACAAGAACACCTCTGATT
  7 AGENOME-ZPMRG-HV-V1.0  99.9%  99.8%       ATCATTCTACTATCAACATTACTAATAAGTGGCTCCTTTAACCTCTCCACCCTTATCACAACACAAGAACACCTCTGATT
  8 AGENOME-ZPMRG-A-V1.0   99.9%  99.8%       ATCATTCTACTATCAACATTACTAATAAGTGGCTCCTTTAACCTCTCCACCCTTATCACAACACAAGAACACCTCTGATT
  9 NC_012920.1            99.9%  99.8%       ATCATTCTACTATCAACATTACTAATAAGTGGCTCCTTTAACCTCTCCACCCTTATCACAACACAAGAACACCTCTGATT
    consensus/100%                            ATCATTCTACTATCAACATTACTAATAAGTGGCTCCTTTAACCTCTCCACCCTTATCACAACACAAGAACACCTCTGATT
    consensus/90%                             ATCATTCTACTATCAACATTACTAATAAGTGGCTCCTTTAACCTCTCCACCCTTATCACAACACAAGAACACCTCTGATT
    consensus/80%                             ATCATTCTACTATCAACATTACTAATAAGTGGCTCCTTTAACCTCTCCACCCTTATCACAACACAAGAACACCTCTGATT
    consensus/70%                             ATCATTCTACTATCAACATTACTAATAAGTGGCTCCTTTAACCTCTCCACCCTTATCACAACACAAGAACACCTCTGATT

                             cov    pid  3841          :         .         .         .         .         9         .         . 3920
  1 AGENOME-ZPMRG-U-V1.0  100.0% 100.0%       ACTCCTGCCATCATGACCCTTGGCCATAATATGATTTATCTCCACACTAGCAGAGACCAACCGAACCCCCTTCGACCTTG
```

```
 2 AGENOME-ZPMCG-V1.0     100.0%  99.9%       ACTCCTGCCATCATGACCCTTGGCCATAATATGATTTATCTCCACACTAGCAGAGACCAACCGAACCCCCTTCGACCTTG
 3 AGENOME-ZPMRG-M-V1.0  100.0%  99.8%       ACTCCTGCCATCATGACCCTTGGCCATAATATGATTTATCTCCACACTAGCAGAGACCAACCGAACCCCCTTCGACCTTG
 4 AGENOME-ZPMRG-F-V1.0   99.9%  99.8%       ACTCCTGCCATCATGACCCTTGGCCATAATATGATTTATCTCCACACTAGCAGAGACCAACCGAACCCCCTTCGACCTTG
 5 AGENOME-ZPMRG-Z-V1.0   99.9%  99.7%       ACTCCTGCCATCATGACCCTTGGCCATAATATGATTTATCTCCACACTAGCAGAGACCAACCGAACCCCCTTCGACCTTG
 6 AGENOME-ZPMRG-T-V1.0   99.9%  99.8%       ACTCCTGCCATCATGACCCTTGGCCATAATATGATTTATCTCCACACTAGCAGAGACCAACCGAACCCCCTTCGACCTTG
 7 AGENOME-ZPMRG-HV-V1.0  99.9%  99.8%       ACTCCTGCCATCATGACCCTTGGCCATAATATGATTTATCTCCACACTAGCAGAGACCAACCGAACCCCCTTCGACCTTG
 8 AGENOME-ZPMRG-A-V1.0   99.9%  99.8%       ACTCCTGCCATCATGACCCTTGGCCATAATATGATTTATCTCCACACTAGCAGAGACCAACCGAACCCCCTTCGACCTTG
 9 NC_012920.1            99.9%  99.8%       ACTCCTGCCATCATGACCCTTGGCCATAATATGATTTATCTCCACACTAGCAGAGACCAACCGAACCCCCTTCGACCTTG
   consensus/100%                            ACTCCTGCCATCATGACCCTTGGCCATAATATGATTTATCTCCACACTAGCAGAGACCAACCGAACCCCCTTCGACCTTG
   consensus/90%                             ACTCCTGCCATCATGACCCTTGGCCATAATATGATTTATCTCCACACTAGCAGAGACCAACCGAACCCCCTTCGACCTTG
   consensus/80%                             ACTCCTGCCATCATGACCCTTGGCCATAATATGATTTATCTCCACACTAGCAGAGACCAACCGAACCCCCTTCGACCTTG
   consensus/70%                             ACTCCTGCCATCATGACCCTTGGCCATAATATGATTTATCTCCACACTAGCAGAGACCAACCGAACCCCCTTCGACCTTG

                            cov    pid  3921          .         .         :         .         .         .         .         0 4000
 1 AGENOME-ZPMRG-U-V1.0  100.0% 100.0%       CCGAAGGGGAGTCCGAACTAGTCTCAGGCTTCAACATCGAATACGCCGCAGGCCCCTTCGCCCTATTCTTCATAGCCGAA
 2 AGENOME-ZPMCG-V1.0     100.0%  99.9%       CCGAAGGGGAGTCCGAACTAGTCTCAGGCTTCAACATCGAATACGCCGCAGGCCCCTTCGCCCTATTCTTCATAGCCGAA
 3 AGENOME-ZPMRG-M-V1.0  100.0%  99.8%       CCGAAGGGGAGTCCGAACTAGTCTCAGGCTTCAACATCGAATACGCCGCAGGCCCCTTCGCCCTATTCTTCATAGCCGAA
 4 AGENOME-ZPMRG-F-V1.0   99.9%  99.8%       CCGAAGGGGAGTCCGAACTAGTCTCAGGCTTCAACATCGAATACGCCGCAGGCCCCTTCGCCTTATTCTTCATAGCCGAA
 5 AGENOME-ZPMRG-Z-V1.0   99.9%  99.7%       CCGAAGGGGAGTCCGAACTAGTCTCAGGCTTCAACATCGAATACGCCGCAGGCCCCTTCGCCCTATTCTTCATAGCCGAA
 6 AGENOME-ZPMRG-T-V1.0   99.9%  99.8%       CCGAAGGGGAGTCCGAACTAGTCTCAGGCTTCAACATCGAATACGCCGCAGGCCCCTTCGCCCTATTCTTCATAGCCGAA
 7 AGENOME-ZPMRG-HV-V1.0  99.9%  99.8%       CCGAAGGGGAGTCCGAACTAGTCTCAGGCTTCAACATCGAATACGCCGCAGGCCCCTTCGCCCTATTCTTCATAGCCGAA
 8 AGENOME-ZPMRG-A-V1.0   99.9%  99.8%       CCGAAGGGGAGTCCGAACTAGTCTCAGGCTTCAACATCGAATACGCCGCAGGCCCCTTCGCCCTATTCTTCATAGCCGAA
 9 NC_012920.1            99.9%  99.8%       CCGAAGGGGAGTCCGAACTAGTCTCAGGCTTCAACATCGAATACGCCGCAGGCCCCTTCGCCCTATTCTTCATAGCCGAA
   consensus/100%                            CCGAAGGGGAGTCCGAACTAGTCTCAGGCTTCAACATCGAATACGCCGCAGGCCCCTTCGCCsTATTCTTCATAGCCGAA
   consensus/90%                             CCGAAGGGGAGTCCGAACTAGTCTCAGGCTTCAACATCGAATACGCCGCAGGCCCCTTCGCCsTATTCTTCATAGCCGAA
   consensus/80%                             CCGAAGGGGAGTCCGAACTAGTCTCAGGCTTCAACATCGAATACGCCGCAGGCCCCTTCGCCCTATTCTTCATAGCCGAA
   consensus/70%                             CCGAAGGGGAGTCCGAACTAGTCTCAGGCTTCAACATCGAATACGCCGCAGGCCCCTTCGCCCTATTCTTCATAGCCGAA

                            cov    pid  4001          .         .         .         .         :         .         .         . 4080
 1 AGENOME-ZPMRG-U-V1.0  100.0% 100.0%       TACACAAACATTATTATAATAAACACCCTCACCACTACAATCTTCCTAGGAACAACATATGACGCACTCTCCCCTGAACT
 2 AGENOME-ZPMCG-V1.0     100.0%  99.9%       TACACAAACATTATTATAATAAACACCCTCACCACTACAATCTTCCTAGGAACAACATATGACGCACTCTCCCCTGAACT
 3 AGENOME-ZPMRG-M-V1.0  100.0%  99.8%       TACACAAACATTATTATAATAAACACCCTCACCACTACAATCTTCCTAGGAACAACATATGACGCACTCTCCCCTGAACT
 4 AGENOME-ZPMRG-F-V1.0   99.9%  99.8%       TACACAAACATCATTATAATAAACACCCTCACCACTACAATCTTCCTAGGAACAACATATGACGCACTCTCCCCTGAACT
 5 AGENOME-ZPMRG-Z-V1.0   99.9%  99.7%       TACACAAACATTATTATAATAAACACCCTCACCACTACAATCTTCCTAGGAACAACATATGACGCACTCTCCCCTGAACT
 6 AGENOME-ZPMRG-T-V1.0   99.9%  99.8%       TACACAAACATTATTATAATAAACACCCTCACCACTACAATCTTCCTAGGAACAACATATGACGCACTCTCCCCTGAACT
 7 AGENOME-ZPMRG-HV-V1.0  99.9%  99.8%       TACACAAACATTATTATAATAAACACCCTCACCACTACAATCTTCCTAGGAACAACATATGACGCACTCTCCCCTGAACT
 8 AGENOME-ZPMRG-A-V1.0   99.9%  99.8%       TACACAAACATTATTATAATAAACACCCTCACCACTACAATCTTCCTAGGAACAACATATGACGCACTCTCCCCTGAACT
 9 NC_012920.1            99.9%  99.8%       TACACAAACATTATTATAATAAACACCCTCACCACTACAATCTTCCTAGGAACAACATATGACGCACTCTCCCCTGAACT
   consensus/100%                            TACACAAACATsATTATAATAAACACCCTCACCACTACAATCTTCCTAGGAACAACATATGACGCACTCTCCCCTGAACT
   consensus/90%                             TACACAAACATsATTATAATAAACACCCTCACCACTACAATCTTCCTAGGAACAACATATGACGCACTCTCCCCTGAACT
   consensus/80%                             TACACAAACATTATTATAATAAACACCCTCACCACTACAATCTTCCTAGGAACAACATATGACGCACTCTCCCCTGAACT
   consensus/70%                             TACACAAACATTATTATAATAAACACCCTCACCACTACAATCTTCCTAGGAACAACATATGACGCACTCTCCCCTGAACT

                            cov    pid  4081          .         1         .         .         .         .         :         . 4160
 1 AGENOME-ZPMRG-U-V1.0  100.0% 100.0%       CTACACAACATATTTTGTCACCAAGACCCTACTTCTAACCTCCCTGTTCTTATGAATTCGAACAGCATACCCCCGATTCC
 2 AGENOME-ZPMCG-V1.0     100.0%  99.9%       CTACACAACATATTTTGTCACCAAGACCCTACTTCTAACCTCCCTGTTCTTATGAATTCGAACAGCATACCCCCGATTCC
 3 AGENOME-ZPMRG-M-V1.0  100.0%  99.8%       CTACACAACATATTTTGTCACCAAGACCCTACTTCTAACCTCCCTGTTCTTATGAATTCGAACAGCATACCCCCGATTCC
 4 AGENOME-ZPMRG-F-V1.0   99.9%  99.8%       CTACACAACATATTTTGTCACCAAGACCCTACTTCTAACCTCCCTGTTCTTATGAATTCGAACAGCATACCCCCGATTCC
 5 AGENOME-ZPMRG-Z-V1.0   99.9%  99.7%       CTACACAACATATTTTGTCACCAAGACCCTACTTCTAACCTCCCTGTTCTTATGAATTCGAACAGCATACCCCCGATTCC
 6 AGENOME-ZPMRG-T-V1.0   99.9%  99.8%       CTACACAACATATTTTGTCACCAAGACCCTACTTCTAACCTCCCTGTTCTTATGAATTCGAACAGCATACCCCCGATTCC
 7 AGENOME-ZPMRG-HV-V1.0  99.9%  99.8%       CTACACAACATATTTTGTCACCAAGACCCTACTTCTAACCTCCCTGTTCTTATGAATTCGAACAGCATACCCCCGATTCC
 8 AGENOME-ZPMRG-A-V1.0   99.9%  99.8%       CTACACAACATATTTTGTCACCAAGACCCTACTTCTAACCTCCCTGTTCTTATGAATTCGAACAGCATACCCCCGATTCC
 9 NC_012920.1            99.9%  99.8%       CTACACAACATATTTTGTCACCAAGACCCTACTTCTAACCTCCCTGTTCTTATGAATTCGAACAGCATACCCCCGATTCC
   consensus/100%                            CTACACAACATATTTTGTCACCAAGACCCTACTTCTAACCTCCCTGTTCTTATGAATTCGAACAGCATACCCCCGATTCC
   consensus/90%                             CTACACAACATATTTTGTCACCAAGACCCTACTTCTAACCTCCCTGTTCTTATGAATTCGAACAGCATACCCCCGATTCC
   consensus/80%                             CTACACAACATATTTTGTCACCAAGACCCTACTTCTAACCTCCCTGTTCTTATGAATTCGAACAGCATACCCCCGATTCC
   consensus/70%                             CTACACAACATATTTTGTCACCAAGACCCTACTTCTAACCTCCCTGTTCTTATGAATTCGAACAGCATACCCCCGATTCC

                            cov    pid  4161          .         .         .         2         .         .         .         . 4240
 1 AGENOME-ZPMRG-U-V1.0  100.0% 100.0%       GCTACGACCAACTCATACACCTCCTATGAAAAACTTCCTACCACTCACCCTAGCATTACTTATATGATATGTCTCCATA
 2 AGENOME-ZPMCG-V1.0     100.0%  99.9%       GCTACGACCAACTCATACACCTCCTATGAAAAACTTCCTACCACTCACCCTAGCATTACTTATATGATATGTCTCCATA
 3 AGENOME-ZPMRG-M-V1.0  100.0%  99.8%       GCTACGACCAACTCATACACCTCCTATGAAAAACTTCCTACCACTCACCCTAGCATTACTTATATGATATGTCTCCATA
 4 AGENOME-ZPMRG-F-V1.0   99.9%  99.8%       GCTACGACCAACTCATACACCTCCTATGAAAAACTTCCTACCACTCACCCTAGCATTACTTATATGATATGTCTCCATA
 5 AGENOME-ZPMRG-Z-V1.0   99.9%  99.7%       GCTACGACCAACTCATACACCTCCTATGAAAAACTTCCTACCACTCACCCTAGCATTACTTATATGATATGTCTCCATA
 6 AGENOME-ZPMRG-T-V1.0   99.9%  99.8%       GCTACGACCAACTCATACACCTCCTATGAAAAACTTCCTACCACTCACCCTAGCATTACTTATATGACATGTCTCCATA
 7 AGENOME-ZPMRG-HV-V1.0  99.9%  99.8%       GCTACGACCAACTCATACACCTCCTATGAAAAACTTCCTACCACTCACCCTAGCATTACTTATATGATATGTCTCCATA
 8 AGENOME-ZPMRG-A-V1.0   99.9%  99.8%       GCTACGACCAACTCATACACCTCCTATGAAAAACTTCCTACCACTCACCCTAGCATTACTTATATGATATGTCTCCATA
 9 NC_012920.1            99.9%  99.8%       GCTACGACCAACTCATACACCTCCTATGAAAAACTTCCTACCACTCACCCTAGCATTACTTATATGATATGTCTCCATA
   consensus/100%                            GCTACGACCAACTCATACACCTCCTATGAAAAACTTCCTACCACTCACCCTAGCATTACTTATATGAsATGTCTCCATA
   consensus/90%                             GCTACGACCAACTCATACACCTCCTATGAAAAACTTCCTACCACTCACCCTAGCATTACTTATATGAsATGTCTCCATA
   consensus/80%                             GCTACGACCAACTCATACACCTCCTATGAAAAACTTCCTACCACTCACCCTAGCATTACTTATATGATATGTCTCCATA
   consensus/70%                             GCTACGACCAACTCATACACCTCCTATGAAAAACTTCCTACCACTCACCCTAGCATTACTTATATGATATGTCTCCATA

                            cov    pid  4241          :         .         .         .         .         3         .         . 4320
 1 AGENOME-ZPMRG-U-V1.0  100.0% 100.0%       CCCATTACAATCTCCAGCATTCCCCCTCAAACCTAAGAAATATGTCTGATAAAAGAGTTACTTTGATAGAGTAAATAATA
 2 AGENOME-ZPMCG-V1.0     100.0%  99.9%       CCCATTACAATCTCCAGCATTCCCCCTCAAACCTAAGAAATATGTCTGATAAAAGAGTTACTTTGATAGAGTAAATAATA
 3 AGENOME-ZPMRG-M-V1.0  100.0%  99.8%       CCCATTACAATCTCCAGCATTCCCCCTCAAACCTAAGAAATATGTCTGATAAAAGAGTTACTTTGATAGAGTAAATAATA
 4 AGENOME-ZPMRG-F-V1.0   99.9%  99.8%       CCCATTACAATCTCCAGCATTCCCCCTCAAACCTAAGAAATATGTCTGATAAAAGAGTTACTTTGATAGAGTAAATAATA
 5 AGENOME-ZPMRG-Z-V1.0   99.9%  99.7%       CCCATTACAATCTCCAGCATTCCCCCTCAAACCTAAGAAATATGTCTGATAAAAGAGTTACTTTGATAGAGTAAATAATA
 6 AGENOME-ZPMRG-T-V1.0   99.9%  99.8%       CCCATTACAATCTCCAGCATTCCCCCTCAAACCTAAGAAATATGTCTGATAAAAGAGTTACTTTGATAGAGTAAATAATA
 7 AGENOME-ZPMRG-HV-V1.0  99.9%  99.8%       CCCATTACAATCTCCAGCATTCCCCCTCAAACCTAAGAAATATGTCTGATAAAAGAGTTACTTTGATAGAGTAAATAATA
 8 AGENOME-ZPMRG-A-V1.0   99.9%  99.8%       CCCATTACAATCTCCAGCATTCCCCCTCAAACCTAAGAAATATGTCTGATAAAAGAGTTACTTTGATAGAGTAAATAATA
 9 NC_012920.1            99.9%  99.8%       CCCATTACAATCTCCAGCATTCCCCCTCAAACCTAAGAAATATGTCTGATAAAAGAGTTACTTTGATAGAGTAAATAATA
   consensus/100%                            CCCATTACAATCTCCAGCATTCCCCCTCAAACCTAAGAAATATGTCTGATAAAAGAGTTACTTTGATAGAGTAAATAATA
   consensus/90%                             CCCATTACAATCTCCAGCATTCCCCCTCAAACCTAAGAAATATGTCTGATAAAAGAGTTACTTTGATAGAGTAAATAATA
   consensus/80%                             CCCATTACAATCTCCAGCATTCCCCCTCAAACCTAAGAAATATGTCTGATAAAAGAGTTACTTTGATAGAGTAAATAATA
   consensus/70%                             CCCATTACAATCTCCAGCATTCCCCCTCAAACCTAAGAAATATGTCTGATAAAAGAGTTACTTTGATAGAGTAAATAATA

                            cov    pid  4321          .         .         :         .         .         .         .         4 4400
 1 AGENOME-ZPMRG-U-V1.0  100.0% 100.0%       GGAGCTTAAACCCCCTTATTTCTAGGACTATGAGAATCGAACCCATCCCTGAGAATCCAAAATTCTCCGTGCCACCTATC
 2 AGENOME-ZPMCG-V1.0     100.0%  99.9%       GGAGCTTAAACCCCCTTATTTCTAGGACTATGAGAATCGAACCCATCCCTGAGAATCCAAAATTCTCCGTGCCACCTATC
```

```
 3 AGENOME-ZPMRG-M-V1.0  100.0%  99.8%       GGAGCTTAAACCCCCTTATTTCTAGGACTATGAGAATCGAACCCATCCCTGAGAATCCAAAATTCTCCGTGCCACCTATC
 4 AGENOME-ZPMRG-F-V1.0   99.9%  99.8%       GGAGCTTAAACCCCCTTATTTCTAGGACTATGAGAATCGAACCCATCCCTGAGAATCCAAAATTCTCCGTGCCACCTATC
 5 AGENOME-ZPMRG-Z-V1.0   99.9%  99.7%       GGAGCTTAAACCCCCTTATTTCTAGGACTATGAGAATCGAACCCATCCCTGAGAATCCAAAATTCTCCGTGCCACCTATC
 6 AGENOME-ZPMRG-T-V1.0   99.9%  99.8%       GGAGCTTAAACCCCCTTATTTCTAGGACTATGAGAATCGAACCCATCCCTGAGAATCCAAAATTCTCCGTGCCACCTATC
 7 AGENOME-ZPMRG-HV-V1.0  99.9%  99.8%       GGAGCTTAAACCCCCTTATTTCTAGGACTATGAGAATCGAACCCATCCCTGAGAATCCAAAATTCTCCGTGCCACCTATC
 8 AGENOME-ZPMRG-A-V1.0   99.9%  99.8%       GGAGCTTAAACCCCCTTATTTCTAGGACTATGAGAATCGAACCCATCCCTGAGAATCCAAAATTCTCCGTGCCACCTATC
 9 NC_012920.1            99.9%  99.8%       GGAGCTTAAACCCCCTTATTTCTAGGACTATGAGAATCGAACCCATCCCTGAGAATCCAAAATTCTCCGTGCCACCTATC
   consensus/100%                            GGAGCTTAAACCCCCTTATTTCTAGGACTATGAGAATCGAACCCATCCCTGAGAATCCAAAATTCTCCGTGCCACCTATC
   consensus/90%                             GGAGCTTAAACCCCCTTATTTCTAGGACTATGAGAATCGAACCCATCCCTGAGAATCCAAAATTCTCCGTGCCACCTATC
   consensus/80%                             GGAGCTTAAACCCCCTTATTTCTAGGACTATGAGAATCGAACCCATCCCTGAGAATCCAAAATTCTCCGTGCCACCTATC
   consensus/70%                             GGAGCTTAAACCCCCTTATTTCTAGGACTATGAGAATCGAACCCATCCCTGAGAATCCAAAATTCTCCGTGCCACCTATC

                            cov    pid  4401          .         .         .         .         :         .         .         . 4480
 1 AGENOME-ZPMRG-U-V1.0  100.0% 100.0%       ACACCCCATCCTAAAGTAAGGTCAGCTAAATAAGCTATCGGGCCCATACCCCGAAAATGTTGGTTATACCCTTCCCGTAC
 2 AGENOME-ZPMCG-V1.0    100.0%  99.9%       ACACCCCATCCTAAAGTAAGGTCAGCTAAATAAGCTATCGGGCCCATACCCCGAAAATGTTGGTTATACCCTTCCCGTAC
 3 AGENOME-ZPMRG-M-V1.0  100.0%  99.8%       ACACCCCATCCTAAAGTAAGGTCAGCTAAATAAGCTATCGGGCCCATACCCCGAAAATGTTGGTTATACCCTTCCCGTAC
 4 AGENOME-ZPMRG-F-V1.0   99.9%  99.8%       ACACCCCATCCTAAAGTAAGGTCAGCTAAATAAGCTATCGGGCCCATACCCCGAAAATGTTGGTTATACCCTTCCCGTAC
 5 AGENOME-ZPMRG-Z-V1.0   99.9%  99.7%       ACACCCCATCCTAAAGTAAGGTCAGCTAAATAAGCTATCGGGCCCATACCCCGAAAATGTTGGTTATACCCTTCCCGTAC
 6 AGENOME-ZPMRG-T-V1.0   99.9%  99.8%       ACACCCCATCCTAAAGTAAGGTCAGCTAAATAAGCTATCGGGCCCATACCCCGAAAATGTTGGTTATACCCTTCCCGTAC
 7 AGENOME-ZPMRG-HV-V1.0  99.9%  99.8%       ACACCCCATCCTAAAGTAAGGTCAGCTAAATAAGCTATCGGGCCCATACCCCGAAAATGTTGGTTATACCCTTCCCGTAC
 8 AGENOME-ZPMRG-A-V1.0   99.9%  99.8%       ACACCCCATCCTAAAGTAAGGTCAGCTAAATAAGCTATCGGGCCCATACCCCGAAAATGTTGGTTATACCCTTCCCGTAC
 9 NC_012920.1            99.9%  99.8%       ACACCCCATCCTAAAGTAAGGTCAGCTAAATAAGCTATCGGGCCCATACCCCGAAAATGTTGGTTATACCCTTCCCGTAC
   consensus/100%                            ACACCCCATCCTAAAGTAAGGTCAGCTAAATAAGCTATCGGGCCCATACCCCGAAAATGTTGGTTATACCCTTCCCGTAC
   consensus/90%                             ACACCCCATCCTAAAGTAAGGTCAGCTAAATAAGCTATCGGGCCCATACCCCGAAAATGTTGGTTATACCCTTCCCGTAC
   consensus/80%                             ACACCCCATCCTAAAGTAAGGTCAGCTAAATAAGCTATCGGGCCCATACCCCGAAAATGTTGGTTATACCCTTCCCGTAC
   consensus/70%                             ACACCCCATCCTAAAGTAAGGTCAGCTAAATAAGCTATCGGGCCCATACCCCGAAAATGTTGGTTATACCCTTCCCGTAC

                            cov    pid  4481          .         5         .         .         .         .         :         . 4560
 1 AGENOME-ZPMRG-U-V1.0  100.0% 100.0%       TAATTAATCCCCTGGCCCAACCCGTCATCTACTCTACCATCTTTGCAGGCACACTCATCACAGCGCTAAGCTCGCACTGA
 2 AGENOME-ZPMCG-V1.0    100.0%  99.9%       TAATTAATCCCCTGGCCCAACCCGTCATCTACTCTACCATCTTTGCAGGCACACTCATCACAGCGCTAAGCTCGCACTGA
 3 AGENOME-ZPMRG-M-V1.0  100.0%  99.8%       TAATTAATCCCCTGGCCCAACCCGTCATCTACTCTACCATCTTTGCAGGCACACTCATCACAGCGCTAAGCTCGCACTGA
 4 AGENOME-ZPMRG-F-V1.0   99.9%  99.8%       TAATTAATCCCCTGGCCCAACCCGTCATCTACTCTACCATCTTTGCAGGCACACTCATCACAGCGCTAAGCTCGCACTGA
 5 AGENOME-ZPMRG-Z-V1.0   99.9%  99.7%       TAATTAATCCCCTGGCCCAACCCGTCATCTACTCTACCATCTTTGCAGGCACACTCATCACAGCGCTAAGCTCGCACTGA
 6 AGENOME-ZPMRG-T-V1.0   99.9%  99.8%       TAATTAATCCCCTGGCCCAACCCGTCATCTACTCTACCATCTTTGCAGGCACACTCATCACAGCGCTAAGCTCGCACTGA
 7 AGENOME-ZPMRG-HV-V1.0  99.9%  99.8%       TAATTAATCCCCTGGCCCAACCCGTCATCTACTCTACCATCTTTGCAGGCACACTCATCACAGCGCTAAGCTCGCACTGA
 8 AGENOME-ZPMRG-A-V1.0   99.9%  99.8%       TAATTAATCCCCTGGCCCAACCCGTCATCTACTCTACCATCTTTGCAGGCACACTCATCACAGCGCTAAGCTCGCACTGA
 9 NC_012920.1            99.9%  99.8%       TAATTAATCCCCTGGCCCAACCCGTCATCTACTCTACCATCTTTGCAGGCACACTCATCACAGCGCTAAGCTCGCACTGA
   consensus/100%                            TAATTAATCCCCTGGCCCAACCCGTCATCTACTCTACCATCTTTGCAGGCACACTCATCACAGCGCTAAGCTCGCACTGA
   consensus/90%                             TAATTAATCCCCTGGCCCAACCCGTCATCTACTCTACCATCTTTGCAGGCACACTCATCACAGCGCTAAGCTCGCACTGA
   consensus/80%                             TAATTAATCCCCTGGCCCAACCCGTCATCTACTCTACCATCTTTGCAGGCACACTCATCACAGCGCTAAGCTCGCACTGA
   consensus/70%                             TAATTAATCCCCTGGCCCAACCCGTCATCTACTCTACCATCTTTGCAGGCACACTCATCACAGCGCTAAGCTCGCACTGA

                            cov    pid  4561          .         .         .         6         .         .         .         . 4640
 1 AGENOME-ZPMRG-U-V1.0  100.0% 100.0%       TTTTTTACCTGAGTAGGCCTAGAAATAAACATGCTAGCTTTTATTCCAGTTCTAACCAAAAAAATAAACCCTCGTTCCAC
 2 AGENOME-ZPMCG-V1.0    100.0%  99.9%       TTTTTTACCTGAGTAGGCCTAGAAATAAACATGCTAGCTTTTATTCCAGTTCTAACCAAAAAAATAAACCCTCGTTCCAC
 3 AGENOME-ZPMRG-M-V1.0  100.0%  99.8%       TTTTTTACCTGAGTAGGCCTAGAAATAAACATGCTAGCTTTTATTCCAGTTCTAACCAAAAAAATAAACCCTCGTTCCAC
 4 AGENOME-ZPMRG-F-V1.0   99.9%  99.8%       TTTTTTACCTGAGTAGGCCTAGAAATAAACATGCTAGCTTTTATTCCAGTTCTAACCAAAAAAATAAACCCTCGTTCCAC
 5 AGENOME-ZPMRG-Z-V1.0   99.9%  99.7%       TTTTTTACCTGAGTAGGCCTAGAAATAAACATGCTAGCTTTTATTCCAGTTCTAACCAAAAAAATAAACCCTCGTTCCAC
 6 AGENOME-ZPMRG-T-V1.0   99.9%  99.8%       TTTTTTACCTGAGTAGGCCTAGAAATAAACATGCTAGCTTTTATTCCAGTTCTAACCAAAAAAATAAACCCTCGTTCCAC
 7 AGENOME-ZPMRG-HV-V1.0  99.9%  99.8%       TTTTTTACCTGAGTAGGCCTAGAAATAAACATGCTAGCTTTTATTCCAGTTCTAACCAAAAAAATAAACCCTCGTTCCAC
 8 AGENOME-ZPMRG-A-V1.0   99.9%  99.8%       TTTTTTACCTGAGTAGGCCTAGAAATAAACATGCTAGCTTTTATTCCAGTTCTAACCAAAAAAATAAACCCTCGTTCCAC
 9 NC_012920.1            99.9%  99.8%       TTTTTTACCTGAGTAGGCCTAGAAATAAACATGCTAGCTTTTATTCCAGTTCTAACCAAAAAAATAAACCCTCGTTCCAC
   consensus/100%                            TTTTTTACCTGAGTAGGCCTAGAAATAAACATGCTAGCTTTTATTCCAGTTCTAACCAAAAAAATAAACCCTCGTTCCAC
   consensus/90%                             TTTTTTACCTGAGTAGGCCTAGAAATAAACATGCTAGCTTTTATTCCAGTTCTAACCAAAAAAATAAACCCTCGTTCCAC
   consensus/80%                             TTTTTTACCTGAGTAGGCCTAGAAATAAACATGCTAGCTTTTATTCCAGTTCTAACCAAAAAAATAAACCCTCGTTCCAC
   consensus/70%                             TTTTTTACCTGAGTAGGCCTAGAAATAAACATGCTAGCTTTTATTCCAGTTCTAACCAAAAAAATAAACCCTCGTTCCAC

                            cov    pid  4641          :         .         .         .         .         7         .         . 4720
 1 AGENOME-ZPMRG-U-V1.0  100.0% 100.0%       AGAAGCTGCCATCAAGTATTTCCTCACGCAAGCAACCGCATCCATAATCCTTCTAATAGCTATCCTCTTCAACAATATAC
 2 AGENOME-ZPMCG-V1.0    100.0%  99.9%       AGAAGCTGCCATCAAGTATTTCCTCACGCAAGCAACCGCATCCATAATCCTTCTAATAGCTATCCTCTTCAACAATATAC
 3 AGENOME-ZPMRG-M-V1.0  100.0%  99.8%       AGAAGCTGCCATCAAGTATTTCCTCACGCAAGCAACCGCATCCATAATCCTTCTAATAGCTATCCTCTTCAACAATATAC
 4 AGENOME-ZPMRG-F-V1.0   99.9%  99.8%       AGAAGCTGCCATCAAGTATTTCCTCACGCAAGCAACCGCATCCATAATCCTTCTAATAGCTATCCTCTTCAACAATATAC
 5 AGENOME-ZPMRG-Z-V1.0   99.9%  99.7%       AGAAGCTGCCATCAAGTATTTCCTCACGCAAGCAACCGCATCCATAATCCTTCTAATAGCTATCCTCTTCAACAATATAC
 6 AGENOME-ZPMRG-T-V1.0   99.9%  99.8%       AGAAGCTGCCATCAAGTATTTCCTCACGCAAGCAACCGCATCCATAATCCTTCTAATAGCTATCCTCTTCAACAATATAC
 7 AGENOME-ZPMRG-HV-V1.0  99.9%  99.8%       AGAAGCTGCCATCAAGTATTTCCTCACGCAAGCAACCGCATCCATAATCCTTCTAATAGCTATCCTCTTCAACAATATAC
 8 AGENOME-ZPMRG-A-V1.0   99.9%  99.8%       AGAAGCTGCCATCAAGTATTTCCTCACGCAAGCAACCGCATCCATAATCCTTCTAATAGCTATCCTCTTCAACAATATAC
 9 NC_012920.1            99.9%  99.8%       AGAAGCTGCCATCAAGTATTTCCTCACGCAAGCAACCGCATCCATAATCCTTCTAATAGCTATCCTCTTCAACAATATAC
   consensus/100%                            AGAAGCTGCCATCAAGTATTTCCTCACGCAAGCAACCGCATCCATAATCCTTCTAATAGCTATCCTCTTCAACAATATAC
   consensus/90%                             AGAAGCTGCCATCAAGTATTTCCTCACGCAAGCAACCGCATCCATAATCCTTCTAATAGCTATCCTCTTCAACAATATAC
   consensus/80%                             AGAAGCTGCCATCAAGTATTTCCTCACGCAAGCAACCGCATCCATAATCCTTCTAATAGCTATCCTCTTCAACAATATAC
   consensus/70%                             AGAAGCTGCCATCAAGTATTTCCTCACGCAAGCAACCGCATCCATAATCCTTCTAATAGCTATCCTCTTCAACAATATAC

                            cov    pid  4721          .         .         :         .         .         .         .         8 4800
 1 AGENOME-ZPMRG-U-V1.0  100.0% 100.0%       TCTCCGGACAATGAACCATAACCAATACTACCAATCAATACTCATCATTAATAATCATAATGGCTATAGCAATAAAACTA
 2 AGENOME-ZPMCG-V1.0    100.0%  99.9%       TCTCCGGACAATGAACCATAACCAATACTACCAATCAATACTCATCATTAATAATCATAATGGCTATAGCAATAAAACTA
 3 AGENOME-ZPMRG-M-V1.0  100.0%  99.8%       TCTCCGGACAATGAACCATAACCAATACTACCAATCAATACTCATCATTAATAATCATAATGGCTATAGCAATAAAACTA
 4 AGENOME-ZPMRG-F-V1.0   99.9%  99.8%       TCTCCGGACAATGAACCATAACCAATACTACCAATCAATACTCATCATTAATAATCATAATGGCTATAGCAATAAAACTA
 5 AGENOME-ZPMRG-Z-V1.0   99.9%  99.7%       TCTCCGGACAATGAACCATAACCAATACTACCAATCAATACTCATCATTAATAATCATAATGGCTATAGCAATAAAACTA
 6 AGENOME-ZPMRG-T-V1.0   99.9%  99.8%       TCTCCGGACAATGAACCATAACCAATACTACCAATCAATACTCATCATTAATAATCATAATGGCTATAGCAATAAAACTA
 7 AGENOME-ZPMRG-HV-V1.0  99.9%  99.8%       TCTCCGGACAATGAACCATAACCAATACTACCAATCAATACTCATCATTAATAATCATAATGGCTATAGCAATAAAACTA
 8 AGENOME-ZPMRG-A-V1.0   99.9%  99.8%       TCTCCGGACAATGAACCATAACCAATACTACCAATCAATACTCATCATTAATAATCATAATGGCTATAGCAATAAAACTA
 9 NC_012920.1            99.9%  99.8%       TCTCCGGACAATGAACCATAACCAATACTACCAATCAATACTCATCATTAATAATCATAATAGCTATAGCAATAAAACTA
   consensus/100%                            TCTCCGGACAATGAACCATAACCAATACTACCAATCAATACTCATCATTAATAATCATAATuGCTATAGCAATAAAACTA
   consensus/90%                             TCTCCGGACAATGAACCATAACCAATACTACCAATCAATACTCATCATTAATAATCATAATuGCTATAGCAATAAAACTA
   consensus/80%                             TCTCCGGACAATGAACCATAACCAATACTACCAATCAATACTCATCATTAATAATCATAATGGCTATAGCAATAAAACTA
   consensus/70%                             TCTCCGGACAATGAACCATAACCAATACTACCAATCAATACTCATCATTAATAATCATAATGGCTATAGCAATAAAACTA

                            cov    pid  4801          .         .         .         .         :         .         .         . 4880
 1 AGENOME-ZPMRG-U-V1.0  100.0% 100.0%       GGAATAGCCCCCTTTCACTTCTGAGTCCCAGAGGTTACCCAAGGCACCCCTCTGACATCCGGCCTGCTTCTTCTCACATG
 2 AGENOME-ZPMCG-V1.0    100.0%  99.9%       GGAATAGCCCCCTTTCACTTCTGAGTCCCAGAGGTTACCCAAGGCACCCCTCTGACATCCGGCCTGCTTCTTCTCACATG
 3 AGENOME-ZPMRG-M-V1.0  100.0%  99.8%       GGAATAGCCCCCTTTCACTTCTGAGTCCCAGAGGTTACCCAAGGCACCCCTCTGACATCCGGCCTGCTTCTTCTCACATG
```

```
  4 AGENOME-ZPMRG-F-V1.0   99.9%  99.8%       GGAATAGCCCCCTTTCACTTCTGAGTCCCAGAGGTTACCCAAGGCACCCCTCTGACATCCGGCCTGCTTCTTCTCACATG
  5 AGENOME-ZPMRG-Z-V1.0   99.9%  99.7%       GGAATAGCCCCCTTTCACTTCTGAGTCCCAGAGGTTACCCAAGGCACCCCTCTGACATCCGGCCTGCTTCTTCTCACATG
  6 AGENOME-ZPMRG-T-V1.0   99.9%  99.8%       GGAATAGCCCCCTTTCACTTCTGAGTCCCAGAGGTTACCCAAGGCACCCCTCTGACATCCGGCCTGCTTCTTCTCACATG
  7 AGENOME-ZPMRG-HV-V1.0  99.9%  99.8%       GGAATAGCCCCCTTTCACTTCTGAGTCCCAGAGGTTACCCAAGGCACCCCTCTGACATCCGGCCTGCTTCTTCTCACATG
  8 AGENOME-ZPMRG-A-V1.0   99.9%  99.8%       GGAATAGCCCCCTTTCACTTCTGAGTCCCAGAGGTTACCCAAGGCACCCCTCTGACATCCGGCCTGCTTCTTCTCACATG
  9 NC_012920.1            99.9%  99.8%       GGAATAGCCCCCTTTCACTTCTGAGTCCCAGAGGTTACCCAAGGCACCCCTCTGACATCCGGCCTGCTTCTTCTCACATG
    consensus/100%                            GGAATAGCCCCCTTTCACTTCTGAGTCCCAGAGGTTACCCAAGGCACCCCTCTGACATCCGGCCTGCTTCTTCTCACATG
    consensus/90%                             GGAATAGCCCCCTTTCACTTCTGAGTCCCAGAGGTTACCCAAGGCACCCCTCTGACATCCGGCCTGCTTCTTCTCACATG
    consensus/80%                             GGAATAGCCCCCTTTCACTTCTGAGTCCCAGAGGTTACCCAAGGCACCCCTCTGACATCCGGCCTGCTTCTTCTCACATG
    consensus/70%                             GGAATAGCCCCCTTTCACTTCTGAGTCCCAGAGGTTACCCAAGGCACCCCTCTGACATCCGGCCTGCTTCTTCTCACATG

                           cov    pid  4881           .         9         .         .         .         .         :         . 4960
  1 AGENOME-ZPMRG-U-V1.0  100.0% 100.0%       ACAAAAACTAGCCCCCATCTCAATCATATACCAAATCTCTCCCTCACTAAACGTAAGCCTTCTCCTCACTCTCTCAATCT
  2 AGENOME-ZPMCG-V1.0    100.0%  99.9%       ACAAAAACTAGCCCCCATCTCAATCATATACCAAATCTCTCCCTCACTAAACGTAAGCCTTCTCCTCACTCTCTCAATCT
  3 AGENOME-ZPMRG-M-V1.0  100.0%  99.8%       ACAAAAACTAGCCCCCATCTCAATCATATACCAAATCTCTCCCTCACTAAACGTAAGCCTTCTCCTCACTCTCTCAATCT
  4 AGENOME-ZPMRG-F-V1.0   99.9%  99.8%       ACAAAAACTAGCCCCCATCTCAATCATATACCAAATCTCTCCCTCACTAAACGTAAGCCTTCTCCTCACTCTCTCAATCT
  5 AGENOME-ZPMRG-Z-V1.0   99.9%  99.7%       ACAAAAACTAGCCCCCATCTCAATCATATACCAAATCTCTCCCTCACTAAACGTAAGCCTTCTCCTCACTCTCTCAATCT
  6 AGENOME-ZPMRG-T-V1.0   99.9%  99.8%       ACAAAAACTAGCCCCCATCTCAATCATATACCAAATCTCTCCCTCACTAGACGTAAGCCTTCTCCTCACTCTCTCAATCT
  7 AGENOME-ZPMRG-HV-V1.0  99.9%  99.8%       ACAAAAACTAGCCCCCATCTCAATCATATACCAAATCTCTCCCTCACTAAACGTAAGCCTTCTCCTCACTCTCTCAATCT
  8 AGENOME-ZPMRG-A-V1.0   99.9%  99.8%       ACAAAAACTAGCCCCCATCTCAATCATATACCAAATCTCTCCCTCACTAAACGTAAGCCTTCTCCTCACTCTCTCAATCT
  9 NC_012920.1            99.9%  99.8%       ACAAAAACTAGCCCCCATCTCAATCATATACCAAATCTCTCCCTCACTAAACGTAAGCCTTCTCCTCACTCTCTCAATCT
    consensus/100%                            ACAAAAACTAGCCCCCATCTCAATCATATACCAAATCTCTCCCTCACTAuACGTAAGCCTTCTCCTCACTCTCTCAATCT
    consensus/90%                             ACAAAAACTAGCCCCCATCTCAATCATATACCAAATCTCTCCCTCACTAuACGTAAGCCTTCTCCTCACTCTCTCAATCT
    consensus/80%                             ACAAAAACTAGCCCCCATCTCAATCATATACCAAATCTCTCCCTCACTAAACGTAAGCCTTCTCCTCACTCTCTCAATCT
    consensus/70%                             ACAAAAACTAGCCCCCATCTCAATCATATACCAAATCTCTCCCTCACTAAACGTAAGCCTTCTCCTCACTCTCTCAATCT

                           cov    pid  4961           .         .         .         0         .         .         .         . 5040
  1 AGENOME-ZPMRG-U-V1.0  100.0% 100.0%       TATCCATCATAGCAGGCAGTTGAGGTGGATTAAACCAAACCCAGCTACGCAAAATCTTAGCATACTCCTCAATTACCCAC
  2 AGENOME-ZPMCG-V1.0    100.0%  99.9%       TATCCATCATAGCAGGCAGTTGAGGTGGATTAAACCAAACCCAGCTACGCAAAATCTTAGCATACTCCTCAATTACCCAC
  3 AGENOME-ZPMRG-M-V1.0  100.0%  99.8%       TATCCATCATAGCAGGCAGTTGAGGTGGATTAAACCAAACCCAGCTACGCAAAATCTTAGCATACTCCTCAATTACCCAC
  4 AGENOME-ZPMRG-F-V1.0   99.9%  99.8%       TATCCATCATAGCAGGCAGTTGAGGTGGATTAAACCAAACCCAGCTACGCAAAATCTTAGCATACTCCTCAATTACCCAC
  5 AGENOME-ZPMRG-Z-V1.0   99.9%  99.7%       TATCCATCATAGCAGGCAGTTGAGGTGGATTAAACCAAACCCAGCTACGCAAAATCTTAGCATACTCCTCAATTACCCAC
  6 AGENOME-ZPMRG-T-V1.0   99.9%  99.8%       TATCCATCATAGCAGGCAGTTGAGGTGGATTAAACCAAACCCAGCTACGCAAAATCTTAGCATACTCCTCAATTACCCAC
  7 AGENOME-ZPMRG-HV-V1.0  99.9%  99.8%       TATCCATCATAGCAGGCAGTTGAGGTGGATTAAACCAAACCCAGCTACGCAAAATCTTAGCATACTCCTCAATTACCCAC
  8 AGENOME-ZPMRG-A-V1.0   99.9%  99.8%       TATCCATCATAGCAGGCAGTTGAGGTGGATTAAACCAAACCCAGCTACGCAAAATCTTAGCATACTCCTCAATTACCCAC
  9 NC_012920.1            99.9%  99.8%       TATCCATCATAGCAGGCAGTTGAGGTGGATTAAACCAAACCCAGCTACGCAAAATCTTAGCATACTCCTCAATTACCCAC
    consensus/100%                            TATCCATCATAGCAGGCAGTTGAGGTGGATTAAACCAAACCCAGCTACGCAAAATCTTAGCATACTCCTCAATTACCCAC
    consensus/90%                             TATCCATCATAGCAGGCAGTTGAGGTGGATTAAACCAAACCCAGCTACGCAAAATCTTAGCATACTCCTCAATTACCCAC
    consensus/80%                             TATCCATCATAGCAGGCAGTTGAGGTGGATTAAACCAAACCCAGCTACGCAAAATCTTAGCATACTCCTCAATTACCCAC
    consensus/70%                             TATCCATCATAGCAGGCAGTTGAGGTGGATTAAACCAAACCCAGCTACGCAAAATCTTAGCATACTCCTCAATTACCCAC

                           cov    pid  5041           :         .         .         .         .         1         .         . 5120
  1 AGENOME-ZPMRG-U-V1.0  100.0% 100.0%       ATAGGATGAATAATAGCAGTTCTACCGTACAACCCTAACATAACCATTCTTAATTTAACTATTTATATTATCCTAACTAC
  2 AGENOME-ZPMCG-V1.0    100.0%  99.9%       ATAGGATGAATAATAGCAGTTCTACCGTACAACCCTAACATAACCATTCTTAATTTAACTATTTATATTATCCTAACTAC
  3 AGENOME-ZPMRG-M-V1.0  100.0%  99.8%       ATAGGATGAATAATAGCAGTTCTACCGTACAACCCTAACATAACCATTCTTAATTTAACTATTTATATTATCCTAACTAC
  4 AGENOME-ZPMRG-F-V1.0   99.9%  99.8%       ATAGGATGAATAATAGCAGTTCTACCGTACAACCCTAACATAACCATTCTTAATTTAACTATTTATATTATCCTAACTAC
  5 AGENOME-ZPMRG-Z-V1.0   99.9%  99.7%       ATAGGATGAATAATAGCAGTTCTACCGTACAACCCTAACATAACCATTCTTAATTTAACTATTTATATTATCCTAACTAC
  6 AGENOME-ZPMRG-T-V1.0   99.9%  99.8%       ATAGGATGAATAATAGCAGTTCTACCGTACAACCCTAACATAACCATTCTTAATTTAACTATTTATATTATCCTAACTAC
  7 AGENOME-ZPMRG-HV-V1.0  99.9%  99.8%       ATAGGATGAATAATAGCAGTTCTACCGTACAACCCTAACATAACCATCCTTAATTTAACTATTTATATTATCCTAACTAC
  8 AGENOME-ZPMRG-A-V1.0   99.9%  99.8%       ATAGGATGAATAATAGCAGTTCTACCGTACAACCCTAACATAACCATTCTTAATTTAACTATTTATATTATCCTAACTAC
  9 NC_012920.1            99.9%  99.8%       ATAGGATGAATAATAGCAGTTCTACCGTACAACCCTAACATAACCATTCTTAATTTAACTATTTATATTATCCTAACTAC
    consensus/100%                            ATAGGATGAATAATAGCAGTTCTACCGTACAACCCTAACATAACCATsCTTAATTTAACTATTTATATTATCCTAACTAC
    consensus/90%                             ATAGGATGAATAATAGCAGTTCTACCGTACAACCCTAACATAACCATsCTTAATTTAACTATTTATATTATCCTAACTAC
    consensus/80%                             ATAGGATGAATAATAGCAGTTCTACCGTACAACCCTAACATAACCATTCTTAATTTAACTATTTATATTATCCTAACTAC
    consensus/70%                             ATAGGATGAATAATAGCAGTTCTACCGTACAACCCTAACATAACCATTCTTAATTTAACTATTTATATTATCCTAACTAC

                           cov    pid  5121           .         .         :         .         .         .         .         2 5200
  1 AGENOME-ZPMRG-U-V1.0  100.0% 100.0%       TACCGCATTCCTACTACTCAACTTAAACTCCAGCACCACGACCCTACTACTATCTCGCACCTGAAACAAGCTAACATGAC
  2 AGENOME-ZPMCG-V1.0    100.0%  99.9%       TACCGCATTCCTACTACTCAACTTAAACTCCAGCACCACGACCCTACTACTATCTCGCACCTGAAACAAGCTAACATGAC
  3 AGENOME-ZPMRG-M-V1.0  100.0%  99.8%       TACCGCATTCCTACTACTCAACTTAAACTCCAGCACCACGACCCTACTACTATCTCGCACCTGAAACAAGCTAACATGAC
  4 AGENOME-ZPMRG-F-V1.0   99.9%  99.8%       TACCGCATTCCTACTACTCAACTTAAACTCCAGCACCACGACCCTACTACTATCTCGCACCTGAAACAAGCTAACATGAC
  5 AGENOME-ZPMRG-Z-V1.0   99.9%  99.7%       TACCGCATTCCTACTACTCAACTTAAACTCCAGCACCACGACCCTACTACTATCTCGCACCTGAAACAAGCTAACATGAC
  6 AGENOME-ZPMRG-T-V1.0   99.9%  99.8%       TACCGCATTCCTACTACTCAACTTAAACTCCAGCACCACGACCCTACTACTATCTCGCACCTGAAACAAGCTAACATGAC
  7 AGENOME-ZPMRG-HV-V1.0  99.9%  99.8%       TACCGCATTCCTACTACTCAACTTAAACTCCAGCACCACGACCCTACTACTATCTCGCACCTGAAACAAGCTAACATGAC
  8 AGENOME-ZPMRG-A-V1.0   99.9%  99.8%       TACCGCATTCCTACTACTCAACTTAAACTCCAGCACCACGACCCTACTACTATCTCGCACCTGAAACAAGCTAACATGAC
  9 NC_012920.1            99.9%  99.8%       TACCGCATTCCTACTACTCAACTTAAACTCCAGCACCACGACCCTACTACTATCTCGCACCTGAAACAAGCTAACATGAC
    consensus/100%                            TACCGCATTCCTACTACTCAACTTAAACTCCAGCACCACGACCCTACTACTATCTCGCACCTGAAACAAGCTAACATGAC
    consensus/90%                             TACCGCATTCCTACTACTCAACTTAAACTCCAGCACCACGACCCTACTACTATCTCGCACCTGAAACAAGCTAACATGAC
    consensus/80%                             TACCGCATTCCTACTACTCAACTTAAACTCCAGCACCACGACCCTACTACTATCTCGCACCTGAAACAAGCTAACATGAC
    consensus/70%                             TACCGCATTCCTACTACTCAACTTAAACTCCAGCACCACGACCCTACTACTATCTCGCACCTGAAACAAGCTAACATGAC

                           cov    pid  5201           .         .         .         .         :         .         .         . 5280
  1 AGENOME-ZPMRG-U-V1.0  100.0% 100.0%       TAACACCCTTAATTCCATCCACCCTCCTCTCCCTAGGAGGCCTGCCCCCGCTAACCGGCTTTTTGCCCAAATGGGCCATT
  2 AGENOME-ZPMCG-V1.0    100.0%  99.9%       TAACACCCTTAATTCCATCCACCCTCCTCTCCCTAGGAGGCCTGCCCCCGCTAACCGGCTTTTTGCCCAAATGGGCCATT
  3 AGENOME-ZPMRG-M-V1.0  100.0%  99.8%       TAACACCCTTAATTCCATCCACCCTCCTCTCCCTAGGAGGCCTGCCCCCGCTAACCGGCTTTTTGCCCAAATGGGCCATT
  4 AGENOME-ZPMRG-F-V1.0   99.9%  99.8%       TAACACCCTTAATTCCATCCACCCTCCTCTCCCTAGGAGGCCTGCCCCCGCTAACCGGCTTTTTGCCCAAATGGGCCATT
  5 AGENOME-ZPMRG-Z-V1.0   99.9%  99.7%       TAACACCCTTAATTCCATCCACCCTCCTCTCCCTAGGAGGCCTGCCCCCGCTAACCGGCTTTTTGCCCAAATGGGCCATT
  6 AGENOME-ZPMRG-T-V1.0   99.9%  99.8%       TAACACCCTTAATTCCATCCACCCTCCTCTCCCTAGGAGGCCTGCCCCCGCTAACCGGCTTTTTGCCCAAATGGGCCATT
  7 AGENOME-ZPMRG-HV-V1.0  99.9%  99.8%       TAACACCCTTAATTCCATCCACCCTCCTCTCCCTAGGAGGCCTGCCCCCGCTAACCGGCTTTTTGCCCAAATGGGCCATT
  8 AGENOME-ZPMRG-A-V1.0   99.9%  99.8%       TAACACCCTTAATTCCATCCACCCTCCTCTCCCTAGGAGGCCTGCCCCCGCTAACCGGCTTTTTGCCCAAATGGGCCATT
  9 NC_012920.1            99.9%  99.8%       TAACACCCTTAATTCCATCCACCCTCCTCTCCCTAGGAGGCCTGCCCCCGCTAACCGGCTTTTTGCCCAAATGGGCCATT
    consensus/100%                            TAACACCCTTAATTCCATCCACCCTCCTCTCCCTAGGAGGCCTGCCCCCGCTAACCGGCTTTTTGCCCAAATGGGCCATT
    consensus/90%                             TAACACCCTTAATTCCATCCACCCTCCTCTCCCTAGGAGGCCTGCCCCCGCTAACCGGCTTTTTGCCCAAATGGGCCATT
    consensus/80%                             TAACACCCTTAATTCCATCCACCCTCCTCTCCCTAGGAGGCCTGCCCCCGCTAACCGGCTTTTTGCCCAAATGGGCCATT
    consensus/70%                             TAACACCCTTAATTCCATCCACCCTCCTCTCCCTAGGAGGCCTGCCCCCGCTAACCGGCTTTTTGCCCAAATGGGCCATT

                           cov    pid  5281           .         3         .         .         .         .         :         . 5360
  1 AGENOME-ZPMRG-U-V1.0  100.0% 100.0%       ATCGAAGAATTCACAAAAAACAATAGCCTCATCATCCCCACCATCATAGCCACCATCACCCTCCTTAACCTCTACTTCTA
  2 AGENOME-ZPMCG-V1.0    100.0%  99.9%       ATCGAAGAATTCACAAAAAACAATAGCCTCATCATCCCCACCATCATAGCCACCATCACCCTCCTTAACCTCTACTTCTA
  3 AGENOME-ZPMRG-M-V1.0  100.0%  99.8%       ATCGAAGAATTCACAAAAAACAATAGCCTCATCATCCCCACCATCATAGCCACCATCACCCTCCTTAACCTCTACTTCTA
  4 AGENOME-ZPMRG-F-V1.0   99.9%  99.8%       ATCGAAGAATTCACAAAAAACAATAGCCTCATCATCCCCACCATCATAGCCACCATCACCCTCCTTAACCTCTACTTCTA
```

```
  5 AGENOME-ZPMRG-Z-V1.0   99.9%  99.7%       ATCGAAGAATTCACAAAAAACAATAGCCTCATCATCCCCACCATCATAGCCACCATCACCCTCCTTAACCTCTACTTCTA
  6 AGENOME-ZPMRG-T-V1.0   99.9%  99.8%       ATCGAAGAATTCACAAAAAACAATAGCCTCATCATCCCCACCATCATAGCCACCATCACCCTCCTTAACCTCTACTTCTA
  7 AGENOME-ZPMRG-HV-V1.0  99.9%  99.8%       ATCGAAGAATTCACAAAAAACAATAGCCTCATCATCCCCACCATCATAGCCACCATCACCCTCCTTAACCTCTACTTCTA
  8 AGENOME-ZPMRG-A-V1.0   99.9%  99.8%       ATCGAAGAATTCACAAAAAACAATAGCCTCATCATCCCCACCATCATAGCCACCATCACCCTCCTTAACCTCTACTTCTA
  9 NC_012920.1            99.9%  99.8%       ATCGAAGAATTCACAAAAAACAATAGCCTCATCATCCCCACCATCATAGCCACCATCACCCTCCTTAACCTCTACTTCTA
    consensus/100%                            ATCGAAGAATTCACAAAAAACAATAGCCTCATCATCCCCACCATCATAGCCACCATCACCCTCCTTAACCTCTACTTCTA
    consensus/90%                             ATCGAAGAATTCACAAAAAACAATAGCCTCATCATCCCCACCATCATAGCCACCATCACCCTCCTTAACCTCTACTTCTA
    consensus/80%                             ATCGAAGAATTCACAAAAAACAATAGCCTCATCATCCCCACCATCATAGCCACCATCACCCTCCTTAACCTCTACTTCTA
    consensus/70%                             ATCGAAGAATTCACAAAAAACAATAGCCTCATCATCCCCACCATCATAGCCACCATCACCCTCCTTAACCTCTACTTCTA

                            cov    pid  5361          .         .         .         4         .         .         .         . 5440
  1 AGENOME-ZPMRG-U-V1.0  100.0% 100.0%       CCTACGCCTAATCTACTCCACCTCAATCACACTACTCCCCATATCTAACAACGTAAAAATAAAATGACAGTTTGAACATA
  2 AGENOME-ZPMCG-V1.0    100.0%  99.9%       CCTACGCCTAATCTACTCCACCTCAATCACACTACTCCCCATATCTAACAACGTAAAAATAAAATGACAGTTTGAACATA
  3 AGENOME-ZPMRG-M-V1.0  100.0%  99.8%       CCTACGCCTAATCTACTCCACCTCAATCACACTACTCCCCATATCTAACAACGTAAAAATAAAATGACAGTTTGAACATA
  4 AGENOME-ZPMRG-F-V1.0   99.9%  99.8%       CCTACGCCTAATCTACTCCACCTCAATCACACTACTCCCCATATCTAACAACGTAAAAATAAAATGACAGTTTGAACATA
  5 AGENOME-ZPMRG-Z-V1.0   99.9%  99.7%       CCTACGCCTAATCTACTCCACCTCAATCACACTACTCCCCATATCTAACAACGTAAAAATAAAATGACAGTTTGAACATA
  6 AGENOME-ZPMRG-T-V1.0   99.9%  99.8%       CCTACGCCTAATCTACTCCACCTCAATCACACTACTCCCCATATCTAACAACGTAAAAATAAAATGACAGTTTGAACATA
  7 AGENOME-ZPMRG-HV-V1.0  99.9%  99.8%       CCTACGCCTAATCTACTCCACCTCAATCACACTACTCCCCATATCTAACAACGTAAAAATAAAATGACAGTTTGAACATA
  8 AGENOME-ZPMRG-A-V1.0   99.9%  99.8%       CCTACGCCTAATCTACTCCACCTCAATCACACTACTCCCCATATCTAACAACGTAAAAATAAAATGACAGTTTGAACATA
  9 NC_012920.1            99.9%  99.8%       CCTACGCCTAATCTACTCCACCTCAATCACACTACTCCCCATATCTAACAACGTAAAAATAAAATGACAGTTTGAACATA
    consensus/100%                            CCTACGCCTAATCTACTCCACCTCAATCACACTACTCCCCATATCTAACAACGTAAAAATAAAATGACAGTTTGAACATA
    consensus/90%                             CCTACGCCTAATCTACTCCACCTCAATCACACTACTCCCCATATCTAACAACGTAAAAATAAAATGACAGTTTGAACATA
    consensus/80%                             CCTACGCCTAATCTACTCCACCTCAATCACACTACTCCCCATATCTAACAACGTAAAAATAAAATGACAGTTTGAACATA
    consensus/70%                             CCTACGCCTAATCTACTCCACCTCAATCACACTACTCCCCATATCTAACAACGTAAAAATAAAATGACAGTTTGAACATA

                            cov    pid  5441          :         .         .         .         .         5         .         . 5520
  1 AGENOME-ZPMRG-U-V1.0  100.0% 100.0%       CAAAACCCACCCCATTCCTCCCCACACTCATCGCCCTTACCACGCTACTCCTACCTATCTCCCCTTTTATACTAATAATC
  2 AGENOME-ZPMCG-V1.0    100.0%  99.9%       CAAAACCCACCCCATTCCTCCCCACACTCATCGCCCTTACCACGCTACTCCTACCTATCTCCCCTTTTATACTAATAATC
  3 AGENOME-ZPMRG-M-V1.0  100.0%  99.8%       CAAAACCCACCCCATTCCTCCCCACACTCATCGCCCTTACCACGCTACTCCTACCTATCTCCCCTTTTATACTAATAATC
  4 AGENOME-ZPMRG-F-V1.0   99.9%  99.8%       CAAAACCCACCCCATTCCTCCCCACACTCATCGCCCTTACCACGCTACTCCTACCTATCTCCCCTTTTATACTAATAATC
  5 AGENOME-ZPMRG-Z-V1.0   99.9%  99.7%       CAAAACCCACCCCATTCCTCCCCACACTCATCGCCCTTACCACGCTACTCCTACCTATCTCCCCTTTTATACTAATAATC
  6 AGENOME-ZPMRG-T-V1.0   99.9%  99.8%       CAAAACCCACCCCATTCCTCCCCACACTCATCGCCCTTACCACGCTACTCCTACCTATCTCCCCTTTTATACTAATAATC
  7 AGENOME-ZPMRG-HV-V1.0  99.9%  99.8%       CAAAACCCACCCCATTCCTCCCCACACTCATCGCCCTTACCACGCTACTCCTACCTATCTCCCCTTTTATACTAATAATC
  8 AGENOME-ZPMRG-A-V1.0   99.9%  99.8%       CAAAACCCACCCCATTCCTCCCCACACTCATCGCCCTTACCACGCTACTCCTACCTATCTCCCCTTTTATACTAATAATC
  9 NC_012920.1            99.9%  99.8%       CAAAACCCACCCCATTCCTCCCCACACTCATCGCCCTTACCACGCTACTCCTACCTATCTCCCCTTTTATACTAATAATC
    consensus/100%                            CAAAACCCACCCCATTCCTCCCCACACTCATCGCCCTTACCACGCTACTCCTACCTATCTCCCCTTTTATACTAATAATC
    consensus/90%                             CAAAACCCACCCCATTCCTCCCCACACTCATCGCCCTTACCACGCTACTCCTACCTATCTCCCCTTTTATACTAATAATC
    consensus/80%                             CAAAACCCACCCCATTCCTCCCCACACTCATCGCCCTTACCACGCTACTCCTACCTATCTCCCCTTTTATACTAATAATC
    consensus/70%                             CAAAACCCACCCCATTCCTCCCCACACTCATCGCCCTTACCACGCTACTCCTACCTATCTCCCCTTTTATACTAATAATC

                            cov    pid  5521          .         .         :         .         .         .         .         6 5600
  1 AGENOME-ZPMRG-U-V1.0  100.0% 100.0%       TTATAGAAATTTAGGTTAAATACAGACCAAGAGCCTTCAAAGCCCTCAGTAAGTTGCAATACTTAATTTCTGTAACAGCT
  2 AGENOME-ZPMCG-V1.0    100.0%  99.9%       TTATAGAAATTTAGGTTAAATACAGACCAAGAGCCTTCAAAGCCCTCAGTAAGTTGCAATACTTAATTTCTGTAACAGCT
  3 AGENOME-ZPMRG-M-V1.0  100.0%  99.8%       TTATAGAAATTTAGGTTAAATACAGACCAAGAGCCTTCAAAGCCCTCAGTAAGTTGCAATACTTAATTTCTGTAACAGCT
  4 AGENOME-ZPMRG-F-V1.0   99.9%  99.8%       TTATAGAAATTTAGGTTAAATACAGACCAAGAGCCTTCAAAGCCCTCAGTAAGTTGCAATACTTAATTTCTGTAACAGCT
  5 AGENOME-ZPMRG-Z-V1.0   99.9%  99.7%       TTATAGAAATTTAGGTTAAATACAGACCAAGAGCCTTCAAAGCCCTCAGTAAGTTGCAATACTTAATTTCTGTAACAGCT
  6 AGENOME-ZPMRG-T-V1.0   99.9%  99.8%       TTATAGAAATTTAGGTTAAATACAGACCAAGAGCCTTCAAAGCCCTCAGTAAGTTGCAATACTTAATTTCTGTAACAGCT
  7 AGENOME-ZPMRG-HV-V1.0  99.9%  99.8%       TTATAGAAATTTAGGTTAAATACAGACCAAGAGCCTTCAAAGCCCTCAGTAAGTTGCAATACTTAATTTCTGTAACAGCT
  8 AGENOME-ZPMRG-A-V1.0   99.9%  99.8%       TTATAGAAATTTAGGTTAAATACAGACCAAGAGCCTTCAAAGCCCTCAGTAAGTTGCAATACTTAATTTCTGTAACAGCT
  9 NC_012920.1            99.9%  99.8%       TTATAGAAATTTAGGTTAAATACAGACCAAGAGCCTTCAAAGCCCTCAGTAAGTTGCAATACTTAATTTCTGTAACAGCT
    consensus/100%                            TTATAGAAATTTAGGTTAAATACAGACCAAGAGCCTTCAAAGCCCTCAGTAAGTTGCAATACTTAATTTCTGTAACAGCT
    consensus/90%                             TTATAGAAATTTAGGTTAAATACAGACCAAGAGCCTTCAAAGCCCTCAGTAAGTTGCAATACTTAATTTCTGTAACAGCT
    consensus/80%                             TTATAGAAATTTAGGTTAAATACAGACCAAGAGCCTTCAAAGCCCTCAGTAAGTTGCAATACTTAATTTCTGTAACAGCT
    consensus/70%                             TTATAGAAATTTAGGTTAAATACAGACCAAGAGCCTTCAAAGCCCTCAGTAAGTTGCAATACTTAATTTCTGTAACAGCT

                            cov    pid  5601          .         .         .         .         :         .         .         . 5680
  1 AGENOME-ZPMRG-U-V1.0  100.0% 100.0%       AAGGACTGCAAAACCCCACTCTGCATCAACTGAACGCAAATCAGCCACTTTAATTAAGCTAAGCCCTTACTAGACCAATG
  2 AGENOME-ZPMCG-V1.0    100.0%  99.9%       AAGGACTGCAAAACCCCACTCTGCATCAACTGAACGCAAATCAGCCACTTTAATTAAGCTAAGCCCTTACTAGACCAATG
  3 AGENOME-ZPMRG-M-V1.0  100.0%  99.8%       AAGGACTGCAAAACCCCACTCTGCATCAACTGAACGCAAATCAGCCACTTTAATTAAGCTAAGCCCTTACTAGACCAATG
  4 AGENOME-ZPMRG-F-V1.0   99.9%  99.8%       AAGGACTGCAAAACCCCACTCTGCATCAACTGAACGCAAATCAGCCACTTTAATTAAGCTAAGCCCTTACTAGACCAATG
  5 AGENOME-ZPMRG-Z-V1.0   99.9%  99.7%       AAGGACTGCAAAACCCCACTCTGCATCAACTGAACGCAAATCAGCCACTTTAATTAAGCTAAGCCCTTACTAGACCAATG
  6 AGENOME-ZPMRG-T-V1.0   99.9%  99.8%       AAGGACTGCAAAACCCCACTCTGCATCAACTGAACGCAAATCAGCCACTTTAATTAAGCTAAGCCCTTACTAGACCAATG
  7 AGENOME-ZPMRG-HV-V1.0  99.9%  99.8%       AAGGACTGCAAAACCCCACTCTGCATCAACTGAACGCAAATCAGCCACTTTAATTAAGCTAAGCCCTTACTAGACCAATG
  8 AGENOME-ZPMRG-A-V1.0   99.9%  99.8%       AAGGACTGCAAAACCCCACTCTGCATCAACTGAACGCAAATCAGCCACTTTAATTAAGCTAAGCCCTTACTAGACCAATG
  9 NC_012920.1            99.9%  99.8%       AAGGACTGCAAAACCCCACTCTGCATCAACTGAACGCAAATCAGCCACTTTAATTAAGCTAAGCCCTTACTAGACCAATG
    consensus/100%                            AAGGACTGCAAAACCCCACTCTGCATCAACTGAACGCAAATCAGCCACTTTAATTAAGCTAAGCCCTTACTAGACCAATG
    consensus/90%                             AAGGACTGCAAAACCCCACTCTGCATCAACTGAACGCAAATCAGCCACTTTAATTAAGCTAAGCCCTTACTAGACCAATG
    consensus/80%                             AAGGACTGCAAAACCCCACTCTGCATCAACTGAACGCAAATCAGCCACTTTAATTAAGCTAAGCCCTTACTAGACCAATG
    consensus/70%                             AAGGACTGCAAAACCCCACTCTGCATCAACTGAACGCAAATCAGCCACTTTAATTAAGCTAAGCCCTTACTAGACCAATG

                            cov    pid  5681          .         7         .         .         .         .         :         . 5760
  1 AGENOME-ZPMRG-U-V1.0  100.0% 100.0%       GGACTTAAACCCACAAACACTTAGTTAACAGCTAAGCACCCTAATCAACTGGCTTCAATCTACTTCTCCCGCCGCCGGGA
  2 AGENOME-ZPMCG-V1.0    100.0%  99.9%       GGACTTAAACCCACAAACACTTAGTTAACAGCTAAGCACCCTAATCAACTGGCTTCAATCTACTTCTCCCGCCGCCGGGA
  3 AGENOME-ZPMRG-M-V1.0  100.0%  99.8%       GGACTTAAACCCACAAACACTTAGTTAACAGCTAAGCACCCTAATCAACTGGCTTCAATCTACTTCTCCCGCCGCCGGGA
  4 AGENOME-ZPMRG-F-V1.0   99.9%  99.8%       GGACTTAAACCCACAAACACTTAGTTAACAGCTAAGCACCCTAATCAACTGGCTTCAATCTACTTCTCCCGCCGCCGGGA
  5 AGENOME-ZPMRG-Z-V1.0   99.9%  99.7%       GGACTTAAACCCACAAACACTTAGTTAACAGCTAAGCACCCTAATCAACTGGCTTCAATCTACTTCTCCCGCCGCCGGGA
  6 AGENOME-ZPMRG-T-V1.0   99.9%  99.8%       GGACTTAAACCCACAAACACTTAGTTAACAGCTAAGCACCCTAATCAACTGGCTTCAATCTACTTCTCCCGCCGCCGGGA
  7 AGENOME-ZPMRG-HV-V1.0  99.9%  99.8%       GGACTTAAACCCACAAACACTTAGTTAACAGCTAAGCACCCTAATCAACTGGCTTCAATCTACTTCTCCCGCCGCCGGGA
  8 AGENOME-ZPMRG-A-V1.0   99.9%  99.8%       GGACTTAAACCCACAAACACTTAGTTAACAGCTAAGCACCCTAATCAACTGGCTTCAATCTACTTCTCCCGCCGCCGGGA
  9 NC_012920.1            99.9%  99.8%       GGACTTAAACCCACAAACACTTAGTTAACAGCTAAGCACCCTAATCAACTGGCTTCAATCTACTTCTCCCGCCGCCGGGA
    consensus/100%                            GGACTTAAACCCACAAACACTTAGTTAACAGCTAAGCACCCTAATCAACTGGCTTCAATCTACTTCTCCCGCCGCCGGGA
    consensus/90%                             GGACTTAAACCCACAAACACTTAGTTAACAGCTAAGCACCCTAATCAACTGGCTTCAATCTACTTCTCCCGCCGCCGGGA
    consensus/80%                             GGACTTAAACCCACAAACACTTAGTTAACAGCTAAGCACCCTAATCAACTGGCTTCAATCTACTTCTCCCGCCGCCGGGA
    consensus/70%                             GGACTTAAACCCACAAACACTTAGTTAACAGCTAAGCACCCTAATCAACTGGCTTCAATCTACTTCTCCCGCCGCCGGGA

                            cov    pid  5761          .         .         .         8         .         .         .         . 5840
  1 AGENOME-ZPMRG-U-V1.0  100.0% 100.0%       AAAAAGGCGGGAGAAGCCCCGGCAGGTTTGAAGCTGCTTCTTCGAATTTGCAATTCAATATGAAAATCACCTCGGAGCTG
  2 AGENOME-ZPMCG-V1.0    100.0%  99.9%       AAAAAGGCGGGAGAAGCCCCGGCAGGTTTGAAGCTGCTTCTTCGAATTTGCAATTCAATATGAAAATCACCTCGGAGCTG
  3 AGENOME-ZPMRG-M-V1.0  100.0%  99.8%       AAAAAGGCGGGAGAAGCCCCGGCAGGTTTGAAGCTGCTTCTTCGAATTTGCAATTCAATATGAAAATCACCTCGGAGCTG
  4 AGENOME-ZPMRG-F-V1.0   99.9%  99.8%       AAAAAGGCGGGAGAAGCCCCGGCAGGTTTGAAGCTGCTTCTTCGAATTTGCAATTCAATATGAAAATCACCTCGGAGCTG
  5 AGENOME-ZPMRG-Z-V1.0   99.9%  99.7%       AAAAAGGCGGGAGAAGCCCCGGCAGGTTTGAAGCTGCTTCTTCGAATTTGCAATTCAATATGAAAATCACCTCGGAGCTG
```

```
  6 AGENOME-ZPMRG-T-V1.0   99.9%  99.8%        AAAAAGGCGGGAGAAGCCCCGGCAGGTTTGAAGCTGCTTCTTCGAATTTGCAATTCAATATGAAAATCACCTCGGAGCTG
  7 AGENOME-ZPMRG-HV-V1.0  99.9%  99.8%        AAAAAGGCGGGAGAAGCCCCGGCAGGTTTGAAGCTGCTTCTTCGAATTTGCAATTCAATATGAAAATCACCTCGGAGCTG
  8 AGENOME-ZPMRG-A-V1.0   99.9%  99.8%        AAAAAGGCGGGAGAAGCCCCGGCAGGTTTGAAGCTGCTTCTTCGAATTTGCAATTCAATATGAAAATCACCTCGGAGCTG
  9 NC_012920.1            99.9%  99.8%        AAAAAGGCGGGAGAAGCCCCGGCAGGTTTGAAGCTGCTTCTTCGAATTTGCAATTCAATATGAAAATCACCTCGGAGCTG
    consensus/100%                             AAAAAGGCGGGAGAAGCCCCGGCAGGTTTGAAGCTGCTTCTTCGAATTTGCAATTCAATATGAAAATCACCTCGGAGCTG
    consensus/90%                              AAAAAGGCGGGAGAAGCCCCGGCAGGTTTGAAGCTGCTTCTTCGAATTTGCAATTCAATATGAAAATCACCTCGGAGCTG
    consensus/80%                              AAAAAGGCGGGAGAAGCCCCGGCAGGTTTGAAGCTGCTTCTTCGAATTTGCAATTCAATATGAAAATCACCTCGGAGCTG
    consensus/70%                              AAAAAGGCGGGAGAAGCCCCGGCAGGTTTGAAGCTGCTTCTTCGAATTTGCAATTCAATATGAAAATCACCTCGGAGCTG

                             cov    pid  5841          :         .         .         .         .         9         .         . 5920
  1 AGENOME-ZPMRG-U-V1.0  100.0% 100.0%        GTAAAAAGAGGCCTAACCCCTGTCTTTAGATTTACAGTCCAATGCTTCACTCAGCCATTTTACCTCACCCCCACTGATGT
  2 AGENOME-ZPMCG-V1.0    100.0%  99.9%        GTAAAAAGAGGCCTAACCCCTGTCTTTAGATTTACAGTCCAATGCTTCACTCAGCCATTTTACCTCACCCCCACTGATGT
  3 AGENOME-ZPMRG-M-V1.0  100.0%  99.8%        GTAAAAAGAGGCCTAACCCCTGTCTTTAGATTTACAGTCCAATGCTTCACTCAGCCATTTTACCTCACCCCCACTGATGT
  4 AGENOME-ZPMRG-F-V1.0   99.9%  99.8%        GTAAAAAGAGGCCTAACCCCTGTCTTTAGATTTACAGTCCAATGCTTCACTCAGCCATTTTACCTCACCCCCACTGATGT
  5 AGENOME-ZPMRG-Z-V1.0   99.9%  99.7%        GTAAAAAGAGGCCTAACCCCTGTCTTTAGATTTACAGTCCAATGCTTCACTCAGCCATTTTACCTCACCCCCACTGATGT
  6 AGENOME-ZPMRG-T-V1.0   99.9%  99.8%        GTAAAAAGAGGCCTAACCCCTGTCTTTAGATTTACAGTCCAATGCTTCACTCAGCCATTTTACCTCACCCCCACTGATGT
  7 AGENOME-ZPMRG-HV-V1.0  99.9%  99.8%        GTAAAAAGAGGCCTAACCCCTGTCTTTAGATTTACAGTCCAATGCTTCACTCAGCCATTTTACCTCACCCCCACTGATGT
  8 AGENOME-ZPMRG-A-V1.0   99.9%  99.8%        GTAAAAAGAGGCCTAACCCCTGTCTTTAGATTTACAGTCCAATGCTTCACTCAGCCATTTTACCTCACCCCCACTGATGT
  9 NC_012920.1            99.9%  99.8%        GTAAAAAGAGGCCTAACCCCTGTCTTTAGATTTACAGTCCAATGCTTCACTCAGCCATTTTACCTCACCCCCACTGATGT
    consensus/100%                             GTAAAAAGAGGCCTAACCCCTGTCTTTAGATTTACAGTCCAATGCTTCACTCAGCCATTTTACCTCACCCCCACTGATGT
    consensus/90%                              GTAAAAAGAGGCCTAACCCCTGTCTTTAGATTTACAGTCCAATGCTTCACTCAGCCATTTTACCTCACCCCCACTGATGT
    consensus/80%                              GTAAAAAGAGGCCTAACCCCTGTCTTTAGATTTACAGTCCAATGCTTCACTCAGCCATTTTACCTCACCCCCACTGATGT
    consensus/70%                              GTAAAAAGAGGCCTAACCCCTGTCTTTAGATTTACAGTCCAATGCTTCACTCAGCCATTTTACCTCACCCCCACTGATGT

                             cov    pid  5921          .         .         :         .         .         .         .         0 6000
  1 AGENOME-ZPMRG-U-V1.0  100.0% 100.0%        TCGCCGACCGTTGACTATTCTCTACAAACCACAAAGACATTGGAACACTATACCTATTATTCGGCGCATGAGCTGGAGTC
  2 AGENOME-ZPMCG-V1.0    100.0%  99.9%        TCGCCGACCGTTGACTATTCTCTACAAACCACAAAGACATTGGAACACTATACCTATTATTCGGCGCATGAGCTGGAGTC
  3 AGENOME-ZPMRG-M-V1.0  100.0%  99.8%        TCGCCGACCGTTGACTATTCTCTACAAACCACAAAGACATTGGAACACTATACCTATTATTCGGCGCATGAGCTGGAGTC
  4 AGENOME-ZPMRG-F-V1.0   99.9%  99.8%        TCGCCGACCGTTGACTATTCTCTACAAACCACAAAGACATTGGAACACTATACCTATTATTCGGCGCATGAGCTGGAGTC
  5 AGENOME-ZPMRG-Z-V1.0   99.9%  99.7%        TCGCCGACCGTTGACTATTCTCTACAAACCACAAAGACATTGGAACACTATACCTATTATTCGGCGCATGAGCTGGAGTC
  6 AGENOME-ZPMRG-T-V1.0   99.9%  99.8%        TCGCCGACCGTTGACTATTCTCTACAAACCACAAAGACATTGGAACACTATACCTATTATTCGGCGCATGAGCTGGAGTC
  7 AGENOME-ZPMRG-HV-V1.0  99.9%  99.8%        TCGCCGACCGTTGACTATTCTCTACAAACCACAAAGACATTGGAACACTATACCTATTATTCGGCGCATGAGCTGGAGTC
  8 AGENOME-ZPMRG-A-V1.0   99.9%  99.8%        TCGCCGACCGTTGACTATTCTCTACAAACCACAAAGACATTGGAACACTATACCTATTATTCGGCGCATGAGCTGGAGTC
  9 NC_012920.1            99.9%  99.8%        TCGCCGACCGTTGACTATTCTCTACAAACCACAAAGACATTGGAACACTATACCTATTATTCGGCGCATGAGCTGGAGTC
    consensus/100%                             TCGCCGACCGTTGACTATTCTCTACAAACCACAAAGACATTGGAACACTATACCTATTATTCGGCGCATGAGCTGGAGTC
    consensus/90%                              TCGCCGACCGTTGACTATTCTCTACAAACCACAAAGACATTGGAACACTATACCTATTATTCGGCGCATGAGCTGGAGTC
    consensus/80%                              TCGCCGACCGTTGACTATTCTCTACAAACCACAAAGACATTGGAACACTATACCTATTATTCGGCGCATGAGCTGGAGTC
    consensus/70%                              TCGCCGACCGTTGACTATTCTCTACAAACCACAAAGACATTGGAACACTATACCTATTATTCGGCGCATGAGCTGGAGTC

                             cov    pid  6001          .         .         .         .         :         .         .         . 6080
  1 AGENOME-ZPMRG-U-V1.0  100.0% 100.0%        CTAGGCACAGCTCTAAGCCTCCTTATTCGAGCCGAGCTGGGCCAGCCAGGCAACCTTCTAGGTAACGACCACATCTACAA
  2 AGENOME-ZPMCG-V1.0    100.0%  99.9%        CTAGGCACAGCTCTAAGCCTCCTTATTCGAGCCGAGCTGGGCCAGCCAGGCAACCTTCTAGGTAACGACCACATCTACAA
  3 AGENOME-ZPMRG-M-V1.0  100.0%  99.8%        CTAGGCACAGCTCTAAGCCTCCTTATTCGAGCCGAGCTGGGCCAGCCAGGCAACCTTCTAGGTAACGACCACATCTACAA
  4 AGENOME-ZPMRG-F-V1.0   99.9%  99.8%        CTAGGCACAGCTCTAAGCCTCCTTATTCGAGCCGAGCTGGGCCAGCCAGGCAACCTTCTAGGTAACGACCACATCTACAA
  5 AGENOME-ZPMRG-Z-V1.0   99.9%  99.7%        CTAGGCACAGCTCTAAGCCTCCTTATTCGAGCCGAGCTGGGCCAGCCAGGCAACCTTCTAGGTAACGACCACATCTACAA
  6 AGENOME-ZPMRG-T-V1.0   99.9%  99.8%        CTAGGCACAGCTCTAAGCCTCCTTATTCGAGCCGAGCTGGGCCAGCCAGGCAACCTTCTAGGTAACGACCACATCTACAA
  7 AGENOME-ZPMRG-HV-V1.0  99.9%  99.8%        CTAGGCACAGCTCTAAGCCTCCTTATTCGAGCCGAGCTGGGCCAGCCAGGCAACCTTCTAGGTAACGACCACATCTACAA
  8 AGENOME-ZPMRG-A-V1.0   99.9%  99.8%        CTAGGCACAGCTCTAAGCCTCCTTATTCGAGCCGAGCTGGGCCAGCCAGGCAACCTTCTAGGTAACGACCACATCTACAA
  9 NC_012920.1            99.9%  99.8%        CTAGGCACAGCTCTAAGCCTCCTTATTCGAGCCGAGCTGGGCCAGCCAGGCAACCTTCTAGGTAACGACCACATCTACAA
    consensus/100%                             CTAGGCACAGCTCTAAGCCTCCTTATTCGAGCCGAGCTGGGCCAGCCAGGCAACCTTCTAGGTAACGACCACATCTACAA
    consensus/90%                              CTAGGCACAGCTCTAAGCCTCCTTATTCGAGCCGAGCTGGGCCAGCCAGGCAACCTTCTAGGTAACGACCACATCTACAA
    consensus/80%                              CTAGGCACAGCTCTAAGCCTCCTTATTCGAGCCGAGCTGGGCCAGCCAGGCAACCTTCTAGGTAACGACCACATCTACAA
    consensus/70%                              CTAGGCACAGCTCTAAGCCTCCTTATTCGAGCCGAGCTGGGCCAGCCAGGCAACCTTCTAGGTAACGACCACATCTACAA

                             cov    pid  6081          .         1         .         .         .         .         :         . 6160
  1 AGENOME-ZPMRG-U-V1.0  100.0% 100.0%        CGTTATCGTCACAGCCCATGCATTTGTAATAATCTTCTTCATAGTAATACCCATCATAATCGGAGGCTTTGGCAACTGAC
  2 AGENOME-ZPMCG-V1.0    100.0%  99.9%        CGTTATCGTCACAGCCCATGCATTTGTAATAATCTTCTTCATAGTAATACCCATCATAATCGGAGGCTTTGGCAACTGAC
  3 AGENOME-ZPMRG-M-V1.0  100.0%  99.8%        CGTTATCGTCACAGCCCATGCATTTGTAATAATCTTCTTCATAGTAATACCCATCATAATCGGAGGCTTTGGCAACTGAC
  4 AGENOME-ZPMRG-F-V1.0   99.9%  99.8%        CGTTATCGTCACAGCCCATGCATTTGTAATAATCTTCTTCATAGTAATACCCATCATAATCGGAGGCTTTGGCAACTGAC
  5 AGENOME-ZPMRG-Z-V1.0   99.9%  99.7%        CGTTATCGTCACAGCCCATGCATTTGTAATAATCTTCTTCATAGTAATACCCATCATAATCGGAGGCTTTGGCAACTGAC
  6 AGENOME-ZPMRG-T-V1.0   99.9%  99.8%        CGTTATCGTCACAGCCCATGCATTTGTAATAATCTTCTTCATAGTAATACCCATCATAATCGGAGGCTTTGGCAACTGAC
  7 AGENOME-ZPMRG-HV-V1.0  99.9%  99.8%        CGTTATCGTCACAGCCCATGCATTTGTAATAATCTTTTTCATAGTAATACCCATCATAATCGGAGGCTTTGGCAACTGAC
  8 AGENOME-ZPMRG-A-V1.0   99.9%  99.8%        CGTTATCGTCACAGCCCATGCATTTGTAATAATCTTCTTCATAGTAATACCCATCATAATCGGAGGCTTTGGCAACTGAC
  9 NC_012920.1            99.9%  99.8%        CGTTATCGTCACAGCCCATGCATTTGTAATAATCTTCTTCATAGTAATACCCATCATAATCGGAGGCTTTGGCAACTGAC
    consensus/100%                             CGTTATCGTCACAGCCCATGCATTTGTAATAATCTTsTTCATAGTAATACCCATCATAATCGGAGGCTTTGGCAACTGAC
    consensus/90%                              CGTTATCGTCACAGCCCATGCATTTGTAATAATCTTsTTCATAGTAATACCCATCATAATCGGAGGCTTTGGCAACTGAC
    consensus/80%                              CGTTATCGTCACAGCCCATGCATTTGTAATAATCTTCTTCATAGTAATACCCATCATAATCGGAGGCTTTGGCAACTGAC
    consensus/70%                              CGTTATCGTCACAGCCCATGCATTTGTAATAATCTTCTTCATAGTAATACCCATCATAATCGGAGGCTTTGGCAACTGAC

                             cov    pid  6161          .         .         .         2         .         .         .         . 6240
  1 AGENOME-ZPMRG-U-V1.0  100.0% 100.0%        TAGTTCCCCTAATAATCGGTGCCCCCGATATGGCGTTTCCCCGCATAAACAACATAAGCTTCTGACTCTTACCTCCCTCT
  2 AGENOME-ZPMCG-V1.0    100.0%  99.9%        TAGTTCCCCTAATAATCGGTGCCCCCGATATGGCGTTTCCCCGCATAAACAACATAAGCTTCTGACTCTTACCTCCCTCT
  3 AGENOME-ZPMRG-M-V1.0  100.0%  99.8%        TAGTTCCCCTAATAATCGGTGCCCCCGATATGGCGTTTCCCCGCATAAACAACATAAGCTTCTGACTCTTACCTCCCTCT
  4 AGENOME-ZPMRG-F-V1.0   99.9%  99.8%        TAGTTCCCCTAATAATCGGTGCCCCCGATATGGCGTTTCCCCGCATAAACAACATAAGCTTCTGACTCTTACCTCCCTCT
  5 AGENOME-ZPMRG-Z-V1.0   99.9%  99.7%        TAGTTCCCCTAATAATCGGTGCCCCCGATATGGCGTTTCCCCGCATAAACAACATAAGCTTCTGACTCTTACCTCCCTCT
  6 AGENOME-ZPMRG-T-V1.0   99.9%  99.8%        TAGTTCCCCTAATAATCGGTGCCCCCGATATGGCGTTTCCCCGCATAAACAACATAAGCTTCTGACTCTTACCTCCCTCT
  7 AGENOME-ZPMRG-HV-V1.0  99.9%  99.8%        TAGTTCCCCTAATAATCGGTGCCCCCGATATAGCGTTTCCCCGCATAAACAACATAAGCTTCTGACTCTTACCTCCCTCT
  8 AGENOME-ZPMRG-A-V1.0   99.9%  99.8%        TAGTTCCCCTAATAATCGGTGCCCCCGATATGGCGTTTCCCCGCATAAACAACATAAGCTTCTGACTCTTACCTCCCTCT
  9 NC_012920.1            99.9%  99.8%        TAGTTCCCCTAATAATCGGTGCCCCCGATATGGCGTTTCCCCGCATAAACAACATAAGCTTCTGACTCTTACCTCCCTCT
    consensus/100%                             TAGTTCCCCTAATAATCGGTGCCCCCGATATuGCGTTTCCCCGCATAAACAACATAAGCTTCTGACTCTTACCTCCCTCT
    consensus/90%                              TAGTTCCCCTAATAATCGGTGCCCCCGATATuGCGTTTCCCCGCATAAACAACATAAGCTTCTGACTCTTACCTCCCTCT
    consensus/80%                              TAGTTCCCCTAATAATCGGTGCCCCCGATATGGCGTTTCCCCGCATAAACAACATAAGCTTCTGACTCTTACCTCCCTCT
    consensus/70%                              TAGTTCCCCTAATAATCGGTGCCCCCGATATGGCGTTTCCCCGCATAAACAACATAAGCTTCTGACTCTTACCTCCCTCT

                             cov    pid  6241          :         .         .         .         .         3         .         . 6320
  1 AGENOME-ZPMRG-U-V1.0  100.0% 100.0%        CTCCTACTCCTGCTCGCATCTGCTATAGTGGAGGCCGGAGCAGGAACAGGTTGAACAGTCTACCCTCCCTTAGCAGGGAA
  2 AGENOME-ZPMCG-V1.0    100.0%  99.9%        CTCCTACTCCTGCTCGCATCTGCTATAGTGGAGGCCGGAGCAGGAACAGGTTGAACAGTCTACCCTCCCTTAGCAGGGAA
  3 AGENOME-ZPMRG-M-V1.0  100.0%  99.8%        CTCCTACTCCTGCTCGCATCTGCTATAGTGGAGGCCGGAGCAGGAACAGGTTGAACAGTCTACCCTCCCTTAGCAGGGAA
  4 AGENOME-ZPMRG-F-V1.0   99.9%  99.8%        CTCCTACTCCTGCTCGCATCTGCTATAGTGGAGGCCGGAGCAGGAACAGGTTGAACAGTCTACCCTCCCTTAGCAGGGAA
  5 AGENOME-ZPMRG-Z-V1.0   99.9%  99.7%        CTCCTACTCCTGCTCGCATCTGCTATAGTGGAGGCCGGAGCAGGAACAGGTTGAACAGTCTACCCTCCCTTAGCAGGGAA
  6 AGENOME-ZPMRG-T-V1.0   99.9%  99.8%        CTCCTACTCCTGCTCGCATCTGCTATAGTGGAGGCCGGAGCAGGAACAGGTTGAACAGTCTACCCTCCCTTAGCAGGGAA
```

```
 7 AGENOME-ZPMRG-HV-V1.0  99.9%  99.8%       CTCCTACTCCTGCTCGCATCTGCTATAGTGGAGGCCGGAGCAGGAACAGGTTGAACAGTCTACCCTCCCTTAGCAGGGAA
 8 AGENOME-ZPMRG-A-V1.0   99.9%  99.8%       CTCCTACTCCTGCTCGCATCTGCTATAGTGGAGGCCGGAGCAGGAACAGGTTGAACAGTCTACCCTCCCTTAGCAGGGAA
 9 NC_012920.1            99.9%  99.8%       CTCCTACTCCTGCTCGCATCTGCTATAGTGGAGGCCGGAGCAGGAACAGGTTGAACAGTCTACCCTCCCTTAGCAGGGAA
   consensus/100%                            CTCCTACTCCTGCTCGCATCTGCTATAGTGGAGGCCGGAGCAGGAACAGGTTGAACAGTCTACCCTCCCTTAGCAGGGAA
   consensus/90%                             CTCCTACTCCTGCTCGCATCTGCTATAGTGGAGGCCGGAGCAGGAACAGGTTGAACAGTCTACCCTCCCTTAGCAGGGAA
   consensus/80%                             CTCCTACTCCTGCTCGCATCTGCTATAGTGGAGGCCGGAGCAGGAACAGGTTGAACAGTCTACCCTCCCTTAGCAGGGAA
   consensus/70%                             CTCCTACTCCTGCTCGCATCTGCTATAGTGGAGGCCGGAGCAGGAACAGGTTGAACAGTCTACCCTCCCTTAGCAGGGAA

                             cov    pid  6321          .         .         :         .         .         .         .         4 6400
 1 AGENOME-ZPMRG-U-V1.0  100.0% 100.0%       CTACTCCCACCCTGGAGCCTCCGTAGACCTAACCATCTTCTCCTTACACCTAGCAGGTGTCTCCTCTATCTTAGGGGCCA
 2 AGENOME-ZPMCG-V1.0    100.0%  99.9%       CTACTCCCACCCTGGAGCCTCCGTAGACCTAACCATCTTCTCCTTACACCTAGCAGGTGTCTCCTCTATCTTAGGGGCCA
 3 AGENOME-ZPMRG-M-V1.0  100.0%  99.8%       CTACTCCCACCCTGGAGCCTCCGTAGACCTAACCATCTTCTCCTTACACCTAGCAGGTGTCTCCTCTATCTTAGGGGCCA
 4 AGENOME-ZPMRG-F-V1.0   99.9%  99.8%       CTACTCCCACCCTGGAGCCTCCGTAGACCTAACCATCTTCTCCTTACACCTAGCAGGTGTCTCCTCTATCTTAGGGGCCA
 5 AGENOME-ZPMRG-Z-V1.0   99.9%  99.7%       CTACTCCCACCCTGGAGCCTCCGTAGACCTAACCATCTTCTCCTTACACCTAGCAGGTGTCTCCTCTATCTTAGGGGCCA
 6 AGENOME-ZPMRG-T-V1.0   99.9%  99.8%       CTACTCCCACCCTGGAGCCTCCGTAGACCTAACCATCTTCTCCTTACACCTAGCAGGTGTCTCCTCTATCTTAGGGGCCA
 7 AGENOME-ZPMRG-HV-V1.0  99.9%  99.8%       CTACTCCCACCCTGGAGCCTCCGTAGACCTAACCATCTTCTCCTTACACCTAGCAGGTGTCTCCTCTATCTTAGGGGCCA
 8 AGENOME-ZPMRG-A-V1.0   99.9%  99.8%       CTACTCCCACCCTGGAGCCTCCGTAGACCTAACCATCTTCTCCTTACACCTAGCAGGTGTCTCCTCTATCTTAGGGGCCA
 9 NC_012920.1            99.9%  99.8%       CTACTCCCACCCTGGAGCCTCCGTAGACCTAACCATCTTCTCCTTACACCTAGCAGGTGTCTCCTCTATCTTAGGGGCCA
   consensus/100%                            CTACTCCCACCCTGGAGCCTCCGTAGACCTAACCATCTTCTCCTTACACCTAGCAGGTGTCTCCTCTATCTTAGGGGCCA
   consensus/90%                             CTACTCCCACCCTGGAGCCTCCGTAGACCTAACCATCTTCTCCTTACACCTAGCAGGTGTCTCCTCTATCTTAGGGGCCA
   consensus/80%                             CTACTCCCACCCTGGAGCCTCCGTAGACCTAACCATCTTCTCCTTACACCTAGCAGGTGTCTCCTCTATCTTAGGGGCCA
   consensus/70%                             CTACTCCCACCCTGGAGCCTCCGTAGACCTAACCATCTTCTCCTTACACCTAGCAGGTGTCTCCTCTATCTTAGGGGCCA

                             cov    pid  6401          .         .         .         .         :         .         .         . 6480
 1 AGENOME-ZPMRG-U-V1.0  100.0% 100.0%       TCAATTTCATCACAACAATTATCAATATAAAACCCCCTGCCATAACCCAATACCAAACGCCCCTCTTCGTCTGATCCGTC
 2 AGENOME-ZPMCG-V1.0    100.0%  99.9%       TCAATTTCATCACAACAATTATCAATATAAAACCCCCTGCCATAACCCAATACCAAACGCCCCTCTTCGTCTGATCCGTC
 3 AGENOME-ZPMRG-M-V1.0  100.0%  99.8%       TCAATTTCATCACAACAATTATCAATATAAAACCCCCTGCCATAACCCAATACCAAACGCCCCTCTTCGTCTGATCCGTC
 4 AGENOME-ZPMRG-F-V1.0   99.9%  99.8%       TCAACTTCATCACAACAATTATCAATATAAAACCCCCTGCCATAACCCAATACCAAACGCCCCTCTTCGTCTGATCCGTC
 5 AGENOME-ZPMRG-Z-V1.0   99.9%  99.7%       TCAATTTCATCACAACAATTATCAATATAAAACCCCCTGCCATAACCCAATACCAAACGCCCCTCTTCGTCTGATCCGTC
 6 AGENOME-ZPMRG-T-V1.0   99.9%  99.8%       TCAATTTCATCACAACAATTATCAATATAAAACCCCCTGCCATAACCCAATACCAAACGCCCCTCTTCGTCTGATCCGTC
 7 AGENOME-ZPMRG-HV-V1.0  99.9%  99.8%       TCAATTTCATCACAACAATTATCAATATAAAACCCCCTGCCATAACCCAATACCAAACGCCCCTCTTCGTCTGATCCGTC
 8 AGENOME-ZPMRG-A-V1.0   99.9%  99.8%       TCAATTTCATCACAACAATTATCAATATAAAACCCCCTGCCATAACCCAATACCAAACGCCCCTCTTCGTCTGATCCGTC
 9 NC_012920.1            99.9%  99.8%       TCAATTTCATCACAACAATTATCAATATAAAACCCCCTGCCATAACCCAATACCAAACGCCCCTCTTCGTCTGATCCGTC
   consensus/100%                            TCAAsTTCATCACAACAATTATCAATATAAAACCCCCTGCCATAACCCAATACCAAACGCCCCTCTTCGTCTGATCCGTC
   consensus/90%                             TCAAsTTCATCACAACAATTATCAATATAAAACCCCCTGCCATAACCCAATACCAAACGCCCCTCTTCGTCTGATCCGTC
   consensus/80%                             TCAATTTCATCACAACAATTATCAATATAAAACCCCCTGCCATAACCCAATACCAAACGCCCCTCTTCGTCTGATCCGTC
   consensus/70%                             TCAATTTCATCACAACAATTATCAATATAAAACCCCCTGCCATAACCCAATACCAAACGCCCCTCTTCGTCTGATCCGTC

                             cov    pid  6481          .         5         .         .         .         .         :         . 6560
 1 AGENOME-ZPMRG-U-V1.0  100.0% 100.0%       CTAATCACAGCAGTCCTACTTCTCCTATCTCTCCCAGTCCTAGCTGCTGGCATCACTATACTACTAACAGACCGCAACCT
 2 AGENOME-ZPMCG-V1.0    100.0%  99.9%       CTAATCACAGCAGTCCTACTTCTCCTATCTCTCCCAGTCCTAGCTGCTGGCATCACTATACTACTAACAGACCGCAACCT
 3 AGENOME-ZPMRG-M-V1.0  100.0%  99.8%       CTAATCACAGCAGTCCTACTTCTCCTATCTCTCCCAGTCCTAGCTGCTGGCATCACTATACTACTAACAGACCGCAACCT
 4 AGENOME-ZPMRG-F-V1.0   99.9%  99.8%       CTAATCACAGCAGTCCTACTTCTCCTATCTCTCCCAGTCCTAGCTGCTGGCATCACTATACTACTAACAGACCGCAACCT
 5 AGENOME-ZPMRG-Z-V1.0   99.9%  99.7%       CTAATCACAGCAGTCCTACTTCTCCTATCTCTCCCAGTCCTAGCTGCTGGCATCACTATACTACTAACAGACCGCAACCT
 6 AGENOME-ZPMRG-T-V1.0   99.9%  99.8%       CTAATCACAGCAGTCCTACTTCTCCTATCTCTCCCAGTCCTAGCTGCTGGCATCACTATACTACTAACAGACCGCAACCT
 7 AGENOME-ZPMRG-HV-V1.0  99.9%  99.8%       CTAATCACAGCAGTCCTACTTCTCCTATCTCTCCCAGTCCTAGCTGCTGGCATCACTATACTACTAACAGACCGCAACCT
 8 AGENOME-ZPMRG-A-V1.0   99.9%  99.8%       CTAATCACAGCAGTCCTACTTCTCCTATCTCTCCCAGTCCTAGCTGCTGGCATCACTATACTACTAACAGACCGCAACCT
 9 NC_012920.1            99.9%  99.8%       CTAATCACAGCAGTCCTACTTCTCCTATCTCTCCCAGTCCTAGCTGCTGGCATCACTATACTACTAACAGACCGCAACCT
   consensus/100%                            CTAATCACAGCAGTCCTACTTCTCCTATCTCTCCCAGTCCTAGCTGCTGGCATCACTATACTACTAACAGACCGCAACCT
   consensus/90%                             CTAATCACAGCAGTCCTACTTCTCCTATCTCTCCCAGTCCTAGCTGCTGGCATCACTATACTACTAACAGACCGCAACCT
   consensus/80%                             CTAATCACAGCAGTCCTACTTCTCCTATCTCTCCCAGTCCTAGCTGCTGGCATCACTATACTACTAACAGACCGCAACCT
   consensus/70%                             CTAATCACAGCAGTCCTACTTCTCCTATCTCTCCCAGTCCTAGCTGCTGGCATCACTATACTACTAACAGACCGCAACCT

                             cov    pid  6561          .         .         .         6         .         .         .         . 6640
 1 AGENOME-ZPMRG-U-V1.0  100.0% 100.0%       CAACACCACCTTCTTCGACCCCGCCGGAGGAGGAGACCCCATTCTATACCAACACCTATTCTGATTTTTCGGTCACCCTG
 2 AGENOME-ZPMCG-V1.0    100.0%  99.9%       CAACACCACCTTCTTCGACCCCGCCGGAGGAGGAGACCCCATTCTATACCAACACCTATTCTGATTTTTCGGTCACCCTG
 3 AGENOME-ZPMRG-M-V1.0  100.0%  99.8%       CAACACCACCTTCTTCGACCCCGCCGGAGGAGGAGACCCCATTCTATACCAACACCTATTCTGATTTTTCGGTCACCCTG
 4 AGENOME-ZPMRG-F-V1.0   99.9%  99.8%       CAACACCACCTTCTTCGACCCCGCCGGAGGAGGAGACCCCATTCTATACCAACACCTATTCTGATTTTTCGGTCACCCTG
 5 AGENOME-ZPMRG-Z-V1.0   99.9%  99.7%       CAACACCACCTTCTTCGACCCCGCCGGAGGAGGAGACCCCATTCTATACCAACACCTATTCTGATTTTTCGGTCACCCTG
 6 AGENOME-ZPMRG-T-V1.0   99.9%  99.8%       CAACACCACCTTCTTCGACCCCGCCGGAGGAGGAGACCCCATTCTATACCAACACCTATTCTGATTTTTCGGTCACCCTG
 7 AGENOME-ZPMRG-HV-V1.0  99.9%  99.8%       CAACACCACCTTCTTCGACCCCGCCGGAGGAGGAGACCCCATTCTATACCAACACCTATTCTGATTTTTCGGTCACCCTG
 8 AGENOME-ZPMRG-A-V1.0   99.9%  99.8%       CAACACCACCTTCTTCGACCCCGCCGGAGGAGGAGACCCCATTCTATACCAACACCTATTCTGATTTTTCGGTCACCCTG
 9 NC_012920.1            99.9%  99.8%       CAACACCACCTTCTTCGACCCCGCCGGAGGAGGAGACCCCATTCTATACCAACACCTATTCTGATTTTTCGGTCACCCTG
   consensus/100%                            CAACACCACCTTCTTCGACCCCGCCGGAGGAGGAGACCCCATTCTATACCAACACCTATTCTGATTTTTCGGTCACCCTG
   consensus/90%                             CAACACCACCTTCTTCGACCCCGCCGGAGGAGGAGACCCCATTCTATACCAACACCTATTCTGATTTTTCGGTCACCCTG
   consensus/80%                             CAACACCACCTTCTTCGACCCCGCCGGAGGAGGAGACCCCATTCTATACCAACACCTATTCTGATTTTTCGGTCACCCTG
   consensus/70%                             CAACACCACCTTCTTCGACCCCGCCGGAGGAGGAGACCCCATTCTATACCAACACCTATTCTGATTTTTCGGTCACCCTG

                             cov    pid  6641          :         .         .         .         .         7         .         . 6720
 1 AGENOME-ZPMRG-U-V1.0  100.0% 100.0%       AAGTTTATATTCTTATCCTACCAGGCTTCGGAATAATCTCCCATATTGTAACTTACTACTCCGGAAAAAAAGAACCATTT
 2 AGENOME-ZPMCG-V1.0    100.0%  99.9%       AAGTTTATATTCTTATCCTACCAGGCTTCGGAATAATCTCCCATATTGTAACTTACTACTCCGGAAAAAAAGAACCATTT
 3 AGENOME-ZPMRG-M-V1.0  100.0%  99.8%       AAGTTTATATTCTTATCCTACCAGGCTTCGGAATAATCTCCCATATTGTAACTTACTACTCCGGAAAAAAAGAACCATTT
 4 AGENOME-ZPMRG-F-V1.0   99.9%  99.8%       AAGTTTATATTCTTATCCTACCAGGCTTCGGAATAATCTCCCATATTGTAACTTACTACTCCGGAAAAAAAGAACCATTT
 5 AGENOME-ZPMRG-Z-V1.0   99.9%  99.7%       AAGTTTATATTCTTATCCTACCAGGCTTCGGAATAATCTCCCATATTGTAACTTACTACTCCGGAAAAAAAGAACCATTT
 6 AGENOME-ZPMRG-T-V1.0   99.9%  99.8%       AAGTTTATATTCTTATCCTACCAGGCTTCGGAATAATCTCCCATATTGTAACTTACTACTCCGGAAAAAAAGAACCATTT
 7 AGENOME-ZPMRG-HV-V1.0  99.9%  99.8%       AAGTTTATATTCTTATCCTACCAGGCTTCGGAATAATCTCCCATATTGTAACTTACTACTCCGGAAAAAAAGAACCATTT
 8 AGENOME-ZPMRG-A-V1.0   99.9%  99.8%       AAGTTTATATTCTTATCCTACCAGGCTTCGGAATAATCTCCCATATTGTAACTTACTACTCCGGAAAAAAAGAACCATTT
 9 NC_012920.1            99.9%  99.8%       AAGTTTATATTCTTATCCTACCAGGCTTCGGAATAATCTCCCATATTGTAACTTACTACTCCGGAAAAAAAGAACCATTT
   consensus/100%                            AAGTTTATATTCTTATCCTACCAGGCTTCGGAATAATCTCCCATATTGTAACTTACTACTCCGGAAAAAAAGAACCATTT
   consensus/90%                             AAGTTTATATTCTTATCCTACCAGGCTTCGGAATAATCTCCCATATTGTAACTTACTACTCCGGAAAAAAAGAACCATTT
   consensus/80%                             AAGTTTATATTCTTATCCTACCAGGCTTCGGAATAATCTCCCATATTGTAACTTACTACTCCGGAAAAAAAGAACCATTT
   consensus/70%                             AAGTTTATATTCTTATCCTACCAGGCTTCGGAATAATCTCCCATATTGTAACTTACTACTCCGGAAAAAAAGAACCATTT

                             cov    pid  6721          .         .         :         .         .         .         .         8 6800
 1 AGENOME-ZPMRG-U-V1.0  100.0% 100.0%       GGATACATAGGTATGGTCTGAGCTATGATATCAATTGGCTTCCTAGGGTTTATCGTGTGAGCACACCATATATTTACAGT
 2 AGENOME-ZPMCG-V1.0    100.0%  99.9%       GGATACATAGGTATGGTCTGAGCTATGATATCAATTGGCTTCCTAGGGTTTATCGTGTGAGCACACCATATATTTACAGT
 3 AGENOME-ZPMRG-M-V1.0  100.0%  99.8%       GGATACATAGGTATGGTCTGAGCTATGATATCAATTGGCTTCCTAGGGTTTATCGTGTGAGCACACCATATATTTACAGT
 4 AGENOME-ZPMRG-F-V1.0   99.9%  99.8%       GGATACATAGGTATGGTCTGAGCTATGATATCAATTGGCTTCCTAGGGTTTATCGTGTGAGCACACCATATATTTACAGT
 5 AGENOME-ZPMRG-Z-V1.0   99.9%  99.7%       GGATACATAGGTATGGTCTGAGCTATGATATCAATTGGCTTCCTAGGGTTTATCGTGTGAGCACACCATATATTTACAGT
 6 AGENOME-ZPMRG-T-V1.0   99.9%  99.8%       GGATACATAGGTATGGTCTGAGCTATGATATCAATTGGCTTCCTAGGGTTTATCGTGTGAGCACACCATATATTTACAGT
 7 AGENOME-ZPMRG-HV-V1.0  99.9%  99.8%       GGATACATAGGTATGGTCTGAGCTATGATATCAATTGGCTTCCTAGGGTTTATCGTGTGAGCACACCATATATTTACAGT
```

```
  8 AGENOME-ZPMRG-A-V1.0   99.9%  99.8%       GGATACATAGGTATGGTCTGAGCTATGATATCAATTGGCTTCCTAGGGTTTATCGTGTGAGCACACCATATATTTACAGT
  9 NC_012920.1           99.9%  99.8%       GGATACATAGGTATGGTCTGAGCTATGATATCAATTGGCTTCCTAGGGTTTATCGTGTGAGCACACCATATATTTACAGT
    consensus/100%                           GGATACATAGGTATGGTCTGAGCTATGATATCAATTGGCTTCCTAGGGTTTATCGTGTGAGCACACCATATATTTACAGT
    consensus/90%                            GGATACATAGGTATGGTCTGAGCTATGATATCAATTGGCTTCCTAGGGTTTATCGTGTGAGCACACCATATATTTACAGT
    consensus/80%                            GGATACATAGGTATGGTCTGAGCTATGATATCAATTGGCTTCCTAGGGTTTATCGTGTGAGCACACCATATATTTACAGT
    consensus/70%                            GGATACATAGGTATGGTCTGAGCTATGATATCAATTGGCTTCCTAGGGTTTATCGTGTGAGCACACCATATATTTACAGT

                            cov    pid  6801          .         .         .         .         :         .         .         . 6880
  1 AGENOME-ZPMRG-U-V1.0  100.0% 100.0%       AGGAATAGACGTAGACACACGAGCATATTTCACCTCCGCTACCATAATCATCGCTATCCCCACCGGCGTCAAAGTATTTA
  2 AGENOME-ZPMCG-V1.0    100.0%  99.9%       AGGAATAGACGTAGACACACGAGCATATTTCACCTCCGCTACCATAATCATCGCTATCCCCACCGGCGTCAAAGTATTTA
  3 AGENOME-ZPMRG-M-V1.0  100.0%  99.8%       AGGAATAGACGTAGACACACGAGCATATTTCACCTCCGCTACCATAATCATCGCTATCCCCACCGGCGTCAAAGTATTTA
  4 AGENOME-ZPMRG-F-V1.0   99.9%  99.8%       AGGAATAGACGTAGACACACGAGCATATTTCACCTCCGCTACCATAATCATCGCTATCCCCACCGGCGTCAAAGTATTTA
  5 AGENOME-ZPMRG-Z-V1.0   99.9%  99.7%       AGGAATAGACGTAGACACACGAGCATATTTCACCTCCGCTACCATAATCATCGCTATCCCCACCGGCGTCAAAGTATTTA
  6 AGENOME-ZPMRG-T-V1.0   99.9%  99.8%       AGGAATAGACGTAGACACACGAGCATATTTCACCTCCGCTACCATAATCATCGCTATCCCCACCGGCGTCAAAGTATTTA
  7 AGENOME-ZPMRG-HV-V1.0  99.9%  99.8%       AGGAATAGACGTAGACACACGAGCATATTTCACCTCCGCTACCATAATCATCGCTATCCCCACCGGCGTCAAAGTATTTA
  8 AGENOME-ZPMRG-A-V1.0   99.9%  99.8%       AGGAATAGACGTAGACACACGAGCATATTTCACCTCCGCTACCATAATCATCGCTATCCCCACCGGCGTCAAAGTATTTA
  9 NC_012920.1           99.9%  99.8%       AGGAATAGACGTAGACACACGAGCATATTTCACCTCCGCTACCATAATCATCGCTATCCCCACCGGCGTCAAAGTATTTA
    consensus/100%                           AGGAATAGACGTAGACACACGAGCATATTTCACCTCCGCTACCATAATCATCGCTATCCCCACCGGCGTCAAAGTATTTA
    consensus/90%                            AGGAATAGACGTAGACACACGAGCATATTTCACCTCCGCTACCATAATCATCGCTATCCCCACCGGCGTCAAAGTATTTA
    consensus/80%                            AGGAATAGACGTAGACACACGAGCATATTTCACCTCCGCTACCATAATCATCGCTATCCCCACCGGCGTCAAAGTATTTA
    consensus/70%                            AGGAATAGACGTAGACACACGAGCATATTTCACCTCCGCTACCATAATCATCGCTATCCCCACCGGCGTCAAAGTATTTA

                            cov    pid  6881          .         9         .         .         .         .         :         . 6960
  1 AGENOME-ZPMRG-U-V1.0  100.0% 100.0%       GCTGACTCGCCACACTCCACGGAAGCAATATGAAATGATCTGCTGCAGTGCTCTGAGCCCTAGGATTCATCTTTCTTTTC
  2 AGENOME-ZPMCG-V1.0    100.0%  99.9%       GCTGACTCGCCACACTCCACGGAAGCAATATGAAATGATCTGCTGCAGTGCTCTGAGCCCTAGGATTCATCTTTCTTTTC
  3 AGENOME-ZPMRG-M-V1.0  100.0%  99.8%       GCTGACTCGCCACACTCCACGGAAGCAATATGAAATGATCTGCTGCAGTGCTCTGAGCCCTAGGATTCATCTTTCTTTTC
  4 AGENOME-ZPMRG-F-V1.0   99.9%  99.8%       GCTGACTCGCCACACTCCACGGAAGCAATATGAAATGATCTGCTGCAGTGCTCTGAGCCCTAGGATTCATCTTTCTTTTC
  5 AGENOME-ZPMRG-Z-V1.0   99.9%  99.7%       GCTGACTCGCCACACTCCACGGAAGCAATATGAAATGATCTGCTGCAGTGCTCTGAGCCCTAGGATTCATCTTTCTTTTC
  6 AGENOME-ZPMRG-T-V1.0   99.9%  99.8%       GCTGACTCGCCACACTCCACGGAAGCAATATGAAATGATCTGCTGCAGTGCTCTGAGCCCTAGGATTCATCTTTCTTTTC
  7 AGENOME-ZPMRG-HV-V1.0  99.9%  99.8%       GCTGACTCGCCACACTCCACGGAAGCAATATGAAATGATCTGCTGCAGTGCTCTGAGCCCTAGGATTCATCTTTCTTTTC
  8 AGENOME-ZPMRG-A-V1.0   99.9%  99.8%       GCTGACTCGCCACACTCCACGGAAGCAATATGAAATGATCTGCTGCAGTGCTCTGAGCCCTAGGATTCATCTTTCTTTTC
  9 NC_012920.1           99.9%  99.8%       GCTGACTCGCCACACTCCACGGAAGCAATATGAAATGATCTGCTGCAGTGCTCTGAGCCCTAGGATTCATCTTTCTTTTC
    consensus/100%                           GCTGACTCGCCACACTCCACGGAAGCAATATGAAATGATCTGCTGCAGTGCTCTGAGCCCTAGGATTCATCTTTCTTTTC
    consensus/90%                            GCTGACTCGCCACACTCCACGGAAGCAATATGAAATGATCTGCTGCAGTGCTCTGAGCCCTAGGATTCATCTTTCTTTTC
    consensus/80%                            GCTGACTCGCCACACTCCACGGAAGCAATATGAAATGATCTGCTGCAGTGCTCTGAGCCCTAGGATTCATCTTTCTTTTC
    consensus/70%                            GCTGACTCGCCACACTCCACGGAAGCAATATGAAATGATCTGCTGCAGTGCTCTGAGCCCTAGGATTCATCTTTCTTTTC

                            cov    pid  6961          .         .         .         0         .         .         .         . 7040
  1 AGENOME-ZPMRG-U-V1.0  100.0% 100.0%       ACCGTAGGTGGCCTGACTGGCATTGTATTAGCAAACTCATCACTAGACATCGTACTACACGACACGTACTACGTTGTAGC
  2 AGENOME-ZPMCG-V1.0    100.0%  99.9%       ACCGTAGGTGGCCTGACTGGCATTGTATTAGCAAACTCATCACTAGACATCGTACTACACGACACGTACTACGTTGTAGC
  3 AGENOME-ZPMRG-M-V1.0  100.0%  99.8%       ACCGTAGGTGGCCTGACTGGCATTGTATTAGCAAACTCATCACTAGACATCGTACTACACGACACGTACTACGTTGTAGC
  4 AGENOME-ZPMRG-F-V1.0   99.9%  99.8%       ACCGTAGGTGGCCTAACTGGCATTGTATTAGCAAACTCATCACTAGACATCGTACTACACGACACGTACTACGTTGTAGC
  5 AGENOME-ZPMRG-Z-V1.0   99.9%  99.7%       ACCGTAGGTGGCCTGACTGGCATTGTATTAGCAAACTCATCACTAGACATCGTACTACACGACACGTACTACGTTGTAGC
  6 AGENOME-ZPMRG-T-V1.0   99.9%  99.8%       ACCGTAGGTGGCCTGACTGGCATTGTATTAGCAAACTCATCACTAGACATCGTACTACACGACACGTACTACGTTGTAGC
  7 AGENOME-ZPMRG-HV-V1.0  99.9%  99.8%       ACCGTAGGTGGCCTGACTGGCATTGTATTAGCAAACTCATCACTAGACATCGTACTACACGACACGTACTACGTTGTAGC
  8 AGENOME-ZPMRG-A-V1.0   99.9%  99.8%       ACCGTAGGTGGCCTGACTGGCATTGTATTAGCAAACTCATCACTAGACATCGTACTACACGACACGTACTACGTTGTAGC
  9 NC_012920.1           99.9%  99.8%       ACCGTAGGTGGCCTGACTGGCATTGTATTAGCAAACTCATCACTAGACATCGTACTACACGACACGTACTACGTTGTAGC
    consensus/100%                           ACCGTAGGTGGCCTuACTGGCATTGTATTAGCAAACTCATCACTAGACATCGTACTACACGACACGTACTACGTTGTAGC
    consensus/90%                            ACCGTAGGTGGCCTuACTGGCATTGTATTAGCAAACTCATCACTAGACATCGTACTACACGACACGTACTACGTTGTAGC
    consensus/80%                            ACCGTAGGTGGCCTGACTGGCATTGTATTAGCAAACTCATCACTAGACATCGTACTACACGACACGTACTACGTTGTAGC
    consensus/70%                            ACCGTAGGTGGCCTGACTGGCATTGTATTAGCAAACTCATCACTAGACATCGTACTACACGACACGTACTACGTTGTAGC

                            cov    pid  7041          :         .         .         .         .         1         .         . 7120
  1 AGENOME-ZPMRG-U-V1.0  100.0% 100.0%       TCACTTCCACTATGTCCTATCAATAGGAGCTGTATTTGCCATCATAGGAGGCTTCATTCACTGATTTCCCCTATTCTCAG
  2 AGENOME-ZPMCG-V1.0    100.0%  99.9%       TCACTTCCACTATGTCCTATCAATAGGAGCTGTATTTGCCATCATAGGAGGCTTCATTCACTGATTTCCCCTATTCTCAG
  3 AGENOME-ZPMRG-M-V1.0  100.0%  99.8%       TCACTTCCACTATGTCCTATCAATAGGAGCTGTATTTGCCATCATAGGAGGCTTCATTCACTGATTTCCCCTATTCTCAG
  4 AGENOME-ZPMRG-F-V1.0   99.9%  99.8%       TCACTTCCACTATGTCCTATCAATAGGAGCTGTATTTGCCATCATAGGAGGCTTCATTCACTGATTTCCCCTATTCTCAG
  5 AGENOME-ZPMRG-Z-V1.0   99.9%  99.7%       TCACTTCCACTATGTCCTATCAATAGGAGCTGTATTTGCCATCATAGGAGGCTTCATTCACTGATTTCCCCTATTCTCAG
  6 AGENOME-ZPMRG-T-V1.0   99.9%  99.8%       TCACTTCCACTATGTCCTATCAATAGGAGCTGTATTTGCCATCATAGGAGGCTTCATTCACTGATTTCCCCTATTCTCAG
  7 AGENOME-ZPMRG-HV-V1.0  99.9%  99.8%       TCACTTCCACTATGTCCTATCAATAGGAGCTGTATTTGCCATCATAGGAGGCTTCATTCACTGATTTCCCCTATTCTCAG
  8 AGENOME-ZPMRG-A-V1.0   99.9%  99.8%       CCACTTCCACTATGTCCTATCAATAGGAGCTGTATTTGCCATCATAGGAGGCTTCATTCACTGATTTCCCCTATTCTCAG
  9 NC_012920.1           99.9%  99.8%       CCACTTCCACTATGTCCTATCAATAGGAGCTGTATTTGCCATCATAGGAGGCTTCATTCACTGATTTCCCCTATTCTCAG
    consensus/100%                           sCACTTCCACTATGTCCTATCAATAGGAGCTGTATTTGCCATCATAGGAGGCTTCATTCACTGATTTCCCCTATTCTCAG
    consensus/90%                            sCACTTCCACTATGTCCTATCAATAGGAGCTGTATTTGCCATCATAGGAGGCTTCATTCACTGATTTCCCCTATTCTCAG
    consensus/80%                            sCACTTCCACTATGTCCTATCAATAGGAGCTGTATTTGCCATCATAGGAGGCTTCATTCACTGATTTCCCCTATTCTCAG
    consensus/70%                            TCACTTCCACTATGTCCTATCAATAGGAGCTGTATTTGCCATCATAGGAGGCTTCATTCACTGATTTCCCCTATTCTCAG

                            cov    pid  7121          .         .         :         .         .         .         .         2 7200
  1 AGENOME-ZPMRG-U-V1.0  100.0% 100.0%       GCTACACCCTAGACCAAACCTACGCCAAAATCCATTTCACTATCATATTCATCGGCGTAAATCTAACTTTCTTCCCACAA
  2 AGENOME-ZPMCG-V1.0    100.0%  99.9%       GCTACACCCTAGACCAAACCTACGCCAAAATCCATTTCACTATCATATTCATCGGCGTAAATCTAACTTTCTTCCCACAA
  3 AGENOME-ZPMRG-M-V1.0  100.0%  99.8%       GCTACACCCTAGACCAAACCTACGCCAAAATCCATTTCACTATCATATTCATCGGCGTAAATCTAACTTTCTTCCCACAA
  4 AGENOME-ZPMRG-F-V1.0   99.9%  99.8%       GCTACACCCTAGACCAAACCTACGCCAAAATCCATTTCACTATCATATTCATCGGCGTAAATCTAACTTTCTTCCCACAA
  5 AGENOME-ZPMRG-Z-V1.0   99.9%  99.7%       GCTACACCCTAGACCAAACCTACGCCAAAATCCATTTCACTATCATATTCATCGGCGTAAATCTAACTTTCTTCCCACAA
  6 AGENOME-ZPMRG-T-V1.0   99.9%  99.8%       GCTACACCCTAGACCAAACCTACGCCAAAATCCATTTCACTATCATATTCATCGGCGTAAATCTAACTTTCTTCCCACAA
  7 AGENOME-ZPMRG-HV-V1.0  99.9%  99.8%       GCTACACCCTAGACCAAACCTACGCCAAAATCCATTTCACTATCATATTCATCGGCGTAAATCTAACTTTCTTCCCACAA
  8 AGENOME-ZPMRG-A-V1.0   99.9%  99.8%       GCTACACCCTAGACCAAACCTACGCCAAAATCCATTTCACTATCATATTCATCGGCGTAAATCTAACTTTCTTCCCACAA
  9 NC_012920.1           99.9%  99.8%       GCTACACCCTAGACCAAACCTACGCCAAAATCCATTTCACTATCATATTCATCGGCGTAAATCTAACTTTCTTCCCACAA
    consensus/100%                           GCTACACCCTAGACCAAACCTACGCCAAAATCCATTTCACTATCATATTCATCGGCGTAAATCTAACTTTCTTCCCACAA
    consensus/90%                            GCTACACCCTAGACCAAACCTACGCCAAAATCCATTTCACTATCATATTCATCGGCGTAAATCTAACTTTCTTCCCACAA
    consensus/80%                            GCTACACCCTAGACCAAACCTACGCCAAAATCCATTTCACTATCATATTCATCGGCGTAAATCTAACTTTCTTCCCACAA
    consensus/70%                            GCTACACCCTAGACCAAACCTACGCCAAAATCCATTTCACTATCATATTCATCGGCGTAAATCTAACTTTCTTCCCACAA

                            cov    pid  7201          .         .         .         .         :         .         .         . 7280
  1 AGENOME-ZPMRG-U-V1.0  100.0% 100.0%       CACTTTCTCGGCCTATCCGGAATGCCCCGACGTTACTCGGACTACCCCGATGCATACACCACATGAAACATCCTATCATC
  2 AGENOME-ZPMCG-V1.0    100.0%  99.9%       CACTTTCTCGGCCTATCCGGAATGCCCCGACGTTACTCGGACTACCCCGATGCATACACCACATGAAACATCCTATCATC
  3 AGENOME-ZPMRG-M-V1.0  100.0%  99.8%       CACTTTCTCGGCCTATCCGGAATGCCCCGACGTTACTCGGACTACCCCGATGCATACACCACATGAAACATCCTATCATC
  4 AGENOME-ZPMRG-F-V1.0   99.9%  99.8%       CACTTTCTCGGCCTATCCGGAATGCCCCGACGTTACTCGGACTACCCCGATGCATACACCACATGAAACATCCTATCATC
  5 AGENOME-ZPMRG-Z-V1.0   99.9%  99.7%       CACTTTCTCGGCCTATCCGGAATGCCCCGACGTTACTCGGACTACCCCGATGCATACACCACATGAAACATCCTATCATC
  6 AGENOME-ZPMRG-T-V1.0   99.9%  99.8%       CACTTTCTCGGCCTATCCGGAATGCCCCGACGTTACTCGGACTACCCCGATGCATACACCACATGAAACATCCTATCATC
  7 AGENOME-ZPMRG-HV-V1.0  99.9%  99.8%       CACTTCCTCGGCCTATCCGGAATGCCCCGACGTTACTCGGACTACCCCGATGCATACACCACATGAAACATCCTATCATC
  8 AGENOME-ZPMRG-A-V1.0   99.9%  99.8%       CACTTTCTCGGCCTATCCGGAATGCCCCGACGTTACTCGGACTACCCCGATGCATACACCACATGAAACATCCTATCATC
```

```
  9 NC_012920.1             99.9%  99.8%       CACTTTCTCGGCCTATCCGGAATGCCCCGACGTTACTCGGACTACCCCGATGCATACACCACATGAAACATCCTATCATC
    consensus/100%                              CACTTsCTCGGCCTATCCGGAATGCCCCGACGTTACTCGGACTACCCCGATGCATACACCACATGAAACATCCTATCATC
    consensus/90%                               CACTTsCTCGGCCTATCCGGAATGCCCCGACGTTACTCGGACTACCCCGATGCATACACCACATGAAACATCCTATCATC
    consensus/80%                               CACTTTCTCGGCCTATCCGGAATGCCCCGACGTTACTCGGACTACCCCGATGCATACACCACATGAAACATCCTATCATC
    consensus/70%                               CACTTTCTCGGCCTATCCGGAATGCCCCGACGTTACTCGGACTACCCCGATGCATACACCACATGAAACATCCTATCATC

                              cov    pid   7281          .         3         .         .         .         .         :         . 7360
  1 AGENOME-ZPMRG-U-V1.0    100.0% 100.0%       TGTAGGCTCATTCATTTCTCTAACAGCAGTAATATTAATAATTTTCATGATTTGAGAAGCCTTCGCTTCGAAGCGAAAAG
  2 AGENOME-ZPMCG-V1.0      100.0%  99.9%       TGTAGGCTCATTCATTTCTCTAACAGCAGTAATATTAATAATTTTCATGATTTGAGAAGCCTTCGCTTCGAAGCGAAAAG
  3 AGENOME-ZPMRG-M-V1.0    100.0%  99.8%       TGTAGGCTCATTCATTTCTCTAACAGCAGTAATATTAATAATTTTCATGATTTGAGAAGCCTTCGCTTCGAAGCGAAAAG
  4 AGENOME-ZPMRG-F-V1.0     99.9%  99.8%       TGTAGGCTCATTCATTTCTCTAACAGCAGTAATATTAATAATTTTCATGATTTGAGAAGCCTTCGCTTCGAAGCGAAAAG
  5 AGENOME-ZPMRG-Z-V1.0     99.9%  99.7%       TGTAGGCTCATTCATTTCTCTAACAGCAGTAATATTAATAATTTTCATGATTTGAGAAGCCTTCGCTTCGAAGCGAAAAG
  6 AGENOME-ZPMRG-T-V1.0     99.9%  99.8%       TGTAGGCTCATTCATTTCTCTAACAGCAGTAATATTAATAATTTTCATGATTTGAGAAGCCTTCGCTTCGAAGCGAAAAG
  7 AGENOME-ZPMRG-HV-V1.0    99.9%  99.8%       TGTAGGCTCATTCATTTCTCTAACAGCAGTAATATTAATAATTTTCATGATTTGAGAAGCCTTCGCTTCGAAGCGAAAAG
  8 AGENOME-ZPMRG-A-V1.0     99.9%  99.8%       TGTAGGCTCATTCATTTCTCTAACAGCAGTAATATTAATAATTTTCATGATTTGAGAAGCCTTCGCTTCGAAGCGAAAAG
  9 NC_012920.1             99.9%  99.8%       TGTAGGCTCATTCATTTCTCTAACAGCAGTAATATTAATAATTTTCATGATTTGAGAAGCCTTCGCTTCGAAGCGAAAAG
    consensus/100%                              TGTAGGCTCATTCATTTCTCTAACAGCAGTAATATTAATAATTTTCATGATTTGAGAAGCCTTCGCTTCGAAGCGAAAAG
    consensus/90%                               TGTAGGCTCATTCATTTCTCTAACAGCAGTAATATTAATAATTTTCATGATTTGAGAAGCCTTCGCTTCGAAGCGAAAAG
    consensus/80%                               TGTAGGCTCATTCATTTCTCTAACAGCAGTAATATTAATAATTTTCATGATTTGAGAAGCCTTCGCTTCGAAGCGAAAAG
    consensus/70%                               TGTAGGCTCATTCATTTCTCTAACAGCAGTAATATTAATAATTTTCATGATTTGAGAAGCCTTCGCTTCGAAGCGAAAAG

                              cov    pid   7361          .         .         .         4         .         .         .         . 7440
  1 AGENOME-ZPMRG-U-V1.0    100.0% 100.0%       TCCTAATAGTAGAAGAACCCTCCATAAACCTGGAGTGACTATATGGATGCCCCCCACCCTACCACACATTCGAAGAACCC
  2 AGENOME-ZPMCG-V1.0      100.0%  99.9%       TCCTAATAGTAGAAGAACCCTCCATAAACCTGGAGTGACTATATGGATGCCCCCCACCCTACCACACATTCGAAGAACCC
  3 AGENOME-ZPMRG-M-V1.0    100.0%  99.8%       TCCTAATAGTAGAAGAACCCTCCATAAACCTGGAGTGACTATATGGATGCCCCCCACCCTACCACACATTCGAAGAACCC
  4 AGENOME-ZPMRG-F-V1.0     99.9%  99.8%       TCCTAATAGTAGAAGAACCCTCCATAAACCTGGAGTGACTATATGGATGCCCCCCACCCTACCACACATTCGAAGAACCC
  5 AGENOME-ZPMRG-Z-V1.0     99.9%  99.7%       TCCTAATAGTAGAAGAACCCTCCATAAACCTGGAGTGACTATATGGATGCCCCCCACCCTACCACACATTCGAAGAACCC
  6 AGENOME-ZPMRG-T-V1.0     99.9%  99.8%       TCCTAATAGTAGAAGAACCCTCCATAAACCTGGAGTGACTATATGGATGCCCCCCACCCTACCACACATTCGAAGAACCC
  7 AGENOME-ZPMRG-HV-V1.0    99.9%  99.8%       TCCTAATAGTAGAAGAACCCTCCATAAACCTGGAGTGACTATATGGATGCCCCCCACCCTACCACACATTCGAAGAACCC
  8 AGENOME-ZPMRG-A-V1.0     99.9%  99.8%       TCCTAATAGTAGAAGAACCCTCCATAAACCTGGAGTGACTATATGGATGCCCCCCACCCTACCACACATTCGAAGAACCC
  9 NC_012920.1             99.9%  99.8%       TCCTAATAGTAGAAGAACCCTCCATAAACCTGGAGTGACTATATGGATGCCCCCCACCCTACCACACATTCGAAGAACCC
    consensus/100%                              TCCTAATAGTAGAAGAACCCTCCATAAACCTGGAGTGACTATATGGATGCCCCCCACCCTACCACACATTCGAAGAACCC
    consensus/90%                               TCCTAATAGTAGAAGAACCCTCCATAAACCTGGAGTGACTATATGGATGCCCCCCACCCTACCACACATTCGAAGAACCC
    consensus/80%                               TCCTAATAGTAGAAGAACCCTCCATAAACCTGGAGTGACTATATGGATGCCCCCCACCCTACCACACATTCGAAGAACCC
    consensus/70%                               TCCTAATAGTAGAAGAACCCTCCATAAACCTGGAGTGACTATATGGATGCCCCCCACCCTACCACACATTCGAAGAACCC

                              cov    pid   7441          :         .         .         .         .         5         .         . 7520
  1 AGENOME-ZPMRG-U-V1.0    100.0% 100.0%       GTATACATAAAATCTAGACAAAAAAGGAAGGAATCGAACCCCCCAAAGCTGGTTTCAAGCCAACCCCATGGCCTCCATGA
  2 AGENOME-ZPMCG-V1.0      100.0%  99.9%       GTATACATAAAATCTAGACAAAAAAGGAAGGAATCGAACCCCCCAAAGCTGGTTTCAAGCCAACCCCATGGCCTCCATGA
  3 AGENOME-ZPMRG-M-V1.0    100.0%  99.8%       GTATACATAAAATCTAGACAAAAAAGGAAGGAATCGAACCCCCCAAAGCTGGTTTCAAGCCAACCCCATGGCCTCCATGA
  4 AGENOME-ZPMRG-F-V1.0     99.9%  99.8%       GTATACATAAAATCTAGACAAAAAAGGAAGGAATCGAACCCCCCAAAGCTGGTTTCAAGCCAACCCCATGGCCTCCATGA
  5 AGENOME-ZPMRG-Z-V1.0     99.9%  99.7%       GTATACATAAAATCTAGACAAAAAAGGAAGGAATCGAACCCCCCAAAGCTGGTTTCAAGCCAACCCCATGGCCTCCATGA
  6 AGENOME-ZPMRG-T-V1.0     99.9%  99.8%       GTATACATAAAATCTAGACAAAAAAGGAAGGAATCGAACCCCCCAAAGCTGGTTTCAAGCCAACCCCATGGCCTCCATGA
  7 AGENOME-ZPMRG-HV-V1.0    99.9%  99.8%       GTATACATAAAATCTAGACAAAAAAGGAAGGAATCGAACCCCCCAAAGCTGGTTTCAAGCCAACCCCATGGCCTCCATGA
  8 AGENOME-ZPMRG-A-V1.0     99.9%  99.8%       GTATACATAAAATCTAGACAAAAAAGGAAGGAATCGAACCCCCCAAAGCTGGTTTCAAGCCAACCCCATGGCCTCCATGA
  9 NC_012920.1             99.9%  99.8%       GTATACATAAAATCTAGACAAAAAAGGAAGGAATCGAACCCCCCAAAGCTGGTTTCAAGCCAACCCCATGGCCTCCATGA
    consensus/100%                              GTATACATAAAATCTAGACAAAAAAGGAAGGAATCGAACCCCCCAAAGCTGGTTTCAAGCCAACCCCATGGCCTCCATGA
    consensus/90%                               GTATACATAAAATCTAGACAAAAAAGGAAGGAATCGAACCCCCCAAAGCTGGTTTCAAGCCAACCCCATGGCCTCCATGA
    consensus/80%                               GTATACATAAAATCTAGACAAAAAAGGAAGGAATCGAACCCCCCAAAGCTGGTTTCAAGCCAACCCCATGGCCTCCATGA
    consensus/70%                               GTATACATAAAATCTAGACAAAAAAGGAAGGAATCGAACCCCCCAAAGCTGGTTTCAAGCCAACCCCATGGCCTCCATGA

                              cov    pid   7521          .         .         :         .         .         .         .         6 7600
  1 AGENOME-ZPMRG-U-V1.0    100.0% 100.0%       CTTTTTCAAAAAGGTATTAGAAAAACCATTTCATAACTTTGTCAAAGTTAAATTATAGGCTAAATCCTATATATCTTAAT
  2 AGENOME-ZPMCG-V1.0      100.0%  99.9%       CTTTTTCAAAAAGGTATTAGAAAAACCATTTCATAACTTTGTCAAAGTTAAATTATAGGCTAAATCCTATATATCTTAAT
  3 AGENOME-ZPMRG-M-V1.0    100.0%  99.8%       CTTTTTCAAAAAGGTATTAGAAAAACCATTTCATAACTTTGTCAAAGTTAAATTATAGGCTAAATCCTATATATCTTAAT
  4 AGENOME-ZPMRG-F-V1.0     99.9%  99.8%       CTTTTTCAAAAAGGTATTAGAAAAACCATTTCATAACTTTGTCAAAGTTAAATTATAGGCTAAATCCTATATATCTTAAT
  5 AGENOME-ZPMRG-Z-V1.0     99.9%  99.7%       CTTTTTCAAAAAGGTATTAGAAAAACCATTTCATAACTTTGTCAAAGTTAAATTATAGGCTAAATCCTATATATCTTAAT
  6 AGENOME-ZPMRG-T-V1.0     99.9%  99.8%       CTTTTTCAAAAAGGTATTAGAAAAACCATTTCATAACTTTGTCAAAGTTAAATTATAGGCTAAATCCTATATATCTTAAT
  7 AGENOME-ZPMRG-HV-V1.0    99.9%  99.8%       CTTTTTCAAAAAGGTATTAGAAAAACCATTTCATAACTTTGTCAAAGTTAAATTATAGGCTAAATCCTATATATCTTAAT
  8 AGENOME-ZPMRG-A-V1.0     99.9%  99.8%       CTTTTTCAAAAAGGTATTAGAAAAACCATTTCATAACTTTGTCAAAGTTAAATTATAGGCTAAATCCTATATATCTTAAT
  9 NC_012920.1             99.9%  99.8%       CTTTTTCAAAAAGGTATTAGAAAAACCATTTCATAACTTTGTCAAAGTTAAATTATAGGCTAAATCCTATATATCTTAAT
    consensus/100%                              CTTTTTCAAAAAGGTATTAGAAAAACCATTTCATAACTTTGTCAAAGTTAAATTATAGGCTAAATCCTATATATCTTAAT
    consensus/90%                               CTTTTTCAAAAAGGTATTAGAAAAACCATTTCATAACTTTGTCAAAGTTAAATTATAGGCTAAATCCTATATATCTTAAT
    consensus/80%                               CTTTTTCAAAAAGGTATTAGAAAAACCATTTCATAACTTTGTCAAAGTTAAATTATAGGCTAAATCCTATATATCTTAAT
    consensus/70%                               CTTTTTCAAAAAGGTATTAGAAAAACCATTTCATAACTTTGTCAAAGTTAAATTATAGGCTAAATCCTATATATCTTAAT

                              cov    pid   7601          .         .         .         .         :         .         .         . 7680
  1 AGENOME-ZPMRG-U-V1.0    100.0% 100.0%       GGCACATGCAGCGCAAGTAGGTCTACAAGACGCTACTTCCCCTATCATAGAAGAGCTTATCACCTTTCATGATCACGCCC
  2 AGENOME-ZPMCG-V1.0      100.0%  99.9%       GGCACATGCAGCGCAAGTAGGTCTACAAGACGCTACTTCCCCTATCATAGAAGAGCTTATCACCTTTCATGATCACGCCC
  3 AGENOME-ZPMRG-M-V1.0    100.0%  99.8%       GGCACATGCAGCGCAAGTAGGTCTACAAGACGCTACTTCCCCTATCATAGAAGAGCTTATCACCTTTCATGATCACGCCC
  4 AGENOME-ZPMRG-F-V1.0     99.9%  99.8%       GGCACATGCAGCGCAAGTAGGTCTACAAGACGCTACTTCCCCTATCATAGAAGAGCTTATCACCTTTCATGATCACGCCC
  5 AGENOME-ZPMRG-Z-V1.0     99.9%  99.7%       GGCACATGCAGCGCAAGTAGGTCTACAAGACGCTACTTCCCCTATCATAGAAGAGCTTATCACCTTTCATGATCACGCCC
  6 AGENOME-ZPMRG-T-V1.0     99.9%  99.8%       GGCACATGCAGCGCAAGTAGGTCTACAAGACGCTACTTCCCCTATCATAGAAGAGCTTATCACCTTTCATGATCACGCCC
  7 AGENOME-ZPMRG-HV-V1.0    99.9%  99.8%       GGCACATGCAGCGCAAGTAGGTCTACAAGACGCTACTTCCCCTATCATAGAAGAGCTTATCACCTTTCATGATCACGCCC
  8 AGENOME-ZPMRG-A-V1.0     99.9%  99.8%       GGCACATGCAGCGCAAGTAGGTCTACAAGACGCTACTTCCCCTATCATAGAAGAGCTTATCACCTTTCATGATCACGCCC
  9 NC_012920.1             99.9%  99.8%       GGCACATGCAGCGCAAGTAGGTCTACAAGACGCTACTTCCCCTATCATAGAAGAGCTTATCACCTTTCATGATCACGCCC
    consensus/100%                              GGCACATGCAGCGCAAGTAGGTCTACAAGACGCTACTTCCCCTATCATAGAAGAGCTTATCACCTTTCATGATCACGCCC
    consensus/90%                               GGCACATGCAGCGCAAGTAGGTCTACAAGACGCTACTTCCCCTATCATAGAAGAGCTTATCACCTTTCATGATCACGCCC
    consensus/80%                               GGCACATGCAGCGCAAGTAGGTCTACAAGACGCTACTTCCCCTATCATAGAAGAGCTTATCACCTTTCATGATCACGCCC
    consensus/70%                               GGCACATGCAGCGCAAGTAGGTCTACAAGACGCTACTTCCCCTATCATAGAAGAGCTTATCACCTTTCATGATCACGCCC

                              cov    pid   7681          .         7         .         .         .         .         :         . 7760
  1 AGENOME-ZPMRG-U-V1.0    100.0% 100.0%       TCATAATCATTTTCCTTATCTGCTTCCTAGTCCTGTATGCCCTTTTCCTAACACTCACAACAAAACTAACTAATACTAAC
  2 AGENOME-ZPMCG-V1.0      100.0%  99.9%       TCATAATCATTTTCCTTATCTGCTTCCTAGTCCTGTATGCCCTTTTCCTAACACTCACAACAAAACTAACTAATACTAAC
  3 AGENOME-ZPMRG-M-V1.0    100.0%  99.8%       TCATAATCATTTTCCTTATCTGCTTCCTAGTCCTGTATGCCCTTTTCCTAACACTCACAACAAAACTAACTAATACTAAC
  4 AGENOME-ZPMRG-F-V1.0     99.9%  99.8%       TCATAATCATTTTCCTTATCTGCTTCCTAGTCCTGTATGCCCTTTTCCTAACACTCACAACAAAACTAACTAATACTAAC
  5 AGENOME-ZPMRG-Z-V1.0     99.9%  99.7%       TCATAATCATTTTCCTTATCTGCTTCCTAGTCCTGTATGCCCTTTTCCTAACACTCACAACAAAACTAACTAATACTAAC
  6 AGENOME-ZPMRG-T-V1.0     99.9%  99.8%       TCATAATCATTTTCCTTATCTGCTTCCTAGTCCTGTATGCCCTTTTCCTAACACTCACAACAAAACTAACTAATACTAAC
  7 AGENOME-ZPMRG-HV-V1.0    99.9%  99.8%       TCATAATCATTTTCCTTATCTGCTTCCTAGTCCTGTATGCCCTTTTCCTAACACTCACAACAAAACTAACTAATACTAAC
  8 AGENOME-ZPMRG-A-V1.0     99.9%  99.8%       TCATAATCATTTTCCTTATCTGCTTCCTAGTCCTGTATGCCCTTTTCCTAACACTCACAACAAAACTAACTAATACTAAC
  9 NC_012920.1             99.9%  99.8%       TCATAATCATTTTCCTTATCTGCTTCCTAGTCCTGTATGCCCTTTTCCTAACACTCACAACAAAACTAACTAATACTAAC
```

```
   consensus/100%                          TCATAATCATTTTCCTTATCTGCTTCCTAGTCCTGTATGCCCTTTTCCTAACACTCACAACAAAACTAACTAATACTAAC
   consensus/90%                           TCATAATCATTTTCCTTATCTGCTTCCTAGTCCTGTATGCCCTTTTCCTAACACTCACAACAAAACTAACTAATACTAAC
   consensus/80%                           TCATAATCATTTTCCTTATCTGCTTCCTAGTCCTGTATGCCCTTTTCCTAACACTCACAACAAAACTAACTAATACTAAC
   consensus/70%                           TCATAATCATTTTCCTTATCTGCTTCCTAGTCCTGTATGCCCTTTTCCTAACACTCACAACAAAACTAACTAATACTAAC

                             cov    pid  7761          .         .         .         8         .         .         .         . 7840
 1 AGENOME-ZPMRG-U-V1.0  100.0% 100.0%       ATCTCAGACGCTCAGGAAATAGAAACCGTCTGAACTATCCTGCCCGCCATCATCCTAGTCCTCATCGCCCTCCCATCCCT
 2 AGENOME-ZPMCG-V1.0    100.0%  99.9%       ATCTCAGACGCTCAGGAAATAGAAACCGTCTGAACTATCCTGCCCGCCATCATCCTAGTCCTCATCGCCCTCCCATCCCT
 3 AGENOME-ZPMRG-M-V1.0  100.0%  99.8%       ATCTCAGACGCTCAGGAAATAGAAACCGTCTGAACTATCCTGCCCGCCATCATCCTAGTCCTCATCGCCCTCCCATCCCT
 4 AGENOME-ZPMRG-F-V1.0   99.9%  99.8%       ATCTCAGACGCTCAGGAAATAGAAACCGTCTGAACTATCCTGCCCGCCATCATCCTAGTCCTCATCGCCCTCCCATCCCT
 5 AGENOME-ZPMRG-Z-V1.0   99.9%  99.7%       ATCTCAGACGCTCAGGAAATAGAAACCGTCTGAACTATCCTGCCCGCCATCATCCTAGTCCTCATCGCCCTCCCATCCCT
 6 AGENOME-ZPMRG-T-V1.0   99.9%  99.8%       ATCTCAGACGCTCAGGAAATAGAAACCGTCTGAACTATCCTGCCCGCCATCATCCTAGTCCTCATCGCCCTCCCATCCCT
 7 AGENOME-ZPMRG-HV-V1.0  99.9%  99.8%       ATCTCAGACGCTCAGGAAATAGAAACCGTCTGAACTATCCTGCCCGCCATCATCCTAGTCCTCATCGCCCTCCCATCCCT
 8 AGENOME-ZPMRG-A-V1.0   99.9%  99.8%       ATCTCAGACGCTCAGGAAATAGAAACCGTCTGAACTATCCTGCCCGCCATCATCCTAGTCCTCATCGCCCTCCCATCCCT
 9 NC_012920.1            99.9%  99.8%       ATCTCAGACGCTCAGGAAATAGAAACCGTCTGAACTATCCTGCCCGCCATCATCCTAGTCCTCATCGCCCTCCCATCCCT
   consensus/100%                            ATCTCAGACGCTCAGGAAATAGAAACCGTCTGAACTATCCTGCCCGCCATCATCCTAGTCCTCATCGCCCTCCCATCCCT
   consensus/90%                             ATCTCAGACGCTCAGGAAATAGAAACCGTCTGAACTATCCTGCCCGCCATCATCCTAGTCCTCATCGCCCTCCCATCCCT
   consensus/80%                             ATCTCAGACGCTCAGGAAATAGAAACCGTCTGAACTATCCTGCCCGCCATCATCCTAGTCCTCATCGCCCTCCCATCCCT
   consensus/70%                             ATCTCAGACGCTCAGGAAATAGAAACCGTCTGAACTATCCTGCCCGCCATCATCCTAGTCCTCATCGCCCTCCCATCCCT

                             cov    pid  7841          :         .         .         .         .         9         .         . 7920
 1 AGENOME-ZPMRG-U-V1.0  100.0% 100.0%       ACGCATCCTTTACATAACAGACGAGGTCAACGATCCCTCCCTTACCATCAAATCAATTGGCCACCAATGGTACTGAACCT
 2 AGENOME-ZPMCG-V1.0    100.0%  99.9%       ACGCATCCTTTACATAACAGACGAGGTCAACGATCCCTCCCTTACCATCAAATCAATTGGCCACCAATGGTACTGAACCT
 3 AGENOME-ZPMRG-M-V1.0  100.0%  99.8%       ACGCATCCTTTACATAACAGACGAGGTCAACGATCCCTCCCTTACCATCAAATCAATTGGCCACCAATGGTACTGAACCT
 4 AGENOME-ZPMRG-F-V1.0   99.9%  99.8%       ACGCATCCTTTACATAACAGACGAGGTCAACGATCCCTCCCTTACCATCAAATCAATTGGCCACCAATGGTACTGAACCT
 5 AGENOME-ZPMRG-Z-V1.0   99.9%  99.7%       ACGCATCCTTTACATAACAGACGAGGTCAACGATCCCTCCCTTACCATCAAATCAATTGGCCACCAATGGTACTGAACCT
 6 AGENOME-ZPMRG-T-V1.0   99.9%  99.8%       ACGCATCCTTTACATAACAGACGAGGTCAACGATCCCTCCCTTACCATCAAATCAATTGGCCACCAATGGTACTGAACCT
 7 AGENOME-ZPMRG-HV-V1.0  99.9%  99.8%       ACGCATCCTTTACATAACAGACGAGGTCAACGATCCCTCCCTTACCATCAAATCAATTGGCCACCAATGGTACTGAACCT
 8 AGENOME-ZPMRG-A-V1.0   99.9%  99.8%       ACGCATCCTTTACATAACAGACGAGGTCAACGATCCCTCCCTTACCATCAAATCAATTGGCCACCAATGGTACTGAACCT
 9 NC_012920.1            99.9%  99.8%       ACGCATCCTTTACATAACAGACGAGGTCAACGATCCCTCCCTTACCATCAAATCAATTGGCCACCAATGGTACTGAACCT
   consensus/100%                            ACGCATCCTTTACATAACAGACGAGGTCAACGATCCCTCCCTTACCATCAAATCAATTGGCCACCAATGGTACTGAACCT
   consensus/90%                             ACGCATCCTTTACATAACAGACGAGGTCAACGATCCCTCCCTTACCATCAAATCAATTGGCCACCAATGGTACTGAACCT
   consensus/80%                             ACGCATCCTTTACATAACAGACGAGGTCAACGATCCCTCCCTTACCATCAAATCAATTGGCCACCAATGGTACTGAACCT
   consensus/70%                             ACGCATCCTTTACATAACAGACGAGGTCAACGATCCCTCCCTTACCATCAAATCAATTGGCCACCAATGGTACTGAACCT

                             cov    pid  7921          .         .         :         .         .         .         .         0 8000
 1 AGENOME-ZPMRG-U-V1.0  100.0% 100.0%       ACGAGTACACCGACTACGGCGGACTAATCTTCAACTCCTACATACTTCCCCCATTATTCCTAGAACCAGGCGACCTGCGA
 2 AGENOME-ZPMCG-V1.0    100.0%  99.9%       ACGAGTACACCGACTACGGCGGACTAATCTTCAACTCCTACATACTTCCCCCATTATTCCTAGAACCAGGCGACCTGCGA
 3 AGENOME-ZPMRG-M-V1.0  100.0%  99.8%       ACGAGTACACCGACTACGGCGGACTAATCTTCAACTCCTACATACTTCCCCCATTATTCCTAGAACCAGGCGACCTGCGA
 4 AGENOME-ZPMRG-F-V1.0   99.9%  99.8%       ACGAGTACACCGACTACGGCGGACTAATCTTCAACTCCTACATACTTCCCCCATTATTCCTAGAACCAGGCGACCTGCGA
 5 AGENOME-ZPMRG-Z-V1.0   99.9%  99.7%       ACGAGTACACCGACTACGGCGGACTAATCTTCAACTCCTACATACTTCCCCCATTATTCCTAGAACCAGGCGACCTGCGA
 6 AGENOME-ZPMRG-T-V1.0   99.9%  99.8%       ACGAGTACACCGACTACGGCGGACTAATCTTCAACTCCTACATACTTCCCCCATTATTCCTAGAACCAGGCGACCTGCGA
 7 AGENOME-ZPMRG-HV-V1.0  99.9%  99.8%       ACGAGTACACCGACTACGGCGGACTAATCTTCAACTCCTACATACTTCCCCCATTATTCCTAGAACCAGGCGACCTGCGA
 8 AGENOME-ZPMRG-A-V1.0   99.9%  99.8%       ACGAGTACACCGACTACGGCGGACTAATCTTCAACTCCTACATACTTCCCCCATTATTCCTAGAACCAGGCGACCTGCGA
 9 NC_012920.1            99.9%  99.8%       ACGAGTACACCGACTACGGCGGACTAATCTTCAACTCCTACATACTTCCCCCATTATTCCTAGAACCAGGCGACCTGCGA
   consensus/100%                            ACGAGTACACCGACTACGGCGGACTAATCTTCAACTCCTACATACTTCCCCCATTATTCCTAGAACCAGGCGACCTGCGA
   consensus/90%                             ACGAGTACACCGACTACGGCGGACTAATCTTCAACTCCTACATACTTCCCCCATTATTCCTAGAACCAGGCGACCTGCGA
   consensus/80%                             ACGAGTACACCGACTACGGCGGACTAATCTTCAACTCCTACATACTTCCCCCATTATTCCTAGAACCAGGCGACCTGCGA
   consensus/70%                             ACGAGTACACCGACTACGGCGGACTAATCTTCAACTCCTACATACTTCCCCCATTATTCCTAGAACCAGGCGACCTGCGA

                             cov    pid  8001          .         .         .         .         :         .         .         . 8080
 1 AGENOME-ZPMRG-U-V1.0  100.0% 100.0%       CTCCTTGACGTTGACAATCGAGTAGTACTCCCGATTGAAGCCCCCATTCGTATAATAATTACATCACAAGACGTCTTGCA
 2 AGENOME-ZPMCG-V1.0    100.0%  99.9%       CTCCTTGACGTTGACAATCGAGTAGTACTCCCGATTGAAGCCCCCATTCGTATAATAATTACATCACAAGACGTCTTGCA
 3 AGENOME-ZPMRG-M-V1.0  100.0%  99.8%       CTCCTTGACGTTGACAATCGAGTAGTACTCCCGATTGAAGCCCCCATTCGTATAATAATTACATCACAAGACGTCTTGCA
 4 AGENOME-ZPMRG-F-V1.0   99.9%  99.8%       CTCCTTGACGTTGACAATCGAGTAGTACTCCCGATTGAAGCCCCCATTCGTATAATAATTACATCACAAGACGTCTTGCA
 5 AGENOME-ZPMRG-Z-V1.0   99.9%  99.7%       CTCCTTGACGTTGACAATCGAGTAGTACTCCCGATTGAAGCCCCCATTCGTATAATAATTACATCACAAGACGTCTTGCA
 6 AGENOME-ZPMRG-T-V1.0   99.9%  99.8%       CTCCTTGACGTTGACAATCGAGTAGTACTCCCGATTGAAGCCCCCATTCGTATAATAATTACATCACAAGACGTCTTGCA
 7 AGENOME-ZPMRG-HV-V1.0  99.9%  99.8%       CTCCTTGACGTTGACAATCGAGTAGTACTCCCGATTGAAGCCCCCATTCGTATAATAATTACATCACAAGACGTCTTGCA
 8 AGENOME-ZPMRG-A-V1.0   99.9%  99.8%       CTCCTTGACGTTGACAATCGAGTAGTACTCCCGATTGAAGCCCCCATTCGTATAATAATTACATCACAAGACGTCTTGCA
 9 NC_012920.1            99.9%  99.8%       CTCCTTGACGTTGACAATCGAGTAGTACTCCCGATTGAAGCCCCCATTCGTATAATAATTACATCACAAGACGTCTTGCA
   consensus/100%                            CTCCTTGACGTTGACAATCGAGTAGTACTCCCGATTGAAGCCCCCATTCGTATAATAATTACATCACAAGACGTCTTGCA
   consensus/90%                             CTCCTTGACGTTGACAATCGAGTAGTACTCCCGATTGAAGCCCCCATTCGTATAATAATTACATCACAAGACGTCTTGCA
   consensus/80%                             CTCCTTGACGTTGACAATCGAGTAGTACTCCCGATTGAAGCCCCCATTCGTATAATAATTACATCACAAGACGTCTTGCA
   consensus/70%                             CTCCTTGACGTTGACAATCGAGTAGTACTCCCGATTGAAGCCCCCATTCGTATAATAATTACATCACAAGACGTCTTGCA

                             cov    pid  8081          .         1         .         .         .         .         :         . 8160
 1 AGENOME-ZPMRG-U-V1.0  100.0% 100.0%       CTCATGAGCTGTCCCCACATTAGGCTTAAAAACAGATGCAATTCCCGGACGTCTAAACCAAACCACTTTCACCGCTACAC
 2 AGENOME-ZPMCG-V1.0    100.0%  99.9%       CTCATGAGCTGTCCCCACATTAGGCTTAAAAACAGATGCAATTCCCGGACGTCTAAACCAAACCACTTTCACCGCTACAC
 3 AGENOME-ZPMRG-M-V1.0  100.0%  99.8%       CTCATGAGCTGTCCCCACATTAGGCTTAAAAACAGATGCAATTCCCGGACGTCTAAACCAAACCACTTTCACCGCTACAC
 4 AGENOME-ZPMRG-F-V1.0   99.9%  99.8%       CTCATGAGCTGTCCCCACATTAGGCTTAAAAACAGATGCAATTCCCGGACGTCTAAACCAAACCACTTTCACCGCTACAC
 5 AGENOME-ZPMRG-Z-V1.0   99.9%  99.7%       CTCATGAGCTGTCCCCACATTAGGCTTAAAAACAGATGCAATTCCCGGACGTCTAAACCAAACCACTTTCACCGCTACAC
 6 AGENOME-ZPMRG-T-V1.0   99.9%  99.8%       CTCATGAGCTGTCCCCACATTAGGCTTAAAAACAGATGCAATTCCCGGACGTCTAAACCAAACCACTTTCACCGCTACAC
 7 AGENOME-ZPMRG-HV-V1.0  99.9%  99.8%       CTCATGAGCTGTCCCCACATTAGGCTTAAAAACAGATGCAATTCCCGGACGTCTAAACCAAACCACTTTCACCGCTACAC
 8 AGENOME-ZPMRG-A-V1.0   99.9%  99.8%       CTCATGAGCTGTCCCCACATTAGGCTTAAAAACAGATGCAATTCCCGGACGTCTAAACCAAACCACTTTCACCGCTACAC
 9 NC_012920.1            99.9%  99.8%       CTCATGAGCTGTCCCCACATTAGGCTTAAAAACAGATGCAATTCCCGGACGTCTAAACCAAACCACTTTCACCGCTACAC
   consensus/100%                            CTCATGAGCTGTCCCCACATTAGGCTTAAAAACAGATGCAATTCCCGGACGTCTAAACCAAACCACTTTCACCGCTACAC
   consensus/90%                             CTCATGAGCTGTCCCCACATTAGGCTTAAAAACAGATGCAATTCCCGGACGTCTAAACCAAACCACTTTCACCGCTACAC
   consensus/80%                             CTCATGAGCTGTCCCCACATTAGGCTTAAAAACAGATGCAATTCCCGGACGTCTAAACCAAACCACTTTCACCGCTACAC
   consensus/70%                             CTCATGAGCTGTCCCCACATTAGGCTTAAAAACAGATGCAATTCCCGGACGTCTAAACCAAACCACTTTCACCGCTACAC

                             cov    pid  8161          .         .         .         2         .         .         .         . 8240
 1 AGENOME-ZPMRG-U-V1.0  100.0% 100.0%       GACCGGGGGTATACTACGGTCAATGCTCTGAAATCTGTGGAGCAAACCACAGTTTCATGCCCATCGTCCTAGAATTAATT
 2 AGENOME-ZPMCG-V1.0    100.0%  99.9%       GACCGGGGGTATACTACGGTCAATGCTCTGAAATCTGTGGAGCAAACCACAGTTTCATGCCCATCGTCCTAGAATTAATT
 3 AGENOME-ZPMRG-M-V1.0  100.0%  99.8%       GACCGGGGGTATACTACGGTCAATGCTCTGAAATCTGTGGAGCAAACCACAGTTTCATGCCCATCGTCCTAGAATTAATT
 4 AGENOME-ZPMRG-F-V1.0   99.9%  99.8%       GACCGGGGGTATACTACGGTCAATGCTCTGAAATCTGTGGAGCAAACCACAGTTTCATGCCCATCGTCCTAGAATTAATT
 5 AGENOME-ZPMRG-Z-V1.0   99.9%  99.7%       GACCGGGGGTATACTACGGTCAATGCTCTGAAATCTGTGGAGCAAACCACAGTTTCATGCCCATCGTCCTAGAATTAATT
 6 AGENOME-ZPMRG-T-V1.0   99.9%  99.8%       GACCGGGGGTATACTACGGTCAATGCTCTGAAATCTGTGGAGCAAACCACAGTTTCATGCCCATCGTCCTAGAATTAATT
 7 AGENOME-ZPMRG-HV-V1.0  99.9%  99.8%       GACCGGGGGTATACTACGGTCAATGCTCTGAAATCTGTGGAGCAAACCACAGTTTCATGCCCATCGTCCTAGAATTAATT
 8 AGENOME-ZPMRG-A-V1.0   99.9%  99.8%       GACCGGGGGTATACTACGGTCAATGCTCTGAAATCTGTGGAGCAAACCACAGTTTCATGCCCATCGTCCTAGAATTAATT
 9 NC_012920.1            99.9%  99.8%       GACCGGGGGTATACTACGGTCAATGCTCTGAAATCTGTGGAGCAAACCACAGTTTCATGCCCATCGTCCTAGAATTAATT
   consensus/100%                            GACCGGGGGTATACTACGGTCAATGCTCTGAAATCTGTGGAGCAAACCACAGTTTCATGCCCATCGTCCTAGAATTAATT
```

```
   consensus/90%                          GACCGGGGGTATACTACGGTCAATGCTCTGAAATCTGTGGAGCAAACCACAGTTTCATGCCCATCGTCCTAGAATTAATT
   consensus/80%                          GACCGGGGGTATACTACGGTCAATGCTCTGAAATCTGTGGAGCAAACCACAGTTTCATGCCCATCGTCCTAGAATTAATT
   consensus/70%                          GACCGGGGGTATACTACGGTCAATGCTCTGAAATCTGTGGAGCAAACCACAGTTTCATGCCCATCGTCCTAGAATTAATT

                          cov    pid  8241          :         .         .         .         .         3         .         . 8320
 1 AGENOME-ZPMRG-U-V1.0  100.0% 100.0%     CCCCTAAAAATCTTTGAAATAGGGCCCGTATTTACCCTATAGCACCCCCTCTACCCCCTCTAGAGCCCACTGTAAAGCTA
 2 AGENOME-ZPMCG-V1.0    100.0%  99.9%     CCCCTAAAAATCTTTGAAATAGGGCCCGTATTTACCCTATAGCACCCCCTCTACCCCCTCTAGAGCCCACTGTAAAGCTA
 3 AGENOME-ZPMRG-M-V1.0  100.0%  99.8%     CCCCTAAAAATCTTTGAAATAGGGCCCGTATTTACCCTATAGCACCCCCTCTACCCCCTCTAGAGCCCACTGTAAAGCTA
 4 AGENOME-ZPMRG-F-V1.0   99.9%  99.8%     CCCCTAAAAATCTTTGAAATAGGGCCCGTATTTACCCTATAGCACCCCCTCTACCCCCTCTAGAGCCCACTGTAAAGCTA
 5 AGENOME-ZPMRG-Z-V1.0   99.9%  99.7%     CCCCTAAAAATCTTTGAAATAGGGCCCGTATTTACCCTATAGCACCCCCTCTACCCCCTCTAGAGCCCACTGTAAAGCTA
 6 AGENOME-ZPMRG-T-V1.0   99.9%  99.8%     CCCCTAAAAATCTTTGAAATAGGGCCCGTATTTACCCTATAGCACCCCCTCTACCCCCTCTAGAGCCCACTGTAAAGCTA
 7 AGENOME-ZPMRG-HV-V1.0  99.9%  99.8%     CCCCTAAAAATCTTTGAAATAGGGCCCGTATTTACCCTATAGCACCCCCTCTACCCCCTCTAGAGCCCACTGTAAAGCTA
 8 AGENOME-ZPMRG-A-V1.0   99.9%  99.8%     CCCCTAAAAATCTTTGAAATAGGGCCCGTATTTACCCTATAGCACCCCCTCTACCCCCTCTAGAGCCCACTGTAAAGCTA
 9 NC_012920.1            99.9%  99.8%     CCCCTAAAAATCTTTGAAATAGGGCCCGTATTTACCCTATAGCACCCCCTCTACCCCCTCTAGAGCCCACTGTAAAGCTA
   consensus/100%                          CCCCTAAAAATCTTTGAAATAGGGCCCGTATTTACCCTATAGCACCCCCTCTACCCCCTCTAGAGCCCACTGTAAAGCTA
   consensus/90%                           CCCCTAAAAATCTTTGAAATAGGGCCCGTATTTACCCTATAGCACCCCCTCTACCCCCTCTAGAGCCCACTGTAAAGCTA
   consensus/80%                           CCCCTAAAAATCTTTGAAATAGGGCCCGTATTTACCCTATAGCACCCCCTCTACCCCCTCTAGAGCCCACTGTAAAGCTA
   consensus/70%                           CCCCTAAAAATCTTTGAAATAGGGCCCGTATTTACCCTATAGCACCCCCTCTACCCCCTCTAGAGCCCACTGTAAAGCTA

                          cov    pid  8321          .         .         :         .         .         .         .         4 8400
 1 AGENOME-ZPMRG-U-V1.0  100.0% 100.0%     ACTTAGCATTAACCTTTTAAGTTAAAGATTAAGAGAACCAACACCTCTTTACAGTGAAATGCCCCAACTAAATACTACCG
 2 AGENOME-ZPMCG-V1.0    100.0%  99.9%     ACTTAGCATTAACCTTTTAAGTTAAAGATTAAGAGAACCAACACCTCTTTACAGTGAAATGCCCCAACTAAATACTACCG
 3 AGENOME-ZPMRG-M-V1.0  100.0%  99.8%     ACTTAGCATTAACCTTTTAAGTTAAAGATTAAGAGAACCAACACCTCTTTACAGTGAAATGCCCCAACTAAATACTACCG
 4 AGENOME-ZPMRG-F-V1.0   99.9%  99.8%     ACTTAGCATTAACCTTTTAAGTTAAAGATTAAGAGAACCAACACCTCTTTACAGTGAAATGCCCCAACTAAATACTACCG
 5 AGENOME-ZPMRG-Z-V1.0   99.9%  99.7%     ACTTAGCATTAACCTTTTAAGTTAAAGATTAAGAGAACCAACACCTCTTTACAGTGAAATGCCCCAACTAAATACTACCG
 6 AGENOME-ZPMRG-T-V1.0   99.9%  99.8%     ACTTAGCATTAACCTTTTAAGTTAAAGATTAAGAGAACCAACACCTCTTTACAGTGAAATGCCCCAACTAAATACTACCG
 7 AGENOME-ZPMRG-HV-V1.0  99.9%  99.8%     ACTTAGCATTAACCTTTTAAGTTAAAGATTAAGAGAACCAACACCTCTTTACAGTGAAATGCCCCAACTAAATACTACCG
 8 AGENOME-ZPMRG-A-V1.0   99.9%  99.8%     ACTTAGCATTAACCTTTTAAGTTAAAGATTAAGAGAACCAACACCTCTTTACAGTGAAATGCCCCAACTAAATACTACCG
 9 NC_012920.1            99.9%  99.8%     ACTTAGCATTAACCTTTTAAGTTAAAGATTAAGAGAACCAACACCTCTTTACAGTGAAATGCCCCAACTAAATACTACCG
   consensus/100%                          ACTTAGCATTAACCTTTTAAGTTAAAGATTAAGAGAACCAACACCTCTTTACAGTGAAATGCCCCAACTAAATACTACCG
   consensus/90%                           ACTTAGCATTAACCTTTTAAGTTAAAGATTAAGAGAACCAACACCTCTTTACAGTGAAATGCCCCAACTAAATACTACCG
   consensus/80%                           ACTTAGCATTAACCTTTTAAGTTAAAGATTAAGAGAACCAACACCTCTTTACAGTGAAATGCCCCAACTAAATACTACCG
   consensus/70%                           ACTTAGCATTAACCTTTTAAGTTAAAGATTAAGAGAACCAACACCTCTTTACAGTGAAATGCCCCAACTAAATACTACCG

                          cov    pid  8401          .         .         .         .         :         .         .         . 8480
 1 AGENOME-ZPMRG-U-V1.0  100.0% 100.0%     TATGGCCCACCATAATTACCCCCATACTCCTTACACTATTCCTCATCACCCAACTAAAAATATTAAACACAAACTACCAC
 2 AGENOME-ZPMCG-V1.0    100.0%  99.9%     TATGGCCCACCATAATTACCCCCATACTCCTTACACTATTCCTCATCACCCAACTAAAAATATTAAACACAAACTACCAC
 3 AGENOME-ZPMRG-M-V1.0  100.0%  99.8%     TATGGCCCACCATAATTACCCCCATACTCCTTACACTATTCCTCATCACCCAACTAAAAATATTAAACACAAACTACCAC
 4 AGENOME-ZPMRG-F-V1.0   99.9%  99.8%     TATGGCCCACCATAATTACCCCCATACTCCTTACACTATTCCTCATCACCCAACTAAAAATATTAAACACAAACTACCAC
 5 AGENOME-ZPMRG-Z-V1.0   99.9%  99.7%     TATGGCCCACCATAATTACCCCCATACTCCTTACACTATTCCTCATCACCCAACTAAAAATATTAAACACAAACTACCAC
 6 AGENOME-ZPMRG-T-V1.0   99.9%  99.8%     TATGGCCCACCATAATTACCCCCATACTCCTTACACTATTCCTCATCACCCAACTAAAAATATTAAACACAAACTACCAC
 7 AGENOME-ZPMRG-HV-V1.0  99.9%  99.8%     TATGGCCCACCATAATTACCCCCATACTCCTTACACTATTCCTCATCACCCAACTAAAAATATTAAACACAAACTACCAC
 8 AGENOME-ZPMRG-A-V1.0   99.9%  99.8%     TATGGCCCACCATAATTACCCCCATACTCCTTACACTATTCCTCATCACCCAACTAAAAATATTAAACACAAACTACCAC
 9 NC_012920.1            99.9%  99.8%     TATGGCCCACCATAATTACCCCCATACTCCTTACACTATTCCTCATCACCCAACTAAAAATATTAAACACAAACTACCAC
   consensus/100%                          TATGGCCCACCATAATTACCCCCATACTCCTTACACTATTCCTCATCACCCAACTAAAAATATTAAACACAAACTACCAC
   consensus/90%                           TATGGCCCACCATAATTACCCCCATACTCCTTACACTATTCCTCATCACCCAACTAAAAATATTAAACACAAACTACCAC
   consensus/80%                           TATGGCCCACCATAATTACCCCCATACTCCTTACACTATTCCTCATCACCCAACTAAAAATATTAAACACAAACTACCAC
   consensus/70%                           TATGGCCCACCATAATTACCCCCATACTCCTTACACTATTCCTCATCACCCAACTAAAAATATTAAACACAAACTACCAC

                          cov    pid  8481          .         5         .         .         .         .         :         . 8560
 1 AGENOME-ZPMRG-U-V1.0  100.0% 100.0%     CTACCTCCCTCACCAAAGCCCATAAAAATAAAAAATTATAACAAACCCTGAGAACCAAAATGAACGAAAATCTGTTCGCT
 2 AGENOME-ZPMCG-V1.0    100.0%  99.9%     CTACCTCCCTCACCAAAGCCCATAAAAATAAAAAATTATAACAAACCCTGAGAACCAAAATGAACGAAAATCTGTTCGCT
 3 AGENOME-ZPMRG-M-V1.0  100.0%  99.8%     CTACCTCCCTCACCAAAGCCCATAAAAATAAAAAATTATAACAAACCCTGAGAACCAAAATGAACGAAAATCTGTTCGCT
 4 AGENOME-ZPMRG-F-V1.0   99.9%  99.8%     CTACCTCCCTCACCAAAGCCCATAAAAATAAAAAATTATAACAAACCCTGAGAACCAAAATGAACGAAAATCTGTTCGCT
 5 AGENOME-ZPMRG-Z-V1.0   99.9%  99.7%     CTACCTCCCTCACCAAAGCCCATAAAAATAAAAAATTATAACAAACCCTGAGAACCAAAATGAACGAAAATCTGTTCGCT
 6 AGENOME-ZPMRG-T-V1.0   99.9%  99.8%     CTACCTCCCTCACCAAAGCCCATAAAAATAAAAAATTATAACAAACCCTGAGAACCAAAATGAACGAAAATCTGTTCGCT
 7 AGENOME-ZPMRG-HV-V1.0  99.9%  99.8%     CTACCTCCCTCACCAAAGCCCATAAAAATAAAAAATTATAACAAACCCTGAGAACCAAAATGAACGAAAATCTGTTCGCT
 8 AGENOME-ZPMRG-A-V1.0   99.9%  99.8%     CTACCTCCCTCACCAAAGCCCATAAAAATAAAAAATTATAACAAACCCTGAGAACCAAAATGAACGAAAATCTGTTCGCT
 9 NC_012920.1            99.9%  99.8%     CTACCTCCCTCACCAAAGCCCATAAAAATAAAAAATTATAACAAACCCTGAGAACCAAAATGAACGAAAATCTGTTCGCT
   consensus/100%                          CTACCTCCCTCACCAAAGCCCATAAAAATAAAAAATTATAACAAACCCTGAGAACCAAAATGAACGAAAATCTGTTCGCT
   consensus/90%                           CTACCTCCCTCACCAAAGCCCATAAAAATAAAAAATTATAACAAACCCTGAGAACCAAAATGAACGAAAATCTGTTCGCT
   consensus/80%                           CTACCTCCCTCACCAAAGCCCATAAAAATAAAAAATTATAACAAACCCTGAGAACCAAAATGAACGAAAATCTGTTCGCT
   consensus/70%                           CTACCTCCCTCACCAAAGCCCATAAAAATAAAAAATTATAACAAACCCTGAGAACCAAAATGAACGAAAATCTGTTCGCT

                          cov    pid  8561          .         .         .         6         .         .         .         . 8640
 1 AGENOME-ZPMRG-U-V1.0  100.0% 100.0%     TCATTCATTGCCCCCACAATCCTAGGCCTACCCGCCGCAGTACTGATCATTCTATTTCCCCCTCTATTGATCCCCACCTC
 2 AGENOME-ZPMCG-V1.0    100.0%  99.9%     TCATTCATTGCCCCCACAATCCTAGGCCTACCCGCCGCAGTACTGATCATTCTATTTCCCCCTCTATTGATCCCCACCTC
 3 AGENOME-ZPMRG-M-V1.0  100.0%  99.8%     TCATTCATTGCCCCCACAATCCTAGGCCTACCCGCCGCAGTACTGATCATTCTATTTCCCCCTCTATTGATCCCCACCTC
 4 AGENOME-ZPMRG-F-V1.0   99.9%  99.8%     TCATTCATTGCCCCCACAATCCTAGGCCTACCCGCCGCAGTGCTGATCATTCTATTTCCCCCTCTATTGATCCCCACCTC
 5 AGENOME-ZPMRG-Z-V1.0   99.9%  99.7%     TCATTCATTGCCCCCACAATCCTAGGCCTACCCGCCGCAGTACTGATCATTCTATTTCCCCCTCTATTGATCCCCACCTC
 6 AGENOME-ZPMRG-T-V1.0   99.9%  99.8%     TCATTCATTGCCCCCACAATCCTAGGCCTACCCGCCGCAGTACTGATCATTCTATTTCCCCCTCTATTGATCCCCACCTC
 7 AGENOME-ZPMRG-HV-V1.0  99.9%  99.8%     TCATTCATTGCCCCCACAATCCTAGGCCTACCCGCCGCAGTACTGATCATTCTATTTCCCCCTCTATTGATCCCCACCTC
 8 AGENOME-ZPMRG-A-V1.0   99.9%  99.8%     TCATTCATTGCCCCCACAATCCTAGGCCTACCCGCCGCAGTACTGATCATTCTATTTCCCCCTCTATTGATCCCCACCTC
 9 NC_012920.1            99.9%  99.8%     TCATTCATTGCCCCCACAATCCTAGGCCTACCCGCCGCAGTACTGATCATTCTATTTCCCCCTCTATTGATCCCCACCTC
   consensus/100%                          TCATTCATTGCCCCCACAATCCTAGGCCTACCCGCCGCAGTuCTGATCATTCTATTTCCCCCTCTATTGATCCCCACCTC
   consensus/90%                           TCATTCATTGCCCCCACAATCCTAGGCCTACCCGCCGCAGTuCTGATCATTCTATTTCCCCCTCTATTGATCCCCACCTC
   consensus/80%                           TCATTCATTGCCCCCACAATCCTAGGCCTACCCGCCGCAGTACTGATCATTCTATTTCCCCCTCTATTGATCCCCACCTC
   consensus/70%                           TCATTCATTGCCCCCACAATCCTAGGCCTACCCGCCGCAGTACTGATCATTCTATTTCCCCCTCTATTGATCCCCACCTC

                          cov    pid  8641          :         .         .         .         .         7         .         . 8720
 1 AGENOME-ZPMRG-U-V1.0  100.0% 100.0%     CAAATATCTCATCAACAACCGACTAATCACCACCCAACAATGACTAATCAAACTAACCTCAAAACAAATGATAACCATAC
 2 AGENOME-ZPMCG-V1.0    100.0%  99.9%     CAAATATCTCATCAACAACCGACTAATCACCACCCAACAATGACTAATCAAACTAACCTCAAAACAAATGATAGCCATAC
 3 AGENOME-ZPMRG-M-V1.0  100.0%  99.8%     CAAATATCTCATCAACAACCGACTAATCACCACCCAACAATGACTAATCAAACTAACCTCAAAACAAATGATAGCCATAC
 4 AGENOME-ZPMRG-F-V1.0   99.9%  99.8%     CAAATATCTCATCAACAACCGACTAATCACCACCCAACAATGACTAATCAAACTAACCTCAAAACAAATGATAACCATAC
 5 AGENOME-ZPMRG-Z-V1.0   99.9%  99.7%     CAAATATCTCATCAACAACCGACTAATCACCACCCAACAATGACTAATCAAACTAACCTCAAAACAAATGATAGCCATAC
 6 AGENOME-ZPMRG-T-V1.0   99.9%  99.8%     CAAATATCTCATCAACAACCGACTAATCACCACCCAACAATGACTAATCAAACTAACCTCAAAACAAATAATAACCATAC
 7 AGENOME-ZPMRG-HV-V1.0  99.9%  99.8%     CAAATATCTCATCAACAACCGACTAATCACCACCCAACAATGACTAATCAAACTAACCTCAAAACAAATGATAACCATAC
 8 AGENOME-ZPMRG-A-V1.0   99.9%  99.8%     CAAATATCTCATCAACAACCGACTAATCACCACCCAACAATGACTAATCAAACTAACCTCAAAACAAATGATAACCATAC
 9 NC_012920.1            99.9%  99.8%     CAAATATCTCATCAACAACCGACTAATCACCACCCAACAATGACTAATCAAACTAACCTCAAAACAAATGATAACCATAC
   consensus/100%                          CAAATATCTCATCAACAACCGACTAATCACCACCCAACAATGACTAATCAAACTAACCTCAAAACAAATuATAuCCATAC
   consensus/90%                           CAAATATCTCATCAACAACCGACTAATCACCACCCAACAATGACTAATCAAACTAACCTCAAAACAAATuATAuCCATAC
```

```
   consensus/80%                               CAAATATCTCATCAACAACCGACTAATCACCACCCAACAATGACTAATCAAACTAACCTCAAAACAAATGATAuCCATAC
   consensus/70%                               CAAATATCTCATCAACAACCGACTAATCACCACCCAACAATGACTAATCAAACTAACCTCAAAACAAATGATAuCCATAC

                             cov    pid  8721          .         .         :         .         .         .         .         8 8800
 1 AGENOME-ZPMRG-U-V1.0  100.0% 100.0%      ACAACACTAAAGGACGAACCTGATCTCTTATACTAGTATCCTTAATCATTTTTATTGCCACAACTAACCTCCTCGGACTC
 2 AGENOME-ZPMCG-V1.0    100.0%  99.9%      ACAACACTAAAGGACGAACCTGATCTCTTATACTAGTATCCTTAATCATTTTTATTGCCACAACTAACCTCCTCGGACTC
 3 AGENOME-ZPMRG-M-V1.0  100.0%  99.8%      ACAACACTAAAGGACGAACCTGATCTCTTATACTAGTATCCTTAATCATTTTTATTGCCACAACTAACCTCCTCGGACTC
 4 AGENOME-ZPMRG-F-V1.0   99.9%  99.8%      ACAACACTAAAGGACGAACCTGATCTCTTATACTAGTATCCTTAATCATTTTTATTGCCACAACTAACCTCCTCGGACTC
 5 AGENOME-ZPMRG-Z-V1.0   99.9%  99.7%      ACAACACTAAAGGACGAACCTGATCTCTTATACTAGTATCCTTAATCATTTTTATTGCCACAACTAACCTCCTCGGACTC
 6 AGENOME-ZPMRG-T-V1.0   99.9%  99.8%      ACAACACTAAAGGACGAACCTGATCTCTTATACTAGTATCCTTAATCATTTTTATTGCCACAACTAACCTCCTCGGACTC
 7 AGENOME-ZPMRG-HV-V1.0  99.9%  99.8%      ACAACACTAAAGGACGAACCTGATCTCTTATACTAGTATCCTTAATCATTTTTATTGCCACAACTAACCTCCTCGGACTC
 8 AGENOME-ZPMRG-A-V1.0   99.9%  99.8%      ACAACACTAAAGGACGAACCTGATCTCTTATACTAGTATCCTTAATCATTTTTATTGCCACAACTAACCTCCTCGGACTC
 9 NC_012920.1            99.9%  99.8%      ACAACACTAAAGGACGAACCTGATCTCTTATACTAGTATCCTTAATCATTTTTATTGCCACAACTAACCTCCTCGGACTC
   consensus/100%                           ACAACACTAAAGGACGAACCTGATCTCTTATACTAGTATCCTTAATCATTTTTATTGCCACAACTAACCTCCTCGGACTC
   consensus/90%                            ACAACACTAAAGGACGAACCTGATCTCTTATACTAGTATCCTTAATCATTTTTATTGCCACAACTAACCTCCTCGGACTC
   consensus/80%                            ACAACACTAAAGGACGAACCTGATCTCTTATACTAGTATCCTTAATCATTTTTATTGCCACAACTAACCTCCTCGGACTC
   consensus/70%                            ACAACACTAAAGGACGAACCTGATCTCTTATACTAGTATCCTTAATCATTTTTATTGCCACAACTAACCTCCTCGGACTC

                             cov    pid  8801          .         .         .         .         :         .         .         . 8880
 1 AGENOME-ZPMRG-U-V1.0  100.0% 100.0%      CTGCCTCACTCATTTACACCAACCACCCAACTATCTATAAACCTAGCCATGGCCATCCCCTTATGAGCGGGCGCAGTGAT
 2 AGENOME-ZPMCG-V1.0    100.0%  99.9%      CTGCCTCACTCATTTACACCAACCACCCAACTATCTATAAACCTAGCCATGGCCATCCCCTTATGAGCGGGCGCAGTGAT
 3 AGENOME-ZPMRG-M-V1.0  100.0%  99.8%      CTGCCTCACTCATTTACACCAACCACCCAACTATCTATAAACCTAGCCATGGCCATCCCCTTATGAGCGGGCGCAGTGAT
 4 AGENOME-ZPMRG-F-V1.0   99.9%  99.8%      CTGCCTCACTCATTTACACCAACCACCCAACTATCTATAAACCTAGCCATGGCCATCCCCTTATGAGCGGGCGCAGTGAT
 5 AGENOME-ZPMRG-Z-V1.0   99.9%  99.7%      CTGCCTCACTCATTTACACCAACCACCCAACTATCTATAAACCTAGCCATGGCCATCCCCTTATGAGCGGGCGCAGTGAT
 6 AGENOME-ZPMRG-T-V1.0   99.9%  99.8%      CTGCCTCACTCATTTACACCAACCACCCAACTATCTATAAACCTAGCCATGGCCATCCCCTTATGAGCGGGCGCAGTGAT
 7 AGENOME-ZPMRG-HV-V1.0  99.9%  99.8%      CTGCCTCACTCATTTACACCAACCACCCAACTATCTATAAACCTAGCCATGGCCATCCCCTTATGAGCGGGCGCAGTGAT
 8 AGENOME-ZPMRG-A-V1.0   99.9%  99.8%      CTGCCTCACTCATTTACACCAACCACCCAACTATCTATAAACCTAGCCATGGCCATCCCCTTATGAGCGGGCGCAGTGAT
 9 NC_012920.1            99.9%  99.8%      CTGCCTCACTCATTTACACCAACCACCCAACTATCTATAAACCTAGCCATGGCCATCCCCTTATGAGCGGGCACAGTGAT
   consensus/100%                           CTGCCTCACTCATTTACACCAACCACCCAACTATCTATAAACCTAGCCATGGCCATCCCCTTATGAGCGGGCuCAGTGAT
   consensus/90%                            CTGCCTCACTCATTTACACCAACCACCCAACTATCTATAAACCTAGCCATGGCCATCCCCTTATGAGCGGGCuCAGTGAT
   consensus/80%                            CTGCCTCACTCATTTACACCAACCACCCAACTATCTATAAACCTAGCCATGGCCATCCCCTTATGAGCGGGCGCAGTGAT
   consensus/70%                            CTGCCTCACTCATTTACACCAACCACCCAACTATCTATAAACCTAGCCATGGCCATCCCCTTATGAGCGGGCGCAGTGAT

                             cov    pid  8881          .         9         .         .         .         .         :         . 8960
 1 AGENOME-ZPMRG-U-V1.0  100.0% 100.0%      TATAGGCTTTCGCTCTAAGATTAAAAATGCCCTAGCCCACTTCTTACCACAAGGCACACCTACACCCCTTATCCCCATAC
 2 AGENOME-ZPMCG-V1.0    100.0%  99.9%      TATAGGCTTTCGCTCTAAGATTAAAAATGCCCTAGCCCACTTCTTACCACAAGGCACACCTACACCCCTTATCCCCATAC
 3 AGENOME-ZPMRG-M-V1.0  100.0%  99.8%      TATAGGCTTTCGCTCTAAGATTAAAAATGCCCTAGCCCACTTCTTACCACAAGGCACACCTACACCCCTTATCCCCATAC
 4 AGENOME-ZPMRG-F-V1.0   99.9%  99.8%      TATAGGCTTTCGCTCTAAGATTAAAAATGCCCTAGCCCACTTCTTACCACAAGGCACACCTACACCCCTTATCCCCATAC
 5 AGENOME-ZPMRG-Z-V1.0   99.9%  99.7%      TATAGGCTTTCGCTCTAAGATTAAAAATGCCCTAGCCCACTTCTTACCACAAGGCACACCTACACCCCTTATCCCCATAC
 6 AGENOME-ZPMRG-T-V1.0   99.9%  99.8%      TATAGGCTTTCGCTCTAAGATTAAAAATGCCCTAGCCCACTTCTTACCACAAGGCACACCTACACCCCTTATCCCCATAC
 7 AGENOME-ZPMRG-HV-V1.0  99.9%  99.8%      TATAGGCTTTCGCTCTAAGATTAAAAATGCCCTAGCCCACTTCTTACCACAAGGCACACCTACACCCCTTATCCCCATAC
 8 AGENOME-ZPMRG-A-V1.0   99.9%  99.8%      TATAGGCTTTCGCTCTAAGATTAAAAATGCCCTAGCCCACTTCTTACCACAAGGCACACCTACACCCCTTATCCCCATAC
 9 NC_012920.1            99.9%  99.8%      TATAGGCTTTCGCTCTAAGATTAAAAATGCCCTAGCCCACTTCTTACCACAAGGCACACCTACACCCCTTATCCCCATAC
   consensus/100%                           TATAGGCTTTCGCTCTAAGATTAAAAATGCCCTAGCCCACTTCTTACCACAAGGCACACCTACACCCCTTATCCCCATAC
   consensus/90%                            TATAGGCTTTCGCTCTAAGATTAAAAATGCCCTAGCCCACTTCTTACCACAAGGCACACCTACACCCCTTATCCCCATAC
   consensus/80%                            TATAGGCTTTCGCTCTAAGATTAAAAATGCCCTAGCCCACTTCTTACCACAAGGCACACCTACACCCCTTATCCCCATAC
   consensus/70%                            TATAGGCTTTCGCTCTAAGATTAAAAATGCCCTAGCCCACTTCTTACCACAAGGCACACCTACACCCCTTATCCCCATAC

                             cov    pid  8961          .         .         .         0         .         .         .         . 9040
 1 AGENOME-ZPMRG-U-V1.0  100.0% 100.0%      TAGTTATTATCGAAACCATCAGCCTACTCATTCAACCAATAGCCCTGGCCGTACGCCTAACCGCTAACATTACTGCAGGC
 2 AGENOME-ZPMCG-V1.0    100.0%  99.9%      TAGTTATTATCGAAACCATCAGCCTACTCATTCAACCAATAGCCCTGGCCGTACGCCTAACCGCTAACATTACTGCAGGC
 3 AGENOME-ZPMRG-M-V1.0  100.0%  99.8%      TAGTTATTATCGAAACCATCAGCCTACTCATTCAACCAATAGCCCTGGCCGTACGCCTAACCGCTAACATTACTGCAGGC
 4 AGENOME-ZPMRG-F-V1.0   99.9%  99.8%      TAGTTATTATCGAAACCATCAGCCTACTCATTCAACCAATAGCCCTGGCCGTACGCCTAACCGCTAACATTACTGCAGGC
 5 AGENOME-ZPMRG-Z-V1.0   99.9%  99.7%      TAGTTATTATCGAAACCATCAGCCTACTCATTCAACCAATAGCCCTGGCCGTACGCCTAACCGCTAACATTACTGCAGGC
 6 AGENOME-ZPMRG-T-V1.0   99.9%  99.8%      TAGTTATTATCGAAACCATCAGCCTACTCATTCAACCAATAGCCCTGGCCGTACGCCTAACCGCTAACATTACTGCAGGC
 7 AGENOME-ZPMRG-HV-V1.0  99.9%  99.8%      TAGTTATTATCGAAACCATCAGCCTACTCATTCAACCAATAGCCCTGGCCGTACGCCTAACCGCTAACATTACTGCAGGC
 8 AGENOME-ZPMRG-A-V1.0   99.9%  99.8%      TAGTTATTATCGAAACCATCAGCCTACTCATTCAACCAATAGCCCTGGCCGTACGCCTAACCGCTAACATTACTGCAGGC
 9 NC_012920.1            99.9%  99.8%      TAGTTATTATCGAAACCATCAGCCTACTCATTCAACCAATAGCCCTGGCCGTACGCCTAACCGCTAACATTACTGCAGGC
   consensus/100%                           TAGTTATTATCGAAACCATCAGCCTACTCATTCAACCAATAGCCCTGGCCGTACGCCTAACCGCTAACATTACTGCAGGC
   consensus/90%                            TAGTTATTATCGAAACCATCAGCCTACTCATTCAACCAATAGCCCTGGCCGTACGCCTAACCGCTAACATTACTGCAGGC
   consensus/80%                            TAGTTATTATCGAAACCATCAGCCTACTCATTCAACCAATAGCCCTGGCCGTACGCCTAACCGCTAACATTACTGCAGGC
   consensus/70%                            TAGTTATTATCGAAACCATCAGCCTACTCATTCAACCAATAGCCCTGGCCGTACGCCTAACCGCTAACATTACTGCAGGC

                             cov    pid  9041          :         .         .         .         .         1         .         . 9120
 1 AGENOME-ZPMRG-U-V1.0  100.0% 100.0%      CACCTACTCATGCACCTAATTGGAAGCGCCACCCTAGCAATATCAACCATTAACCTTCCCTCTACACTTATCATCTTCAC
 2 AGENOME-ZPMCG-V1.0    100.0%  99.9%      CACCTACTCATGCACCTAATTGGAAGCGCCACCCTAGCAATATCAACCATTAACCTTCCCTCTACACTTATCATCTTCAC
 3 AGENOME-ZPMRG-M-V1.0  100.0%  99.8%      CACCTACTCATGCACCTAATTGGAAGCGCCACCCTAGCAATATCAACCATTAACCTTCCCTCTACACTTATCATCTTCAC
 4 AGENOME-ZPMRG-F-V1.0   99.9%  99.8%      CACCTACTCATGCACCTAATTGGAAGCGCCACCCTAGCAATATCAACCATTAACCTTCCCTCTACACTTATCATCTTCAC
 5 AGENOME-ZPMRG-Z-V1.0   99.9%  99.7%      CACCTACTCATGCACCTAATTGGAAGCGCCACCCTAGCAATATCAACCATTAACCTTCCCTCTACACTTATCATCTTCAC
 6 AGENOME-ZPMRG-T-V1.0   99.9%  99.8%      CACCTACTCATGCACCTAATTGGAAGCGCCACCCTAGCAATATCAACCATTAACCTTCCCTCTACACTTATCATCTTCAC
 7 AGENOME-ZPMRG-HV-V1.0  99.9%  99.8%      CACCTACTCATGCACCTAATTGGAAGCGCCACCCTAGCAATATCAACCATTAACCTTCCCTCTACACTTATCATCTTCAC
 8 AGENOME-ZPMRG-A-V1.0   99.9%  99.8%      CACCTACTCATGCACCTAATTGGAAGCGCCACCCTAGCAATATCAACCATTAACCTTCCCTCTACACTTATCATCTTCAC
 9 NC_012920.1            99.9%  99.8%      CACCTACTCATGCACCTAATTGGAAGCGCCACCCTAGCAATATCAACCATTAACCTTCCCTCTACACTTATCATCTTCAC
   consensus/100%                           CACCTACTCATGCACCTAATTGGAAGCGCCACCCTAGCAATATCAACCATTAACCTTCCCTCTACACTTATCATCTTCAC
   consensus/90%                            CACCTACTCATGCACCTAATTGGAAGCGCCACCCTAGCAATATCAACCATTAACCTTCCCTCTACACTTATCATCTTCAC
   consensus/80%                            CACCTACTCATGCACCTAATTGGAAGCGCCACCCTAGCAATATCAACCATTAACCTTCCCTCTACACTTATCATCTTCAC
   consensus/70%                            CACCTACTCATGCACCTAATTGGAAGCGCCACCCTAGCAATATCAACCATTAACCTTCCCTCTACACTTATCATCTTCAC

                             cov    pid  9121          .         .         :         .         .         .         .         2 9200
 1 AGENOME-ZPMRG-U-V1.0  100.0% 100.0%      AATTCTAATTCTACTGACTATCCTAGAAATCGCTGTCGCCTTAATCCAAGCCTACGTTTTCACACTTCTAGTAAGCCTCT
 2 AGENOME-ZPMCG-V1.0    100.0%  99.9%      AATTCTAATTCTACTGACTATCCTAGAAATCGCTGTCGCCTTAATCCAAGCCTACGTTTTCACACTTCTAGTAAGCCTCT
 3 AGENOME-ZPMRG-M-V1.0  100.0%  99.8%      AATTCTAATTCTACTGACTATCCTAGAAATCGCTGTCGCCTTAATCCAAGCCTACGTTTTCACACTTCTAGTAAGCCTCT
 4 AGENOME-ZPMRG-F-V1.0   99.9%  99.8%      AATTCTAATTCTACTGACTATCCTAGAAATCGCTGTCGCCTTAATCCAAGCCTACGTTTTCACACTTCTAGTAAGCCTCT
 5 AGENOME-ZPMRG-Z-V1.0   99.9%  99.7%      AATTCTAATTCTACTGACTATCCTAGAAATCGCTGTCGCCTTAATCCAAGCCTACGTTTTCACACTTCTAGTAAGCCTCT
 6 AGENOME-ZPMRG-T-V1.0   99.9%  99.8%      AATTCTAATTCTACTGACTATCCTAGAAATCGCTGTCGCCTTAATCCAAGCCTACGTTTTCACACTTCTAGTAAGCCTCT
 7 AGENOME-ZPMRG-HV-V1.0  99.9%  99.8%      AATTCTAATTCTACTGACTATCCTAGAAATCGCTGTCGCCTTAATCCAAGCCTACGTTTTCACACTTCTAGTAAGCCTCT
 8 AGENOME-ZPMRG-A-V1.0   99.9%  99.8%      AATTCTAATTCTACTGACTATCCTAGAAATCGCTGTCGCCTTAATCCAAGCCTACGTTTTCACACTTCTAGTAAGCCTCT
 9 NC_012920.1            99.9%  99.8%      AATTCTAATTCTACTGACTATCCTAGAAATCGCTGTCGCCTTAATCCAAGCCTACGTTTTCACACTTCTAGTAAGCCTCT
   consensus/100%                           AATTCTAATTCTACTGACTATCCTAGAAATCGCTGTCGCCTTAATCCAAGCCTACGTTTTCACACTTCTAGTAAGCCTCT
   consensus/90%                            AATTCTAATTCTACTGACTATCCTAGAAATCGCTGTCGCCTTAATCCAAGCCTACGTTTTCACACTTCTAGTAAGCCTCT
   consensus/80%                            AATTCTAATTCTACTGACTATCCTAGAAATCGCTGTCGCCTTAATCCAAGCCTACGTTTTCACACTTCTAGTAAGCCTCT
```

```
   consensus/70%                         AATTCTAATTCTACTGACTATCCTAGAAATCGCTGTCGCCTTAATCCAAGCCTACGTTTTCACACTTCTAGTAAGCCTCT

                          cov    pid  9201          .         .         .         .         :         .         .         . 9280
 1 AGENOME-ZPMRG-U-V1.0  100.0% 100.0%      ACCTGCACGACAACACATAATGACCCACCAATCACATGCCTATCATATAGTAAAACCCAGCCCATGACCCCTAACAGGGG
 2 AGENOME-ZPMCG-V1.0    100.0%  99.9%      ACCTGCACGACAACACATAATGACCCACCAATCACATGCCTATCATATAGTAAAACCCAGCCCATGACCCCTAACAGGGG
 3 AGENOME-ZPMRG-M-V1.0  100.0%  99.8%      ACCTGCACGACAACACATAATGACCCACCAATCACATGCCTATCATATAGTAAAACCCAGCCCATGACCCCTAACAGGGG
 4 AGENOME-ZPMRG-F-V1.0   99.9%  99.8%      ACCTGCACGACAACACATAATGACCCACCAATCACATGCCTATCATATAGTAAAACCCAGCCCATGACCCCTAACAGGGG
 5 AGENOME-ZPMRG-Z-V1.0   99.9%  99.7%      ACCTGCACGACAACACATAATGACCCACCAATCACATGCCTATCATATAGTAAAACCCAGCCCATGACCCCTAACAGGGG
 6 AGENOME-ZPMRG-T-V1.0   99.9%  99.8%      ACCTGCACGACAACACATAATGACCCACCAATCACATGCCTATCATATAGTAAAACCCAGCCCATGACCCCTAACAGGGG
 7 AGENOME-ZPMRG-HV-V1.0  99.9%  99.8%      ACCTGCACGACAACACATAATGACCCACCAATCACATGCCTATCATATAGTAAAACCCAGCCCATGACCCCTAACAGGGG
 8 AGENOME-ZPMRG-A-V1.0   99.9%  99.8%      ACCTGCACGACAACACATAATGACCCACCAATCACATGCCTATCATATAGTAAAACCCAGCCCATGACCCCTAACAGGGG
 9 NC_012920.1            99.9%  99.8%      ACCTGCACGACAACACATAATGACCCACCAATCACATGCCTATCATATAGTAAAACCCAGCCCATGACCCCTAACAGGGG
   consensus/100%                       ACCTGCACGACAACACATAATGACCCACCAATCACATGCCTATCATATAGTAAAACCCAGCCCATGACCCCTAACAGGGG
   consensus/90%                        ACCTGCACGACAACACATAATGACCCACCAATCACATGCCTATCATATAGTAAAACCCAGCCCATGACCCCTAACAGGGG
   consensus/80%                        ACCTGCACGACAACACATAATGACCCACCAATCACATGCCTATCATATAGTAAAACCCAGCCCATGACCCCTAACAGGGG
   consensus/70%                        ACCTGCACGACAACACATAATGACCCACCAATCACATGCCTATCATATAGTAAAACCCAGCCCATGACCCCTAACAGGGG

                          cov    pid  9281          .         3         .         .         .         .         :         . 9360
 1 AGENOME-ZPMRG-U-V1.0  100.0% 100.0%      CCCTCTCAGCCCTCCTAATGACCTCCGGCCTAGCCATGTGATTTCACTTCCACTCCATAACGCTCCTCATACTAGGCCTA
 2 AGENOME-ZPMCG-V1.0    100.0%  99.9%      CCCTCTCAGCCCTCCTAATGACCTCCGGCCTAGCCATGTGATTTCACTTCCACTCCATAACGCTCCTCATACTAGGCCTA
 3 AGENOME-ZPMRG-M-V1.0  100.0%  99.8%      CCCTCTCAGCCCTCCTAATGACCTCCGGCCTAGCCATGTGATTTCACTTCCACTCCATAACGCTCCTCATACTAGGCCTA
 4 AGENOME-ZPMRG-F-V1.0   99.9%  99.8%      CCCTCTCAGCCCTCCTAATGACCTCCGGCCTAGCCATGTGATTTCACTTCCACTCCATAACGCTCCTCATACTAGGCCTA
 5 AGENOME-ZPMRG-Z-V1.0   99.9%  99.7%      CCCTCTCAGCCCTCCTAATGACCTCCGGCCTAGCCATGTGATTTCACTTCCACTCCATAACGCTCCTCATACTAGGCCTA
 6 AGENOME-ZPMRG-T-V1.0   99.9%  99.8%      CCCTCTCAGCCCTCCTAATGACCTCCGGCCTAGCCATGTGATTTCACTTCCACTCCATAACGCTCCTCATACTAGGCCTA
 7 AGENOME-ZPMRG-HV-V1.0  99.9%  99.8%      CCCTCTCAGCCCTCCTAATGACCTCCGGCCTAGCCATGTGATTTCACTTCCACTCCATAACGCTCCTCGTACTAGGCCTA
 8 AGENOME-ZPMRG-A-V1.0   99.9%  99.8%      CCCTCTCAGCCCTCCTAATGACCTCCGGCCTAGCCATGTGATTTCACTTCCACTCCATAACGCTCCTCATACTAGGCCTA
 9 NC_012920.1            99.9%  99.8%      CCCTCTCAGCCCTCCTAATGACCTCCGGCCTAGCCATGTGATTTCACTTCCACTCCATAACGCTCCTCATACTAGGCCTA
   consensus/100%                       CCCTCTCAGCCCTCCTAATGACCTCCGGCCTAGCCATGTGATTTCACTTCCACTCCATAACGCTCCTCuTACTAGGCCTA
   consensus/90%                        CCCTCTCAGCCCTCCTAATGACCTCCGGCCTAGCCATGTGATTTCACTTCCACTCCATAACGCTCCTCuTACTAGGCCTA
   consensus/80%                        CCCTCTCAGCCCTCCTAATGACCTCCGGCCTAGCCATGTGATTTCACTTCCACTCCATAACGCTCCTCATACTAGGCCTA
   consensus/70%                        CCCTCTCAGCCCTCCTAATGACCTCCGGCCTAGCCATGTGATTTCACTTCCACTCCATAACGCTCCTCATACTAGGCCTA

                          cov    pid  9361          .         .         .         4         .         .         .         . 9440
 1 AGENOME-ZPMRG-U-V1.0  100.0% 100.0%      CTAACCAACACACTAACCATATACCAATGATGGCGCGATGTAACACGAGAAAGCACATACCAAGGCCACCACACACCACC
 2 AGENOME-ZPMCG-V1.0    100.0%  99.9%      CTAACCAACACACTAACCATATACCAATGATGGCGCGATGTAACACGAGAAAGCACATACCAAGGCCACCACACACCACC
 3 AGENOME-ZPMRG-M-V1.0  100.0%  99.8%      CTAACCAACACACTAACCATATACCAATGATGGCGCGATGTAACACGAGAAAGCACATACCAAGGCCACCACACACCACC
 4 AGENOME-ZPMRG-F-V1.0   99.9%  99.8%      CTAACCAACACACTAACCATATACCAATGATGGCGCGATGTAACACGAGAAAGCACATACCAAGGCCACCACACACCACC
 5 AGENOME-ZPMRG-Z-V1.0   99.9%  99.7%      CTAACCAACACACTAACCATATACCAATGATGGCGCGATGTAACACGAGAAAGCACATACCAAGGCCACCACACACCACC
 6 AGENOME-ZPMRG-T-V1.0   99.9%  99.8%      CTAACCAACACACTAACCATATACCAATGATGGCGCGATGTAACACGAGAAAGCACATACCAAGGCCACCACACACCACC
 7 AGENOME-ZPMRG-HV-V1.0  99.9%  99.8%      CTAACCAACACACTAACCATATACCAATGATGGCGCGATGTAACACGAGAAAGCACATACCAAGGCCACCACACACCACC
 8 AGENOME-ZPMRG-A-V1.0   99.9%  99.8%      CTAACCAACACACTAACCATATACCAATGATGGCGCGATGTAACACGAGAAAGCACATACCAAGGCCACCACACACCACC
 9 NC_012920.1            99.9%  99.8%      CTAACCAACACACTAACCATATACCAATGATGGCGCGATGTAACACGAGAAAGCACATACCAAGGCCACCACACACCACC
   consensus/100%                       CTAACCAACACACTAACCATATACCAATGATGGCGCGATGTAACACGAGAAAGCACATACCAAGGCCACCACACACCACC
   consensus/90%                        CTAACCAACACACTAACCATATACCAATGATGGCGCGATGTAACACGAGAAAGCACATACCAAGGCCACCACACACCACC
   consensus/80%                        CTAACCAACACACTAACCATATACCAATGATGGCGCGATGTAACACGAGAAAGCACATACCAAGGCCACCACACACCACC
   consensus/70%                        CTAACCAACACACTAACCATATACCAATGATGGCGCGATGTAACACGAGAAAGCACATACCAAGGCCACCACACACCACC

                          cov    pid  9441          :         .         .         .         .         5         .         . 9520
 1 AGENOME-ZPMRG-U-V1.0  100.0% 100.0%      TGTCCAAAAAGGCCTTCGATACGGGATAATCCTATTTATTACCTCAGAAGTTTTTTTCTTCGCAGGATTTTTCTGAGCCT
 2 AGENOME-ZPMCG-V1.0    100.0%  99.9%      TGTCCAAAAAGGCCTTCGATACGGGATAATCCTATTTATTACCTCAGAAGTTTTTTTCTTCGCAGGATTTTTCTGAGCCT
 3 AGENOME-ZPMRG-M-V1.0  100.0%  99.8%      TGTCCAAAAAGGCCTTCGATACGGGATAATCCTATTTATTACCTCAGAAGTTTTTTTCTTCGCAGGATTTTTCTGAGCCT
 4 AGENOME-ZPMRG-F-V1.0   99.9%  99.8%      TGTCCAAAAAGGCCTTCGATACGGGATAATCCTATTTATTACCTCAGAAGTTTTTTTCTTCGCAGGATTTTTCTGAGCCT
 5 AGENOME-ZPMRG-Z-V1.0   99.9%  99.7%      TGTCCAAAAAGGCCTTCGATACGGGATAATCCTATTTATTACCTCAGAAGTTTTTTTCTTCGCAGGATTTTTCTGAGCCT
 6 AGENOME-ZPMRG-T-V1.0   99.9%  99.8%      TGTCCAAAAAGGCCTTCGATACGGGATAATCCTATTTATTACCTCAGAAGTTTTTTTCTTCGCAGGATTTTTCTGAGCCT
 7 AGENOME-ZPMRG-HV-V1.0  99.9%  99.8%      TGTCCAAAAAGGCCTTCGATACGGGATAATCCTATTTATTACCTCAGAAGTTTTTTTCTTCGCAGGATTTTTCTGAGCCT
 8 AGENOME-ZPMRG-A-V1.0   99.9%  99.8%      TGTCCAAAAAGGCCTTCGATACGGGATAATCCTATTTATTACCTCAGAAGTTTTTTTCTTCGCAGGATTTTTCTGAGCCT
 9 NC_012920.1            99.9%  99.8%      TGTCCAAAAAGGCCTTCGATACGGGATAATCCTATTTATTACCTCAGAAGTTTTTTTCTTCGCAGGATTTTTCTGAGCCT
   consensus/100%                       TGTCCAAAAAGGCCTTCGATACGGGATAATCCTATTTATTACCTCAGAAGTTTTTTTCTTCGCAGGATTTTTCTGAGCCT
   consensus/90%                        TGTCCAAAAAGGCCTTCGATACGGGATAATCCTATTTATTACCTCAGAAGTTTTTTTCTTCGCAGGATTTTTCTGAGCCT
   consensus/80%                        TGTCCAAAAAGGCCTTCGATACGGGATAATCCTATTTATTACCTCAGAAGTTTTTTTCTTCGCAGGATTTTTCTGAGCCT
   consensus/70%                        TGTCCAAAAAGGCCTTCGATACGGGATAATCCTATTTATTACCTCAGAAGTTTTTTTCTTCGCAGGATTTTTCTGAGCCT

                          cov    pid  9521          .         .         :         .         .         .         .         6 9600
 1 AGENOME-ZPMRG-U-V1.0  100.0% 100.0%      TTTACCACTCCAGCCTAGCCCCTACCCCCCAATTAGGAGGGCACTGGCCCCCAACAGGCATCACCCCGCTAAATCCCCTA
 2 AGENOME-ZPMCG-V1.0    100.0%  99.9%      TTTACCACTCCAGCCTAGCCCCTACCCCCCAACTAGGAGGGCACTGGCCCCCAACAGGCATCACCCCGCTAAATCCCCTA
 3 AGENOME-ZPMRG-M-V1.0  100.0%  99.8%      TTTACCACTCCAGCCTAGCCCCTACCCCCCAACTAGGAGGGCACTGGCCCCCAACAGGCATCACCCCGCTAAATCCCCTA
 4 AGENOME-ZPMRG-F-V1.0   99.9%  99.8%      TTTACCACTCCAGCCTAGCCCCTACCCCCCAATTAGGAGGGCACTGGCCCCCAACAGGCATCACCCCGCTAAATCCCCTA
 5 AGENOME-ZPMRG-Z-V1.0   99.9%  99.7%      TTTACCACTCCAGCCTAGCCCCTACCCCCCAACTAGGAGGGCACTGGCCCCCAACAGGCATCACCCCGCTAAATCCCCTA
 6 AGENOME-ZPMRG-T-V1.0   99.9%  99.8%      TTTACCACTCCAGCCTAGCCCCTACCCCCCAATTAGGAGGGCACTGGCCCCCAACAGGCATCACCCCGCTAAATCCCCTA
 7 AGENOME-ZPMRG-HV-V1.0  99.9%  99.8%      TTTACCACTCCAGCCTAGCCCCTACCCCCCAATTAGGAGGGCACTGGCCCCCAACAGGCATCACCCCGCTAAATCCCCTA
 8 AGENOME-ZPMRG-A-V1.0   99.9%  99.8%      TTTACCACTCCAGCCTAGCCCCTACCCCCCAATTAGGAGGGCACTGGCCCCCAACAGGCATCACCCCGCTAAATCCCCTA
 9 NC_012920.1            99.9%  99.8%      TTTACCACTCCAGCCTAGCCCCTACCCCCCAATTAGGAGGGCACTGGCCCCCAACAGGCATCACCCCGCTAAATCCCCTA
   consensus/100%                       TTTACCACTCCAGCCTAGCCCCTACCCCCCAAsTAGGAGGGCACTGGCCCCCAACAGGCATCACCCCGCTAAATCCCCTA
   consensus/90%                        TTTACCACTCCAGCCTAGCCCCTACCCCCCAAsTAGGAGGGCACTGGCCCCCAACAGGCATCACCCCGCTAAATCCCCTA
   consensus/80%                        TTTACCACTCCAGCCTAGCCCCTACCCCCCAAsTAGGAGGGCACTGGCCCCCAACAGGCATCACCCCGCTAAATCCCCTA
   consensus/70%                        TTTACCACTCCAGCCTAGCCCCTACCCCCCAAsTAGGAGGGCACTGGCCCCCAACAGGCATCACCCCGCTAAATCCCCTA

                          cov    pid  9601          .         .         .         .         :         .         .         . 9680
 1 AGENOME-ZPMRG-U-V1.0  100.0% 100.0%      GAAGTCCCACTCCTAAACACATCCGTATTACTCGCATCAGGAGTATCAATCACCTGAGCTCACCATAGTCTAATAGAAAA
 2 AGENOME-ZPMCG-V1.0    100.0%  99.9%      GAAGTCCCACTCCTAAACACATCCGTATTACTCGCATCAGGAGTATCAATCACCTGAGCTCACCATAGTCTAATAGAAAA
 3 AGENOME-ZPMRG-M-V1.0  100.0%  99.8%      GAAGTCCCACTCCTAAACACATCCGTATTACTCGCATCAGGAGTATCAATCACCTGAGCTCACCATAGTCTAATAGAAAA
 4 AGENOME-ZPMRG-F-V1.0   99.9%  99.8%      GAAGTCCCACTCCTAAACACATCCGTATTACTCGCATCAGGAGTATCAATCACCTGAGCTCACCATAGTCTAATAGAAAA
 5 AGENOME-ZPMRG-Z-V1.0   99.9%  99.7%      GAAGTCCCACTCCTAAACACATCCGTATTACTCGCATCAGGAGTATCAATCACCTGAGCTCACCATAGTCTAATAGAAAA
 6 AGENOME-ZPMRG-T-V1.0   99.9%  99.8%      GAAGTCCCACTCCTAAACACATCCGTATTACTCGCATCAGGAGTATCAATCACCTGAGCTCACCATAGTCTAATAGAAAA
 7 AGENOME-ZPMRG-HV-V1.0  99.9%  99.8%      GAAGTCCCACTCCTAAACACATCCGTATTACTCGCATCAGGAGTATCAATCACCTGAGCTCACCATAGTCTAATAGAAAA
 8 AGENOME-ZPMRG-A-V1.0   99.9%  99.8%      GAAGTCCCACTCCTAAACACATCCGTATTACTCGCATCAGGAGTATCAATCACCTGAGCTCACCATAGTCTAATAGAAAA
 9 NC_012920.1            99.9%  99.8%      GAAGTCCCACTCCTAAACACATCCGTATTACTCGCATCAGGAGTATCAATCACCTGAGCTCACCATAGTCTAATAGAAAA
   consensus/100%                       GAAGTCCCACTCCTAAACACATCCGTATTACTCGCATCAGGAGTATCAATCACCTGAGCTCACCATAGTCTAATAGAAAA
   consensus/90%                        GAAGTCCCACTCCTAAACACATCCGTATTACTCGCATCAGGAGTATCAATCACCTGAGCTCACCATAGTCTAATAGAAAA
   consensus/80%                        GAAGTCCCACTCCTAAACACATCCGTATTACTCGCATCAGGAGTATCAATCACCTGAGCTCACCATAGTCTAATAGAAAA
   consensus/70%                        GAAGTCCCACTCCTAAACACATCCGTATTACTCGCATCAGGAGTATCAATCACCTGAGCTCACCATAGTCTAATAGAAAA
```

```
                              cov    pid  9681          .          7          .          .          .          .          :          . 9760
1 AGENOME-ZPMRG-U-V1.0  100.0% 100.0%       CAACCGAAACCAAATAATTCAAGCACTGCTTATTACAATTTTACTGGGTCTCTATTTTACCCTCCTACAAGCCTCAGAGT
2 AGENOME-ZPMCG-V1.0    100.0%  99.9%       CAACCGAAACCAAATAATTCAAGCACTGCTTATTACAATTTTACTGGGTCTCTATTTTACCCTCCTACAAGCCTCAGAGT
3 AGENOME-ZPMRG-M-V1.0  100.0%  99.8%       CAACCGAAACCAAATAATTCAAGCACTGCTTATTACAATTTTACTGGGTCTCTATTTTACCCTCCTACAAGCCTCAGAGT
4 AGENOME-ZPMRG-F-V1.0   99.9%  99.8%       CAACCGAAACCAAATAATTCAAGCACTGCTTATTACAATTTTACTGGGTCTCTATTTTACCCTCCTACAAGCCTCAGAGT
5 AGENOME-ZPMRG-Z-V1.0   99.9%  99.7%       CAACCGAAACCAAATAATTCAAGCACTGCTTATTACAATTTTACTGGGTCTCTATTTTACCCTCCTACAAGCCTCAGAGT
6 AGENOME-ZPMRG-T-V1.0   99.9%  99.8%       CAACCGAAACCAAATAATTCAAGCACTGCTTATTACAATTTTACTGGGTCTCTATTTTACCCTCCTACAAGCCTCAGAGT
7 AGENOME-ZPMRG-HV-V1.0  99.9%  99.8%       CAACCGAAACCAAATAATTCAAGCACTGCTTATTACAATTTTACTGGGTCTCTATTTTACCCTCCTACAAGCCTCAGAGT
8 AGENOME-ZPMRG-A-V1.0   99.9%  99.8%       CAACCGAAACCAAATAATTCAAGCACTGCTTATTACAATTTTACTGGGTCTCTATTTTACCCTCCTACAAGCCTCAGAGT
9 NC_012920.1            99.9%  99.8%       CAACCGAAACCAAATAATTCAAGCACTGCTTATTACAATTTTACTGGGTCTCTATTTTACCCTCCTACAAGCCTCAGAGT
  consensus/100%                            CAACCGAAACCAAATAATTCAAGCACTGCTTATTACAATTTTACTGGGTCTCTATTTTACCCTCCTACAAGCCTCAGAGT
  consensus/90%                             CAACCGAAACCAAATAATTCAAGCACTGCTTATTACAATTTTACTGGGTCTCTATTTTACCCTCCTACAAGCCTCAGAGT
  consensus/80%                             CAACCGAAACCAAATAATTCAAGCACTGCTTATTACAATTTTACTGGGTCTCTATTTTACCCTCCTACAAGCCTCAGAGT
  consensus/70%                             CAACCGAAACCAAATAATTCAAGCACTGCTTATTACAATTTTACTGGGTCTCTATTTTACCCTCCTACAAGCCTCAGAGT

                              cov    pid  9761          .          .          .          8          .          .          .          . 9840
1 AGENOME-ZPMRG-U-V1.0  100.0% 100.0%       ACTTCGAGTCTCCCTTCACCATTTCCGACGGCATCTACGGCTCAACATTTTTTGTAGCCACAGGCTTCCACGGACTTCAC
2 AGENOME-ZPMCG-V1.0    100.0%  99.9%       ACTTCGAGTCTCCCTTCACCATTTCCGACGGCATCTACGGCTCAACATTTTTTGTAGCCACAGGCTTCCACGGACTTCAC
3 AGENOME-ZPMRG-M-V1.0  100.0%  99.8%       ACTTCGAGTCTCCCTTCACCATTTCCGACGGCATCTACGGCTCAACATTTTTTGTAGCCACAGGCTTCCACGGACTTCAC
4 AGENOME-ZPMRG-F-V1.0   99.9%  99.8%       ACTTCGAGTCTCCCTTCACCATTTCCGACGGCATCTACGGCTCAACATTTTTTGTAGCCACAGGCTTCCACGGACTTCAC
5 AGENOME-ZPMRG-Z-V1.0   99.9%  99.7%       ACTTCGAGTCTCCCTTCACCATTTCCGACGGCATCTACGGCTCAACATTTTTTGTAGCCACAGGCTTCCACGGACTTCAC
6 AGENOME-ZPMRG-T-V1.0   99.9%  99.8%       ACTTCGAGTCTCCCTTCACCATTTCCGACGGCATCTACGGCTCAACATTTTTTGTAGCCACAGGCTTCCACGGACTTCAC
7 AGENOME-ZPMRG-HV-V1.0  99.9%  99.8%       ACTTCGAGTCTCCCTTCACCATTTCCGACGGCATCTACGGCTCAACATTTTTTGTAGCCACAGGCTTCCACGGACTTCAC
8 AGENOME-ZPMRG-A-V1.0   99.9%  99.8%       ACTTCGAGTCTCCCTTCACCATTTCCGACGGCATCTACGGCTCAACATTTTTTGTAGCCACAGGCTTCCACGGACTTCAC
9 NC_012920.1            99.9%  99.8%       ACTTCGAGTCTCCCTTCACCATTTCCGACGGCATCTACGGCTCAACATTTTTTGTAGCCACAGGCTTCCACGGACTTCAC
  consensus/100%                            ACTTCGAGTCTCCCTTCACCATTTCCGACGGCATCTACGGCTCAACATTTTTTGTAGCCACAGGCTTCCACGGACTTCAC
  consensus/90%                             ACTTCGAGTCTCCCTTCACCATTTCCGACGGCATCTACGGCTCAACATTTTTTGTAGCCACAGGCTTCCACGGACTTCAC
  consensus/80%                             ACTTCGAGTCTCCCTTCACCATTTCCGACGGCATCTACGGCTCAACATTTTTTGTAGCCACAGGCTTCCACGGACTTCAC
  consensus/70%                             ACTTCGAGTCTCCCTTCACCATTTCCGACGGCATCTACGGCTCAACATTTTTTGTAGCCACAGGCTTCCACGGACTTCAC

                              cov    pid  9841          :          .          .          .          .          9          .          . 9920
1 AGENOME-ZPMRG-U-V1.0  100.0% 100.0%       GTCATTATTGGCTCAACTTTCCTCACTATCTGCTTCATCCGCCAACTAATATTTCACTTTACATCCAAACATCACTTTGG
2 AGENOME-ZPMCG-V1.0    100.0%  99.9%       GTCATTATTGGCTCAACTTTCCTCACTATCTGCTTCATCCGCCAACTAATATTTCACTTTACATCCAAACATCACTTTGG
3 AGENOME-ZPMRG-M-V1.0  100.0%  99.8%       GTCATTATTGGCTCAACTTTCCTCACTATCTGCTTCATCCGCCAACTAATATTTCACTTTACATCCAAACATCACTTTGG
4 AGENOME-ZPMRG-F-V1.0   99.9%  99.8%       GTCATTATTGGCTCAACTTTCCTCACTATCTGCTTCATCCGCCAACTAATATTTCACTTTACATCCAAACATCACTTTGG
5 AGENOME-ZPMRG-Z-V1.0   99.9%  99.7%       GTCATTATTGGCTCAACTTTCCTCACTATCTGCTTCATCCGCCAACTAATATTTCACTTTACATCCAAACATCACTTTGG
6 AGENOME-ZPMRG-T-V1.0   99.9%  99.8%       GTCATTATTGGCTCAACTTTCCTCACTATCTGCTTCATCCGCCAACTAATATTTCACTTTACATCCAAACATCACTTTGG
7 AGENOME-ZPMRG-HV-V1.0  99.9%  99.8%       GTCATTATTGGCTCAACTTTCCTCACTATCTGCTTCATCCGCCAACTAATATTTCACTTTACATCCAAACATCACTTTGG
8 AGENOME-ZPMRG-A-V1.0   99.9%  99.8%       GTCATTATTGGCTCAACTTTCCTCACTATCTGCTTCATCCGCCAACTAATATTTCACTTTACATCCAAACATCACTTTGG
9 NC_012920.1            99.9%  99.8%       GTCATTATTGGCTCAACTTTCCTCACTATCTGCTTCATCCGCCAACTAATATTTCACTTTACATCCAAACATCACTTTGG
  consensus/100%                            GTCATTATTGGCTCAACTTTCCTCACTATCTGCTTCATCCGCCAACTAATATTTCACTTTACATCCAAACATCACTTTGG
  consensus/90%                             GTCATTATTGGCTCAACTTTCCTCACTATCTGCTTCATCCGCCAACTAATATTTCACTTTACATCCAAACATCACTTTGG
  consensus/80%                             GTCATTATTGGCTCAACTTTCCTCACTATCTGCTTCATCCGCCAACTAATATTTCACTTTACATCCAAACATCACTTTGG
  consensus/70%                             GTCATTATTGGCTCAACTTTCCTCACTATCTGCTTCATCCGCCAACTAATATTTCACTTTACATCCAAACATCACTTTGG

                              cov    pid  9921          .          .          :          .          .          .          .          0 10000
1 AGENOME-ZPMRG-U-V1.0  100.0% 100.0%       CTTCGAAGCCGCCGCCTGATACTGGCATTTTGTAGATGTGGTTTGACTATTTCTGTATGTCTCCATCTATTGATGAGGGT
2 AGENOME-ZPMCG-V1.0    100.0%  99.9%       CTTCGAAGCCGCCGCCTGATACTGGCATTTTGTAGATGTGGTTTGACTATTTCTGTATGTCTCCATCTATTGATGAGGGT
3 AGENOME-ZPMRG-M-V1.0  100.0%  99.8%       CTTCGAAGCCGCCGCCTGATACTGGCATTTTGTAGATGTGGTTTGACTATTTCTGTATGTCTCCATCTATTGATGAGGGT
4 AGENOME-ZPMRG-F-V1.0   99.9%  99.8%       CTTCGAAGCCGCCGCCTGATACTGGCATTTTGTAGATGTGGTTTGACTATTTCTGTATGTCTCCATCTATTGATGAGGGT
5 AGENOME-ZPMRG-Z-V1.0   99.9%  99.7%       CTTCGAAGCCGCCGCCTGATACTGGCATTTTGTAGATGTGGTTTGACTATTTCTGTATGTCTCCATCTATTGATGAGGGT
6 AGENOME-ZPMRG-T-V1.0   99.9%  99.8%       CTTCGAAGCCGCCGCCTGATACTGGCATTTTGTAGATGTGGTTTGACTATTTCTGTATGTCTCCATCTATTGATGAGGGT
7 AGENOME-ZPMRG-HV-V1.0  99.9%  99.8%       CTTCGAAGCCGCCGCCTGATACTGGCATTTTGTAGATGTGGTTTGACTATTTCTGTATGTCTCCATCTATTGATGAGGGT
8 AGENOME-ZPMRG-A-V1.0   99.9%  99.8%       CTTCGAAGCCGCCGCCTGATACTGGCATTTTGTAGATGTGGTTTGACTATTTCTGTATGTCTCCATCTATTGATGAGGGT
9 NC_012920.1            99.9%  99.8%       CTTCGAAGCCGCCGCCTGATACTGGCATTTTGTAGATGTGGTTTGACTATTTCTGTATGTCTCCATCTATTGATGAGGGT
  consensus/100%                            CTTCGAAGCCGCCGCCTGATACTGGCATTTTGTAGATGTGGTTTGACTATTTCTGTATGTCTCCATCTATTGATGAGGGT
  consensus/90%                             CTTCGAAGCCGCCGCCTGATACTGGCATTTTGTAGATGTGGTTTGACTATTTCTGTATGTCTCCATCTATTGATGAGGGT
  consensus/80%                             CTTCGAAGCCGCCGCCTGATACTGGCATTTTGTAGATGTGGTTTGACTATTTCTGTATGTCTCCATCTATTGATGAGGGT
  consensus/70%                             CTTCGAAGCCGCCGCCTGATACTGGCATTTTGTAGATGTGGTTTGACTATTTCTGTATGTCTCCATCTATTGATGAGGGT

                              cov    pid 10001          .          .          .          .          :          .          .          . 10080
1 AGENOME-ZPMRG-U-V1.0  100.0% 100.0%       CTTACTCTTTTAGTATAAATAGTACCGTTAACTTCCAATTAACTAGTTTTGACAACATTCAAAAAAGAGTAATAAACTTC
2 AGENOME-ZPMCG-V1.0    100.0%  99.9%       CTTACTCTTTTAGTATAAATAGTACCGTTAACTTCCAATTAACTAGTTTTGACAACATTCAAAAAAGAGTAATAAACTTC
3 AGENOME-ZPMRG-M-V1.0  100.0%  99.8%       CTTACTCTTTTAGTATAAATAGTACCGTTAACTTCCAATTAACTAGTTTTGACAACATTCAAAAAAGAGTAATAAACTTC
4 AGENOME-ZPMRG-F-V1.0   99.9%  99.8%       CTTACTCTTTTAGTATAAATAGTACCGTTAACTTCCAATTAACTAGTTTTGACAACATTCAAAAAAGAGTAATAAACTTC
5 AGENOME-ZPMRG-Z-V1.0   99.9%  99.7%       CTTACTCTTTTAGTATAAATAGTACCGTTAACTTCCAATTAACTAGTTTTGACAACATTCAAAAAAGAGTAATAAACTTC
6 AGENOME-ZPMRG-T-V1.0   99.9%  99.8%       CTTACTCTTTTAGTATAAATAGTACCGTTAACTTCCAATTAACTAGTTTTGACAACATTCAAAAAAGAGTAATAAACTTC
7 AGENOME-ZPMRG-HV-V1.0  99.9%  99.8%       CTTACTCTTTTAGTATAAATAGTACCGTTAACTTCCAATTAACTAGTTTTGACAACATTCAAAAAAGAGTAATAAACTTC
8 AGENOME-ZPMRG-A-V1.0   99.9%  99.8%       CTTACTCTTTTAGTATAAATAGTACCGTTAACTTCCAATTAACTAGTTTTGACAACATTCAAAAAAGAGTAATAAACTTC
9 NC_012920.1            99.9%  99.8%       CTTACTCTTTTAGTATAAATAGTACCGTTAACTTCCAATTAACTAGTTTTGACAACATTCAAAAAAGAGTAATAAACTTC
  consensus/100%                            CTTACTCTTTTAGTATAAATAGTACCGTTAACTTCCAATTAACTAGTTTTGACAACATTCAAAAAAGAGTAATAAACTTC
  consensus/90%                             CTTACTCTTTTAGTATAAATAGTACCGTTAACTTCCAATTAACTAGTTTTGACAACATTCAAAAAAGAGTAATAAACTTC
  consensus/80%                             CTTACTCTTTTAGTATAAATAGTACCGTTAACTTCCAATTAACTAGTTTTGACAACATTCAAAAAAGAGTAATAAACTTC
  consensus/70%                             CTTACTCTTTTAGTATAAATAGTACCGTTAACTTCCAATTAACTAGTTTTGACAACATTCAAAAAAGAGTAATAAACTTC

                              cov    pid 10081          .          1          .          .          .          .          :          . 10160
1 AGENOME-ZPMRG-U-V1.0  100.0% 100.0%       GCCTTAATTTTAATAATCAACACCCTCCTAGCCTTACTACTAATAATTATTACATTTTGACTACCACAACTCAACGGCTA
2 AGENOME-ZPMCG-V1.0    100.0%  99.9%       GCCTTAATTTTAATAATCAACACCCTCCTAGCCTTACTACTAATAATTATTACATTTTGACTACCACAACTCAACGGCTA
3 AGENOME-ZPMRG-M-V1.0  100.0%  99.8%       GCCTTAATTTTAATAATCAACACCCTCCTAGCCTTACTACTAATAATTATTACATTTTGACTACCACAACTCAACGGCTA
4 AGENOME-ZPMRG-F-V1.0   99.9%  99.8%       GCCTTAATTTTAATAATCAACACCCTCCTAGCCTTACTACTAATAATTATTACATTTTGACTACCACAACTCAACGGCTA
5 AGENOME-ZPMRG-Z-V1.0   99.9%  99.7%       GCCTTAATTTTAATAATCAACACCCTCCTAGCCTTACTACTAATAATTATTACATTTTGACTACCACAACTCAACGGCTA
6 AGENOME-ZPMRG-T-V1.0   99.9%  99.8%       GCCTTAATTTTAATAATCAACACCCTCCTAGCCTTACTACTAATAATTATTACATTTTGACTACCACAACTCAACGGCTA
7 AGENOME-ZPMRG-HV-V1.0  99.9%  99.8%       GCCTTAATTTTAATAATCAACACCCTCCTAGCCTTACTACTAATAATTATTACATTTTGACTACCACAACTCAACGGCTA
8 AGENOME-ZPMRG-A-V1.0   99.9%  99.8%       GCCTTAATTTTAATAATCAACACCCTCCTAGCCTTACTACTAATAATTATTACATTTTGACTACCACAACTCAACGGCTA
9 NC_012920.1            99.9%  99.8%       GCCTTAATTTTAATAATCAACACCCTCCTAGCCTTACTACTAATAATTATTACATTTTGACTACCACAACTCAACGGCTA
  consensus/100%                            GCCTTAATTTTAATAATCAACACCCTCCTAGCCTTACTACTAATAATTATTACATTTTGACTACCACAACTCAACGGCTA
  consensus/90%                             GCCTTAATTTTAATAATCAACACCCTCCTAGCCTTACTACTAATAATTATTACATTTTGACTACCACAACTCAACGGCTA
  consensus/80%                             GCCTTAATTTTAATAATCAACACCCTCCTAGCCTTACTACTAATAATTATTACATTTTGACTACCACAACTCAACGGCTA
  consensus/70%                             GCCTTAATTTTAATAATCAACACCCTCCTAGCCTTACTACTAATAATTATTACATTTTGACTACCACAACTCAACGGCTA
```

```
                         cov    pid 10161          .         .         .         2         .         .         .         . 10240
1 AGENOME-ZPMRG-U-V1.0  100.0% 100.0%       CATAGAAAAATCCACCCCTTACGAGTGCGGCTTCGACCCTATATCCCCCGCCCGCGTCCCTTTCTCCATAAAATTCTTCT
2 AGENOME-ZPMCG-V1.0    100.0%  99.9%       CATAGAAAAATCCACCCCTTACGAGTGCGGCTTCGACCCTATATCCCCCGCCCGCGTCCCTTTCTCCATAAAATTCTTCT
3 AGENOME-ZPMRG-M-V1.0  100.0%  99.8%       CATAGAAAAATCCACCCCTTACGAGTGCGGCTTCGACCCTATATCCCCCGCCCGCGTCCCTTTCTCCATAAAATTCTTCT
4 AGENOME-ZPMRG-F-V1.0   99.9%  99.8%       CATAGAAAAATCCACCCCTTACGAGTGCGGCTTCGACCCTATATCCCCCGCCCGCGTCCCTTTCTCCATAAAATTCTTCT
5 AGENOME-ZPMRG-Z-V1.0   99.9%  99.7%       CTTAGAAAAATCCACCCCTTACGAGTGCGGCTTCGACCCTATATCCCCCGCCCGCGTCCCTTTCTCCATAAAATTCTTCT
6 AGENOME-ZPMRG-T-V1.0   99.9%  99.8%       CATAGAAAAATCCACCCCTTACGAGTGCGGCTTCGACCCTATATCCCCCGCCCGCGTCCCTTTCTCCATAAAATTCTTCT
7 AGENOME-ZPMRG-HV-V1.0  99.9%  99.8%       CATAGAAAAATCCACCCCTTACGAGTGCGGCTTCGACCCTATATCCCCCGCCCGCGTCCCTTTCTCCATAAAATTCTTCT
8 AGENOME-ZPMRG-A-V1.0   99.9%  99.8%       CATAGAAAAATCCACCCCTTACGAGTGCGGCTTCGACCCTATATCCCCCGCCCGCGTCCCTTTCTCCATAAAATTCTTCT
9 NC_012920.1            99.9%  99.8%       CATAGAAAAATCCACCCCTTACGAGTGCGGCTTCGACCCTATATCCCCCGCCCGCGTCCCTTTCTCCATAAAATTCTTCT
  consensus/100%                            CsTAGAAAAATCCACCCCTTACGAGTGCGGCTTCGACCCTATATCCCCCGCCCGCGTCCCTTTCTCCATAAAATTCTTCT
  consensus/90%                             CsTAGAAAAATCCACCCCTTACGAGTGCGGCTTCGACCCTATATCCCCCGCCCGCGTCCCTTTCTCCATAAAATTCTTCT
  consensus/80%                             CATAGAAAAATCCACCCCTTACGAGTGCGGCTTCGACCCTATATCCCCCGCCCGCGTCCCTTTCTCCATAAAATTCTTCT
  consensus/70%                             CATAGAAAAATCCACCCCTTACGAGTGCGGCTTCGACCCTATATCCCCCGCCCGCGTCCCTTTCTCCATAAAATTCTTCT

                         cov    pid 10241          :         .         .         .         .         3         .         . 10320
1 AGENOME-ZPMRG-U-V1.0  100.0% 100.0%       TAGTAGCTATTACCTTCTTATTATTTGATCTAGAAATTGCCCTCCTTTTACCCCTACCATGAGCCCTACAAACAACTAAC
2 AGENOME-ZPMCG-V1.0    100.0%  99.9%       TAGTAGCTATTACCTTCTTATTATTTGATCTAGAAATTGCCCTCCTTTTACCCCTACCATGAGCCCTACAAACAACTAAC
3 AGENOME-ZPMRG-M-V1.0  100.0%  99.8%       TAGTAGCTATTACCTTCTTATTATTTGATCTAGAAATTGCCCTCCTTTTACCCCTACCATGAGCCCTACAAACAACTAAC
4 AGENOME-ZPMRG-F-V1.0   99.9%  99.8%       TAGTAGCTATTACCTTCTTATTATTTGATCTAGAAATTGCCCTCCTTTTACCCCTACCATGAGCCCTACAAACAACTAAC
5 AGENOME-ZPMRG-Z-V1.0   99.9%  99.7%       TAGTAGCTATTACCTTCTTATTATTTGATCTAGAAATTGCCCTCCTTTTACCCCTACCATGAGCCCTACAAACAACTAAC
6 AGENOME-ZPMRG-T-V1.0   99.9%  99.8%       TAGTAGCTATTACCTTCTTATTATTTGATCTAGAAATTGCCCTCCTTTTACCCCTACCATGAGCCCTACAAACAACTAAC
7 AGENOME-ZPMRG-HV-V1.0  99.9%  99.8%       TAGTAGCTATTACCTTCTTATTATTTGATCTAGAAATTGCCCTCCTTTTACCCCTACCATGAGCCCTACAAACAACTAAC
8 AGENOME-ZPMRG-A-V1.0   99.9%  99.8%       TAGTAGCTATTACCTTCTTATTATTTGATCTAGAAATTGCCCTCCTTTTACCCCTACCATGAGCCCTACAAACAACTAAC
9 NC_012920.1            99.9%  99.8%       TAGTAGCTATTACCTTCTTATTATTTGATCTAGAAATTGCCCTCCTTTTACCCCTACCATGAGCCCTACAAACAACTAAC
  consensus/100%                            TAGTAGCTATTACCTTCTTATTATTTGATCTAGAAATTGCCCTCCTTTTACCCCTACCATGAGCCCTACAAACAACTAAC
  consensus/90%                             TAGTAGCTATTACCTTCTTATTATTTGATCTAGAAATTGCCCTCCTTTTACCCCTACCATGAGCCCTACAAACAACTAAC
  consensus/80%                             TAGTAGCTATTACCTTCTTATTATTTGATCTAGAAATTGCCCTCCTTTTACCCCTACCATGAGCCCTACAAACAACTAAC
  consensus/70%                             TAGTAGCTATTACCTTCTTATTATTTGATCTAGAAATTGCCCTCCTTTTACCCCTACCATGAGCCCTACAAACAACTAAC

                         cov    pid 10321          .         .         :         .         .         .         .         4 10400
1 AGENOME-ZPMRG-U-V1.0  100.0% 100.0%       CTGCCACTAATAGTTATGTCATCCCTCTTATTAATCATCATCCTAGCCCTAAGTCTGGCCTATGAGTGACTACAAAAAGG
2 AGENOME-ZPMCG-V1.0    100.0%  99.9%       CTGCCACTAATAGTTATGTCATCCCTCTTATTAATCATCATCCTAGCCCTAAGTCTGGCCTATGAGTGACTACAAAAAGG
3 AGENOME-ZPMRG-M-V1.0  100.0%  99.8%       CTGCCACTAATAGTTATGTCATCCCTCTTATTAATCATCATCCTAGCCCTAAGTCTGGCCTATGAGTGACTACAAAAAGG
4 AGENOME-ZPMRG-F-V1.0   99.9%  99.8%       CTACCACTAATAGTTATGTCATCCCTCTTATTAATCATCATCCTAGCCCTAAGTCTGGCCTATGAGTGACTACAAAAAGG
5 AGENOME-ZPMRG-Z-V1.0   99.9%  99.7%       CTGCCACTAATAGTTATGTCATCCCTCTTATTAATCATCATCCTAGCCCTAAGTCTGGCCTATGAGTGACTACAAAAAGG
6 AGENOME-ZPMRG-T-V1.0   99.9%  99.8%       CTGCCACTAATAGTTATGTCATCCCTCTTATTAATCATCATCCTAGCCCTAAGTCTGGCCTATGAGTGACTACAAAAAGG
7 AGENOME-ZPMRG-HV-V1.0  99.9%  99.8%       CTGCCACTAATAGTTATGTCATCCCTCTTATTAATCATCATCCTAGCCCTAAGTCTGGCCTATGAGTGACTACAAAAAGG
8 AGENOME-ZPMRG-A-V1.0   99.9%  99.8%       CTGCCACTAATAGTTATGTCATCCCTCTTATTAATCATCATCCTAGCCCTAAGTCTGGCCTATGAGTGACTACAAAAAGG
9 NC_012920.1            99.9%  99.8%       CTGCCACTAATAGTTATGTCATCCCTCTTATTAATCATCATCCTAGCCCTAAGTCTGGCCTATGAGTGACTACAAAAAGG
  consensus/100%                            CTuCCACTAATAGTTATGTCATCCCTCTTATTAATCATCATCCTAGCCCTAAGTCTGGCCTATGAGTGACTACAAAAAGG
  consensus/90%                             CTuCCACTAATAGTTATGTCATCCCTCTTATTAATCATCATCCTAGCCCTAAGTCTGGCCTATGAGTGACTACAAAAAGG
  consensus/80%                             CTGCCACTAATAGTTATGTCATCCCTCTTATTAATCATCATCCTAGCCCTAAGTCTGGCCTATGAGTGACTACAAAAAGG
  consensus/70%                             CTGCCACTAATAGTTATGTCATCCCTCTTATTAATCATCATCCTAGCCCTAAGTCTGGCCTATGAGTGACTACAAAAAGG

                         cov    pid 10401          .         .         .         .         :         .         .         . 10480
1 AGENOME-ZPMRG-U-V1.0  100.0% 100.0%       ATTAGACTGAACCGAATTGGTATATAGTTTAAACAAAACGAATGATTTCGACTCATTAAATTATGATAATCATATTTACC
2 AGENOME-ZPMCG-V1.0    100.0%  99.9%       ATTAGACTGAGCTGAATTGGTATATAGTTTAAACAAAACGAATGATTTCGACTCATTAAATTATGATAATCATATTTACC
3 AGENOME-ZPMRG-M-V1.0  100.0%  99.8%       ATTAGACTGAGCTGAATTGGTATATAGTTTAAACAAAACGAATGATTTCGACTCATTAAATTATGATAATCATATTTACC
4 AGENOME-ZPMRG-F-V1.0   99.9%  99.8%       ATTAGACTGAACCGAATTGGTATATAGTTTAAACAAAACGAATGATTTCGACTCATTAAATTATGATAATCATATTTACC
5 AGENOME-ZPMRG-Z-V1.0   99.9%  99.7%       ATTAGACTGAGCTGAATTGGTATATAGTTTAAACAAAACGAATGATTTCGACTCATTAAATTATGATAATCATATTTACC
6 AGENOME-ZPMRG-T-V1.0   99.9%  99.8%       ATTAGACTGAACCGAATTGGTATATAGTTTAAACAAAACGAATGATTTCGACTCATTAAATTATGATAATCATATCTACC
7 AGENOME-ZPMRG-HV-V1.0  99.9%  99.8%       ATTAGACTGAACCGAATTGGTACATAGTTTAAACAAAACGAATGATTTCGACTCATTAAATTATGATAATCATATTTACC
8 AGENOME-ZPMRG-A-V1.0   99.9%  99.8%       ATTAGACTGAACCGAATTGGTATATAGTTTAAACAAAACGAATGATTTCGACTCATTAAATTATGATAATCATATTTACC
9 NC_012920.1            99.9%  99.8%       ATTAGACTGAACCGAATTGGTATATAGTTTAAACAAAACGAATGATTTCGACTCATTAAATTATGATAATCATATTTACC
  consensus/100%                            ATTAGACTGAuCsGAATTGGTAsATAGTTTAAACAAAACGAATGATTTCGACTCATTAAATTATGATAATCATATsTACC
  consensus/90%                             ATTAGACTGAuCsGAATTGGTAsATAGTTTAAACAAAACGAATGATTTCGACTCATTAAATTATGATAATCATATsTACC
  consensus/80%                             ATTAGACTGAuCsGAATTGGTATATAGTTTAAACAAAACGAATGATTTCGACTCATTAAATTATGATAATCATATTTACC
  consensus/70%                             ATTAGACTGAuCsGAATTGGTATATAGTTTAAACAAAACGAATGATTTCGACTCATTAAATTATGATAATCATATTTACC

                         cov    pid 10481          .         5         .         .         .         .         :         . 10560
1 AGENOME-ZPMRG-U-V1.0  100.0% 100.0%       AAATGCCCCTCATTTACATAAATATTATACTAGCATTTACCATCTCACTTCTAGGAATACTAGTATATCGCTCACACCTC
2 AGENOME-ZPMCG-V1.0    100.0%  99.9%       AAATGCCCCTCATTTACATAAATATTATACTAGCATTTACCATCTCACTTCTAGGAATACTAGTATATCGCTCACACCTC
3 AGENOME-ZPMRG-M-V1.0  100.0%  99.8%       AAATGCCCCTCATTTACATAAATATTATACTAGCATTTACCATCTCACTTCTAGGAATACTAGTATATCGCTCACACCTC
4 AGENOME-ZPMRG-F-V1.0   99.9%  99.8%       AAATGCCCCTCATTTACATAAATATTATACTAGCATTTACCATCTCACTTCTAGGAATACTAGTATATCGCTCACACCTC
5 AGENOME-ZPMRG-Z-V1.0   99.9%  99.7%       AAATGCCCCTCATTTACATAAATATTATACTAGCATTTACCATCTCACTTCTAGGAATACTAGTATATCGCTCACACCTC
6 AGENOME-ZPMRG-T-V1.0   99.9%  99.8%       AAATGCCCCTCATTTACATAAATATTATACTAGCATTTACCATCTCACTTCTAGGAATACTAGTATATCGCTCACACCTC
7 AGENOME-ZPMRG-HV-V1.0  99.9%  99.8%       AAATGCCCCTCATTTACATAAATATTATACTAGCATTTACCATCTCACTTCTAGGAATACTAGTATATCGCTCACACCTC
8 AGENOME-ZPMRG-A-V1.0   99.9%  99.8%       AAATGCCCCTCATTTACATAAATATTATACTAGCATTTACCATCTCACTTCTAGGAATACTAGTATATCGCTCACACCTC
9 NC_012920.1            99.9%  99.8%       AAATGCCCCTCATTTACATAAATATTATACTAGCATTTACCATCTCACTTCTAGGAATACTAGTATATCGCTCACACCTC
  consensus/100%                            AAATGCCCCTCATTTACATAAATATTATACTAGCATTTACCATCTCACTTCTAGGAATACTAGTATATCGCTCACACCTC
  consensus/90%                             AAATGCCCCTCATTTACATAAATATTATACTAGCATTTACCATCTCACTTCTAGGAATACTAGTATATCGCTCACACCTC
  consensus/80%                             AAATGCCCCTCATTTACATAAATATTATACTAGCATTTACCATCTCACTTCTAGGAATACTAGTATATCGCTCACACCTC
  consensus/70%                             AAATGCCCCTCATTTACATAAATATTATACTAGCATTTACCATCTCACTTCTAGGAATACTAGTATATCGCTCACACCTC

                         cov    pid 10561          .         .         .         6         .         .         .         . 10640
1 AGENOME-ZPMRG-U-V1.0  100.0% 100.0%       ATATCCTCCCTACTATGCCTAGAAGGAATAATACTATCGCTGTTCATTATAGCTACTCTCATAACCCTCAACACCCACTC
2 AGENOME-ZPMCG-V1.0    100.0%  99.9%       ATATCCTCCCTACTATGCCTAGAAGGAATAATACTATCGCTGTTCATTATAGCTACTCTCATAACCCTCAACACCCACTC
3 AGENOME-ZPMRG-M-V1.0  100.0%  99.8%       ATATCCTCCCTACTATGCCTAGAAGGAATAATACTATCGCTGTTCATTATAGCTACTCTCATAACCCTCAACACCCACTC
4 AGENOME-ZPMRG-F-V1.0   99.9%  99.8%       ATATCCTCCCTACTATGCCTAGAAGGAATAATACTATCGCTGTTCATTATAGCTACTCTCACAACCCTCAACACCCACTC
5 AGENOME-ZPMRG-Z-V1.0   99.9%  99.7%       ATATCCTCTCTACTATGCCTAGAAGGAATAATACTATCGCTGTTCATTATAGCTACTCTCATAACCCTCAACACCCACTC
6 AGENOME-ZPMRG-T-V1.0   99.9%  99.8%       ATATCCTCCCTACTATGCCTAGAAGGAATAATACTATCGCTGTTCATTATAGCTACTCTCATAACCCTCAACACCCACTC
7 AGENOME-ZPMRG-HV-V1.0  99.9%  99.8%       ATATCCTCCCTACTATGCCTAGAAGGAATAATACTATCGCTGTTCATTATAGCTACTCTCATAACCCTCAACACCCACTC
8 AGENOME-ZPMRG-A-V1.0   99.9%  99.8%       ATATCCTCCCTACTATGCCTAGAAGGAATAATACTATCGCTGTTCATTATAGCTACTCTCATAACCCTCAACACCCACTC
9 NC_012920.1            99.9%  99.8%       ATATCCTCCCTACTATGCCTAGAAGGAATAATACTATCGCTGTTCATTATAGCTACTCTCATAACCCTCAACACCCACTC
  consensus/100%                            ATATCCTCsCTACTATGCCTAGAAGGAATAATACTATCGCTGTTCATTATAGCTACTCTCAsAACCCTCAACACCCACTC
  consensus/90%                             ATATCCTCsCTACTATGCCTAGAAGGAATAATACTATCGCTGTTCATTATAGCTACTCTCAsAACCCTCAACACCCACTC
  consensus/80%                             ATATCCTCCCTACTATGCCTAGAAGGAATAATACTATCGCTGTTCATTATAGCTACTCTCATAACCCTCAACACCCACTC
  consensus/70%                             ATATCCTCCCTACTATGCCTAGAAGGAATAATACTATCGCTGTTCATTATAGCTACTCTCATAACCCTCAACACCCACTC

                         cov    pid 10641          :         .         .         .         .         7         .         . 10720
```

```
 1 AGENOME-ZPMRG-U-V1.0  100.0% 100.0%      CCTCTTAGCCAATATTGTGCCTATTGCCATACTAGTCTTTGCCGCCTGCGAAGCAGCGGTGGGCCTAGCCCTACTAGTCT
 2 AGENOME-ZPMCG-V1.0    100.0%  99.9%      CCTCTTAGCCAATATTGTGCCTATTGCCATACTAGTCTTTGCCGCCTGCGAAGCAGCGGTGGGCCTAGCCCTACTAGTCT
 3 AGENOME-ZPMRG-M-V1.0  100.0%  99.8%      CCTCTTAGCCAATATTGTGCCTATTGCCATACTAGTCTTTGCCGCCTGCGAAGCAGCGGTGGGCCTAGCCCTACTAGTCT
 4 AGENOME-ZPMRG-F-V1.0   99.9%  99.8%      CCTCTTAGCCAATATTGTGCCTATTGCCATACTAGTCTTTGCCGCCTGCGAAGCAGCGGTGGGCCTAGCCCTACTAGTCT
 5 AGENOME-ZPMRG-Z-V1.0   99.9%  99.7%      CCTCTTAGCCAATATTGTGCCTATTGCCATACTAGTCTTTGCCGCCTGCGAAGCAGCGGTGGGCCTAGCCCTACTAGTCT
 6 AGENOME-ZPMRG-T-V1.0   99.9%  99.8%      CCTCTTAGCCAATATTGTGCCTATTGCCATACTAGTCTTTGCCGCCTGCGAAGCAGCGGTGGGCCTAGCCCTACTAGTCT
 7 AGENOME-ZPMRG-HV-V1.0  99.9%  99.8%      CCTCTTAGCCAATATTGTGCCTATTGCCATACTAGTCTTTGCCGCCTGCGAAGCAGCGGTGGGCCTAGCCCTACTAGTCT
 8 AGENOME-ZPMRG-A-V1.0   99.9%  99.8%      CCTCTTAGCCAATATTGTGCCTATTGCCATACTAGTCTTTGCCGCCTGCGAAGCAGCGGTGGGCCTAGCCCTACTAGTCT
 9 NC_012920.1            99.9%  99.8%      CCTCTTAGCCAATATTGTGCCTATTGCCATACTAGTCTTTGCCGCCTGCGAAGCAGCGGTGGGCCTAGCCCTACTAGTCT
   consensus/100%                           CCTCTTAGCCAATATTGTGCCTATTGCCATACTAGTCTTTGCCGCCTGCGAAGCAGCGGTGGGCCTAGCCCTACTAGTCT
   consensus/90%                            CCTCTTAGCCAATATTGTGCCTATTGCCATACTAGTCTTTGCCGCCTGCGAAGCAGCGGTGGGCCTAGCCCTACTAGTCT
   consensus/80%                            CCTCTTAGCCAATATTGTGCCTATTGCCATACTAGTCTTTGCCGCCTGCGAAGCAGCGGTGGGCCTAGCCCTACTAGTCT
   consensus/70%                            CCTCTTAGCCAATATTGTGCCTATTGCCATACTAGTCTTTGCCGCCTGCGAAGCAGCGGTGGGCCTAGCCCTACTAGTCT

                          cov    pid 10721          .          .          :          .          .          .          .          8 10800
 1 AGENOME-ZPMRG-U-V1.0  100.0% 100.0%      CAATCTCCAACACATATGGCCTAGACTACGTACATAACCTAAACCTACTCCAATGCTAAAACTAATCGTCCCAACAATTA
 2 AGENOME-ZPMCG-V1.0    100.0%  99.9%      CAATCTCCAACACATATGGCCTAGACTACGTACATAACCTAAACCTACTCCAATGCTAAAACTAATCGTCCCAACAATTA
 3 AGENOME-ZPMRG-M-V1.0  100.0%  99.8%      CAATCTCCAACACATATGGCCTAGACTACGTACATAACCTAAACCTACTCCAATGCTAAAACTAATCGTCCCAACAATTA
 4 AGENOME-ZPMRG-F-V1.0   99.9%  99.8%      CAATCTCCAACACATATGGCCTAGACTACGTACATAACCTAAACCTACTCCAATGCTAAAACTAATCGTCCCAACAATTA
 5 AGENOME-ZPMRG-Z-V1.0   99.9%  99.7%      CAATCTCCAACACATATGGCCTAGACTACGTACATAACCTAAACCTACTCCAATGCTAAAACTAATCGTCCCAACAATTA
 6 AGENOME-ZPMRG-T-V1.0   99.9%  99.8%      CAATCTCCAACACATATGGCCTAGACTACGTACATAACCTAAACCTACTCCAATGCTAAAACTAATCGTCCCAACAATTA
 7 AGENOME-ZPMRG-HV-V1.0  99.9%  99.8%      CAATCTCCAACACATATGGCCTAGACTACGTACATAACCTAAACCTACTCCAATGCTAAAACTAATCGTCCCAACAATTA
 8 AGENOME-ZPMRG-A-V1.0   99.9%  99.8%      CAATCTCCAACACATATGGCCTAGACTACGTACATAACCTAAACCTACTCCAATGCTAAAACTAATCGTCCCAACAATTA
 9 NC_012920.1            99.9%  99.8%      CAATCTCCAACACATATGGCCTAGACTACGTACATAACCTAAACCTACTCCAATGCTAAAACTAATCGTCCCAACAATTA
   consensus/100%                           CAATCTCCAACACATATGGCCTAGACTACGTACATAACCTAAACCTACTCCAATGCTAAAACTAATCGTCCCAACAATTA
   consensus/90%                            CAATCTCCAACACATATGGCCTAGACTACGTACATAACCTAAACCTACTCCAATGCTAAAACTAATCGTCCCAACAATTA
   consensus/80%                            CAATCTCCAACACATATGGCCTAGACTACGTACATAACCTAAACCTACTCCAATGCTAAAACTAATCGTCCCAACAATTA
   consensus/70%                            CAATCTCCAACACATATGGCCTAGACTACGTACATAACCTAAACCTACTCCAATGCTAAAACTAATCGTCCCAACAATTA

                          cov    pid 10801          .          .          .          .          :          .          .          . 10880
 1 AGENOME-ZPMRG-U-V1.0  100.0% 100.0%      TATTACTACCACTGACATGACTTTCCAAAAAACACATAATTTGAATCAACACAACCACCCACAGCCTAATTATTAGCATC
 2 AGENOME-ZPMCG-V1.0    100.0%  99.9%      TATTACTACCACTGACATGACTTTCCAAAAAACACATAATTTGAATCAACACAACCACCCACAGCCTAATTATTAGCATC
 3 AGENOME-ZPMRG-M-V1.0  100.0%  99.8%      TATTACTACCACTGACATGACTTTCCAAAAAACACATAATTTGAATCAACACAACCACCCACAGCCTAATTATTAGCATC
 4 AGENOME-ZPMRG-F-V1.0   99.9%  99.8%      TATTACTACCACTGACATGACTTTCCAAAAAACACATAATTTGAATCAACACAACCACCCACAGCCTAATTATTAGCATC
 5 AGENOME-ZPMRG-Z-V1.0   99.9%  99.7%      TATTACTACCACTGACATGACTTTCCAAAAAACACATAATTTGAATCAACACAACCACCCACAGCCTAATTATTAGCATC
 6 AGENOME-ZPMRG-T-V1.0   99.9%  99.8%      TATTACTACCACTGACATGACTTTCCAAAAAACACATAATTTGAATCAACACAACCACCCACAGCCTAATTATTAGCATC
 7 AGENOME-ZPMRG-HV-V1.0  99.9%  99.8%      TATTACTACCACTGACATGACTTTCCAAAAAACACATAATTTGAATCAACACAACCACCCACAGCCTAATTATTAGCATC
 8 AGENOME-ZPMRG-A-V1.0   99.9%  99.8%      TATTACTACCACTGACATGACTTTCCAAAAAACACATAATTTGAATCAACACAACCACCCACAGCCTAATTATTAGCATC
 9 NC_012920.1            99.9%  99.8%      TATTACTACCACTGACATGACTTTCCAAAAAACACATAATTTGAATCAACACAACCACCCACAGCCTAATTATTAGCATC
   consensus/100%                           TATTACTACCACTGACATGACTTTCCAAAAAACACATAATTTGAATCAACACAACCACCCACAGCCTAATTATTAGCATC
   consensus/90%                            TATTACTACCACTGACATGACTTTCCAAAAAACACATAATTTGAATCAACACAACCACCCACAGCCTAATTATTAGCATC
   consensus/80%                            TATTACTACCACTGACATGACTTTCCAAAAAACACATAATTTGAATCAACACAACCACCCACAGCCTAATTATTAGCATC
   consensus/70%                            TATTACTACCACTGACATGACTTTCCAAAAAACACATAATTTGAATCAACACAACCACCCACAGCCTAATTATTAGCATC

                          cov    pid 10881          .          9          .          .          .          .          :          . 10960
 1 AGENOME-ZPMRG-U-V1.0  100.0% 100.0%      ATCCCTCTACTATTTTTTAACCAAATCAACAACAACCTATTTAGCTGTTCCCCAACCTTTTCCTCCGACCCCCTAACAAC
 2 AGENOME-ZPMCG-V1.0    100.0%  99.9%      ATCCCCCTACTATTTTTTAACCAAATCAACAACAACCTATTTAGCTGTTCCCCAACCTTTTCCTCCGACCCCCTAACAAC
 3 AGENOME-ZPMRG-M-V1.0  100.0%  99.8%      ATCCCCCTACTATTTTTTAACCAAATCAACAACAACCTATTTAGCTGTTCCCCAACCTTTTCCTCCGACCCCCTAACAAC
 4 AGENOME-ZPMRG-F-V1.0   99.9%  99.8%      ATCCCTCTACTATTTTTTAACCAAATCAACAACAACCTATTTAGCTGTTCCCCAACCTTTTCCTCCGACCCCCTAACAAC
 5 AGENOME-ZPMRG-Z-V1.0   99.9%  99.7%      ATCCCCCTACTATTTTTTAACCAAATCAACAACAACCTATTTAGCTGTTCCCCAACCTTTTCCTCCGACCCCCTAACAAC
 6 AGENOME-ZPMRG-T-V1.0   99.9%  99.8%      ATCCCTCTACTATTTTTTAACCAAATCAACAACAACCTATTTAGCTGTTCCCCAACCTTTTCCTCCGACCCCCTAACAAC
 7 AGENOME-ZPMRG-HV-V1.0  99.9%  99.8%      ATCCCTCTACTATTTTTTAACCAAATCAACAACAACCTATTTAGCTGTTCCCCAACCTTTTCCTCCGACCCCCTAACAAC
 8 AGENOME-ZPMRG-A-V1.0   99.9%  99.8%      ATCCCTCTACTATTTTTTAACCAAATCAACAACAACCTATTTAGCTGTTCCCCAACCTTTTCCTCCGACCCCCTAACAAC
 9 NC_012920.1            99.9%  99.8%      ATCCCTCTACTATTTTTTAACCAAATCAACAACAACCTATTTAGCTGTTCCCCAACCTTTTCCTCCGACCCCCTAACAAC
   consensus/100%                           ATCCCsCTACTATTTTTTAACCAAATCAACAACAACCTATTTAGCTGTTCCCCAACCTTTTCCTCCGACCCCCTAACAAC
   consensus/90%                            ATCCCsCTACTATTTTTTAACCAAATCAACAACAACCTATTTAGCTGTTCCCCAACCTTTTCCTCCGACCCCCTAACAAC
   consensus/80%                            ATCCCsCTACTATTTTTTAACCAAATCAACAACAACCTATTTAGCTGTTCCCCAACCTTTTCCTCCGACCCCCTAACAAC
   consensus/70%                            ATCCCsCTACTATTTTTTAACCAAATCAACAACAACCTATTTAGCTGTTCCCCAACCTTTTCCTCCGACCCCCTAACAAC

                          cov    pid 10961          .          .          .          0          .          .          .          . 11040
 1 AGENOME-ZPMRG-U-V1.0  100.0% 100.0%      CCCCCTCCTAATACTAACTACCTGACTCCTACCCCTCACAATCATGGCAAGCCAACGCCACTTATCCAGTGAACCACTAT
 2 AGENOME-ZPMCG-V1.0    100.0%  99.9%      CCCCCTCCTAATACTAACTACCTGACTCCTACCCCTCACAATCATGGCAAGCCAACGCCACTTATCCAGTGAACCACTAT
 3 AGENOME-ZPMRG-M-V1.0  100.0%  99.8%      CCCCCTCCTAATACTAACTACCTGACTCCTACCCCTCACAATCATGGCAAGCCAACGCCACTTATCCAGTGAACCACTAT
 4 AGENOME-ZPMRG-F-V1.0   99.9%  99.8%      CCCCCTCCTAATACTAACTACCTGACTCCTACCCCTCACAATCATGGCAAGCCAACGCCACTTATCCAGTGAACCACTAT
 5 AGENOME-ZPMRG-Z-V1.0   99.9%  99.7%      CCCCCTCCTAATACTAACTACCTGACTCCTACCCCTCACAATCATGGCAAGCCAACGCCACTTATCCAGTGAACCACTAT
 6 AGENOME-ZPMRG-T-V1.0   99.9%  99.8%      CCCCCTCCTAATACTAACTACCTGACTCCTACCCCTCACAATCATGGCAAGCCAACGCCACTTATCCAGTGAACCACTAT
 7 AGENOME-ZPMRG-HV-V1.0  99.9%  99.8%      CCCCCTCCTAATACTAACTACCTGACTCCTACCCCTCACAATCATGGCAAGCCAACGCCACTTATCCAATGAACCACTAT
 8 AGENOME-ZPMRG-A-V1.0   99.9%  99.8%      CCCCCTCCTAATACTAACTACCTGACTCCTACCCCTCACAATCATGGCAAGCCAACGCCACTTATCCAGTGAACCACTAT
 9 NC_012920.1            99.9%  99.8%      CCCCCTCCTAATACTAACTACCTGACTCCTACCCCTCACAATCATGGCAAGCCAACGCCACTTATCCAGTGAACCACTAT
   consensus/100%                           CCCCCTCCTAATACTAACTACCTGACTCCTACCCCTCACAATCATGGCAAGCCAACGCCACTTATCCAuTGAACCACTAT
   consensus/90%                            CCCCCTCCTAATACTAACTACCTGACTCCTACCCCTCACAATCATGGCAAGCCAACGCCACTTATCCAuTGAACCACTAT
   consensus/80%                            CCCCCTCCTAATACTAACTACCTGACTCCTACCCCTCACAATCATGGCAAGCCAACGCCACTTATCCAGTGAACCACTAT
   consensus/70%                            CCCCCTCCTAATACTAACTACCTGACTCCTACCCCTCACAATCATGGCAAGCCAACGCCACTTATCCAGTGAACCACTAT

                          cov    pid 11041          :          .          .          .          .          1          .          . 11120
 1 AGENOME-ZPMRG-U-V1.0  100.0% 100.0%      CACGAAAAAAACTCTACCTCTCTATACTAATCTCCCTACAAATCTCCTTAATTATAACATTCACAGCCACAGAACTAATC
 2 AGENOME-ZPMCG-V1.0    100.0%  99.9%      CACGAAAAAAACTCTACCTCTCTATACTAATCTCCCTACAAATCTCCTTAATTATAACATTCACAGCCACAGAACTAATC
 3 AGENOME-ZPMRG-M-V1.0  100.0%  99.8%      CACGAAAAAAACTCTACCTCTCTATACTAATCTCCCTACAAATCTCCTTAATTATAACATTCACAGCCACAGAACTAATC
 4 AGENOME-ZPMRG-F-V1.0   99.9%  99.8%      CACGAAAAAAACTCTACCTCTCTATACTAATCTCCCTACAAATCTCCTTAATTATAACATTCACAGCCACAGAACTAATC
 5 AGENOME-ZPMRG-Z-V1.0   99.9%  99.7%      CACGAAAAAAACTCTACCTCTCTATACTAATCTCCCTACAAATCTCCTTAATTATAACATTCACAGCCACAGAACTAATC
 6 AGENOME-ZPMRG-T-V1.0   99.9%  99.8%      CACGAAAAAAACTCTACCTCTCTATACTAATCTCCCTACAAATCTCCTTAATTATAACATTCACAGCCACAGAACTAATC
 7 AGENOME-ZPMRG-HV-V1.0  99.9%  99.8%      CACGAAAAAAACTCTACCTCTCTATACTAATCTCCCTACAAATCTCCTTAATTATAACATTCACAGCCACAGAACTAATC
 8 AGENOME-ZPMRG-A-V1.0   99.9%  99.8%      CACGAAAAAAACTCTACCTCTCTATACTAATCTCCCTACAAATCTCCTTAATTATAACATTCACAGCCACAGAACTAATC
 9 NC_012920.1            99.9%  99.8%      CACGAAAAAAACTCTACCTCTCTATACTAATCTCCCTACAAATCTCCTTAATTATAACATTCACAGCCACAGAACTAATC
   consensus/100%                           CACGAAAAAAACTCTACCTCTCTATACTAATCTCCCTACAAATCTCCTTAATTATAACATTCACAGCCACAGAACTAATC
   consensus/90%                            CACGAAAAAAACTCTACCTCTCTATACTAATCTCCCTACAAATCTCCTTAATTATAACATTCACAGCCACAGAACTAATC
   consensus/80%                            CACGAAAAAAACTCTACCTCTCTATACTAATCTCCCTACAAATCTCCTTAATTATAACATTCACAGCCACAGAACTAATC
   consensus/70%                            CACGAAAAAAACTCTACCTCTCTATACTAATCTCCCTACAAATCTCCTTAATTATAACATTCACAGCCACAGAACTAATC

                          cov    pid 11121          .          .          :          .          .          .          .          2 11200
 1 AGENOME-ZPMRG-U-V1.0  100.0% 100.0%      ATATTTTATATCTTCTTCGAAACCACACTTATCCCCACCTTGGCTATCATCACCCGATGAGGCAACCAGCCAGAACGCCT
```

```
  2 AGENOME-ZPMCG-V1.0    100.0%  99.9%       ATATTTTATATCTTCTTCGAAACCACACTTATCCCCACCTTGGCTATCATCACCCGATGAGGCAACCAGCCAGAACGCCT
  3 AGENOME-ZPMRG-M-V1.0  100.0%  99.8%       ATATTTTATATCTTCTTCGAAACCACACTTATCCCCACCTTGGCTATCATCACCCGATGAGGCAACCAGCCAGAACGCCT
  4 AGENOME-ZPMRG-F-V1.0   99.9%  99.8%       ATATTTTATATCTTCTTCGAAACCACACTTATCCCCACCTTGGCTATCATCACCCGATGAGGCAACCAGCCAGAACGCCT
  5 AGENOME-ZPMRG-Z-V1.0   99.9%  99.7%       ATATTTTATATCTTCTTCGAAACCACACTTATCCCCACCTTGGCTATCATCACCCGATGAGGCAACCAGCCAGAACGCCT
  6 AGENOME-ZPMRG-T-V1.0   99.9%  99.8%       ATATTTTATATCTTCTTCGAAACCACACTTATCCCCACCTTGGCTATCATCACCCGATGAGGCAACCAGCCAGAACGCCT
  7 AGENOME-ZPMRG-HV-V1.0  99.9%  99.8%       ATATTTTATATCTTCTTCGAAACCACACTTATCCCCACCTTGGCTATCATCACCCGATGAGGCAACCAGCCAGAACGCCT
  8 AGENOME-ZPMRG-A-V1.0   99.9%  99.8%       ATATTTTATATCTTCTTCGAAACCACACTTATCCCCACCTTGGCTATCATCACCCGATGAGGCAACCAGCCAGAACGCCT
  9 NC_012920.1            99.9%  99.8%       ATATTTTATATCTTCTTCGAAACCACACTTATCCCCACCTTGGCTATCATCACCCGATGAGGCAACCAGCCAGAACGCCT
    consensus/100%                            ATATTTTATATCTTCTTCGAAACCACACTTATCCCCACCTTGGCTATCATCACCCGATGAGGCAACCAGCCAGAACGCCT
    consensus/90%                             ATATTTTATATCTTCTTCGAAACCACACTTATCCCCACCTTGGCTATCATCACCCGATGAGGCAACCAGCCAGAACGCCT
    consensus/80%                             ATATTTTATATCTTCTTCGAAACCACACTTATCCCCACCTTGGCTATCATCACCCGATGAGGCAACCAGCCAGAACGCCT
    consensus/70%                             ATATTTTATATCTTCTTCGAAACCACACTTATCCCCACCTTGGCTATCATCACCCGATGAGGCAACCAGCCAGAACGCCT

                             cov    pid 11201          .         .         .         .         :         .         .         . 11280
  1 AGENOME-ZPMRG-U-V1.0  100.0% 100.0%       GAACGCAGGCACATACTTCCTATTCTACACCCTAGTAGGCTCCCTTCCCCTACTCATCGCACTAATTTACACTCACAACA
  2 AGENOME-ZPMCG-V1.0    100.0%  99.9%       GAACGCAGGCACATACTTCCTATTCTACACCCTAGTAGGCTCCCTTCCCCTACTCATCGCACTAATTTACACTCACAACA
  3 AGENOME-ZPMRG-M-V1.0  100.0%  99.8%       GAACGCAGGCACATACTTCCTATTCTACACCCTAGTAGGCTCCCTTCCCCTACTCATCGCACTAATTTACACTCACAACA
  4 AGENOME-ZPMRG-F-V1.0   99.9%  99.8%       GAACGCAGGCACATACTTCCTATTCTACACCCTAGTAGGCTCCCTTCCCCTACTCATCGCACTAATTTACACTCACAACA
  5 AGENOME-ZPMRG-Z-V1.0   99.9%  99.7%       GAACGCAGGCACATACTTCCTATTCTACACCCTAGTAGGCTCCCTTCCCCTACTCATCGCACTAATTTACACTCACAACA
  6 AGENOME-ZPMRG-T-V1.0   99.9%  99.8%       GAACGCAGGCACATACTTCCTATTCTACACCCTAGTAGGCTCCCTTCCCCTACTCATCGCACTGATTTACACTCACAACA
  7 AGENOME-ZPMRG-HV-V1.0  99.9%  99.8%       GAACGCAGGCACATACTTCCTATTCTACACCCTAGTAGGCTCCCTTCCCCTACTCATCGCACTAATTTACACTCACAACA
  8 AGENOME-ZPMRG-A-V1.0   99.9%  99.8%       GAACGCAGGCACATACTTCCTATTCTACACCCTAGTAGGCTCCCTTCCCCTACTCATCGCACTAATTTACACTCACAACA
  9 NC_012920.1            99.9%  99.8%       GAACGCAGGCACATACTTCCTATTCTACACCCTAGTAGGCTCCCTTCCCCTACTCATCGCACTAATTTACACTCACAACA
    consensus/100%                            GAACGCAGGCACATACTTCCTATTCTACACCCTAGTAGGCTCCCTTCCCCTACTCATCGCACTuATTTACACTCACAACA
    consensus/90%                             GAACGCAGGCACATACTTCCTATTCTACACCCTAGTAGGCTCCCTTCCCCTACTCATCGCACTuATTTACACTCACAACA
    consensus/80%                             GAACGCAGGCACATACTTCCTATTCTACACCCTAGTAGGCTCCCTTCCCCTACTCATCGCACTAATTTACACTCACAACA
    consensus/70%                             GAACGCAGGCACATACTTCCTATTCTACACCCTAGTAGGCTCCCTTCCCCTACTCATCGCACTAATTTACACTCACAACA

                             cov    pid 11281          .         3         .         .         .         .         :         . 11360
  1 AGENOME-ZPMRG-U-V1.0  100.0% 100.0%       CCCTAGGCTCACTAAACATTCTACTACTCACTCTCACTGCCCAAGAACTATCAAACTCCTGAGCTAACAACTTAATATGA
  2 AGENOME-ZPMCG-V1.0    100.0%  99.9%       CCCTAGGCTCACTAAACATTCTACTACTCACTCTCACTGCCCAAGAACTATCAAACTCCTGAGCCAACAACTTAATATGA
  3 AGENOME-ZPMRG-M-V1.0  100.0%  99.8%       CCCTAGGCTCACTAAACATTCTACTACTCACTCTCACTGCCCAAGAACTATCAAACTCCTGAGCCAACAACTTAATATGA
  4 AGENOME-ZPMRG-F-V1.0   99.9%  99.8%       CCCTAGGCTCACTAAACATTCTACTACTCACTCTCACTGCCCAAGAACTATCAAACTCCTGAGCCAACAACTTAATATGA
  5 AGENOME-ZPMRG-Z-V1.0   99.9%  99.7%       CCCTAGGCTCACTAAACATTCTACTACTCACTCTCACTGCCCAAGAACTATCAAACTCCTGAGCCAACAACTTAATATGA
  6 AGENOME-ZPMRG-T-V1.0   99.9%  99.8%       CCCTAGGCTCACTAAACATTCTACTACTCACTCTCACTGCCCAAGAACTATCAAACTCCTGAGCCAACAACTTAATATGA
  7 AGENOME-ZPMRG-HV-V1.0  99.9%  99.8%       CCCTAGGCTCACTAAACATTCTACTACTCACTCTCACTGCCCAAGAACTATCAAACTCCTGAGCCAACAACTTAATATGA
  8 AGENOME-ZPMRG-A-V1.0   99.9%  99.8%       CCCTAGGCTCACTAAACATTCTACTACTCACTCTCACTGCCCAAGAACTATCAAACTCCTGAGCCAACAACTTAATATGA
  9 NC_012920.1            99.9%  99.8%       CCCTAGGCTCACTAAACATTCTACTACTCACTCTCACTGCCCAAGAACTATCAAACTCCTGAGCCAACAACTTAATATGA
    consensus/100%                            CCCTAGGCTCACTAAACATTCTACTACTCACTCTCACTGCCCAAGAACTATCAAACTCCTGAGCsAACAACTTAATATGA
    consensus/90%                             CCCTAGGCTCACTAAACATTCTACTACTCACTCTCACTGCCCAAGAACTATCAAACTCCTGAGCsAACAACTTAATATGA
    consensus/80%                             CCCTAGGCTCACTAAACATTCTACTACTCACTCTCACTGCCCAAGAACTATCAAACTCCTGAGCCAACAACTTAATATGA
    consensus/70%                             CCCTAGGCTCACTAAACATTCTACTACTCACTCTCACTGCCCAAGAACTATCAAACTCCTGAGCCAACAACTTAATATGA

                             cov    pid 11361          .         .         .         4         .         .         .         . 11440
  1 AGENOME-ZPMRG-U-V1.0  100.0% 100.0%       CTAGCTTACACAATAGCTTTTATAGTAAAGATACCTCTTTACGGACTCCACTTATGACTCCCTAAAGCCCATGTCGAAGC
  2 AGENOME-ZPMCG-V1.0    100.0%  99.9%       CTAGCTTACACAATAGCTTTTATAGTAAAGATACCTCTTTACGGACTCCACTTATGACTCCCTAAAGCCCATGTCGAAGC
  3 AGENOME-ZPMRG-M-V1.0  100.0%  99.8%       CTAGCTTACACAATAGCTTTTATAGTAAAGATACCTCTTTACGGACTCCACTTATGACTCCCTAAAGCCCATGTCGAAGC
  4 AGENOME-ZPMRG-F-V1.0   99.9%  99.8%       CTAGCTTACACAATAGCTTTTATAGTAAAGATACCTCTTTACGGACTCCACTTATGACTCCCTAAAGCCCATGTCGAAGC
  5 AGENOME-ZPMRG-Z-V1.0   99.9%  99.7%       CTAGCTTACACAATAGCTTTTATAGTAAAGATACCTCTTTACGGACTCCACTTATGACTCCCTAAAGCCCATGTCGAAGC
  6 AGENOME-ZPMRG-T-V1.0   99.9%  99.8%       CTAGCTTACACAATAGCTTTTATAGTAAAGATACCTCTTTACGGACTCCACTTATGACTCCCTAAAGCCCATGTCGAAGC
  7 AGENOME-ZPMRG-HV-V1.0  99.9%  99.8%       CTAGCTTACACAATAGCTTTTATAGTAAAGATACCTCTTTACGGACTCCACTTATGACTCCCTAAAGCCCATGTCGAAGC
  8 AGENOME-ZPMRG-A-V1.0   99.9%  99.8%       CTAGCTTACACAATAGCTTTTATAGTAAAGATACCTCTTTACGGACTCCACTTATGACTCCCTAAAGCCCATGTCGAAGC
  9 NC_012920.1            99.9%  99.8%       CTAGCTTACACAATAGCTTTTATAGTAAAGATACCTCTTTACGGACTCCACTTATGACTCCCTAAAGCCCATGTCGAAGC
    consensus/100%                            CTAGCTTACACAATAGCTTTTATAGTAAAGATACCTCTTTACGGACTCCACTTATGACTCCCTAAAGCCCATGTCGAAGC
    consensus/90%                             CTAGCTTACACAATAGCTTTTATAGTAAAGATACCTCTTTACGGACTCCACTTATGACTCCCTAAAGCCCATGTCGAAGC
    consensus/80%                             CTAGCTTACACAATAGCTTTTATAGTAAAGATACCTCTTTACGGACTCCACTTATGACTCCCTAAAGCCCATGTCGAAGC
    consensus/70%                             CTAGCTTACACAATAGCTTTTATAGTAAAGATACCTCTTTACGGACTCCACTTATGACTCCCTAAAGCCCATGTCGAAGC

                             cov    pid 11441          :         .         .         .         .         5         .         . 11520
  1 AGENOME-ZPMRG-U-V1.0  100.0% 100.0%       CCCCATCGCTGGGTCAATAGTACTTGCCGCAGTACTCTTGAAACTAGGCGGCTATGGTATAATACGCCTCACACTCATTC
  2 AGENOME-ZPMCG-V1.0    100.0%  99.9%       CCCCATCGCTGGGTCAATAGTACTTGCCGCAGTACTCTTAAAACTAGGCGGCTATGGTATAATACGCCTCACACTCATTC
  3 AGENOME-ZPMRG-M-V1.0  100.0%  99.8%       CCCCATCGCTGGGTCAATAGTACTTGCCGCAGTACTCTTAAAACTAGGCGGCTATGGTATAATACGCCTCACACTCATTC
  4 AGENOME-ZPMRG-F-V1.0   99.9%  99.8%       CCCCATCGCTGGGTCAATAGTACTTGCCGCAGTACTCTTAAAACTAGGCGGCTATGGTATAATACGCCTCACACTCATTC
  5 AGENOME-ZPMRG-Z-V1.0   99.9%  99.7%       CCCCATCGCTGGGTCAATAGTACTTGCCGCAGTACTCTTAAAACTAGGCGGCTATGGTATAATACGCCTCACACTCATTC
  6 AGENOME-ZPMRG-T-V1.0   99.9%  99.8%       CCCCATCGCTGGGTCAATAGTACTTGCCGCAGTACTCTTAAAACTAGGCGGCTATGGTATAATACGCCTCACACTCATTC
  7 AGENOME-ZPMRG-HV-V1.0  99.9%  99.8%       CCCCATCGCTGGGTCAATAGTACTTGCCGCAGTACTCTTAAAACTAGGCGGCTATGGTATAATACGCCTCACACTCATTC
  8 AGENOME-ZPMRG-A-V1.0   99.9%  99.8%       CCCCATCGCTGGGTCAATAGTACTTGCCGCAGTACTCTTAAAACTAGGCGGCTATGGTATAATACGCCTCACACTCATTC
  9 NC_012920.1            99.9%  99.8%       CCCCATCGCTGGGTCAATAGTACTTGCCGCAGTACTCTTAAAACTAGGCGGCTATGGTATAATACGCCTCACACTCATTC
    consensus/100%                            CCCCATCGCTGGGTCAATAGTACTTGCCGCAGTACTCTTuAAACTAGGCGGCTATGGTATAATACGCCTCACACTCATTC
    consensus/90%                             CCCCATCGCTGGGTCAATAGTACTTGCCGCAGTACTCTTuAAACTAGGCGGCTATGGTATAATACGCCTCACACTCATTC
    consensus/80%                             CCCCATCGCTGGGTCAATAGTACTTGCCGCAGTACTCTTAAAACTAGGCGGCTATGGTATAATACGCCTCACACTCATTC
    consensus/70%                             CCCCATCGCTGGGTCAATAGTACTTGCCGCAGTACTCTTAAAACTAGGCGGCTATGGTATAATACGCCTCACACTCATTC

                             cov    pid 11521          .         .         :         .         .         .         .         6 11600
  1 AGENOME-ZPMRG-U-V1.0  100.0% 100.0%       TCAACCCCCTGACAAAACACATAGCCTACCCCTTCCTTGTACTATCCCTATGAGGCATAATTATAACAAGCTCCATCTGC
  2 AGENOME-ZPMCG-V1.0    100.0%  99.9%       TCAACCCCCTGACAAAACACATAGCCTACCCCTTCCTTGTACTATCCCTATGAGGCATAATTATAACAAGCTCCATCTGC
  3 AGENOME-ZPMRG-M-V1.0  100.0%  99.8%       TCAACCCCCTGACAAAACACATAGCCTACCCCTTCCTTGTACTATCCCTATGAGGCATAATTATAACAAGCTCCATCTGC
  4 AGENOME-ZPMRG-F-V1.0   99.9%  99.8%       TCAACCCCCTGACAAAACACATAGCCTACCCCTTCCTTGTACTATCCCTATGAGGCATAATTATAACAAGCTCCATCTGC
  5 AGENOME-ZPMRG-Z-V1.0   99.9%  99.7%       TCAACCCCCTGACAAAACACATAGCCTACCCCTTCCTTGTACTATCCCTATGAGGCATAATTATAACAAGCTCCATCTGC
  6 AGENOME-ZPMRG-T-V1.0   99.9%  99.8%       TCAACCCCCTGACAAAACACATAGCCTACCCCTTCCTTGTACTATCCCTATGAGGCATAATTATAACAAGCTCCATCTGC
  7 AGENOME-ZPMRG-HV-V1.0  99.9%  99.8%       TCAACCCCCTGACAAAACACATAGCCTACCCCTTCCTTGTACTATCCCTATGAGGCATAATTATAACAAGCTCCATCTGC
  8 AGENOME-ZPMRG-A-V1.0   99.9%  99.8%       TCAACCCCCTGACAAAACACATAGCCTACCCCTTCCTTGTACTATCCCTATGAGGCATAATTATAACAAGCTCCATCTGC
  9 NC_012920.1            99.9%  99.8%       TCAACCCCCTGACAAAACACATAGCCTACCCCTTCCTTGTACTATCCCTATGAGGCATAATTATAACAAGCTCCATCTGC
    consensus/100%                            TCAACCCCCTGACAAAACACATAGCCTACCCCTTCCTTGTACTATCCCTATGAGGCATAATTATAACAAGCTCCATCTGC
    consensus/90%                             TCAACCCCCTGACAAAACACATAGCCTACCCCTTCCTTGTACTATCCCTATGAGGCATAATTATAACAAGCTCCATCTGC
    consensus/80%                             TCAACCCCCTGACAAAACACATAGCCTACCCCTTCCTTGTACTATCCCTATGAGGCATAATTATAACAAGCTCCATCTGC
    consensus/70%                             TCAACCCCCTGACAAAACACATAGCCTACCCCTTCCTTGTACTATCCCTATGAGGCATAATTATAACAAGCTCCATCTGC

                             cov    pid 11601          .         .         .         .         :         .         .         . 11680
  1 AGENOME-ZPMRG-U-V1.0  100.0% 100.0%       CTACGACAAACAGACCTAAAATCGCTCATTGCATACTCTTCAATCAGCCACATAGCCCTCGTAGTAACAGCCATTCTCAT
  2 AGENOME-ZPMCG-V1.0    100.0%  99.9%       CTACGACAAACAGACCTAAAATCGCTCATTGCATACTCTTCAATCAGCCACATAGCCCTCGTAGTAACAGCCATTCTCAT
```

```
  3 AGENOME-ZPMRG-M-V1.0  100.0%  99.8%       CTACGACAAACAGACCTAAAATCGCTCATTGCATACTCTTCAATCAGCCACATAGCCCTCGTAGTAACAGCCATTCTCAT
  4 AGENOME-ZPMRG-F-V1.0   99.9%  99.8%       CTACGACAAACAGACCTAAAATCGCTCATTGCATACTCTTCAATCAGCCACATAGCCCTCGTAGTAACAGCCATTCTCAT
  5 AGENOME-ZPMRG-Z-V1.0   99.9%  99.7%       CTACGACAAACAGACCTAAAATCGCTCATTGCATACTCTTCAATCAGCCACATAGCCCTCGTAGTAACAGCCATTCTCAT
  6 AGENOME-ZPMRG-T-V1.0   99.9%  99.8%       CTACGACAAACAGACCTAAAATCGCTCATTGCATACTCTTCAATCAGCCACATAGCCCTCGTAGTAACAGCCATTCTCAT
  7 AGENOME-ZPMRG-HV-V1.0  99.9%  99.8%       CTACGACAAACAGACCTAAAATCGCTCATTGCATACTCTTCAATCAGCCACATAGCCCTCGTAGTAACAGCCATTCTCAT
  8 AGENOME-ZPMRG-A-V1.0   99.9%  99.8%       CTACGACAAACAGACCTAAAATCGCTCATTGCATACTCTTCAATCAGCCACATAGCCCTCGTAGTAACAGCCATTCTCAT
  9 NC_012920.1            99.9%  99.8%       CTACGACAAACAGACCTAAAATCGCTCATTGCATACTCTTCAATCAGCCACATAGCCCTCGTAGTAACAGCCATTCTCAT
    consensus/100%                            CTACGACAAACAGACCTAAAATCGCTCATTGCATACTCTTCAATCAGCCACATAGCCCTCGTAGTAACAGCCATTCTCAT
    consensus/90%                             CTACGACAAACAGACCTAAAATCGCTCATTGCATACTCTTCAATCAGCCACATAGCCCTCGTAGTAACAGCCATTCTCAT
    consensus/80%                             CTACGACAAACAGACCTAAAATCGCTCATTGCATACTCTTCAATCAGCCACATAGCCCTCGTAGTAACAGCCATTCTCAT
    consensus/70%                             CTACGACAAACAGACCTAAAATCGCTCATTGCATACTCTTCAATCAGCCACATAGCCCTCGTAGTAACAGCCATTCTCAT

                          cov    pid 11681          .         7         .         .         .         .         :         . 11760
  1 AGENOME-ZPMRG-U-V1.0  100.0% 100.0%       CCAAACCCCCTGAAGCTTCACCGGCGCAGTCATTCTCATAATCGCCCACGGACTTACATCCTCATTACTATTCTGCCTAG
  2 AGENOME-ZPMCG-V1.0    100.0%  99.9%       CCAAACCCCCTGAAGCTTCACCGGCGCAGTCATTCTCATAATCGCCCACGGACTTACATCCTCATTACTATTCTGCCTAG
  3 AGENOME-ZPMRG-M-V1.0  100.0%  99.8%       CCAAACCCCCTGAAGCTTCACCGGCGCAGTCATTCTCATAATCGCCCACGGACTTACATCCTCATTACTATTCTGCCTAG
  4 AGENOME-ZPMRG-F-V1.0   99.9%  99.8%       CCAAACCCCCTGAAGCTTCACCGGCGCAGTCATTCTCATAATCGCCCACGGACTTACATCCTCATTACTATTCTGCCTAG
  5 AGENOME-ZPMRG-Z-V1.0   99.9%  99.7%       CCAAACCCCCTGAAGCTTCACCGGCGCAGTCATTCTCATAATCGCCCACGGACTTACATCCTCATTACTATTCTGCCTAG
  6 AGENOME-ZPMRG-T-V1.0   99.9%  99.8%       CCAAACCCCCTGAAGCTTCACCGGCGCAGTCATTCTCATAATCGCCCACGGACTTACATCCTCATTACTATTCTGCCTAG
  7 AGENOME-ZPMRG-HV-V1.0  99.9%  99.8%       CCAAACCCCCTGAAGCTTCACCGGCGCAGTCATTCTCATAATCGCCCACGGGCTTACATCCTCATTACTATTCTGCCTAG
  8 AGENOME-ZPMRG-A-V1.0   99.9%  99.8%       CCAAACCCCCTGAAGCTTCACCGGCGCAGTCATTCTCATAATCGCCCACGGGCTTACATCCTCATTACTATTCTGCCTAG
  9 NC_012920.1            99.9%  99.8%       CCAAACCCCCTGAAGCTTCACCGGCGCAGTCATTCTCATAATCGCCCACGGGCTTACATCCTCATTACTATTCTGCCTAG
    consensus/100%                            CCAAACCCCCTGAAGCTTCACCGGCGCAGTCATTCTCATAATCGCCCACGGuCTTACATCCTCATTACTATTCTGCCTAG
    consensus/90%                             CCAAACCCCCTGAAGCTTCACCGGCGCAGTCATTCTCATAATCGCCCACGGuCTTACATCCTCATTACTATTCTGCCTAG
    consensus/80%                             CCAAACCCCCTGAAGCTTCACCGGCGCAGTCATTCTCATAATCGCCCACGGuCTTACATCCTCATTACTATTCTGCCTAG
    consensus/70%                             CCAAACCCCCTGAAGCTTCACCGGCGCAGTCATTCTCATAATCGCCCACGGuCTTACATCCTCATTACTATTCTGCCTAG

                          cov    pid 11761          .         .         .         8         .         .         .         . 11840
  1 AGENOME-ZPMRG-U-V1.0  100.0% 100.0%       CAAACTCAAACTACGAACGCACTCACAGTCGCATCATAATCCTCTCTCAAGGACTTCAAACTCTACTCCCACTAATAGCT
  2 AGENOME-ZPMCG-V1.0    100.0%  99.9%       CAAACTCAAACTACGAACGCACTCACAGTCGCATCATAATCCTCTCTCAAGGACTTCAAACTCTACTCCCACTAATAGCT
  3 AGENOME-ZPMRG-M-V1.0  100.0%  99.8%       CAAACTCAAACTACGAACGCACTCACAGTCGCATCATAATCCTCTCTCAAGGACTTCAAACTCTACTCCCACTAATAGCT
  4 AGENOME-ZPMRG-F-V1.0   99.9%  99.8%       CAAACTCAAACTACGAACGCACTCACAGTCGCATCATAATCCTCTCTCAAGGACTTCAAACTCTACTCCCACTAATAGCT
  5 AGENOME-ZPMRG-Z-V1.0   99.9%  99.7%       CAAACTCAAACTACGAACGCACTCACAGTCGCATCATAATCCTCTCTCAAGGACTTCAAACTCTACTCCCACTAATAGCT
  6 AGENOME-ZPMRG-T-V1.0   99.9%  99.8%       CAAACTCAAACTACGAACGCACTCACAGTCGCATCATAATCCTCTCTCAAGGACTTCAAACTCTACTCCCACTAATAGCT
  7 AGENOME-ZPMRG-HV-V1.0  99.9%  99.8%       CAAACTCAAACTACGAACGCACTCACAGTCGCATCATAATCCTCTCTCAAGGACTTCAAACTCTACTCCCACTAATAGCT
  8 AGENOME-ZPMRG-A-V1.0   99.9%  99.8%       CAAACTCAAACTACGAACGCACTCACAGTCGCATCATAATCCTCTCTCAAGGACTTCAAACTCTACTCCCACTAATAGCT
  9 NC_012920.1            99.9%  99.8%       CAAACTCAAACTACGAACGCACTCACAGTCGCATCATAATCCTCTCTCAAGGACTTCAAACTCTACTCCCACTAATAGCT
    consensus/100%                            CAAACTCAAACTACGAACGCACTCACAGTCGCATCATAATCCTCTCTCAAGGACTTCAAACTCTACTCCCACTAATAGCT
    consensus/90%                             CAAACTCAAACTACGAACGCACTCACAGTCGCATCATAATCCTCTCTCAAGGACTTCAAACTCTACTCCCACTAATAGCT
    consensus/80%                             CAAACTCAAACTACGAACGCACTCACAGTCGCATCATAATCCTCTCTCAAGGACTTCAAACTCTACTCCCACTAATAGCT
    consensus/70%                             CAAACTCAAACTACGAACGCACTCACAGTCGCATCATAATCCTCTCTCAAGGACTTCAAACTCTACTCCCACTAATAGCT

                          cov    pid 11841          :         .         .         .         .         9         .         . 11920
  1 AGENOME-ZPMRG-U-V1.0  100.0% 100.0%       TTTTGATGACTTCTAGCAAGCCTCGCTAACCTCGCCTTACCCCCCACTATTAACCTACTGGGAGAACTCTCTGTGCTAGT
  2 AGENOME-ZPMCG-V1.0    100.0%  99.9%       TTTTGATGACTTCTAGCAAGCCTCGCTAACCTCGCCTTACCCCCCACTATTAACCTACTGGGAGAACTCTCTGTGCTAGT
  3 AGENOME-ZPMRG-M-V1.0  100.0%  99.8%       TTTTGATGACTTCTAGCAAGCCTCGCTAACCTCGCCTTACCCCCCACTATTAACCTACTGGGAGAACTCTCTGTGCTAGT
  4 AGENOME-ZPMRG-F-V1.0   99.9%  99.8%       TTTTGATGACTTCTAGCAAGCCTCGCTAACCTCGCCTTACCCCCCACTATTAACCTACTGGGAGAACTCTCTGTGCTAGT
  5 AGENOME-ZPMRG-Z-V1.0   99.9%  99.7%       TTTTGATGACTTCTAGCAAGCCTCGCTAACCTCGCCTTACCCCCCACTATTAACCTACTGGGAGAACTCTCTGTGCTAGT
  6 AGENOME-ZPMRG-T-V1.0   99.9%  99.8%       TTTTGATGACTTCTAGCAAGCCTCGCTAACCTCGCCTTACCCCCCACTATTAACCTACTGGGAGAACTCTCTGTGCTAGT
  7 AGENOME-ZPMRG-HV-V1.0  99.9%  99.8%       TTTTGATGACTTCTAGCAAGCCTCGCTAACCTCGCCTTACCCCCCACTATTAACCTACTGGGAGAACTCTCTGTGCTAGT
  8 AGENOME-ZPMRG-A-V1.0   99.9%  99.8%       TTTTGATGACTTCTAGCAAGCCTCGCTAACCTCGCCTTACCCCCCACTATTAATCTACTGGGAGAACTCTCTGTGCTAGT
  9 NC_012920.1            99.9%  99.8%       TTTTGATGACTTCTAGCAAGCCTCGCTAACCTCGCCTTACCCCCCACTATTAACCTACTGGGAGAACTCTCTGTGCTAGT
    consensus/100%                            TTTTGATGACTTCTAGCAAGCCTCGCTAACCTCGCCTTACCCCCCACTATTAAsCTACTGGGAGAACTCTCTGTGCTAGT
    consensus/90%                             TTTTGATGACTTCTAGCAAGCCTCGCTAACCTCGCCTTACCCCCCACTATTAAsCTACTGGGAGAACTCTCTGTGCTAGT
    consensus/80%                             TTTTGATGACTTCTAGCAAGCCTCGCTAACCTCGCCTTACCCCCCACTATTAACCTACTGGGAGAACTCTCTGTGCTAGT
    consensus/70%                             TTTTGATGACTTCTAGCAAGCCTCGCTAACCTCGCCTTACCCCCCACTATTAACCTACTGGGAGAACTCTCTGTGCTAGT

                          cov    pid 11921          .         .         :         .         .         .         .         0 12000
  1 AGENOME-ZPMRG-U-V1.0  100.0% 100.0%       AACCACGTTCTCCTGATCAAATATCACTCTCCTACTTACAGGACTCAACATACTAGTCACAGCCCTATACTCCCTCTACA
  2 AGENOME-ZPMCG-V1.0    100.0%  99.9%       AACCACGTTCTCCTGATCAAATATCACTCTCCTACTTACAGGACTCAACATACTAGTCACAGCCCTATACTCCCTCTACA
  3 AGENOME-ZPMRG-M-V1.0  100.0%  99.8%       AACCACGTTCTCCTGATCAAATATCACTCTCCTACTTACAGGACTCAACATACTAGTCACAGCCCTATACTCCCTCTACA
  4 AGENOME-ZPMRG-F-V1.0   99.9%  99.8%       AACCACGTTCTCCTGATCAAATATCACTCTCCTACTTACAGGACTCAACATACTAGTCACAGCCCTATACTCCCTCTACA
  5 AGENOME-ZPMRG-Z-V1.0   99.9%  99.7%       AACCACGTTCTCCTGATCAAATATCACTCTCCTACTTACAGGACTCAACATACTAGTCACAGCCCTATACTCCCTCTACA
  6 AGENOME-ZPMRG-T-V1.0   99.9%  99.8%       AACCACGTTCTCCTGATCAAATATCACTCTCCTACTTACAGGACTCAACATACTAGTCACAGCCCTATACTCCCTCTACA
  7 AGENOME-ZPMRG-HV-V1.0  99.9%  99.8%       AACCACGTTCTCCTGATCAAATATCACCCTCCTACTTACAGGACTCAACATACTAGTCACAGCCCTATACTCCCTCTACA
  8 AGENOME-ZPMRG-A-V1.0   99.9%  99.8%       AACCACGTTCTCCTGATCAAATATCACTCTCCTACTTACAGGACTCAACATACTAGTCACAGCCCTATACTCCCTCTACA
  9 NC_012920.1            99.9%  99.8%       AACCACGTTCTCCTGATCAAATATCACTCTCCTACTTACAGGACTCAACATACTAGTCACAGCCCTATACTCCCTCTACA
    consensus/100%                            AACCACGTTCTCCTGATCAAATATCACsCTCCTACTTACAGGACTCAACATACTAGTCACAGCCCTATACTCCCTCTACA
    consensus/90%                             AACCACGTTCTCCTGATCAAATATCACsCTCCTACTTACAGGACTCAACATACTAGTCACAGCCCTATACTCCCTCTACA
    consensus/80%                             AACCACGTTCTCCTGATCAAATATCACTCTCCTACTTACAGGACTCAACATACTAGTCACAGCCCTATACTCCCTCTACA
    consensus/70%                             AACCACGTTCTCCTGATCAAATATCACTCTCCTACTTACAGGACTCAACATACTAGTCACAGCCCTATACTCCCTCTACA

                          cov    pid 12001          .         .         .         .         :         .         .         . 12080
  1 AGENOME-ZPMRG-U-V1.0  100.0% 100.0%       TATTTACCACAACACAATGGGGCTCACTCACCCACCACATTAACAACATAAAACCCTCATTCACACGAGAAAACACCCTC
  2 AGENOME-ZPMCG-V1.0    100.0%  99.9%       TATTTACCACAACACAATGGGGCTCACTCACCCACCACATTAACAACATAAAACCCTCATTCACACGAGAAAACACCCTC
  3 AGENOME-ZPMRG-M-V1.0  100.0%  99.8%       TATTTACCACAACACAATGGGGCTCACTCACCCACCACATTAACAACATAAAACCCTCATTCACACGAGAAAACACCCTC
  4 AGENOME-ZPMRG-F-V1.0   99.9%  99.8%       TATTTACCACAACACAATGGGGCTCACTCACCCACCACATTAACAACATAAAACCCTCATTCACACGAGAAAACACCCTC
  5 AGENOME-ZPMRG-Z-V1.0   99.9%  99.7%       TATTTACCACAACACAATGAGGCTCACTCACCCACCACATTAACAACATAAAACCCTCATTCACACGAGAAAACACCCTC
  6 AGENOME-ZPMRG-T-V1.0   99.9%  99.8%       TATTTACCACAACACAATGGGGCTCACTCACCCACCACATTAACAACATAAAACCCTCATTCACACGAGAAAACACCCTC
  7 AGENOME-ZPMRG-HV-V1.0  99.9%  99.8%       TATTTACCACAACACAATGGGGCTCACTCACCCACCACATTAACAACATAAAACCCTCATTCACACGAGAAAATACCCTC
  8 AGENOME-ZPMRG-A-V1.0   99.9%  99.8%       TATTTACCACAACACAATGGGGCTCACTCACCCACCACATTAACAACATAAAACCCTCATTCACACGAGAAAACACCCTC
  9 NC_012920.1            99.9%  99.8%       TATTTACCACAACACAATGGGGCTCACTCACCCACCACATTAACAACATAAAACCCTCATTCACACGAGAAAACACCCTC
    consensus/100%                            TATTTACCACAACACAATGuGGCTCACTCACCCACCACATTAACAACATAAAACCCTCATTCACACGAGAAAAsACCCTC
    consensus/90%                             TATTTACCACAACACAATGuGGCTCACTCACCCACCACATTAACAACATAAAACCCTCATTCACACGAGAAAAsACCCTC
    consensus/80%                             TATTTACCACAACACAATGGGGCTCACTCACCCACCACATTAACAACATAAAACCCTCATTCACACGAGAAAACACCCTC
    consensus/70%                             TATTTACCACAACACAATGGGGCTCACTCACCCACCACATTAACAACATAAAACCCTCATTCACACGAGAAAACACCCTC

                          cov    pid 12081          .         1         .         .         .         .         :         . 12160
  1 AGENOME-ZPMRG-U-V1.0  100.0% 100.0%       ATGTTCATACACCTATCCCCCATTCTCCTCCTATCCCTCAACCCCGACATCATTACCGGGTTTTCCTCTTGTAAATATAG
  2 AGENOME-ZPMCG-V1.0    100.0%  99.9%       ATGTTCATACACCTATCCCCCATTCTCCTCCTATCCCTCAACCCCGACATCATTACCGGGTTTTCCTCTTGTAAATATAG
  3 AGENOME-ZPMRG-M-V1.0  100.0%  99.8%       ATGTTCATACACCTATCCCCCATTCTCCTCCTATCCCTCAACCCCGACATCATTACCGGGTTTTCCTCTTGTAAATATAG
```

```
 4 AGENOME-ZPMRG-F-V1.0   99.9%  99.8%       ATGTTCATACACCTATCCCCCATTCTCCTCCTATCCCTCAACCCCGACATCATTACCGGGTTTTCCTCTTGTAAATATAG
 5 AGENOME-ZPMRG-Z-V1.0   99.9%  99.7%       ATGTTCATACACCTATCCCCCATTCTCCTCCTATCCCTCAACCCCGACATCATTACCGGGTTTTCCTCTTGTAAATATAG
 6 AGENOME-ZPMRG-T-V1.0   99.9%  99.8%       ATGTTCATACACCTATCCCCCATTCTCCTCCTATCCCTCAACCCCGACATCATTACCGGGTTTTCCTCTTGTAAATATAG
 7 AGENOME-ZPMRG-HV-V1.0  99.9%  99.8%       ATGTTCATACACCTATCCCCCATTCTCCTCCTATCCCTCAACCCCGACATCATTACCGGGTTTTCCTCTTGTAAATATAG
 8 AGENOME-ZPMRG-A-V1.0   99.9%  99.8%       ATGTTCATACACCTATCCCCCATTCTCCTCCTATCCCTCAACCCCGACATCATTACCGGGTTTTCCTCTTGTAAATATAG
 9 NC_012920.1            99.9%  99.8%       ATGTTCATACACCTATCCCCCATTCTCCTCCTATCCCTCAACCCCGACATCATTACCGGGTTTTCCTCTTGTAAATATAG
   consensus/100%                            ATGTTCATACACCTATCCCCCATTCTCCTCCTATCCCTCAACCCCGACATCATTACCGGGTTTTCCTCTTGTAAATATAG
   consensus/90%                             ATGTTCATACACCTATCCCCCATTCTCCTCCTATCCCTCAACCCCGACATCATTACCGGGTTTTCCTCTTGTAAATATAG
   consensus/80%                             ATGTTCATACACCTATCCCCCATTCTCCTCCTATCCCTCAACCCCGACATCATTACCGGGTTTTCCTCTTGTAAATATAG
   consensus/70%                             ATGTTCATACACCTATCCCCCATTCTCCTCCTATCCCTCAACCCCGACATCATTACCGGGTTTTCCTCTTGTAAATATAG

                            cov    pid 12161          .          .          .          2          .          .          .          . 12240
 1 AGENOME-ZPMRG-U-V1.0  100.0% 100.0%       TTTAACCAAAACATCAGATTGTGAATCTGACAACAGAGGCTTACGACCCCTTATTTACCGAGAAAGCTCACAAGAACTGC
 2 AGENOME-ZPMCG-V1.0    100.0%  99.9%       TTTAACCAAAACATCAGATTGTGAATCTGACAACAGAGGCTTACGACCCCTTATTTACCGAGAAAGCTCACAAGAACTGC
 3 AGENOME-ZPMRG-M-V1.0  100.0%  99.8%       TTTAACCAAAACATCAGATTGTGAATCTGACAACAGAGGCTTACGACCCCTTATTTACCGAGAAAGCTCACAAGAACTGC
 4 AGENOME-ZPMRG-F-V1.0   99.9%  99.8%       TTTAACCAAAACATCAGATTGTGAATCTGACAACAGAGGCTTACGACCCCTTATTTACCGAGAAAGCTCACAAGAACTGC
 5 AGENOME-ZPMRG-Z-V1.0   99.9%  99.7%       TTTAACCAAAACATCAGATTGTGAATCTGACAACAGAGGCTTACGACCCCTTATTTACCGAGAAAGCTCACAAGAACTGC
 6 AGENOME-ZPMRG-T-V1.0   99.9%  99.8%       TTTAACCAAAACATCAGATTGTGAATCTGACAACAGAGGCTTACGACCCCTTATTTACCGAGAAAGCTCACAAGAACTGC
 7 AGENOME-ZPMRG-HV-V1.0  99.9%  99.8%       TTTAACCAAAACATCAGATTGTGAATCTGACAACAGAGGCTTACGACCCCTTATTTACCGAGAAAGCTCACAAGAACTGC
 8 AGENOME-ZPMRG-A-V1.0   99.9%  99.8%       TTTAACCAAAACATCAGATTGTGAATCTGACAACAGAGGCTTACGACCCCTTATTTACCGAGAAAGCTCACAAGAACTGC
 9 NC_012920.1            99.9%  99.8%       TTTAACCAAAACATCAGATTGTGAATCTGACAACAGAGGCTTACGACCCCTTATTTACCGAGAAAGCTCACAAGAACTGC
   consensus/100%                            TTTAACCAAAACATCAGATTGTGAATCTGACAACAGAGGCTTACGACCCCTTATTTACCGAGAAAGCTCACAAGAACTGC
   consensus/90%                             TTTAACCAAAACATCAGATTGTGAATCTGACAACAGAGGCTTACGACCCCTTATTTACCGAGAAAGCTCACAAGAACTGC
   consensus/80%                             TTTAACCAAAACATCAGATTGTGAATCTGACAACAGAGGCTTACGACCCCTTATTTACCGAGAAAGCTCACAAGAACTGC
   consensus/70%                             TTTAACCAAAACATCAGATTGTGAATCTGACAACAGAGGCTTACGACCCCTTATTTACCGAGAAAGCTCACAAGAACTGC

                            cov    pid 12241          :          .          .          .          .          3          .          . 12320
 1 AGENOME-ZPMRG-U-V1.0  100.0% 100.0%       TAACTCATGCCCCCATGTCTAACAACATGGCTTTCTCAACTTTTAAAGGATAACAGCTATCCATTGGTCTTAGGCCCCAA
 2 AGENOME-ZPMCG-V1.0    100.0%  99.9%       TAACTCATGCCCCCATGTCTAACAACATGGCTTTCTCAACTTTTAAAGGATAACAGCTATCCATTGGTCTTAGGCCCCAA
 3 AGENOME-ZPMRG-M-V1.0  100.0%  99.8%       TAACTCATGCCCCCATGTCTAACAACATGGCTTTCTCAACTTTTAAAGGATAACAGCTATCCATTGGTCTTAGGCCCCAA
 4 AGENOME-ZPMRG-F-V1.0   99.9%  99.8%       TAACTCATGCCCCCATGTCTAACAACATGGCTTTCTCAACTTTTAAAGGATAACAGCTATCCATTGGTCTTAGGCCCCAA
 5 AGENOME-ZPMRG-Z-V1.0   99.9%  99.7%       TAACTCATGCCCCCATGTCTAACAACATGGCTTTCTCAACTTTTAAAGGATAACAGCTATCCATTGGTCTTAGGCCCCAA
 6 AGENOME-ZPMRG-T-V1.0   99.9%  99.8%       TAACTCATGCCCCCATGTCTAACAACATGGCTTTCTCAACTTTTAAAGGATAACAGCTATCCATTGGTCTTAGGCCCCAA
 7 AGENOME-ZPMRG-HV-V1.0  99.9%  99.8%       TAACTCATGCCCCCATGTCTAACAACATGGCTTTCTCAACTTTTAAAGGATAACAGCTATCCATTGGTCTTAGGCCCCAA
 8 AGENOME-ZPMRG-A-V1.0   99.9%  99.8%       TAACTCATGCCCCCATGTCTAACAACATGGCTTTCTCAACTTTTAAAGGATAACAGCTATCCATTGGTCTTAGGCCCCAA
 9 NC_012920.1            99.9%  99.8%       TAACTCATGCCCCCATGTCTAACAACATGGCTTTCTCAACTTTTAAAGGATAACAGCTATCCATTGGTCTTAGGCCCCAA
   consensus/100%                            TAACTCATGCCCCCATGTCTAACAACATGGCTTTCTCAACTTTTAAAGGATAACAGCTATCCATTGGTCTTAGGCCCCAA
   consensus/90%                             TAACTCATGCCCCCATGTCTAACAACATGGCTTTCTCAACTTTTAAAGGATAACAGCTATCCATTGGTCTTAGGCCCCAA
   consensus/80%                             TAACTCATGCCCCCATGTCTAACAACATGGCTTTCTCAACTTTTAAAGGATAACAGCTATCCATTGGTCTTAGGCCCCAA
   consensus/70%                             TAACTCATGCCCCCATGTCTAACAACATGGCTTTCTCAACTTTTAAAGGATAACAGCTATCCATTGGTCTTAGGCCCCAA

                            cov    pid 12321          .          .          :          .          .          .          .          4 12400
 1 AGENOME-ZPMRG-U-V1.0  100.0% 100.0%       GAATTTTGGTGCAACTCCAAATAAAAGTAATAACCATGCACACTACTATAACCACCCTAACCTAACTTCCCTAATTCCC
 2 AGENOME-ZPMCG-V1.0    100.0%  99.9%       AAATTTTGGTGCAACTCCAAATAAAAGTAATAACCATGCACACTACTATAACCACCCTAACCCTGACTTCCCTAATTCCC
 3 AGENOME-ZPMRG-M-V1.0  100.0%  99.8%       AAATTTTGGTGCAACTCCAAATAAAAGTAATAACCATGCACACTACTATAACCACCCTAACCCTGACTTCCCTAATTCCC
 4 AGENOME-ZPMRG-F-V1.0   99.9%  99.8%       AAATTTTGGTGCAACTCCAAATAAAAGTAATAACCATGCACACTACTATAACCACCCTAACCCTGACTTCCCTAATTCCC
 5 AGENOME-ZPMRG-Z-V1.0   99.9%  99.7%       AAATTTTGGTGCAACTCCAAATAAAAGTAATAACCATGCACACTACTATAACCACCCTAACCCTGACTTCCCTAATTCCC
 6 AGENOME-ZPMRG-T-V1.0   99.9%  99.8%       AAATTTTGGTGCAACTCCAAATAAAAGTAATAACCATGCACACTACTATAACCACCCTAACCCTGACTTCCCTAATTCCC
 7 AGENOME-ZPMRG-HV-V1.0  99.9%  99.8%       AAATTTTGGTGCAACTCCAAATAAAAGTAATAACCATGCACACTACTATAACCACCCTAACCCTGACTTCCCTAATTCCC
 8 AGENOME-ZPMRG-A-V1.0   99.9%  99.8%       AAATTTTGGTGCAACTCCAAATAAAAGTAATAACCATGCACACTACTATAACCACCCTAACCCTGACTTCCCTAATTCCC
 9 NC_012920.1            99.9%  99.8%       AAATTTTGGTGCAACTCCAAATAAAAGTAATAACCATGCACACTACTATAACCACCCTAACCCTGACTTCCCTAATTCCC
   consensus/100%                            uAATTTTGGTGCAACTCCAAATAAAAGTAATAACCATGCACACTACTATAACCACCCTAACCCTuACTTCCCTAATTCCC
   consensus/90%                             uAATTTTGGTGCAACTCCAAATAAAAGTAATAACCATGCACACTACTATAACCACCCTAACCCTuACTTCCCTAATTCCC
   consensus/80%                             AAATTTTGGTGCAACTCCAAATAAAAGTAATAACCATGCACACTACTATAACCACCCTAACCCTGACTTCCCTAATTCCC
   consensus/70%                             AAATTTTGGTGCAACTCCAAATAAAAGTAATAACCATGCACACTACTATAACCACCCTAACCCTGACTTCCCTAATTCCC

                            cov    pid 12401          .          .          .          .          :          .          .          . 12480
 1 AGENOME-ZPMRG-U-V1.0  100.0% 100.0%       CCCATCCTTACCACCCTCGTTAACCCTAACAAAAAAAACTCATACCCCCATTATGTAAAATCCATTGTCGCATCCACCTT
 2 AGENOME-ZPMCG-V1.0    100.0%  99.9%       CCCATCCTTACCACCCTCGTTAACCCTAACAAAAAAAACTCATACCCCCATTATGTAAAATCCATTGTCGCATCCACCTT
 3 AGENOME-ZPMRG-M-V1.0  100.0%  99.8%       CCCATCCTTACCACCCTCGTTAACCCTAACAAAAAAAACTCATACCCCCATTATGTAAAATCCATTGTCGCATCCACCTT
 4 AGENOME-ZPMRG-F-V1.0   99.9%  99.8%       CCCATCCTTACCACCCTCATTAACCCTAACAAAAAAAACTCATACCCCCATTATGTAAAATCCATTGTCGCATCCACCTT
 5 AGENOME-ZPMRG-Z-V1.0   99.9%  99.7%       CCCATCCTTACCACCCTCGTTAACCCTAACAAAAAAAACTCATACCCCCATTATGTAAAATCCATTGTCGCATCCACCTT
 6 AGENOME-ZPMRG-T-V1.0   99.9%  99.8%       CCCATCCTTACCACCCTCGTTAACCCTAACAAAAAAAACTCATACCCCCATTATGTAAAATCCATTGTCGCATCCACCTT
 7 AGENOME-ZPMRG-HV-V1.0  99.9%  99.8%       CCCATCCTTACCACCCTCGTTAACCCTAACAAAAAAAACTCATACCCCCATTATGTAAAATCCATTGTCGCATCCACCTT
 8 AGENOME-ZPMRG-A-V1.0   99.9%  99.8%       CCCATCCTTACCACCCTCGTTAACCCTAACAAAAAAAACTCATACCCCCATTATGTAAAATCCATTGTCGCATCCACCTT
 9 NC_012920.1            99.9%  99.8%       CCCATCCTTACCACCCTCGTTAACCCTAACAAAAAAAACTCATACCCCCATTATGTAAAATCCATTGTCGCATCCACCTT
   consensus/100%                            CCCATCCTTACCACCCTCuTTAACCCTAACAAAAAAAACTCATACCCCCATTATGTAAAATCCATTGTCGCATCCACCTT
   consensus/90%                             CCCATCCTTACCACCCTCuTTAACCCTAACAAAAAAAACTCATACCCCCATTATGTAAAATCCATTGTCGCATCCACCTT
   consensus/80%                             CCCATCCTTACCACCCTCGTTAACCCTAACAAAAAAAACTCATACCCCCATTATGTAAAATCCATTGTCGCATCCACCTT
   consensus/70%                             CCCATCCTTACCACCCTCGTTAACCCTAACAAAAAAAACTCATACCCCCATTATGTAAAATCCATTGTCGCATCCACCTT

                            cov    pid 12481          .          5          .          .          .          .          :          . 12560
 1 AGENOME-ZPMRG-U-V1.0  100.0% 100.0%       TATTATCAGTCTCTTCCCCACAACAATATTCATGTGCCTAGACCAAGAAGTTATTATCTCGAACTGACACTGAGCCACAA
 2 AGENOME-ZPMCG-V1.0    100.0%  99.9%       TATTATCAGTCTCTTCCCCACAACAATATTCATGTGCCTAGACCAAGAAGTTATTATCTCGAACTGACACTGAGCCACAA
 3 AGENOME-ZPMRG-M-V1.0  100.0%  99.8%       TATTATCAGTCTCTTCCCCACAACAATATTCATGTGCCTAGACCAAGAAGTTATTATCTCGAACTGACACTGAGCCACAA
 4 AGENOME-ZPMRG-F-V1.0   99.9%  99.8%       TATTATCAGTCTCTTCCCCACAACAATATTCATGTGCCTAGACCAAGAAGTTATTATCTCGAACTGACACTGAGCCACAA
 5 AGENOME-ZPMRG-Z-V1.0   99.9%  99.7%       TATTATCAGTCTCTTCCCCACAACAATATTCATGTGCCTAGACCAAGAAGTTATTATCTCGAACTGACACTGAGCCACAA
 6 AGENOME-ZPMRG-T-V1.0   99.9%  99.8%       TATTATCAGTCTCTTCCCCACAACAATATTCATGTGCCTAGACCAAGAAGTTATTATCTCGAACTGACACTGAGCCACAA
 7 AGENOME-ZPMRG-HV-V1.0  99.9%  99.8%       TATTATCAGTCTCTTCCCCACAACAATATTCATGTGCCTAGACCAAGAAGTTATTATCTCGAACTGACACTGAGCCACAA
 8 AGENOME-ZPMRG-A-V1.0   99.9%  99.8%       TATTATCAGTCTCTTCCCCACAACAATATTCATGTGCCTAGACCAAGAAGTTATTATCTCGAACTGACACTGAGCCACAA
 9 NC_012920.1            99.9%  99.8%       TATTATCAGTCTCTTCCCCACAACAATATTCATGTGCCTAGACCAAGAAGTTATTATCTCGAACTGACACTGAGCCACAA
   consensus/100%                            TATTATCAGTCTCTTCCCCACAACAATATTCATGTGCCTAGACCAAGAAGTTATTATCTCGAACTGACACTGAGCCACAA
   consensus/90%                             TATTATCAGTCTCTTCCCCACAACAATATTCATGTGCCTAGACCAAGAAGTTATTATCTCGAACTGACACTGAGCCACAA
   consensus/80%                             TATTATCAGTCTCTTCCCCACAACAATATTCATGTGCCTAGACCAAGAAGTTATTATCTCGAACTGACACTGAGCCACAA
   consensus/70%                             TATTATCAGTCTCTTCCCCACAACAATATTCATGTGCCTAGACCAAGAAGTTATTATCTCGAACTGACACTGAGCCACAA

                            cov    pid 12561          .          .          .          6          .          .          .          . 12640
 1 AGENOME-ZPMRG-U-V1.0  100.0% 100.0%       CCCAAACAACCCAGCTCTCCCTAAGCTTCAAACTAGACTACTTCTCCATAATATTCATCCCTGTAGCATTGTTCGTTACA
 2 AGENOME-ZPMCG-V1.0    100.0%  99.9%       CCCAAACAACCCAGCTCTCCCTAAGCTTCAAACTAGACTACTTCTCCATAATATTCATCCCTGTAGCATTGTTCGTTACA
 3 AGENOME-ZPMRG-M-V1.0  100.0%  99.8%       CCCAAACAACCCAGCTCTCCCTAAGCTTCAAACTAGACTACTTCTCCATAATATTCATCCCTGTAGCATTGTTCGTTACA
 4 AGENOME-ZPMRG-F-V1.0   99.9%  99.8%       CCCAAACAACCCAGCTCTCCCTAAGCTTCAAACTAGACTACTTCTCCATAATATTCATCCCTGTAGCATTGTTCGTTACA
```

```
 5 AGENOME-ZPMRG-Z-V1.0   99.9%  99.7%        CCCAAACAACCCAGCTCTCCCTAAGCTTCAAACTAGACTACTTCTCCATAATATTCATCCCTGTAGCATTGTTCGTTACA
 6 AGENOME-ZPMRG-T-V1.0   99.9%  99.8%        CCCAAACAACCCAGCTCTCCCTAAGCTTCAAACTAGACTACTTCTCCATAATATTCATCCCTGTAGCATTGTTCGTTACA
 7 AGENOME-ZPMRG-HV-V1.0  99.9%  99.8%        CCCAAACAACCCAGCTCTCCCTAAGCTTCAAACTAGACTACTTCTCCATAATATTCATCCCTGTAGCATTGTTCGTTACA
 8 AGENOME-ZPMRG-A-V1.0   99.9%  99.8%        CCCAAACAACCCAGCTCTCCCTAAGCTTCAAACTAGACTACTTCTCCATAATATTCATCCCTGTAGCATTGTTCGTTACA
 9 NC_012920.1            99.9%  99.8%        CCCAAACAACCCAGCTCTCCCTAAGCTTCAAACTAGACTACTTCTCCATAATATTCATCCCTGTAGCATTGTTCGTTACA
   consensus/100%                             CCCAAACAACCCAGCTCTCCCTAAGCTTCAAACTAGACTACTTCTCCATAATATTCATCCCTGTAGCATTGTTCGTTACA
   consensus/90%                              CCCAAACAACCCAGCTCTCCCTAAGCTTCAAACTAGACTACTTCTCCATAATATTCATCCCTGTAGCATTGTTCGTTACA
   consensus/80%                              CCCAAACAACCCAGCTCTCCCTAAGCTTCAAACTAGACTACTTCTCCATAATATTCATCCCTGTAGCATTGTTCGTTACA
   consensus/70%                              CCCAAACAACCCAGCTCTCCCTAAGCTTCAAACTAGACTACTTCTCCATAATATTCATCCCTGTAGCATTGTTCGTTACA

                         cov    pid 12641          :         .         .         .         .         7         .         . 12720
 1 AGENOME-ZPMRG-U-V1.0  100.0% 100.0%        TGGTCCATCATAGAATTCTCACTGTGATATATAAACTCAGACCCAAACATTAATCAGTTCTTCAAATATCTACTCATCTT
 2 AGENOME-ZPMCG-V1.0    100.0%  99.9%        TGGTCCATCATAGAATTCTCACTGTGATATATAAACTCAGACCCAAACATTAATCAGTTCTTCAAATATCTACTCATCTT
 3 AGENOME-ZPMRG-M-V1.0  100.0%  99.8%        TGGTCCATCATAGAATTCTCACTGTGATATATAAACTCAGACCCAAACATTAATCAGTTCTTCAAATATCTACTCATTTT
 4 AGENOME-ZPMRG-F-V1.0   99.9%  99.8%        TGGTCCATCATAGAATTCTCACTGTGATATATAAACTCAGACCCAAACATTAATCAGTTCTTCAAATATCTACTCATCTT
 5 AGENOME-ZPMRG-Z-V1.0   99.9%  99.7%        TGGTCCATCATAGAATTCTCACTGTGATATATAAACTCAGACCCAAACATTAATCAGTTCTTCAAATATCTACTCATTTT
 6 AGENOME-ZPMRG-T-V1.0   99.9%  99.8%        TGGTCCATCATAGAATTCTCACTGTGATATATAAACTCAGACCCAAACATTAATCAGTTCTTCAAATATCTACTCATCTT
 7 AGENOME-ZPMRG-HV-V1.0  99.9%  99.8%        TGGTCCATCATAGAATTCTCACTGTGATATATAAACTCAGACCCAAACATTAATCAGTTCTTCAAATATCTACTCATCTT
 8 AGENOME-ZPMRG-A-V1.0   99.9%  99.8%        TGGTCCATCATAGAATTCTCACTGTGATATATAAACTCAGACCCAAACATTAATCAGTTCTTCAAATATCTACTCATCTT
 9 NC_012920.1            99.9%  99.8%        TGGTCCATCATAGAATTCTCACTGTGATATATAAACTCAGACCCAAACATTAATCAGTTCTTCAAATATCTACTCATCTT
   consensus/100%                             TGGTCCATCATAGAATTCTCACTGTGATATATAAACTCAGACCCAAACATTAATCAGTTCTTCAAATATCTACTCATsTT
   consensus/90%                              TGGTCCATCATAGAATTCTCACTGTGATATATAAACTCAGACCCAAACATTAATCAGTTCTTCAAATATCTACTCATsTT
   consensus/80%                              TGGTCCATCATAGAATTCTCACTGTGATATATAAACTCAGACCCAAACATTAATCAGTTCTTCAAATATCTACTCATsTT
   consensus/70%                              TGGTCCATCATAGAATTCTCACTGTGATATATAAACTCAGACCCAAACATTAATCAGTTCTTCAAATATCTACTCATsTT

                         cov    pid 12721          .         .         :         .         .         .         .         8 12800
 1 AGENOME-ZPMRG-U-V1.0  100.0% 100.0%        CCTAATTACCATACTAATCTTAGTTACCGCTAACAACCTATTCCAACTGTTCATCGGCTGAGAGGGCGTAGGAATTATAT
 2 AGENOME-ZPMCG-V1.0    100.0%  99.9%        CCTAATTACCATACTAATCTTAGTTACCGCTAACAACCTATTCCAACTGTTCATCGGCTGAGAGGGCGTAGGAATTATAT
 3 AGENOME-ZPMRG-M-V1.0  100.0%  99.8%        CCTAATTACCATACTAATCTTAGTTACCGCTAACAACCTATTCCAACTGTTCATCGGCTGAGAGGGCGTAGGAATTATAT
 4 AGENOME-ZPMRG-F-V1.0   99.9%  99.8%        CCTAATTACCATACTAATCTTAGTTACCGCTAACAACCTATTCCAACTGTTCATCGGCTGAGAGGGCGTAGGAATTATAT
 5 AGENOME-ZPMRG-Z-V1.0   99.9%  99.7%        CCTAATTACCATACTAATCTTAGTTACCGCTAACAACCTATTCCAACTGTTCATCGGCTGAGAGGGCGTAGGAATTATAT
 6 AGENOME-ZPMRG-T-V1.0   99.9%  99.8%        CCTAATTACCATACTAATCTTAGTTACCGCTAACAACCTATTCCAACTGTTCATCGGCTGAGAGGGCGTAGGAATTATAT
 7 AGENOME-ZPMRG-HV-V1.0  99.9%  99.8%        CCTAATTACCATACTAATCTTAGTTACCGCTAACAACCTATTCCAACTGTTCATCGGCTGAGAGGGCGTAGGAATTATAT
 8 AGENOME-ZPMRG-A-V1.0   99.9%  99.8%        CCTAATTACCATACTAATCTTAGTTACCGCTAACAACCTATTCCAACTGTTCATCGGCTGAGAGGGCGTAGGAATTATAT
 9 NC_012920.1            99.9%  99.8%        CCTAATTACCATACTAATCTTAGTTACCGCTAACAACCTATTCCAACTGTTCATCGGCTGAGAGGGCGTAGGAATTATAT
   consensus/100%                             CCTAATTACCATACTAATCTTAGTTACCGCTAACAACCTATTCCAACTGTTCATCGGCTGAGAGGGCGTAGGAATTATAT
   consensus/90%                              CCTAATTACCATACTAATCTTAGTTACCGCTAACAACCTATTCCAACTGTTCATCGGCTGAGAGGGCGTAGGAATTATAT
   consensus/80%                              CCTAATTACCATACTAATCTTAGTTACCGCTAACAACCTATTCCAACTGTTCATCGGCTGAGAGGGCGTAGGAATTATAT
   consensus/70%                              CCTAATTACCATACTAATCTTAGTTACCGCTAACAACCTATTCCAACTGTTCATCGGCTGAGAGGGCGTAGGAATTATAT

                         cov    pid 12801          .         .         .         .         :         .         .         . 12880
 1 AGENOME-ZPMRG-U-V1.0  100.0% 100.0%        CCTTCTTGCTCATCAGTTGATGATACGCCCGAGCAGATGCCAACACAGCAGCCATTCAAGCAATCCTATACAACCGTATC
 2 AGENOME-ZPMCG-V1.0    100.0%  99.9%        CCTTCTTGCTCATCAGTTGATGATACGCCCGAGCAGATGCCAACACAGCAGCCATTCAAGCAATCCTATACAACCGTATC
 3 AGENOME-ZPMRG-M-V1.0  100.0%  99.8%        CCTTCTTGCTCATCAGTTGATGATACGCCCGAGCAGATGCCAACACAGCAGCCATTCAAGCAATCCTATACAACCGTATC
 4 AGENOME-ZPMRG-F-V1.0   99.9%  99.8%        CCTTCTTGCTCATCAGTTGATGATACGCCCGAGCAGATGCCAACACAGCAGCCATTCAAGCAATCCTATACAACCGTATC
 5 AGENOME-ZPMRG-Z-V1.0   99.9%  99.7%        CCTTCTTGCTCATCAGTTGATGATACGCCCGAGCAGATGCCAACACAGCAGCCATTCAAGCAATCCTATACAACCGTATC
 6 AGENOME-ZPMRG-T-V1.0   99.9%  99.8%        CCTTCTTGCTCATCAGTTGATGATACGCCCGAGCAGATGCCAACACAGCAGCCATTCAAGCAATCCTATACAACCGTATC
 7 AGENOME-ZPMRG-HV-V1.0  99.9%  99.8%        CCTTCTTGCTCATCAGTTGATGATACGCCCGAGCAGATGCCAACACAGCAGCCATTCAAGCAATCCTATACAACCGTATC
 8 AGENOME-ZPMRG-A-V1.0   99.9%  99.8%        CCTTCTTGCTCATCAGTTGATGATACGCCCGAGCAGATGCCAACACAGCAGCCATTCAAGCAATCCTATACAACCGTATC
 9 NC_012920.1            99.9%  99.8%        CCTTCTTGCTCATCAGTTGATGATACGCCCGAGCAGATGCCAACACAGCAGCCATTCAAGCAATCCTATACAACCGTATC
   consensus/100%                             CCTTCTTGCTCATCAGTTGATGATACGCCCGAGCAGATGCCAACACAGCAGCCATTCAAGCAATCCTATACAACCGTATC
   consensus/90%                              CCTTCTTGCTCATCAGTTGATGATACGCCCGAGCAGATGCCAACACAGCAGCCATTCAAGCAATCCTATACAACCGTATC
   consensus/80%                              CCTTCTTGCTCATCAGTTGATGATACGCCCGAGCAGATGCCAACACAGCAGCCATTCAAGCAATCCTATACAACCGTATC
   consensus/70%                              CCTTCTTGCTCATCAGTTGATGATACGCCCGAGCAGATGCCAACACAGCAGCCATTCAAGCAATCCTATACAACCGTATC

                         cov    pid 12881          .         9         .         .         .         .         :         . 12960
 1 AGENOME-ZPMRG-U-V1.0  100.0% 100.0%        GGCGATATCGGTTTCATCCTCGCCTTAGCATGATTTATCCTACACTCCAACTCATGAGACCCACAACAAATAGCCCTTCT
 2 AGENOME-ZPMCG-V1.0    100.0%  99.9%        GGCGATATCGGTTTCATCCTCGCCTTAGCATGATTTATCCTACACTCCAACTCATGAGACCCACAACAAATAGCCCTTCT
 3 AGENOME-ZPMRG-M-V1.0  100.0%  99.8%        GGCGATATCGGTTTCATCCTCGCCTTAGCATGATTTATCCTACACTCCAACTCATGAGACCCACAACAAATAGCCCTTCT
 4 AGENOME-ZPMRG-F-V1.0   99.9%  99.8%        GGCGATATCGGTTTTATCCTCGCCTTAGCATGATTTATCCTACACTCCAACTCATGAGACCCACAACAAATAGCCCTTCT
 5 AGENOME-ZPMRG-Z-V1.0   99.9%  99.7%        GGCGATATCGGTTTCATCCTCGCCTTAGCATGATTTATCCTACACTCCAACTCATGAGACCCACAACAAATAGCCCTTCT
 6 AGENOME-ZPMRG-T-V1.0   99.9%  99.8%        GGCGATATCGGTTTCATCCTCGCCTTAGCATGATTTATCCTACACTCCAACTCATGAGACCCACAACAAATAGCCCTTCT
 7 AGENOME-ZPMRG-HV-V1.0  99.9%  99.8%        GGCGATATCGGTTTCATCCTCGCCTTAGCATGATTTATCCTACACTCCAACTCATGAGACCCACAACAAATAGCCCTTCT
 8 AGENOME-ZPMRG-A-V1.0   99.9%  99.8%        GGCGATATCGGTTTCATCCTCGCCTTAGCATGATTTATCCTACACTCCAACTCATGAGACCCACAACAAATAGCCCTTCT
 9 NC_012920.1            99.9%  99.8%        GGCGATATCGGTTTCATCCTCGCCTTAGCATGATTTATCCTACACTCCAACTCATGAGACCCACAACAAATAGCCCTTCT
   consensus/100%                             GGCGATATCGGTTTsATCCTCGCCTTAGCATGATTTATCCTACACTCCAACTCATGAGACCCACAACAAATAGCCCTTCT
   consensus/90%                              GGCGATATCGGTTTsATCCTCGCCTTAGCATGATTTATCCTACACTCCAACTCATGAGACCCACAACAAATAGCCCTTCT
   consensus/80%                              GGCGATATCGGTTTCATCCTCGCCTTAGCATGATTTATCCTACACTCCAACTCATGAGACCCACAACAAATAGCCCTTCT
   consensus/70%                              GGCGATATCGGTTTCATCCTCGCCTTAGCATGATTTATCCTACACTCCAACTCATGAGACCCACAACAAATAGCCCTTCT

                         cov    pid 12961          .         .         .         0         .         .         .         . 13040
 1 AGENOME-ZPMRG-U-V1.0  100.0% 100.0%        AAACGCTAATCCAAGCCTCACCCCACTACTAGGCCTCCTCCTAGCAGCAGCAGGCAAATCAGCCCAATTAGGTCTCCACC
 2 AGENOME-ZPMCG-V1.0    100.0%  99.9%        AAACGCTAATCCAAGCCTCACCCCACTACTAGGCCTCCTCCTAGCAGCAGCAGGCAAATCAGCCCAATTAGGTCTCCACC
 3 AGENOME-ZPMRG-M-V1.0  100.0%  99.8%        AAACGCTAATCCAAGCCTCACCCCACTACTAGGCCTCCTCCTAGCAGCAGCAGGCAAATCAGCCCAATTAGGTCTCCACC
 4 AGENOME-ZPMRG-F-V1.0   99.9%  99.8%        AAACGCTAATCCAAGCCTCACCCCACTACTAGGCCTCCTCCTAGCAGCAGCAGGCAAATCAGCCCAATTAGGTCTCCACC
 5 AGENOME-ZPMRG-Z-V1.0   99.9%  99.7%        AAACGCTAATCCAAGCCTCACCCCACTACTAGGCCTCCTCCTAGCAGCAGCAGGCAAATCAGCCCAATTAGGTCTCCACC
 6 AGENOME-ZPMRG-T-V1.0   99.9%  99.8%        AAACGCTAATCCAAGCCTCACCCCACTACTAGGCCTCCTCCTAGCAGCAGCAGGCAAATCAGCCCAATTAGGTCTCCACC
 7 AGENOME-ZPMRG-HV-V1.0  99.9%  99.8%        AAACGCTAATCCAAGCCTCACCCCACTACTAGGCCTCCTCCTAGCAGCAGCAGGCAAATCAGCCCAATTAGGTCTCCACC
 8 AGENOME-ZPMRG-A-V1.0   99.9%  99.8%        AAACGCTAATCCAAGCCTCACCCCACTACTAGGCCTCCTCCTAGCAGCAGCAGGCAAATCAGCCCAATTAGGTCTCCACC
 9 NC_012920.1            99.9%  99.8%        AAACGCTAATCCAAGCCTCACCCCACTACTAGGCCTCCTCCTAGCAGCAGCAGGCAAATCAGCCCAATTAGGTCTCCACC
   consensus/100%                             AAACGCTAATCCAAGCCTCACCCCACTACTAGGCCTCCTCCTAGCAGCAGCAGGCAAATCAGCCCAATTAGGTCTCCACC
   consensus/90%                              AAACGCTAATCCAAGCCTCACCCCACTACTAGGCCTCCTCCTAGCAGCAGCAGGCAAATCAGCCCAATTAGGTCTCCACC
   consensus/80%                              AAACGCTAATCCAAGCCTCACCCCACTACTAGGCCTCCTCCTAGCAGCAGCAGGCAAATCAGCCCAATTAGGTCTCCACC
   consensus/70%                              AAACGCTAATCCAAGCCTCACCCCACTACTAGGCCTCCTCCTAGCAGCAGCAGGCAAATCAGCCCAATTAGGTCTCCACC

                         cov    pid 13041          :         .         .         .         .         1         .         . 13120
 1 AGENOME-ZPMRG-U-V1.0  100.0% 100.0%        CCTGACTCCCCTCAGCCATAGAAGGCCCCACCCCAGTCTCAGCCCTACTCCACTCAAGCACTATAGTTGTAGCAGGAATC
 2 AGENOME-ZPMCG-V1.0    100.0%  99.9%        CCTGACTCCCCTCAGCCATAGAAGGCCCCACCCCAGTCTCAGCCCTACTCCACTCAAGCACTATAGTTGTAGCAGGAATC
 3 AGENOME-ZPMRG-M-V1.0  100.0%  99.8%        CCTGACTCCCCTCAGCCATAGAAGGCCCCACCCCAGTCTCAGCCCTACTCCACTCAAGCACTATAGTTGTAGCAGGAATC
 4 AGENOME-ZPMRG-F-V1.0   99.9%  99.8%        CCTGACTCCCCTCAGCCATAGAAGGCCCCACCCCAGTCTCAGCCCTACTCCACTCAAGCACTATAGTTGTAGCAGGAATC
 5 AGENOME-ZPMRG-Z-V1.0   99.9%  99.7%        CCTGACTCCCCTCAGCCATAGAAGGCCCCACCCCAGTCTCAGCCCTACTCCACTCAAGCACTATAGTTGTAGCAGGAATC
```

```
 6 AGENOME-ZPMRG-T-V1.0    99.9%  99.8%       CCTGACTCCCCTCAGCCATAGAAGGCCCCACCCCAGTCTCAGCCCTACTCCACTCAAGCACTATAGTTGTAGCAGGAATC
 7 AGENOME-ZPMRG-HV-V1.0   99.9%  99.8%       CCTGACTCCCCTCAGCCATAGAAGGCCCCACCCCAGTCTCAGCCCTACTCCACTCAAGCACTATAGTTGTAGCAGGAATC
 8 AGENOME-ZPMRG-A-V1.0    99.9%  99.8%       CCTGACTCCCCTCAGCCATAGAAGGCCCCACCCCAGTCTCAGCCCTACTCCACTCAAGCACTATAGTTGTAGCAGGAATC
 9 NC_012920.1             99.9%  99.8%       CCTGACTCCCCTCAGCCATAGAAGGCCCCACCCCAGTCTCAGCCCTACTCCACTCAAGCACTATAGTTGTAGCAGGAATC
   consensus/100%                             CCTGACTCCCCTCAGCCATAGAAGGCCCCACCCCAGTCTCAGCCCTACTCCACTCAAGCACTATAGTTGTAGCAGGAATC
   consensus/90%                              CCTGACTCCCCTCAGCCATAGAAGGCCCCACCCCAGTCTCAGCCCTACTCCACTCAAGCACTATAGTTGTAGCAGGAATC
   consensus/80%                              CCTGACTCCCCTCAGCCATAGAAGGCCCCACCCCAGTCTCAGCCCTACTCCACTCAAGCACTATAGTTGTAGCAGGAATC
   consensus/70%                              CCTGACTCCCCTCAGCCATAGAAGGCCCCACCCCAGTCTCAGCCCTACTCCACTCAAGCACTATAGTTGTAGCAGGAATC

                            cov    pid 13121          .         .         :         .         .         .         .         2 13200
 1 AGENOME-ZPMRG-U-V1.0   100.0% 100.0%       TTCTTACTCATCCGCTTCCACCCCCTAGCAGAAAATAGCCCACTAATCCAAACTCTAACACTATGCTTAGGCGCTATCAC
 2 AGENOME-ZPMCG-V1.0    100.0%  99.9%       TTCTTACTCATCCGCTTCCACCCCCTAGCAGAAAATAGCCCACTAATCCAAACTCTAACACTATGCTTAGGCGCTATCAC
 3 AGENOME-ZPMRG-M-V1.0  100.0%  99.8%       TTCTTACTCATCCGCTTCCACCCCCTAGCAGAAAATAGCCCACTAATCCAAACTCTAACACTATGCTTAGGCGCTATCAC
 4 AGENOME-ZPMRG-F-V1.0    99.9%  99.8%       TTCTTACTCATCCGCTTCCACCCCCTAGCAGAAAATAGCCCACTAATCCAAACTCTAACACTATGCTTAGGCGCTATCAC
 5 AGENOME-ZPMRG-Z-V1.0    99.9%  99.7%       TTCTTACTCATCCGCTTCCACCCCCTAGCAGAAAATAGCCCACTAATCCAAACTCTAACACTATGCTTAGGCGCTATCAC
 6 AGENOME-ZPMRG-T-V1.0    99.9%  99.8%       TTCTTACTCATCCGCTTCCACCCCCTAGCAGAAAATAGCCCACTAATCCAAACTCTAACACTATGCTTAGGCGCTATCAC
 7 AGENOME-ZPMRG-HV-V1.0   99.9%  99.8%       TTCTTACTCATCCGCTTCCACCCCCTAGCAGAAAATAGCCCACTAATCCAAACTCTAACACTATGCTTAGGCGCTATCAC
 8 AGENOME-ZPMRG-A-V1.0    99.9%  99.8%       TTCTTACTCATCCGCTTCCACCCCCTAGCAGAAAATAGCCCACTAATCCAAACTCTAACACTATGCTTAGGCGCTATCAC
 9 NC_012920.1             99.9%  99.8%       TTCTTACTCATCCGCTTCCACCCCCTAGCAGAAAATAGCCCACTAATCCAAACTCTAACACTATGCTTAGGCGCTATCAC
   consensus/100%                             TTCTTACTCATCCGCTTCCACCCCCTAGCAGAAAATAGCCCACTAATCCAAACTCTAACACTATGCTTAGGCGCTATCAC
   consensus/90%                              TTCTTACTCATCCGCTTCCACCCCCTAGCAGAAAATAGCCCACTAATCCAAACTCTAACACTATGCTTAGGCGCTATCAC
   consensus/80%                              TTCTTACTCATCCGCTTCCACCCCCTAGCAGAAAATAGCCCACTAATCCAAACTCTAACACTATGCTTAGGCGCTATCAC
   consensus/70%                              TTCTTACTCATCCGCTTCCACCCCCTAGCAGAAAATAGCCCACTAATCCAAACTCTAACACTATGCTTAGGCGCTATCAC

                            cov    pid 13201          .         .         .         .         :         .         .         . 13280
 1 AGENOME-ZPMRG-U-V1.0   100.0% 100.0%       CACTCTGTTCGCAGCAGTCTGCGCCCTTACACAAAATGACATCAAAAAAATCGTAGCCTTCTCCACTTCAAGTCAACTAG
 2 AGENOME-ZPMCG-V1.0    100.0%  99.9%       CACTCTGTTCGCAGCAGTCTGCGCCCTTACACAAAATGACATCAAAAAAATCGTAGCCTTCTCCACTTCAAGTCAACTAG
 3 AGENOME-ZPMRG-M-V1.0  100.0%  99.8%       CACTCTGTTCGCAGCAGTCTGCGCCCTTACACAAAATGACATCAAAAAAATCGTAGCCTTCTCCACTTCAAGTCAACTAG
 4 AGENOME-ZPMRG-F-V1.0    99.9%  99.8%       CACTCTGTTCGCAGCAGTCTGCGCCCTTACACAAAATGACATCAAAAAAATCGTAGCCTTCTCCACTTCAAGTCAACTAG
 5 AGENOME-ZPMRG-Z-V1.0    99.9%  99.7%       CACTCTGTTCGCAGCAGTCTGCGCCCTTACACAAAATGACATCAAAAAAATCGTAGCCTTCTCCACTTCAAGTCAACTAG
 6 AGENOME-ZPMRG-T-V1.0    99.9%  99.8%       CACTCTGTTCGCAGCAGTCTGCGCCCTTACACAAAATGACATCAAAAAAATCGTAGCCTTCTCCACTTCAAGTCAACTAG
 7 AGENOME-ZPMRG-HV-V1.0   99.9%  99.8%       CACTCTGTTCGCAGCAGTCTGCGCCCTTACACAAAATGACATCAAAAAAATCGTAGCCTTCTCCACTTCAAGTCAACTAG
 8 AGENOME-ZPMRG-A-V1.0    99.9%  99.8%       CACTCTGTTCGCAGCAGTCTGCGCCCTTACACAAAATGACATCAAAAAAATCGTAGCCTTCTCCACTTCAAGTCAACTAG
 9 NC_012920.1             99.9%  99.8%       CACTCTGTTCGCAGCAGTCTGCGCCCTTACACAAAATGACATCAAAAAAATCGTAGCCTTCTCCACTTCAAGTCAACTAG
   consensus/100%                             CACTCTGTTCGCAGCAGTCTGCGCCCTTACACAAAATGACATCAAAAAAATCGTAGCCTTCTCCACTTCAAGTCAACTAG
   consensus/90%                              CACTCTGTTCGCAGCAGTCTGCGCCCTTACACAAAATGACATCAAAAAAATCGTAGCCTTCTCCACTTCAAGTCAACTAG
   consensus/80%                              CACTCTGTTCGCAGCAGTCTGCGCCCTTACACAAAATGACATCAAAAAAATCGTAGCCTTCTCCACTTCAAGTCAACTAG
   consensus/70%                              CACTCTGTTCGCAGCAGTCTGCGCCCTTACACAAAATGACATCAAAAAAATCGTAGCCTTCTCCACTTCAAGTCAACTAG

                            cov    pid 13281          .         3         .         .         .         .         :         . 13360
 1 AGENOME-ZPMRG-U-V1.0   100.0% 100.0%       GACTCATAATAGTTACAATCGGCATCAACCAACCACACCTAGCATTCCTGCACATCTGTACCCACGCCTTCTTCAAAGCC
 2 AGENOME-ZPMCG-V1.0    100.0%  99.9%       GACTCATAATAGTTACAATCGGCATCAACCAACCACACCTAGCATTCCTGCACATCTGTACCCACGCCTTCTTCAAAGCC
 3 AGENOME-ZPMRG-M-V1.0  100.0%  99.8%       GACTCATAATAGTTACAATCGGCATCAACCAACCACACCTAGCATTCCTGCACATCTGTACCCACGCCTTCTTCAAAGCC
 4 AGENOME-ZPMRG-F-V1.0    99.9%  99.8%       GACTCATAATAGTTACAATCGGCATCAACCAACCACACCTAGCATTCCTGCACATCTGTACCCACGCCTTCTTCAAAGCC
 5 AGENOME-ZPMRG-Z-V1.0    99.9%  99.7%       GACTCATAATAGTTACAATCGGCATCAACCAACCACACCTAGCATTCCTGCACATCTGTACCCACGCCTTCTTCAAAGCC
 6 AGENOME-ZPMRG-T-V1.0    99.9%  99.8%       GACTCATAATAGTTACAATCGGCATCAACCAACCACACCTAGCATTCCTGCACATCTGTACCCACGCCTTCTTCAAAGCC
 7 AGENOME-ZPMRG-HV-V1.0   99.9%  99.8%       GACTCATAATAGTTACAATCGGCATCAACCAACCACACCTAGCATTCCTGCACATCTGTACCCACGCCTTCTTCAAAGCC
 8 AGENOME-ZPMRG-A-V1.0    99.9%  99.8%       GACTCATAATAGTTACAATCGGCATCAACCAACCACACCTAGCATTCCTGCACATCTGTACCCACGCCTTCTTCAAAGCC
 9 NC_012920.1             99.9%  99.8%       GACTCATAATAGTTACAATCGGCATCAACCAACCACACCTAGCATTCCTGCACATCTGTACCCACGCCTTCTTCAAAGCC
   consensus/100%                             GACTCATAATAGTTACAATCGGCATCAACCAACCACACCTAGCATTCCTGCACATCTGTACCCACGCCTTCTTCAAAGCC
   consensus/90%                              GACTCATAATAGTTACAATCGGCATCAACCAACCACACCTAGCATTCCTGCACATCTGTACCCACGCCTTCTTCAAAGCC
   consensus/80%                              GACTCATAATAGTTACAATCGGCATCAACCAACCACACCTAGCATTCCTGCACATCTGTACCCACGCCTTCTTCAAAGCC
   consensus/70%                              GACTCATAATAGTTACAATCGGCATCAACCAACCACACCTAGCATTCCTGCACATCTGTACCCACGCCTTCTTCAAAGCC

                            cov    pid 13361          .         .         .         4         .         .         .         . 13440
 1 AGENOME-ZPMRG-U-V1.0   100.0% 100.0%       ATACTATTTATGTGCTCCGGGTCCATCATCCACAACCTTAACAATGAACAAGATATTCGAAAAATAGGAGGACTACTCAA
 2 AGENOME-ZPMCG-V1.0    100.0%  99.9%       ATACTATTTATGTGCTCCGGGTCCATCATCCACAACCTTAACAATGAACAAGATATTCGAAAAATAGGAGGACTACTCAA
 3 AGENOME-ZPMRG-M-V1.0  100.0%  99.8%       ATACTATTTATGTGCTCCGGGTCCATCATCCACAACCTTAACAATGAACAAGATATTCGAAAAATAGGAGGACTACTCAA
 4 AGENOME-ZPMRG-F-V1.0    99.9%  99.8%       ATACTATTTATGTGCTCCGGGTCCATCATCCACAACCTTAACAATGAACAAGATATTCGAAAAATAGGAGGACTACTCAA
 5 AGENOME-ZPMRG-Z-V1.0    99.9%  99.7%       ATACTATTTATGTGCTCCGGGTCCATCATCCACAACCTTAACAATGAACAAGATATTCGAAAAATAGGAGGACTACTCAA
 6 AGENOME-ZPMRG-T-V1.0    99.9%  99.8%       ATACTATTTATGTGCTCCGGATCCATCATCCACAACCTTAACAATGAACAAGATATTCGAAAAATAGGAGGACTACTCAA
 7 AGENOME-ZPMRG-HV-V1.0   99.9%  99.8%       ATACTATTTATGTGCTCCGGGTCCATCATCCACAACCTTAACAATGAACAAGATATTCGAAAAATAGGAGGACTACTCAA
 8 AGENOME-ZPMRG-A-V1.0    99.9%  99.8%       ATACTATTTATGTGCTCCGGGTCCATCATCCACAACCTTAACAATGAACAAGATATTCGAAAAATAGGAGGACTACTCAA
 9 NC_012920.1             99.9%  99.8%       ATACTATTTATGTGCTCCGGGTCCATCATCCACAACCTTAACAATGAACAAGATATTCGAAAAATAGGAGGACTACTCAA
   consensus/100%                             ATACTATTTATGTGCTCCGGuTCCATCATCCACAACCTTAACAATGAACAAGATATTCGAAAAATAGGAGGACTACTCAA
   consensus/90%                              ATACTATTTATGTGCTCCGGuTCCATCATCCACAACCTTAACAATGAACAAGATATTCGAAAAATAGGAGGACTACTCAA
   consensus/80%                              ATACTATTTATGTGCTCCGGGTCCATCATCCACAACCTTAACAATGAACAAGATATTCGAAAAATAGGAGGACTACTCAA
   consensus/70%                              ATACTATTTATGTGCTCCGGGTCCATCATCCACAACCTTAACAATGAACAAGATATTCGAAAAATAGGAGGACTACTCAA

                            cov    pid 13441          :         .         .         .         .         5         .         . 13520
 1 AGENOME-ZPMRG-U-V1.0   100.0% 100.0%       AACCATACCTCTCACTTCAACCTCCCTCACCATTGGCAGCCTAGCATTAGCAGGAATACCTTTCCTCACAGGTTTCTACT
 2 AGENOME-ZPMCG-V1.0    100.0%  99.9%       AACCATACCTCTCACTTCAACCTCCCTCACCATTGGCAGCCTAGCATTAGCAGGAATACCTTTCCTCACAGGTTTCTACT
 3 AGENOME-ZPMRG-M-V1.0  100.0%  99.8%       AACCATACCTCTCACTTCAACCTCCCTCACCATTGGCAGCCTAGCATTAGCAGGAATACCTTTCCTCACAGGTTTCTACT
 4 AGENOME-ZPMRG-F-V1.0    99.9%  99.8%       AACCATACCTCTCACTTCAACCTCCCTCACCATTGGCAGCCTAGCATTAGCAGGAATACCTTTCCTCACAGGTTTCTACT
 5 AGENOME-ZPMRG-Z-V1.0    99.9%  99.7%       AACCATACCTCTCACTTCAACCTCCCTCACCATTGGCAGCCTAGCATTAGCAGGAATACCTTTCCTCACAGGTTTCTACT
 6 AGENOME-ZPMRG-T-V1.0    99.9%  99.8%       AACCATACCTCTCACTTCAACCTCCCTCACCATTGGCAGCCTAGCATTAGCAGGAATACCTTTCCTCACAGGTTTCTACT
 7 AGENOME-ZPMRG-HV-V1.0   99.9%  99.8%       AACCATACCTCTCACTTCAACCTCCCTCACCATTGGCAGCCTAGCATTAGCAGGAATACCTTTCCTCACAGGTTTCTACT
 8 AGENOME-ZPMRG-A-V1.0    99.9%  99.8%       AACCATACCTCTCACTTCAACCTCCCTCACCATTGGCAGCCTAGCATTAGCAGGAATACCTTTCCTCACAGGTTTCTACT
 9 NC_012920.1             99.9%  99.8%       AACCATACCTCTCACTTCAACCTCCCTCACCATTGGCAGCCTAGCATTAGCAGGAATACCTTTCCTCACAGGTTTCTACT
   consensus/100%                             AACCATACCTCTCACTTCAACCTCCCTCACCATTGGCAGCCTAGCATTAGCAGGAATACCTTTCCTCACAGGTTTCTACT
   consensus/90%                              AACCATACCTCTCACTTCAACCTCCCTCACCATTGGCAGCCTAGCATTAGCAGGAATACCTTTCCTCACAGGTTTCTACT
   consensus/80%                              AACCATACCTCTCACTTCAACCTCCCTCACCATTGGCAGCCTAGCATTAGCAGGAATACCTTTCC
```
