## Appendix 6 for "The First Complete Zoroastrian-Parsi Mitochondrial Reference Genome and genetic signatures of an endogamous non-smoking population"

| PMID | GENE | MUTATION | TISSUE | DISEASE |
| --- | --- | --- | --- | --- |
| 16892079 | NA | m.16183C>A | cancer cell lines | Lung cancer |
| 16892079 | NA | m.16187C>T | cancer cell lines | Lung cancer |
| 16892079 | 16S ribosomal RNA | m.2664T>C | cancer cell lines | Lung cancer |
| 16892079 | ND5 | m.12345G>A | cancer cell lines | Lung cancer |
| 16892079 | tRNAtryptophan | m.5521G>A | cancer cell lines | Lung cancer |
| 29138417 | NA | m.3398T>C | NA | non-small cell lung cancer; colon cancer |
| 29138417 | NA | m.C3497T | NA | non-small cell lung cancer; colon cancer |
| 29138417 | NA | m.T12338C | NA | non-small cell lung cancer; colon cancer |
| 29138417 | NA | m.T3394C | NA | non-small cell lung cancer; colon cancer |
| 29138417 | ND6 | m.13885insC | NA | lung carcinoma metastasis |

Appendix : 6

**Reported Lung Cancer Associated Variants absent in the Zoroastrian-Parsi Population**
